# Learning the Relationship Between Variants, Metabolic Fluxes and Phenotypes

**DOI:** 10.1101/2024.03.04.577140

**Authors:** Dongin Kim, Seokjin Han, Seong-Hyeuk Nam, Tae Yong Kim

**Author notes:** These authors contributed equally to this work (Dongin Kim, Seokjin Han).

## Abstract

In the dynamic field of cancer research, the fusion of genetics and machine learning presents a groundbreaking opportunity to understand the intricate links between genomic variations and cancer phenotypes. Despite the wealth of genetic data, translating it into actionable insights remains a challenge, particularly in the context of complex cellular metabolism. VaMP (**<u>Va</u>**riants, **<u>M</u>**etabolic Fluxes & **<u>P</u>**henotypes), our novel approach, addresses this by integrating neural networks and metabolic models, offering a comprehensive framework for deciphering cancer biology. The developed method leverages a novel end-to-end neural network architecture, integrating genome-scale metabolic models (GSMs) to capture the intricate relationship between genetic variations and cellular phenotypes. VaMP’s encoder maps genetic variants to metabolic fluxes through a series of carefully designed steps utilizing neural network and GSM, while the decoder predicts phenotype probabilities using the differences between input and reference fluxes. The training data comprise mutation information and phenotypes, eliminating the need for explicit metabolic flux data preparation. Validation experiments on five cancer types demonstrate VaMP’s ability to identify significant genes and metabolic signatures. Further utilizing SCREENER and co-occurrence analyses, the assessment reveals VaMP’s capacity to anticipate established gene-disease relationships. The identified metabolic signatures are robustly substantiated by diverse literature grounded in experimental studies. Furthermore, an in-depth exploration of the outcomes from five VaMP models involved the correlation analysis for variants and fluxes to establish connections between significant genes and cancer-causing variants. Overall, VaMP can be used as a promising tool for unraveling the complex interplay between genetic alterations and cancer phenotypes, with implications for understanding disease mechanisms and identifying novel therapeutic targets.

## 1 Introduction

In the current landscape of cancer research, the convergence of genetic studies and machine learning presents an unprecedented opportunity to decode the intricate relationship between genomic variations and cancer phenotypes [1–3]. The genomic era has witnessed an exponential increase in the availability of genetic data, offering a comprehensive catalog of mutations associated with various cancers [4–6]. However, translating this wealth of information into actionable insights remains a formidable challenge.

Traditional methodologies for interpreting genetic variants often fall short in capturing the nuanced interplay of alterations within cellular systems, particularly in the context of metabolism [7, 8]. The dynamic and context-dependent nature of metabolic networks adds an additional layer of complexity to deciphering the consequences of genetic mutations. As machine learning continues to demonstrate its prowess in unraveling complex patterns in biological data [9, 10], there is a compelling need for specialized tools that bridge the gap between genetic variations and metabolic phenotypes.

As a response to this pressing need, we developed VaMP, leveraging cutting-edge neural network architectures and genome-scale metabolic models (GSMs) [11]. By providing a holistic framework that integrates genetic information with metabolic fluxes, VaMP aims to enhance our understanding of cancer biology. The urgency of this endeavor is underscored by the potential impact on precision medicine, where a deeper comprehension of the molecular underpinnings of cancer can pave the way for personalized therapeutic strategies.

Moreover, in the broader context of machine learning research, VaMP contributes to the evolving landscape of interpretable models in genomics. As black-box algorithms pose challenges in elucidating the rationale behind predictions [10], VaMP offers a transparent and interpretable approach, shedding light on the functional consequences of genetic variants within the realm of cellular metabolism.

## 2 VaMP: <u>Va</u>riants, <u>M</u>etabolic Fluxes & <u>P</u>henotypes

VaMP is an end-to-end neural network for predicting the phenotype from given patient’s variants, where the latent space represents the metabolic flux calculated from the GSM (see Figure 1).

**Figure 1:**
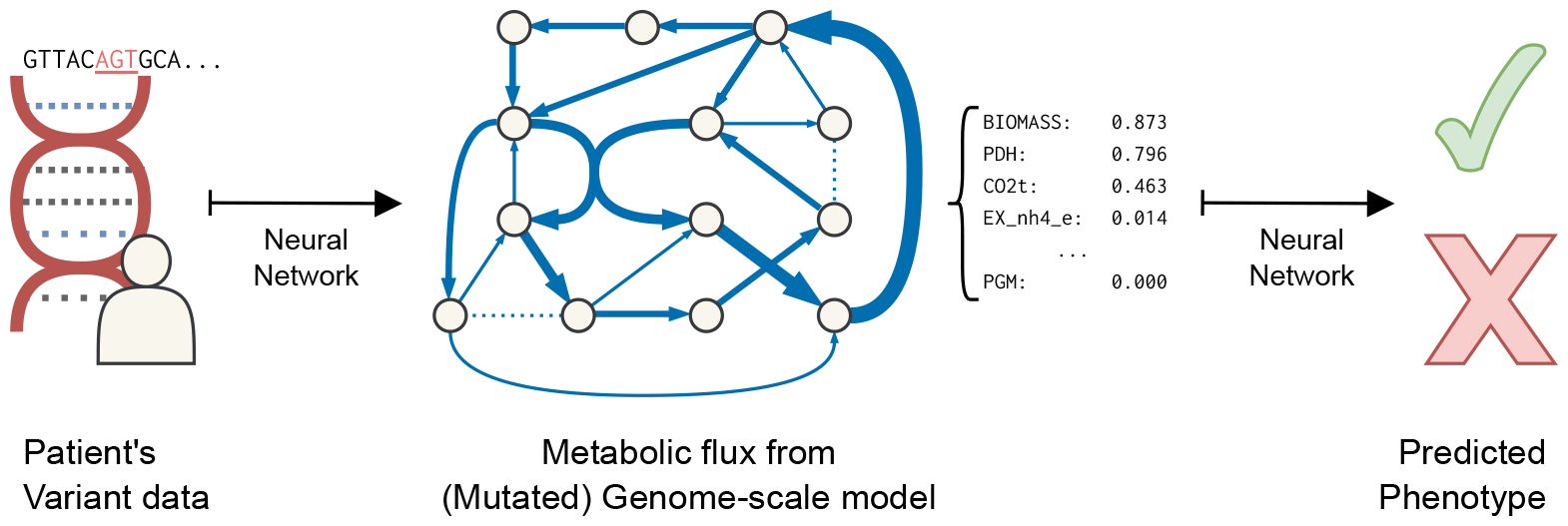
Overview of VaMP

Note that the training data for VaMP is the mutation information and the phenotypes, so there is no need to prepare training data for the metabolic flux.

### 2.1 Encoder

VaMP maps variants to the flux via the following steps (see Figure 2):

i. For each genes, compute gene damage from the variants.
ii. Derive reaction damages from gene damages, where the relationship is encoded as the gene product association format in the GSM.
iii. Solve the minimization of metabolic adjustment (MOMA) [12] to get the flux, where the reference flux *v*_0_ is the solution of the original GSM and the flux bound *l* and *u* are changed according to the reaction damage *r*.

**Figure 2:**
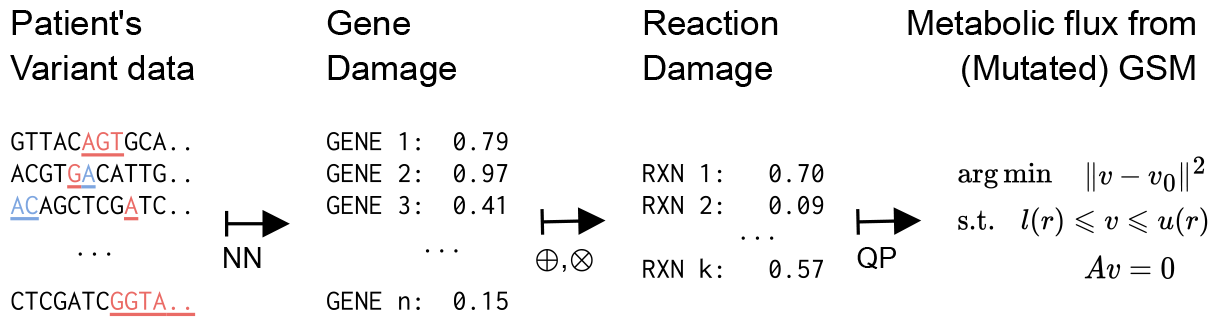
Encoder of VaMP

#### 2.1.1 Input Data

VaMP requires the number of each variant consequence (defined in [13]) of every genes as an input. In other words, 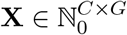 where *C* is the number of consequence types and *G* is the number of genes in the GSM.

#### 2.1.2 Gene Damage

From the variant consequence counts, VaMP computes a damage for each gene: Δ = *β*^*T*^ log(X + 1) *∈ ℝ*^*G*^, where *β* is a *C*-dimensional parameter.

Notice that each element of *β* should be positive since we assume that every variants may cause functional damage. Furthermore, because we assume that different types of consequences will have different severities, further constraints are added: *β*_*k*_ *≥ β*_*k*+1_ for all *k ∈{* 1, …, *C −* 1*}*.

To establish this constraint, we derive *β* from *β*^(0)^ *∈ ℝ*^*C*^ as follows:

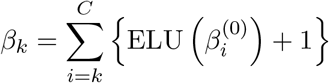

where ELU is defined as

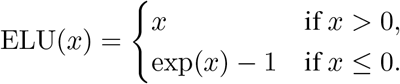

#### 2.1.3 Reaction Damage

Reaction damages are derived from the gene damages, combined with the gene product association encoded in the GSM. Gene product association refers to the mapping of genes to their associated metabolic reactions, where the association is written as a boolean expression using `or` and `and`, implying one of the unit are required to catalyze the reaction (*e*.*g*., they encode isozymes) and all units are required to catalzye the reaction (*e*.*g*., they are subunits of enzymatic complex).

Usually, the logical expression in GSM is represented as a positive disjunctive normal form (DNF). In other words, the logical expression can be formulated as a nested list of genes, where the inner-level genes are connected by and and the outer-level conjunctions are connected by or. For instance, `[[A, B]`, `[A, C, E]]` is interpreted to `(A and B) or (A and C and E)`.

In terms of damage, `or` and `and` in the association can be interpreted as min and max operations, respectively. If every genes involved in a reaction are in an `or` expression, then even if some genes are heavily damaged, another gene can replace it. Therefore, the actual damage to the reaction can be modeled via min function. Similarly, if every genes involved in a reaction are in an `and` expression, then a large damage to one gene will affect the reaction by that much, even if all the other genes are normal. Therefore, the actual damage to the reaction can be modeled via max function.

Formally, let *δ ∈ ℝ*^*R*^ be the reaction damage, where *R* is the number of reactions in the GSM. Define *α* : *{* 1, …, *R} → 𝒫* (*𝒫* (*{* 1, …, *G}*)) be the gene product association map for each relation. From the DNF *α*(*r*) = *{ A*_1_, …, *A*_*T*_ *}, δ*_*r*_ can be computed as follows:

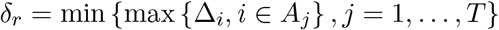

However, this expression is difficult to parallelize on a GPU, so VaMP uses softmax to efficiently approximate the min/max expressions.

#### 2.1.4 Metabolic Flux

After computing the reaction damage *δ*, VaMP solves the minimization of metabolic adjustment (MOMA) [12] to get the flux *v*^*∗*^ of damaged GSM:

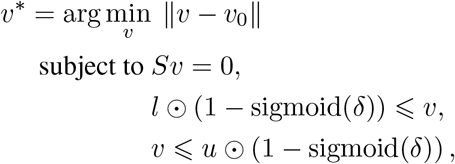

where *S* is the stoichiometry matrix of GSM, *v*_0_ is the reference flux, and the *l* and *u* are the flux bounds. Note that VaMP uses the solution of parsimonious flux balance analysis (pFBA) [14] from the original GSM as the reference flux.

VaMP currently uses OSQP [15] to solve every optimization problems. Note that this is not compatible to the usual automatic differentiation frameworks (*e*.*g*., TensorFlow [16], PyTorch [17], …) so we have to implement the backpropagation algorithm manually. Based on OptNet [18], we’ve developed a package to support the backpropagation of the specific form of the quadratic programming (see Appendix A for details).

### 2.2 Decoder

To predict the phenotype probability, VaMP first calculates the difference between the input flux and the reference flux. Then, the differences are fed into a neural network to predict the final probability (see Figure 3).

**Figure 3:**
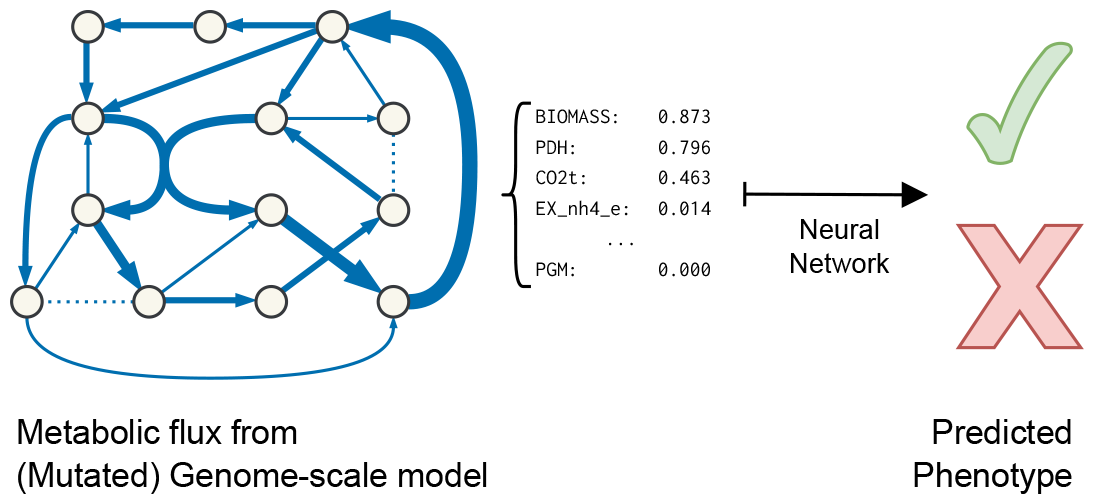
Decoder of VaMP

VaMP first calculates the difference between the flux *v*^*∗*^ and the reference flux *v*_0_. Since *Sv*^*∗*^ = *Sv*_0_ = 0, we can reduce the dimension by using bases of kernel space of *S*. Finally, the reduced difference is fed into a neural network to predict the final probability.

## 3 Experiment Setup

### 3.1 Preprocessing and Training

We used a human GSM from Metabolic Atlas [19], and genetic variants data from Genomics England (GE) [20] and gnomAD [21]. We chose five cancer types (breast, pancreatic, lung, renal, and colorectal cancer) from GE and trained model for each cancer type. We removed variants that not passed quality control filter and not mapped to genes in GSM.

We used Adam optimizer [22] to train VaMP, accompanied with the gradient clipping [23] and OneCycleLR learning rate scheduler [24]. Furthermore, we applied *𝓁*_1_ penalty on the encoder and the decoder respectively.

### 3.2 Post-hoc Analysis

We followed post-hoc analysis method suggested by [25]. First, we tried to find significant reactions based on reaction fluxes. We used *t*-test to compute *p*-values between the normal group and the cancer group, and adjusted using Bonferroni correction. Furthermore, we calculated log_2_ FC and choose significant reactions based on adjusted *p*-values and |log_2_ FC|.

### 3.3 Pathway enrichment Analysis

After the significant reactions are obtained, we collected all genes related to that reaction. Using these genes, we did pathway enrichment analysis via hypergeometric tests and Benjamini-Hochberg procedure. We identified significantly enriched pathways having adjusted *p*-value less than 0.01.

### 3.4 SCREENER Analysis

We used a web service platform named SCREENER1 [26], which extracts the gene-disease relations from the biomedical literature in real-time to identify the pairs of a significant gene and a cancer type having the known relationship based on literature evidence.

### 3.5 Co-occurrence Analysis

We used Biopython [27], freely available Python tools for computational molecular biology and bioinformatics to count the number of literature in PubMed mentioning each pair of a significant gene and a cancer type simultaneously.

### 3.6 LOR-correlation Analysis

We followed LOR-correlation analysis method suggested by [25]. We conducted statistical tests to assess the cancer-causing effect of variants based on the flux difference of the significant reactions between cancer and normal GSMs. It was examined by fitting a linear regression line between the log odds ratio (LOR) of variants and fluxes. The LOR for each variant concerning a specific cancer type was calculated as LOR = log_10_ ((PC*/*PN) */* (AC*/*AN)), where PR, PS, AR, and AS represent the counts of the individual with the variant and the type of cancer (PC), have the variants and are normal (PN), do not have the variant and the type of cancer (AC), and do not have the variant and are normal (AN), respectively. If any of these counts were 0, 0.5 was added to each to ensure non-zero values when computing the logarithm. The fluxes associated with each variant were defined as the set of fluxes in individuals containing that variants. We used *t*-test to compute *p*-values between each pair of variants in a significant reaction.

## 4 Result

### 4.1 Identification of significant genes for five cancer types

We trained VaMP models of five cancer types to predict the cancer phenotype based on the metabolic flux adjusted by the information of variant consequences of metabolic genes. Then, we performed post-hoc analysis on each VaMP model of five cancer types to identify significant reactions of which fluxes are remarkably discriminating between cancer and normal phenotype, that is the cancer-causing reactions. During this process, we calculated log_2_ FC and adjusted *p*-values for every reaction in GSM and selected significant reactions using the threshold value of 0.5 for |log_2_ FC| and 0.01 for adjusted *p*-values. Additionally, the reactions with the mean flux values below 0.1 in both cancer and normal GSMs are excluded to avoid the influence by insignificant flux values. Furthermore, we extracted all genes which have variant information and are also related to the significant reactions in GSM simultaneously and these genes are defined as significant genes causing the cancer phenotype. Using these genes, we did pathway enrichment analysis using KEGG pathways to explore overall metabolic pathways where significant genes are located (see Figure 4–Figure 8). As a result, the pathways in purine metabolism, carbohydrate digestion and absorption, starch and sucrose metabolism, adipocytokine signaling and thermogenesis are significantly enriched in five types of cancers.

**Figure 4:**
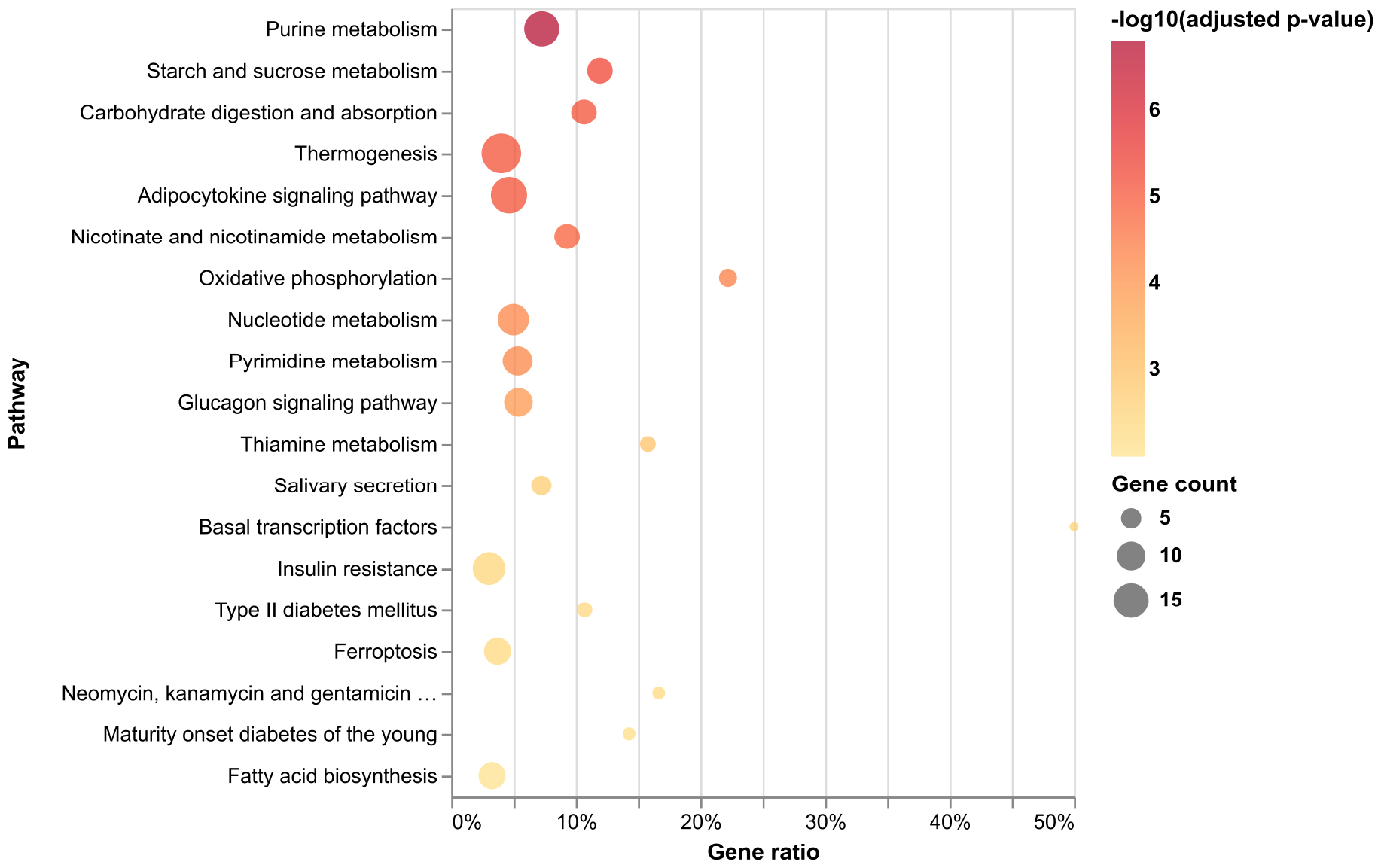
Pathway enrichment analysis of significant genes in breast cancer

**Figure 5:**
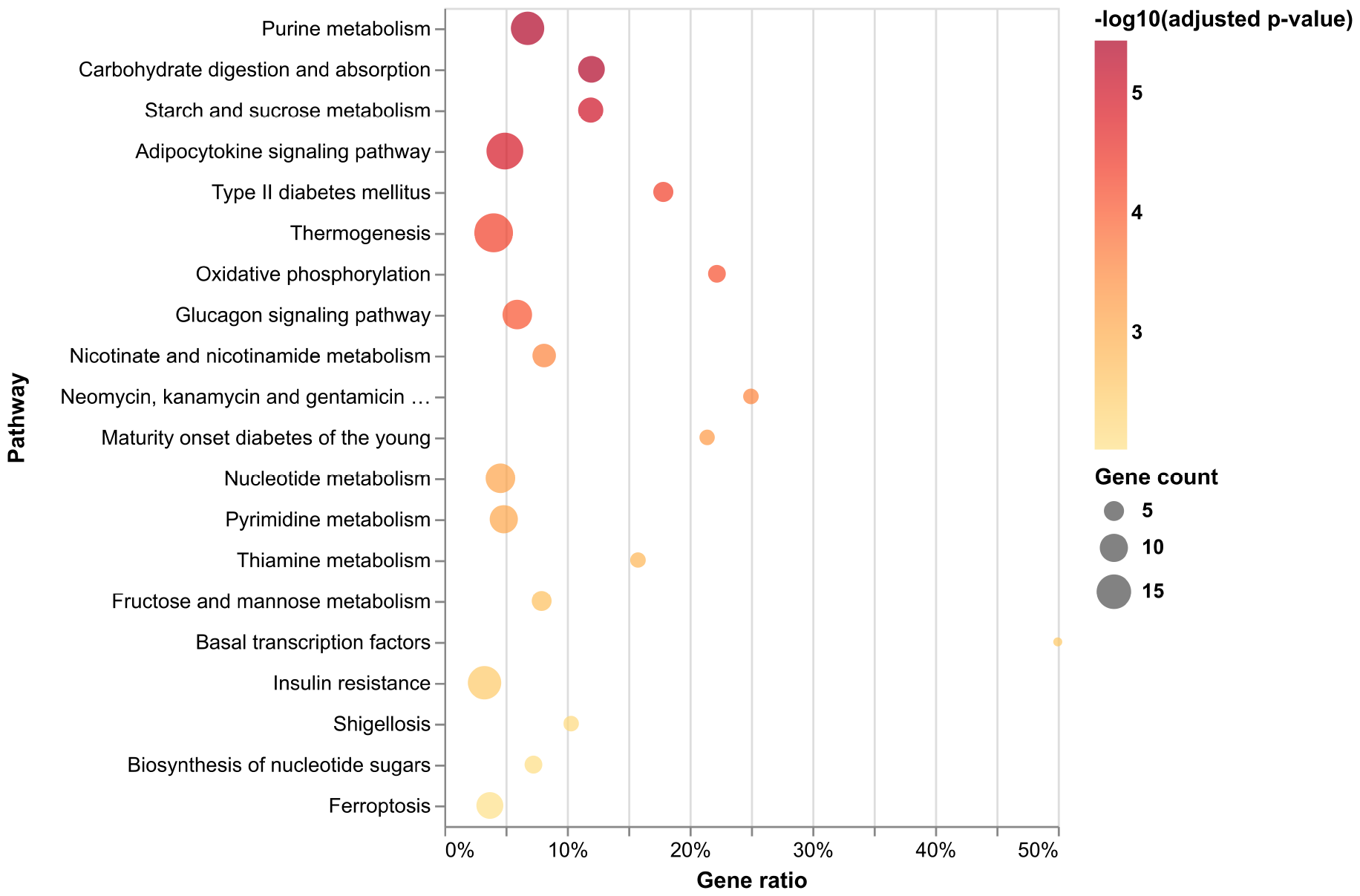
Pathway enrichment analysis of significant genes in colorectal cancer

**Figure 6:**
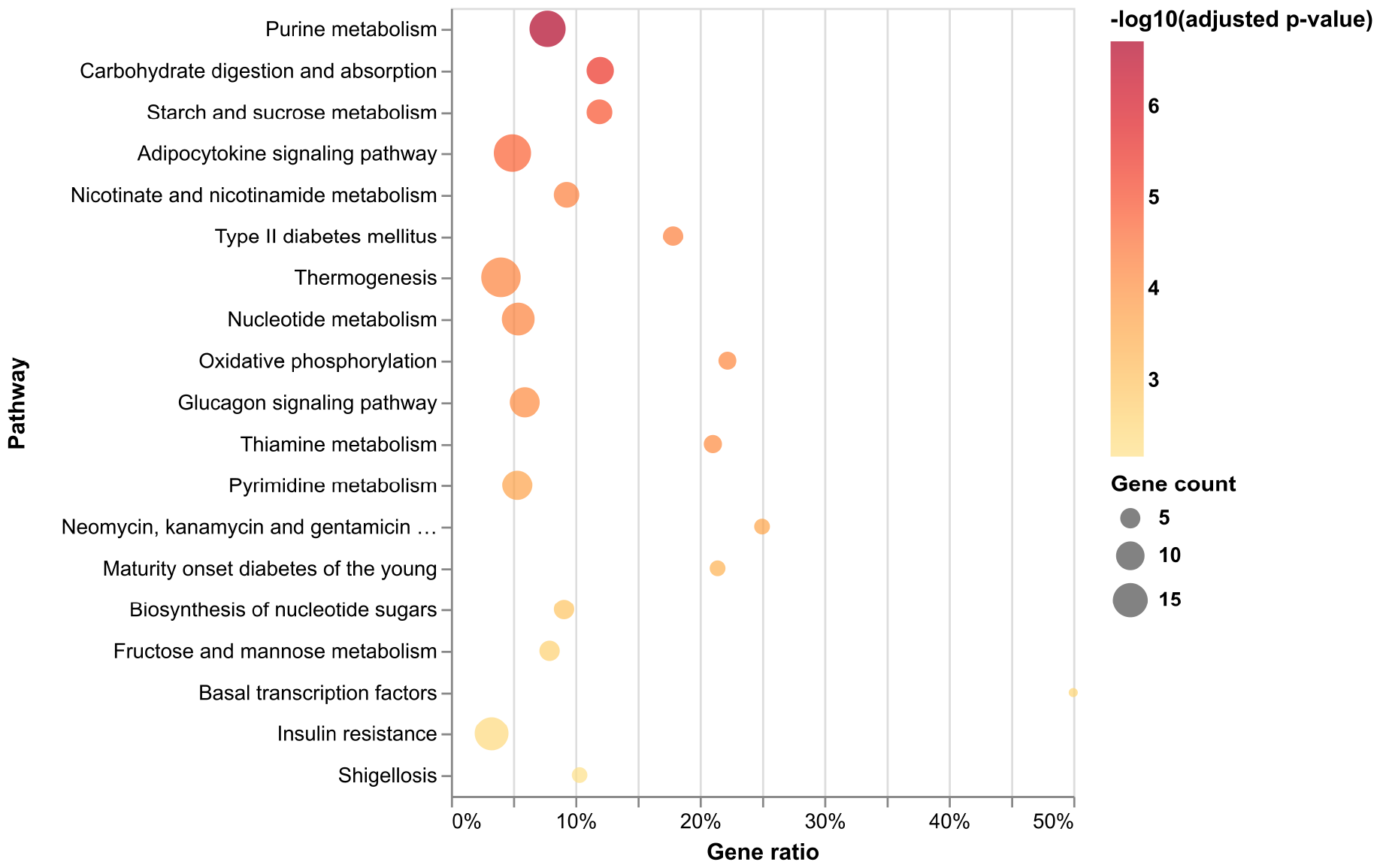
Pathway enrichment analysis of significant genes in lung cancer

**Figure 7:**
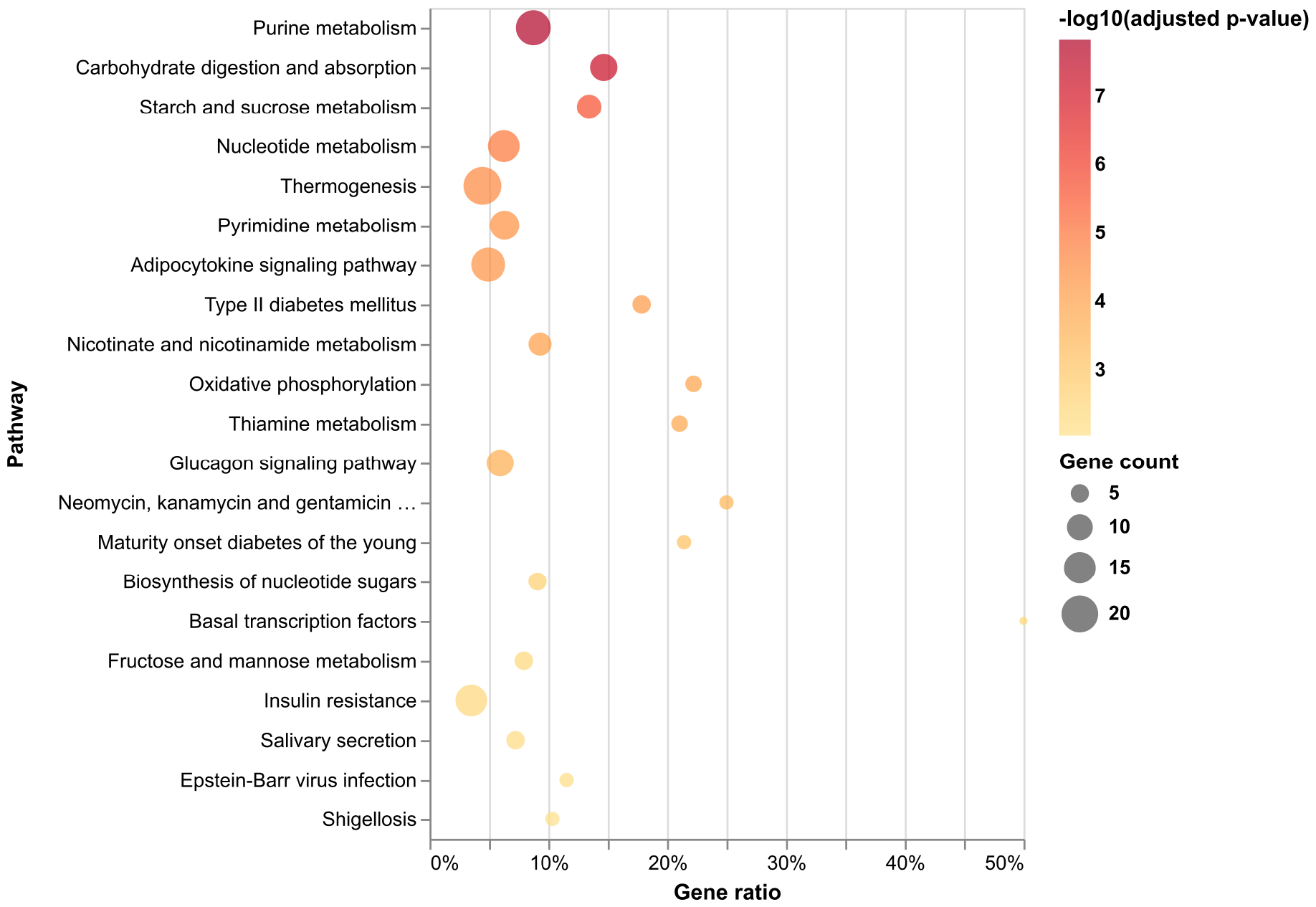
Pathway enrichment analysis of significant genes in pancreatic cancer

**Figure 8:**
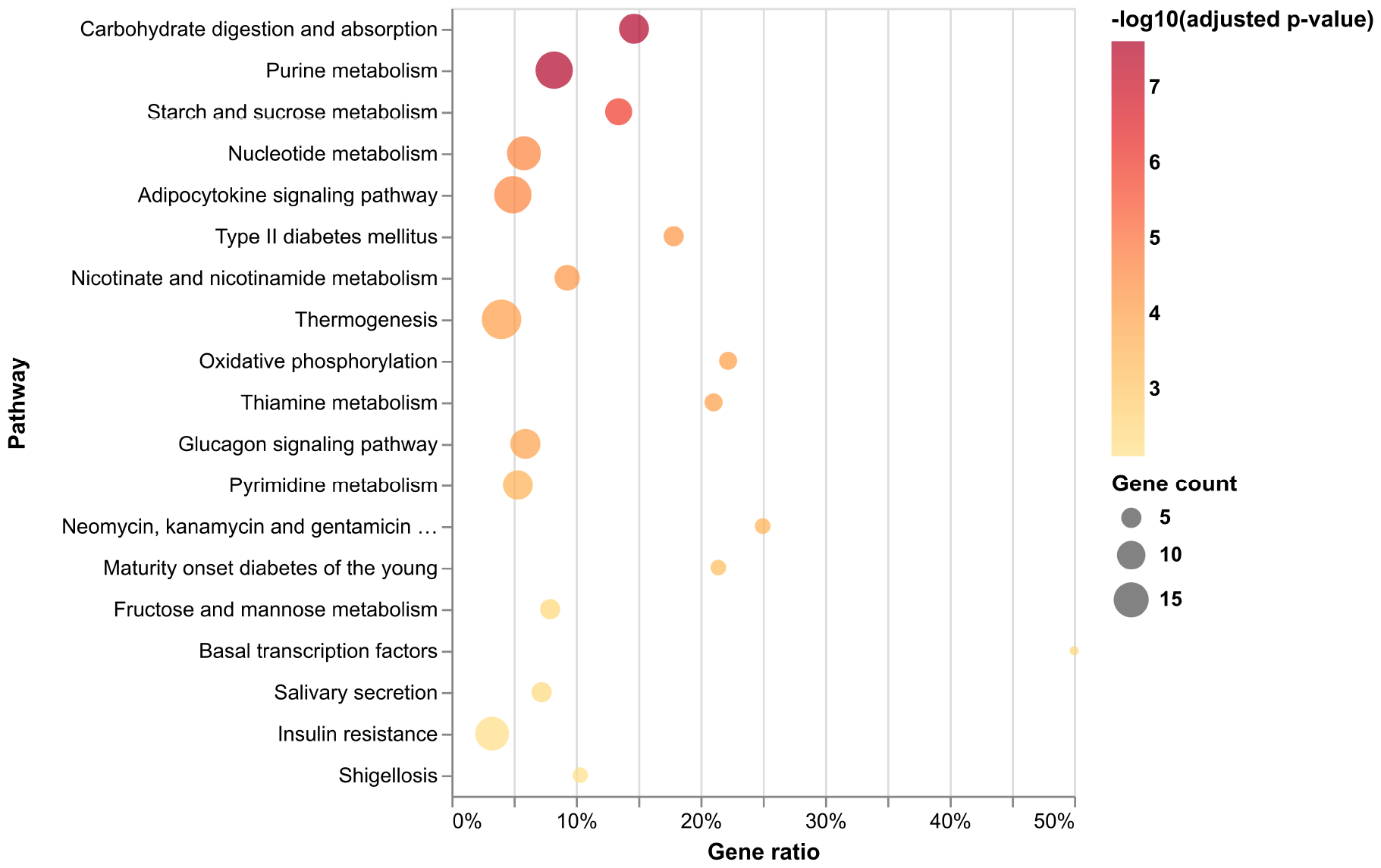
Pathway enrichment analysis of significant genes in renal cancer

### 4.2 Evaluation of identified significant genes based on literature evidence

After post-hos analysis, we further performed SCREENER analysis and co-occurrence analysis to evaluate the association of these significant genes with each cancer type. SCREENER is an AI-based text-mining model that learns the document-level relations between genes and diseases using an attention mechanism and identifies biological connections between target genes and diseases [26]. We did SCREENER analysis by counting the number of PubMed literature identified to have biological relationship between each pair of a significant gene and a cancer type by SCREENER model. Co-occurrence analysis was performed by counting the number of PubMed literature which mentioned each pair of significant gene and cancer type simultaneously. Based on these analysis, we compared the significant genes with SCREENER or co-occurrence evidence among five cancer types (see Figure 9 and Figure 10). Of total 146 significant genes, 87 genes (60%) have literature evidence at least with one cancer types by SCREENER analysis and 125 genes (86%) by co-occurrence analysis, meaning VaMP models can predict the existing known targets at a fairly reasonable level. The list of PubMed IDs for the pairs of significant genes and cancer types which are turned out to have literature evidence by SCREENER and co-occurrence analysis (see Supplementary File 1) could serve as supporting evidences to affirm the performance of VaMP. In SCREENER analysis, the more cancer types a gene is associated with, the more literature evidence there is while this correlation is not clear in co-occurrence analysis as SCREENER analysis. Because SCREENER captures semantic relationship of gene and disease in literature, SCREENER analysis is more reliable than co-occurrence analysis considering the fact that co-occurrence analysis just detects the coexistence of a pair of gene and disease in literature. If a gene is associated with various cancer types, it might be well studied and known previously and the correlation above reflects this phenomenon.

**Figure 9:**
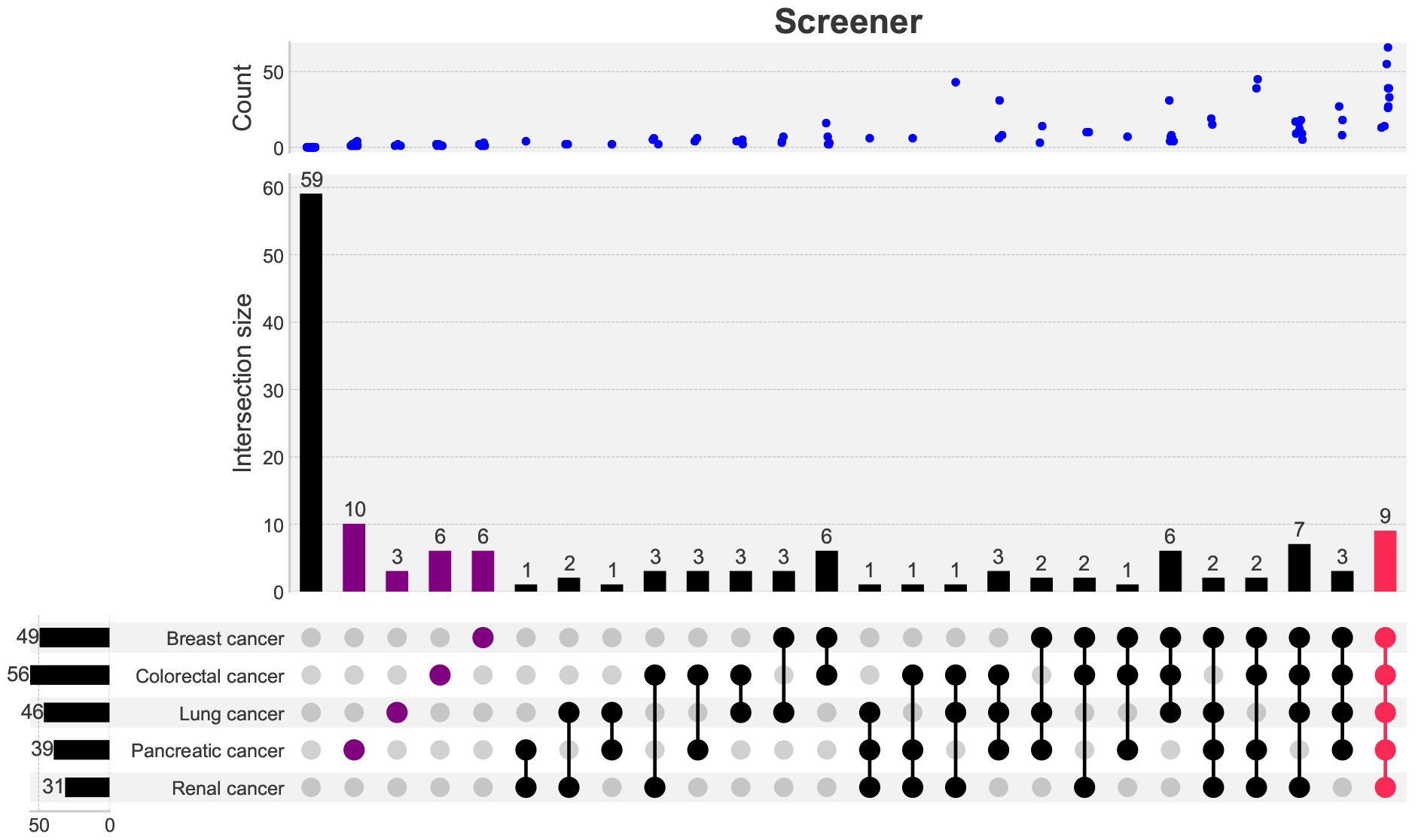
Upset plot of significant genes across five cancer types filtered by SCREENER analysis

**Figure 10:**
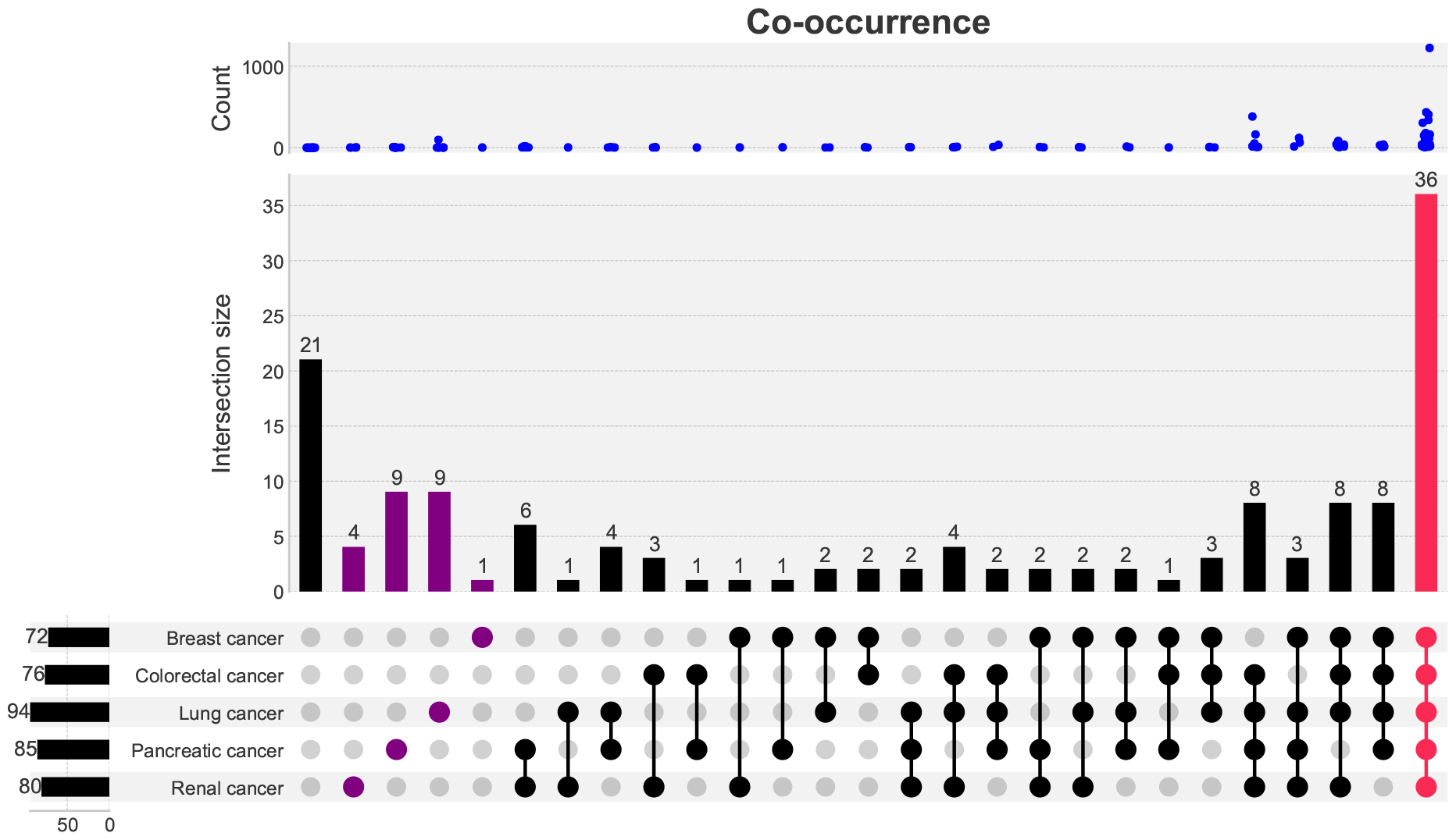
Upset plot of significant genes across five cancer types filtered by co-occurrence analysis

### 4.3 Metabolic signatures of significant genes by post-hoc analysis

Next, significant genes with post-hoc analysis results (log_2_ FC and adjusted *p*-values) and SCREENER and co-occurrence count information for five VaMP models are further studied. (see Figure 11–Figure 15). The *y*-axis value represents each significant gene, and the number of PubMed literature calculated through SCREENER and cooccurrence analysis is shown in two bar plots. The lower *x*-axis value of the heatmap with divided cells indicates the reaction IDs of the significant reactions to which a significant gene is related and the upper *x*-axis value indicates the metabolic subsystems in which the reactions participate. The lower and upper triangle of the rectangle in the heatmap represent log_2_ FC and adjusted *p*-value of each significant reaction respectively.

**Figure 11:**
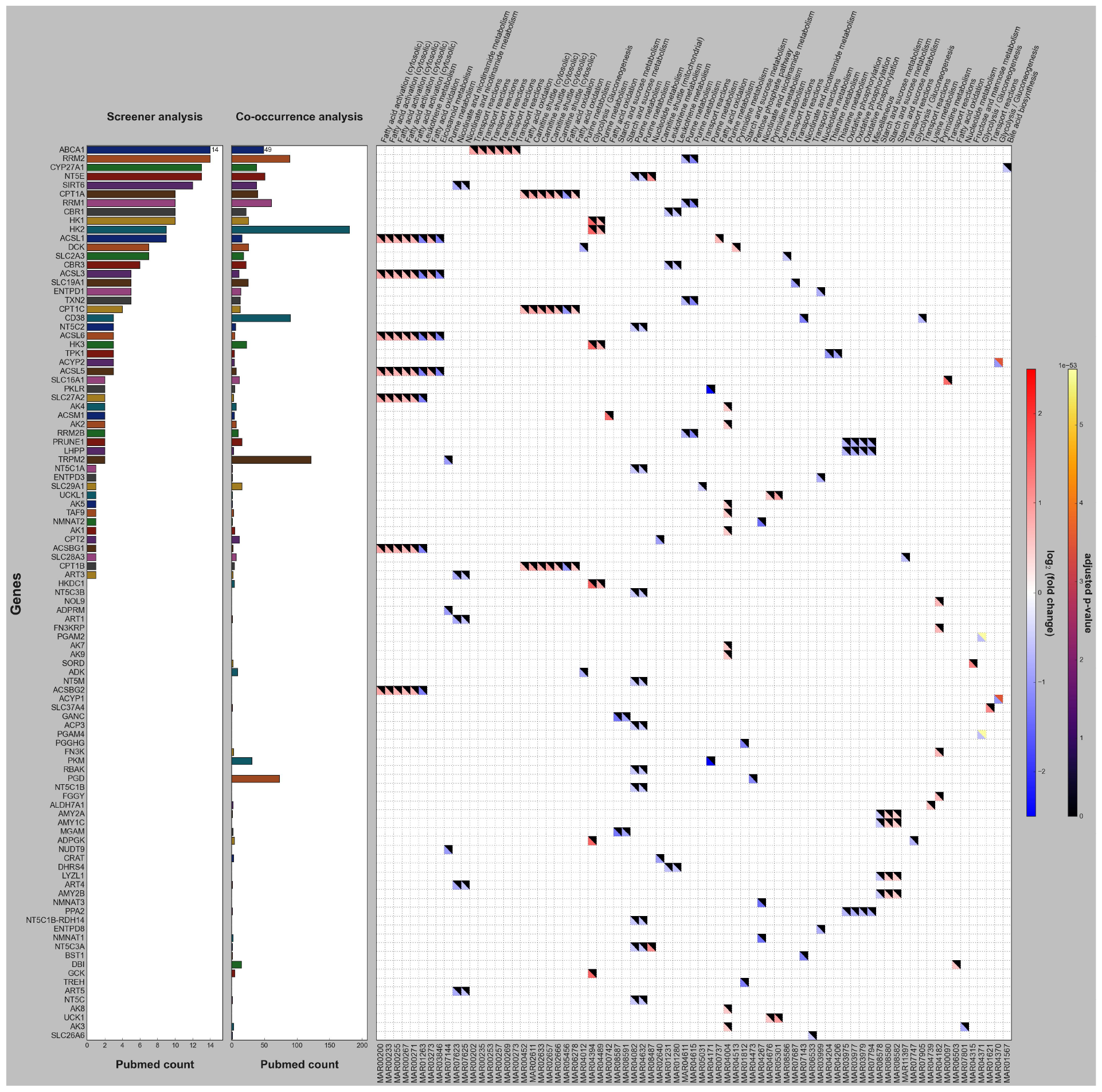
Post-hoc analysis of VaMP of breast cancer

**Figure 12:**
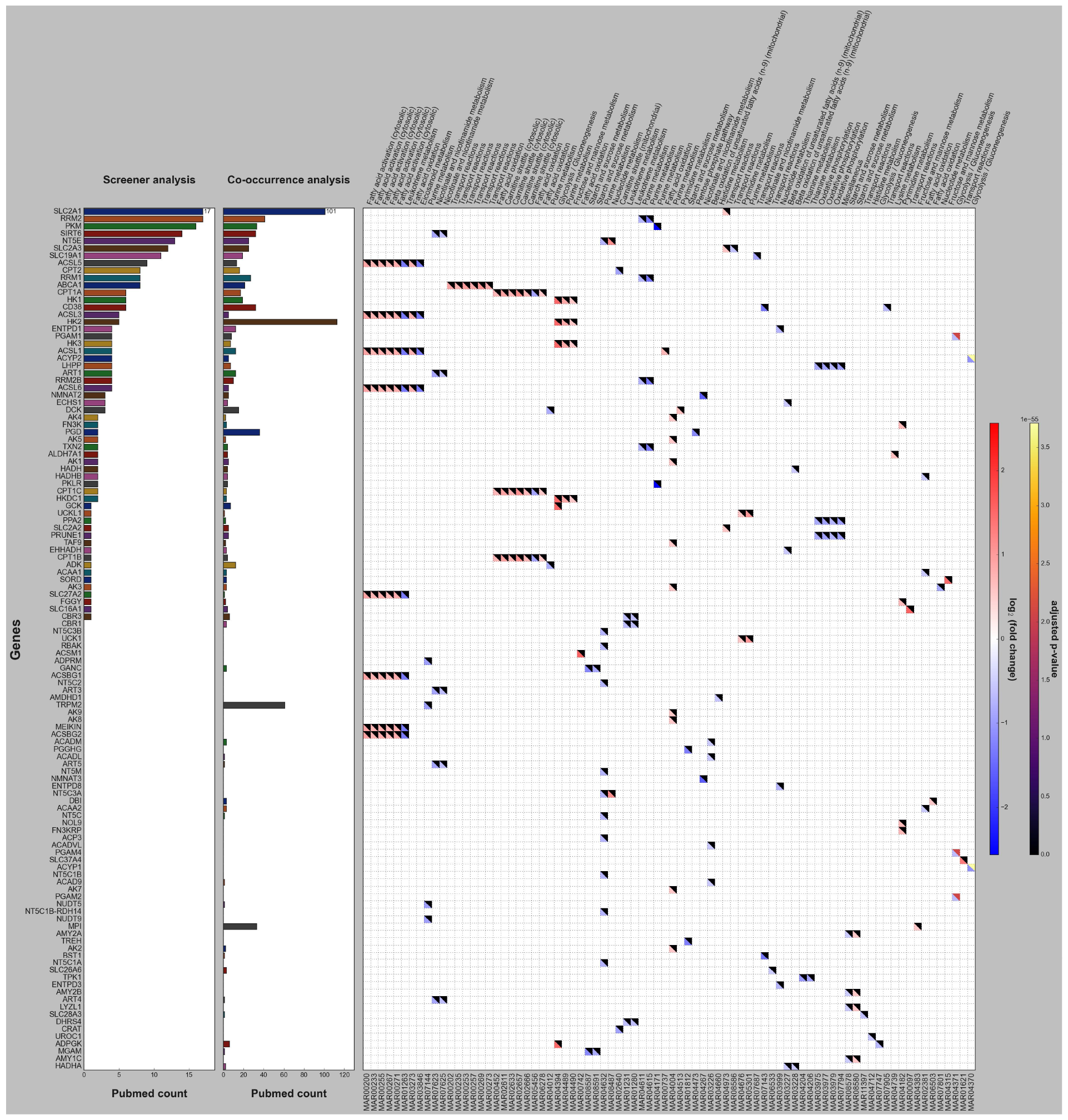
Post-hoc analysis of VaMP of colorectal cancer

**Figure 13:**
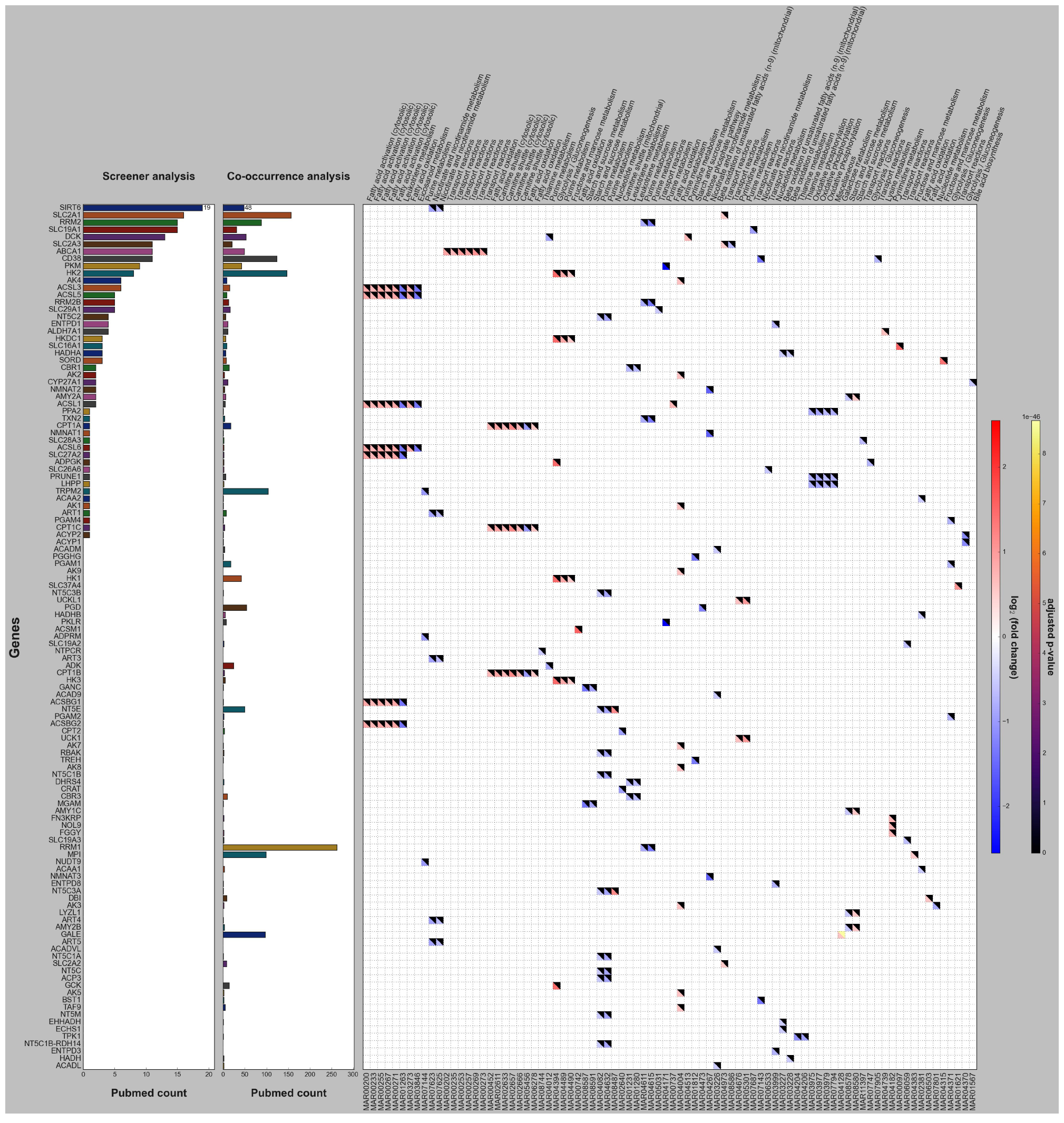
Post-hoc analysis of VaMP of lung cancer

**Figure 14:**
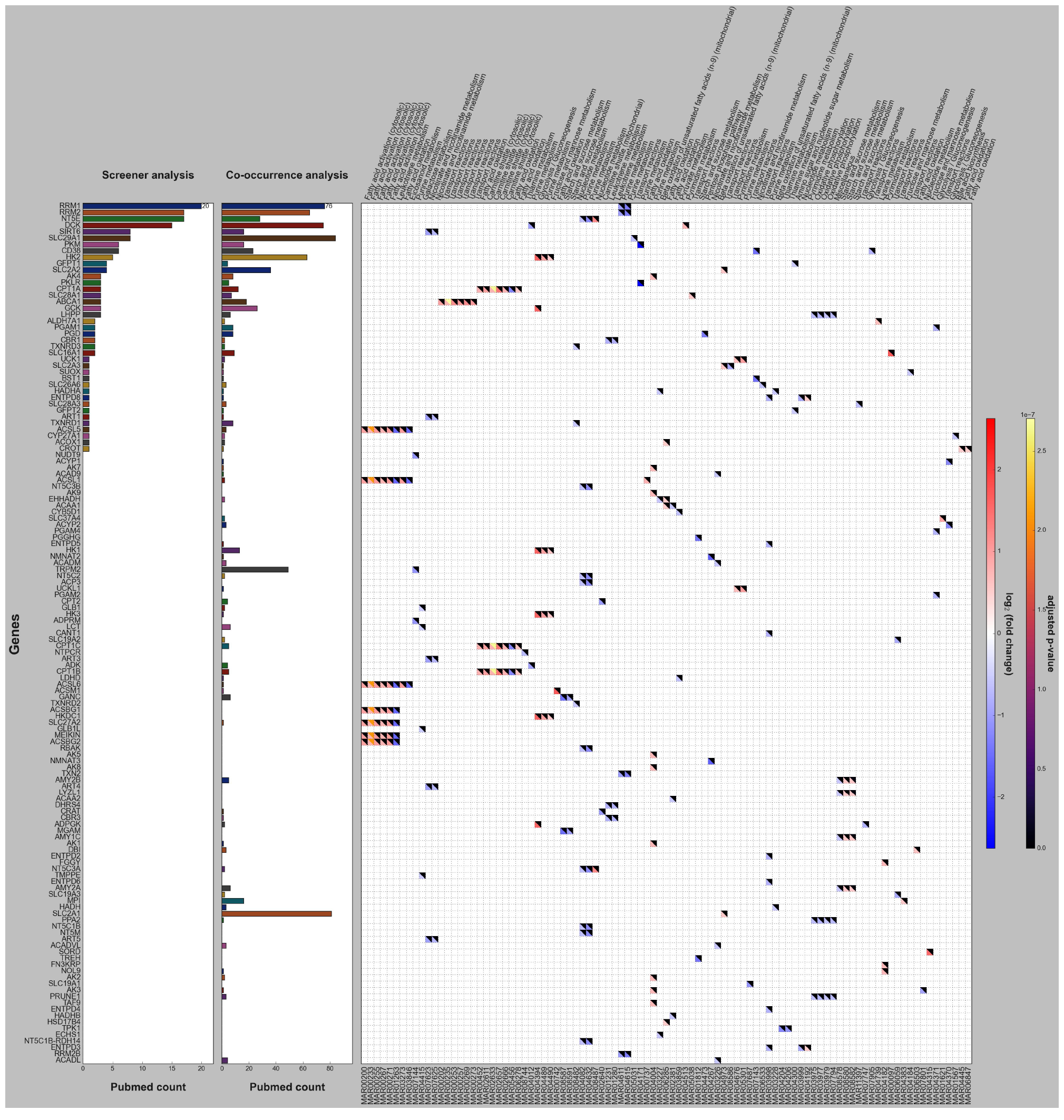
Post-hoc analysis of VaMP of pancreatic cancer

**Figure 15:**
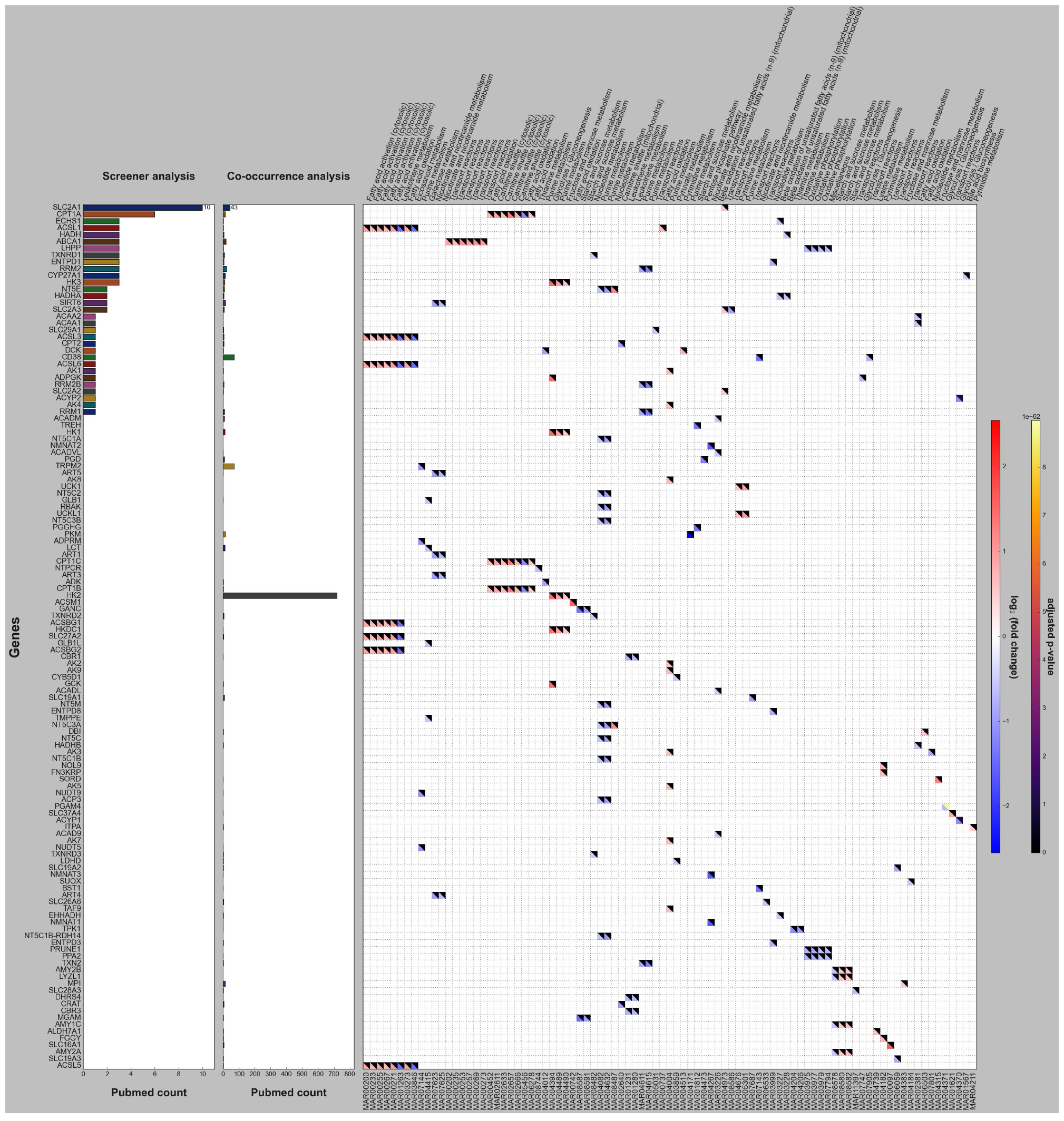
Post-hoc analysis of VaMP of renal cancer

#### 4.3.1 Breast cancer

As for breast cancer, top 5 significant genes by SCREENER count include *ACBA1, RRM2, CYP27A1, NT5E* and *SIRT6*. Each gene participates in the significant metabolic reactions related to transport reactions, purine metabolism, bile acid biosynthesis, nucleotide metabolism and nicotinate and nicotinamide metabolism (see Figure 11). As a result of post-hoc analysis, *ACBA1*-related reactions appear to have greater fluxes in cancer GSMs than normal GSMs meaning its up-regulation in breast cancer. The up-regulation pattern identified by VaMP is previously revealed in the study exploring the relationship of breast cancer and *ACBA1* [28]. Meanwhile, *RRM2, CYP27A1* and *SIRT6*-related reactions show smaller fluxes in cancer GSMs indicating their down-regulation in breast cancer. As for the case of *SIRT6*, metabolic signature identified VaMP is consistent with the study showing breast cancer patient’s survival positively correlated with *SIRT6* abundance [29]. In the case of *NT5E*, purine metabolism-related reactions are up-regulated and while nucleotide metabolism-related reaction is upregulated.

#### 4.3.2 Colorectal cancer

In colorectal cancer, top 5 significant genes are *SLC2A1, RRM2, PKM, SIRT6* and *NT5E. SLC2A1* is related to transport reactions and *PKM* is related to purine metabolism (see Figure 12). *SLC2A1*- related reaction is up-regulated and *RRM2, PKM* and *SIRT6*-related reactions are down-regulated while *NT5E*-related reactions show the same pattern in the case of breast cancer. The metabolic pattern that *SLC2A1* is up-regulated in colorectal cancer well recapitulates the previous experimental study showing *SLC2A1* is highly expressed in colorectal cancer patients [30] and the down-regulation pattern of *PKM* is also supported by the study describing that the *PKM1*, an isoform of *PKM* is decreased in human colorectal carcinomas as compared to non-cancerous tissue [31].

#### 4.3.3 Lung cancer

Next, top 5 significant genes in lung cancer turn out to be *SIRT6, SLC2A1, RRM2, SLC19A1* and *DCK* and the last two genes are related to transport reactions, purine metabolism and pyrimidine metabolism (see Figure 13). *SIRT6, RRM2* and *SLC19A1* are down-regulated and *SLC2A1* are upregulated in lung cancer. The metabolic relationships of *SIRT2* and *SLC2A1* with lung cancer captured by VaMP are previously validated by two experimental studies, the one describing that *SIRT6* is generally down-regulated and linked to tumorigenesis in non-small cell lung carcinoma [32] and the other observing that strong up-regulation of *SLC2A1* in patients with lung adenocarcinoma and unveiling that *SLC2A1* was significantly correlated with prognosis [33]. *DCK* shows down-regulated pattern in purine metabolism-related reaction while up-regulated pattern is observed in pyrimidine metabolism-related reaction.

#### 4.3.4 Pancreatic cancer

Top 5 significant genes in pancreatic cancer includes *RRM1, RRM2, NT5E, DCK* and *SIRT6* (see Figure 14). *RRM1, RRM2* and *SIRT6* appear to be down-regulated in pancreatic cancer and *NT5E* and *DCK* shows the same mixed pattern in the cases of breast and lung cancer. The down-regulation pattern of *SIRT6* in pancreatic cancer identified by VaMP is consistent with the experimental study finding *SIRT6* inactivation accelerates pancreatic ductal adenocarcinoma progression and metastasis *via* up-regulation of *LIN28B*, a negative regulator of the let-7 microRNA [34]. In the case of *NT5E*, among two mixed patterns, the up-regulation pattern in pancreatic cancer is aligned with the study describing *NT5E* gene was almost absent in normal pancreas but strongly up-regulated in pancreatic tumors [35].

#### 4.3.5 Renal cancer

Lastly, *SLC2A1, CPT1A, ECHS1, ACSL1* and *HADH* are placed at top 5 significant genes in renal cancer (see Figure 15). *CPT1A* is related to fatty acid oxidation and carnitine shuttle reactions and *ECHS1* is related to beta-oxidation of unsaturated fatty acids reaction. *ACSL1* participates in the significant reactions related to fatty acid activiation/oxidation, leukotriene metabolism and eicosanoid metabolism and *HADH* is related to beta-oxidation of unsaturated fatty acids reaction. According to post-hoc analysis, *SLC2A1* shows up-regulated pattern while *ECHS1* and *HADH* have down-regulated pattern in renal cancer. These metabolic patterns of *SLC2A1, ECHS1* and *HADH* in renal cancer identified by VaMP recapitulate the previous experimental studies well [36–39]. In the case of *CPT1A*, it is up-regulated in carnitine shuttle-related reactions but shows mixed pattern in fatty acid oxidation reactions. As for *ACSL1*, it shows up-regulated pattern in fatty acid activiation/oxidation-related reactions while it is down-regulated in leukotriene and eicosanoid metabolism-related reactions. No-tably, the conflicting pattern of *ACSL1* in different metabolic subsystems (up-regulation in fatty acid metabolism and down-regulation in leukotriene and eicosanoid metabolism) precisely recapitulate the study showing that the *ACSL1* high-expression sub- group in clear cell renal cell carcinoma had enriched fatty acid metabolism-related pathways and high expression of ferroptosis-related gene while the *ACSL1* low-expression subgroup exhibited higher immune and microenvironment scores, elevated expression of immune checkpoints PDCD1, CTLA4, LAG3, and TIGIT [40]. Considering the fact that eicosanoids and leukotrienes are lipid-based signaling molecules that play a unique role in innate immune responses [41], the down-regulation pattern of *ACSL1* in eicosanoid and leukotriene metabolism captured by VaMP can be reasonably related to the influence of *ACSL1* low expression on immune system in renal cancer as described by the previous study.

Additionally, the pattern of *RRM1/2* and *SLC2A1* appearing to prevail across five cancer types can be explained by the previous study showing these two genes’ pan-cancer role as an oncogene or a cancer biomarker [42, 43].

Overall, smaller adjusted *p*-values are observed in significant genes which have SCREENER evidence (SCREENER count > 0) implying that five VaMP models successfully recapitulate previously known relationships of cancer-causing genes and the cancer types. Therefore, other significant genes with no SCREENER evidence but low adjusted *p*-values can be potential novel cancer-causing genes and also be therapeutic targets for cancer treatment.

### 4.4 Linking significant genes to cancer-causing variants

For identifying what type of variant contribute to causing the certain type of cancer, the correlation between the fluxes of significant reactions and their related significant genes’ variants are further analyzed by LOR-correlation (see §3.6). LOR-correlation plot of five cancer types and adjusted *p*-values for each pair of variants in significant reactions can be found in Supplementary File 2. Based on these data, the greater LOR means more frequent presence of the variant in the cancer patients. Therefore, the greater the difference in flux between a variant with a larger LOR value and a variant with a smaller LOR value, the more likely it is to contribute to causing cancer.

## 5 Discussion

We have developed a novel approach, VaMP, which employs an end-to-end neural network for predicting the phenotype of a given patient based on their genetic variants. The key innovation lies in the representation of the latent space, which captures metabolic fluxes derived from a GSM. The training data for VaMP consists of mutation information and phenotypes, eliminating the need for preparing training data for metabolic flux. The encoder of VaMP maps genetic variants to flux through a series of steps involving gene damage computation, derivation of reaction damages, and solving the MOMA to obtain the flux. The gene damage computation employs a parameterized approach with constraints, ensuring positive values for each element. The reaction damage, derived from gene damage, considers gene product associations en-coded in the GSM. The resulting metabolic flux is obtained by solving the MOMA, incorporating a quadratic programming framework. The decoder of VaMP predicts phenotype probabilities by calculating differences between input flux and reference flux. These differences are then fed into a neural network for the final probability prediction.

We further utilized a human GSM from Metabolic Atlas and genetic variant data from Genomics England and gnomAD for training VaMP models for five cancer types. Results reveal significant genes for each cancer type, obtained through post-hoc analysis, and their corresponding metabolic signatures. Evaluation using SCREENER and co-occurrence analyses demonstrates the ability of VaMP to predict known gene-disease relationships and their identified metabolic signatures are found to be strongly supported by various literature based on experimental studies. Additionally, we further analyzed the results of five VaMP models to link significant genes to cancer-causing variants through LOR-correlation analysis.

On the other side, several improvements can be considered as the future research topics. One of the most obvious ideas is improving the encoder. Currently, the encoder is based on a linear regression model with the number of variants per consequence as input. This structure is oversimplified, and even the input features are too summarized. Thus, current encoder is not enough information to inspect specific variants. To overcome this issue, we should make the structure and the input features more complex. One can use protein language models (*e*.*g*., ESM2 [44]) to compute the embedding vectors of reference protein sequence and mutated protein sequence, and use the difference of embedding vectors as the input features. This approach seems natural and promising, but it will require huge amounts of resources since it should handle millions of tokens per each sample. Another approach is to train a lightweight proxy model to simulate the protein language model, and use the model to compute the mutated embedding vectors. We consider two possible approaches for training the proxy model: (a) use knowledge distillation [45] (b) use another set of labels (*e*.*g*., annotated labels from AlphaMissense [46])

In summary, the development of VaMP is not merely an academic pursuit but a crucial response to the evolving demands of cancer research, genetic exploration, and the growing intersection of machine learning with the life sciences. VaMP has the capacity to emerge as a potent tool for predicting cancer phenotypes based on genetic variants, showcasing its effectiveness in capturing known relationships and highlighting its potential for uncovering novel gene-disease associations.

## Supporting information

The list of PubMed IDs for the pairs of significant genes and cancer types analyzed by SCREENER and co-occurrence analysis

LOR-correlation plots of five cancer types and adjusted p-values for each pair of variants in significant reactions

## Acknowledgements

This research was made possible through access to data in the National Genomic Research Library, which is managed by Genomics England Limited (a wholly owned company of the Department of Health and Social Care). The National Genomic Research Library holds data provided by patients and collected by the NHS as part of their care and data collected as part of their participation in research. The National Genomic Research Library is funded by the National Institute for Health Research and NHS England. The Wellcome Trust, Cancer Research UK and the Medical Research Council have also funded research infrastructure.

## A Differentiate the Quadratic Programming

One can follow [18] to derive the efficient forward and backward differentiation rules below. Consider the following optimization problem:

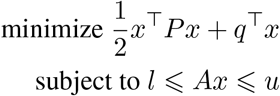

where *x ∈ ℝ*^*n*^ is the variable and *P ∈ 𝕊*^*n×n*^,*A∈ ℝ*^*m×n*^, *q ∈ ℝ*^*n*^, *l ∈*(*{ −∞} ∪ ℝ*)^*m*^ and *u∈* (*ℝ∪ { ∞}*)^*m*^ are parameters. Note that this is the canonical input form of [15].

Let *x*^*∗*^ *∈ ℝ*^*n*^ be the optimal solution of the problem and *y*^*∗*^ *∈ ℝ*^*m*^ be the corresponding dual solution. In addition, define index sets *I*_+_, *I*_*−*_, *I*_0_ *⊂ {* 1, .., *m}* as

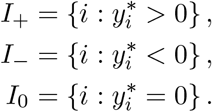

Furthermore, for any vector *v ∈ ℝ*^*m*^, define *v*_+_ = *v*_*I*+_ and vice versa. Similary, define 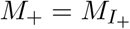. for the matrix *M ∈ ℝ*^*m×n*^. Then, the optimality conditions for the problem can be stated as followed:

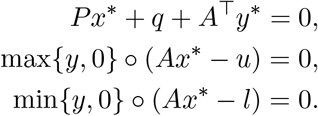

Taking the differentiate with respect to *x*^*∗*^, *y*^*∗*^, *q, l, u* will give

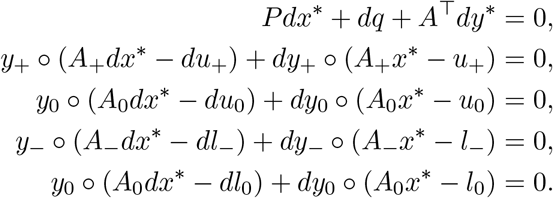

Note that *A*_+_*x*^*∗*^ *− u*_+_ = 0 and *A*_*−*_*x*^*∗*^ *− l*_*−*_ = 0, since *y*_+_, *y*_*−*_≠ 0. Also, by the definition, *y*_0_ = 0.

Therefore,

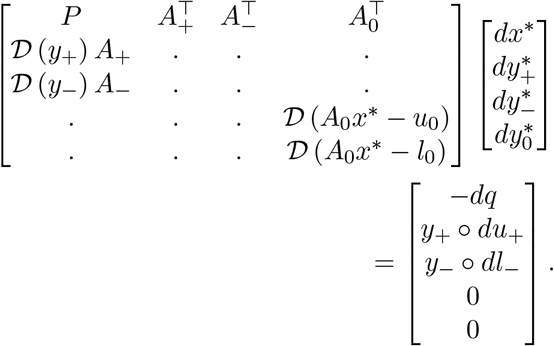

where *𝒟* (.) is the diagonal matrix operator.

Last two rows implies that 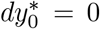 so corresponding rows and columns can be removed and this gives our core rule for the forward/reverse AD.

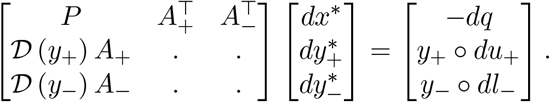

### A.1 Forward Rule

The goal is to calculate *dx*^*∗*^*/dθ* while *dq/dθ, dl/dθ, du/dθ* are given, which can be accomplished by solving the following equation.

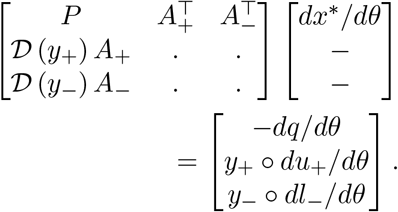

Since *𝒟* (*I, y*_+_, *y*_*−*_) is invertible, the above equation can be converted into

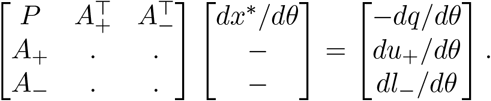

### A.2 Reverse Rule

The goal is to calculate *df/dq, df/dl, df/du* while *df/dx*^*∗*^ is given. To derive the reverse rule, first convert the core rule as in the matrix form:

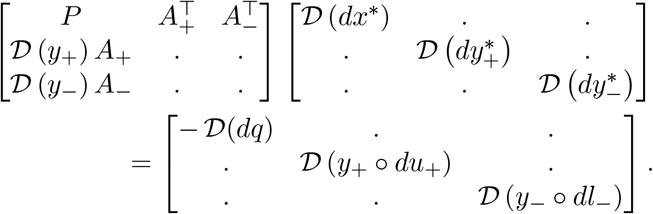

Inverting and transposing the equation will give

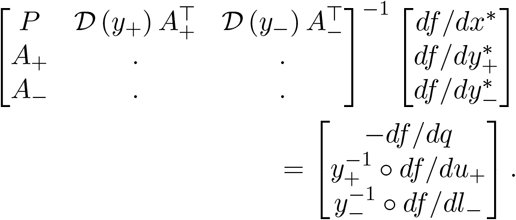

Similarly, above equation can be converted into the final form

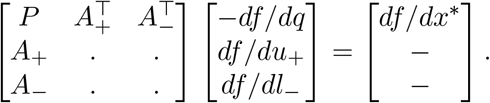

Notice that the coefficient matrix of the both forward rule and reverse rule are same and symmetric, so one can cache the efficient factorization (*e*.*g*., *LDL*^*T*^ factorization) of the coefficient matrix and reuse to speed up the computation.

## Footnotes

1 https://ican.standigm.com

