## Supplementary figures and images for "Learning the Relationship Between Variants, Metabolic Fluxes and Phenotypes"

### MAR00200_ENSG00000123983.png

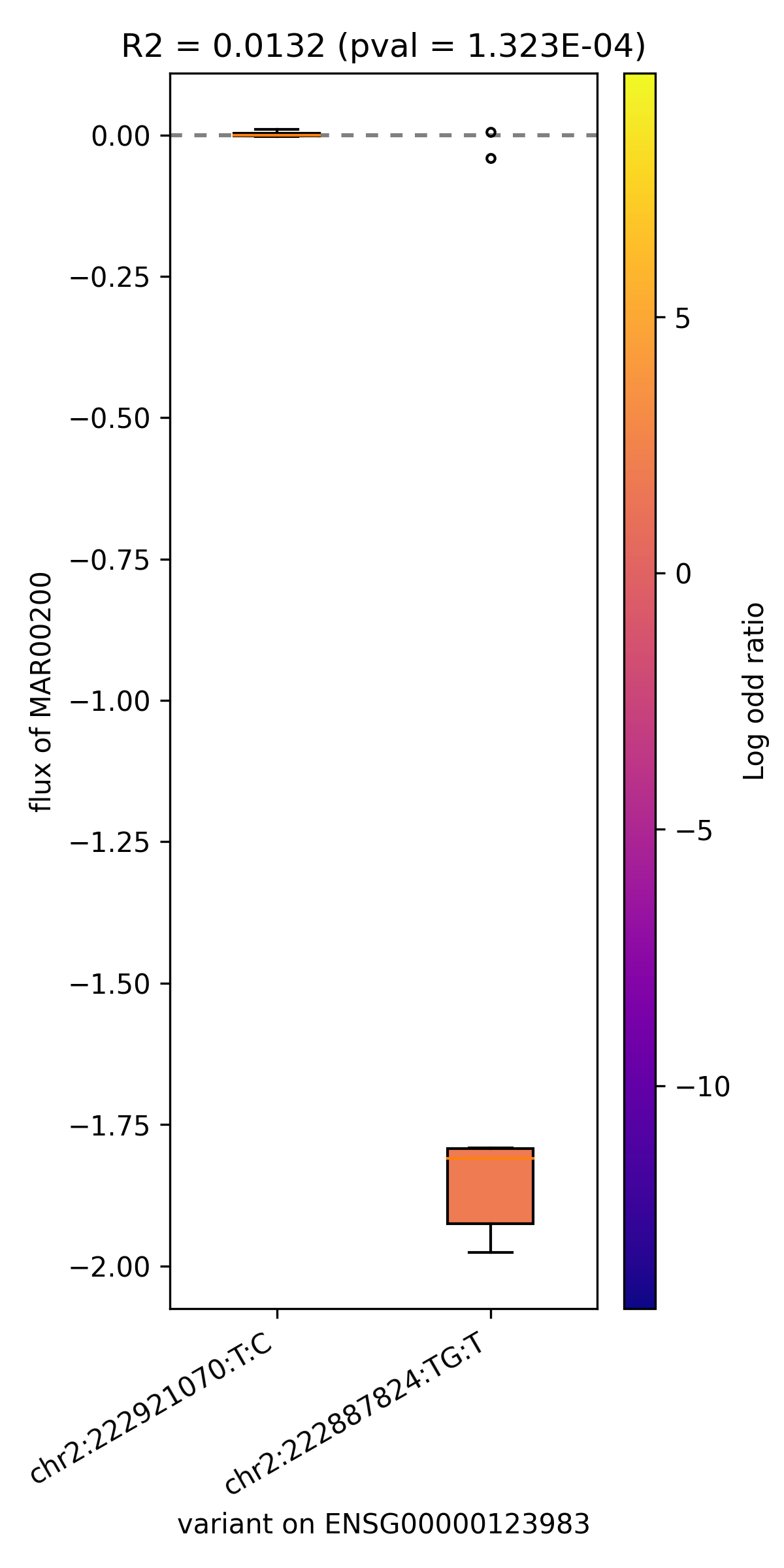

### MAR00200_ENSG00000151726.png

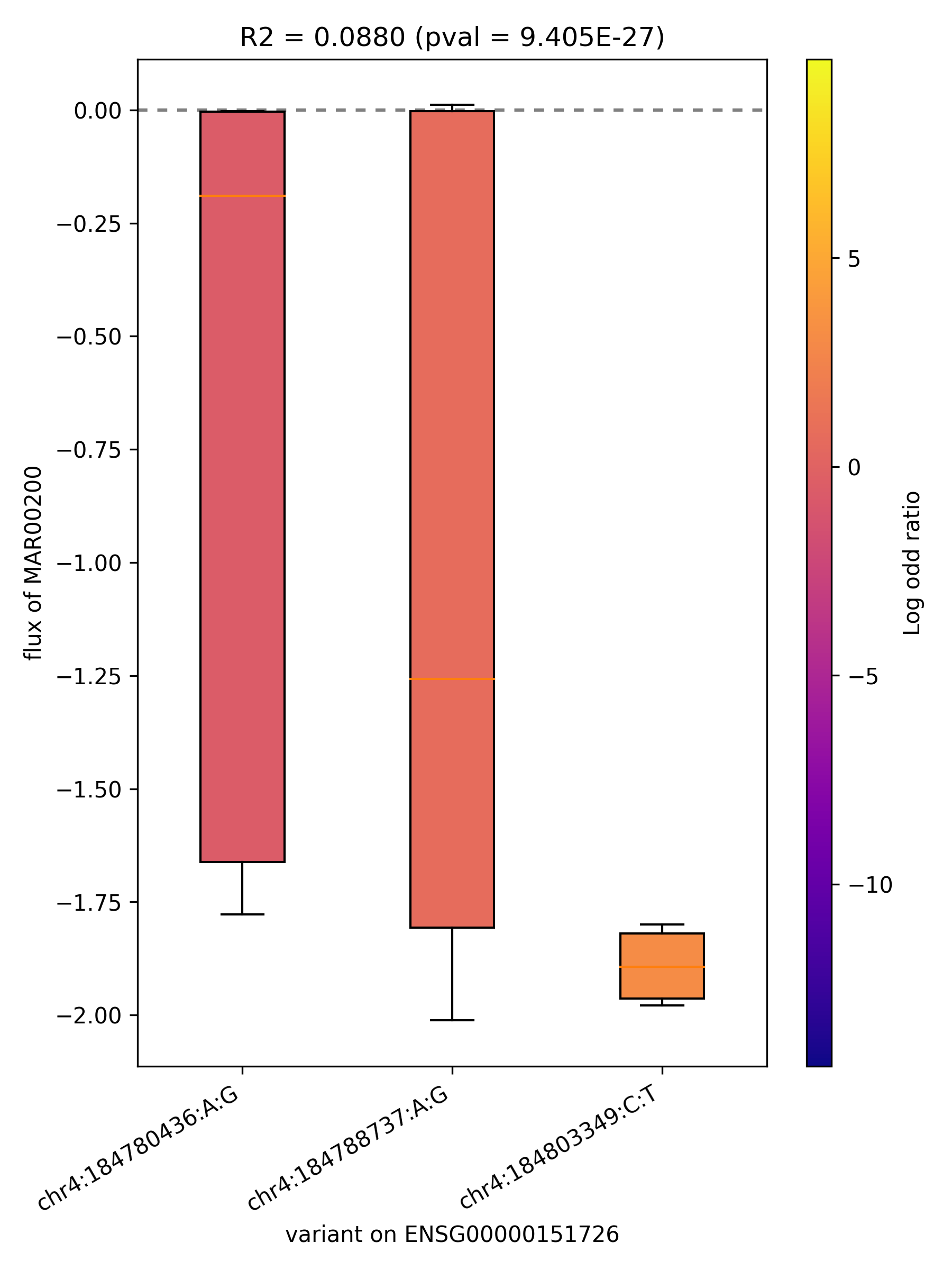

### MAR00200_ENSG00000164398.png

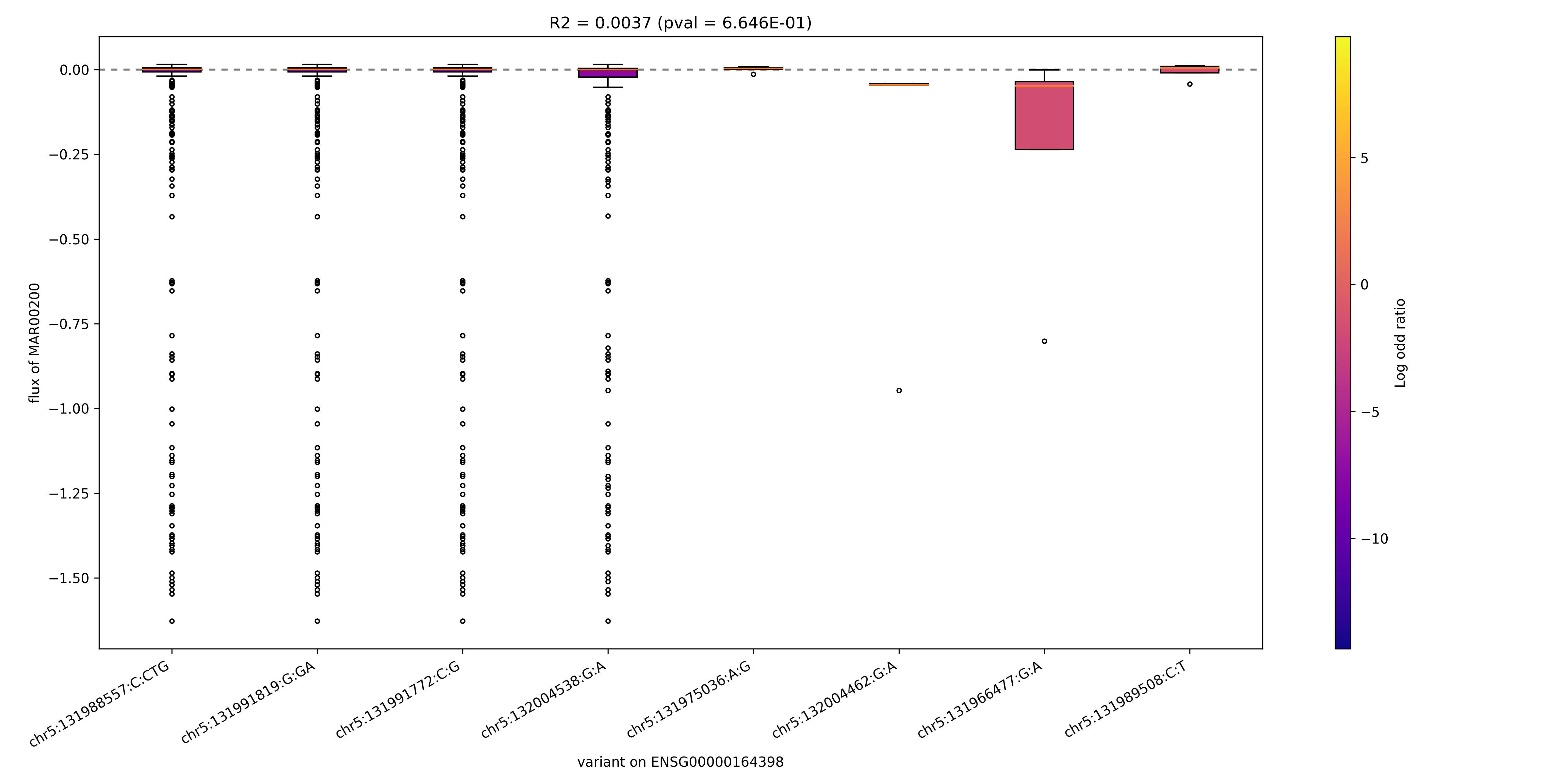

### MAR00200_ENSG00000239642.png

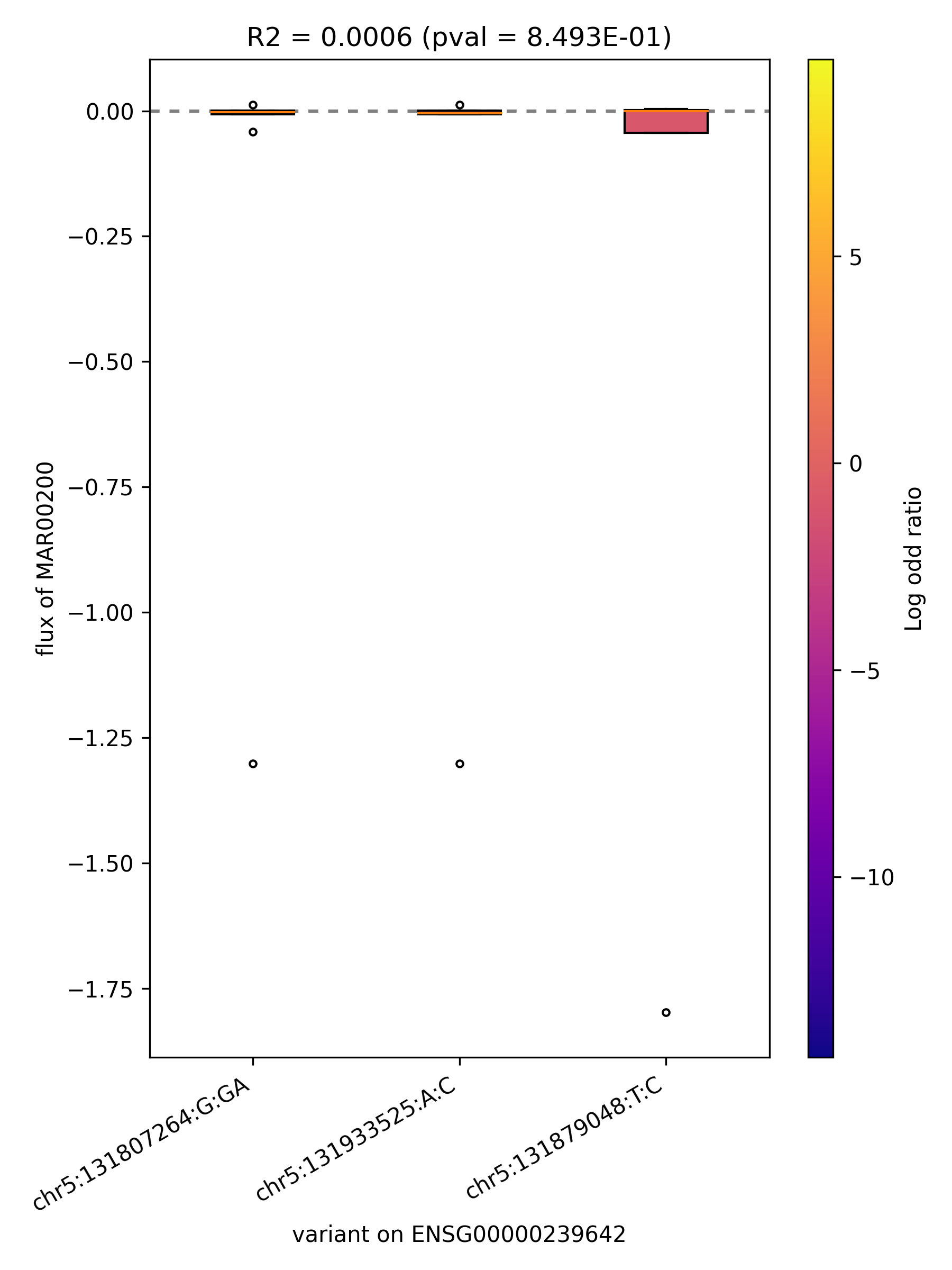

### MAR00233_ENSG00000103740.png

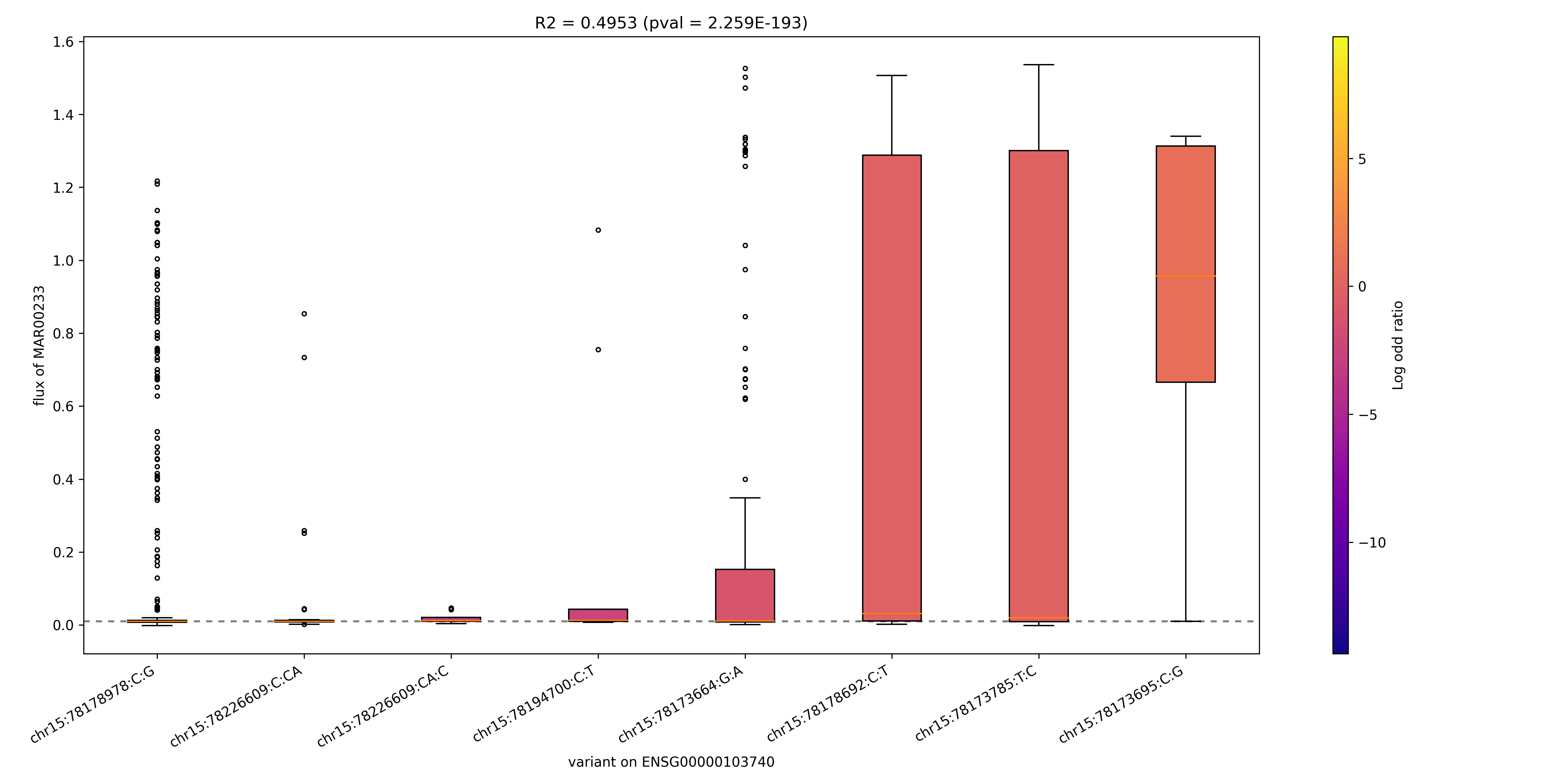

### MAR00233_ENSG00000130377.png

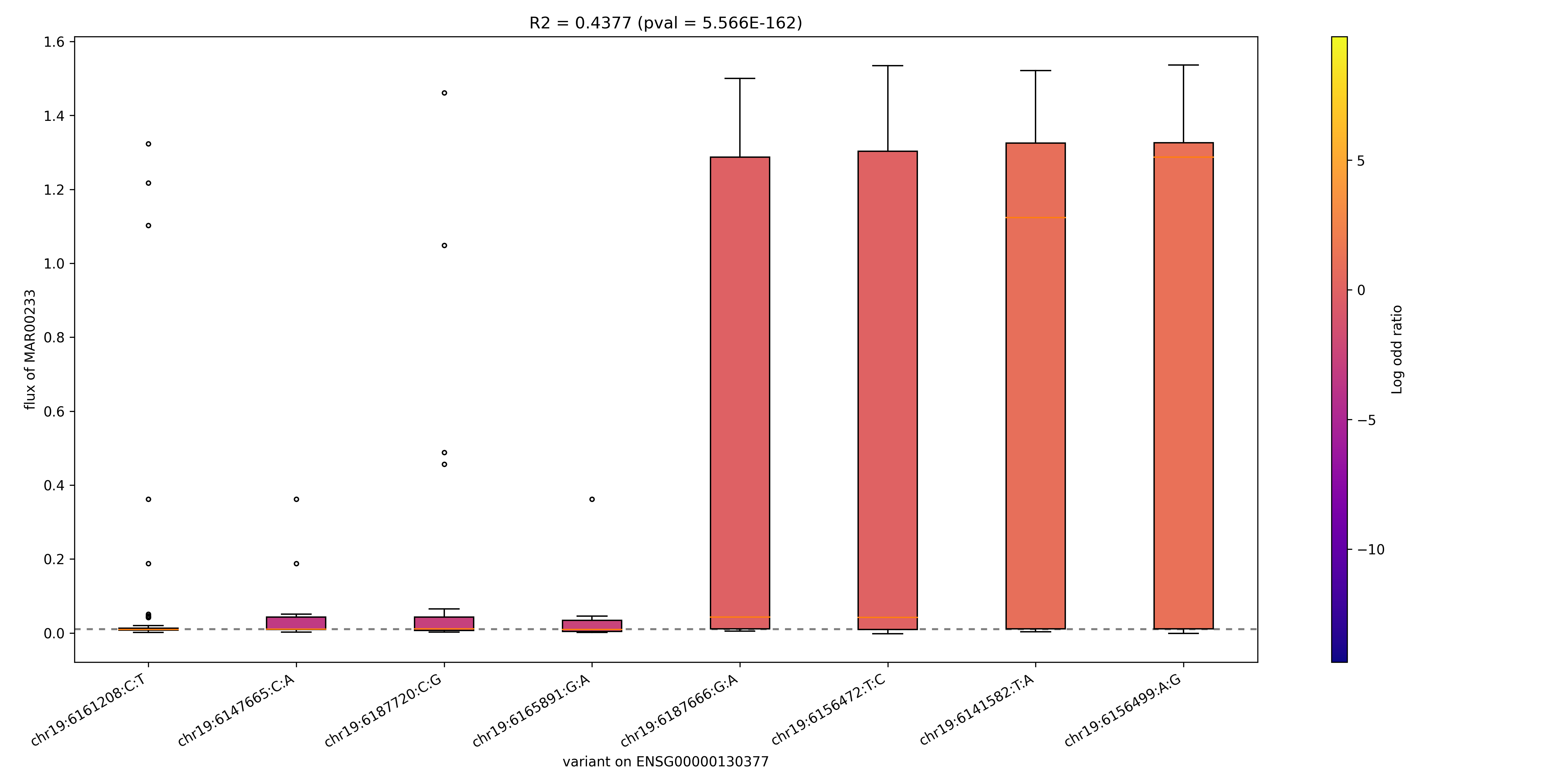

### MAR00233_ENSG00000140284.png

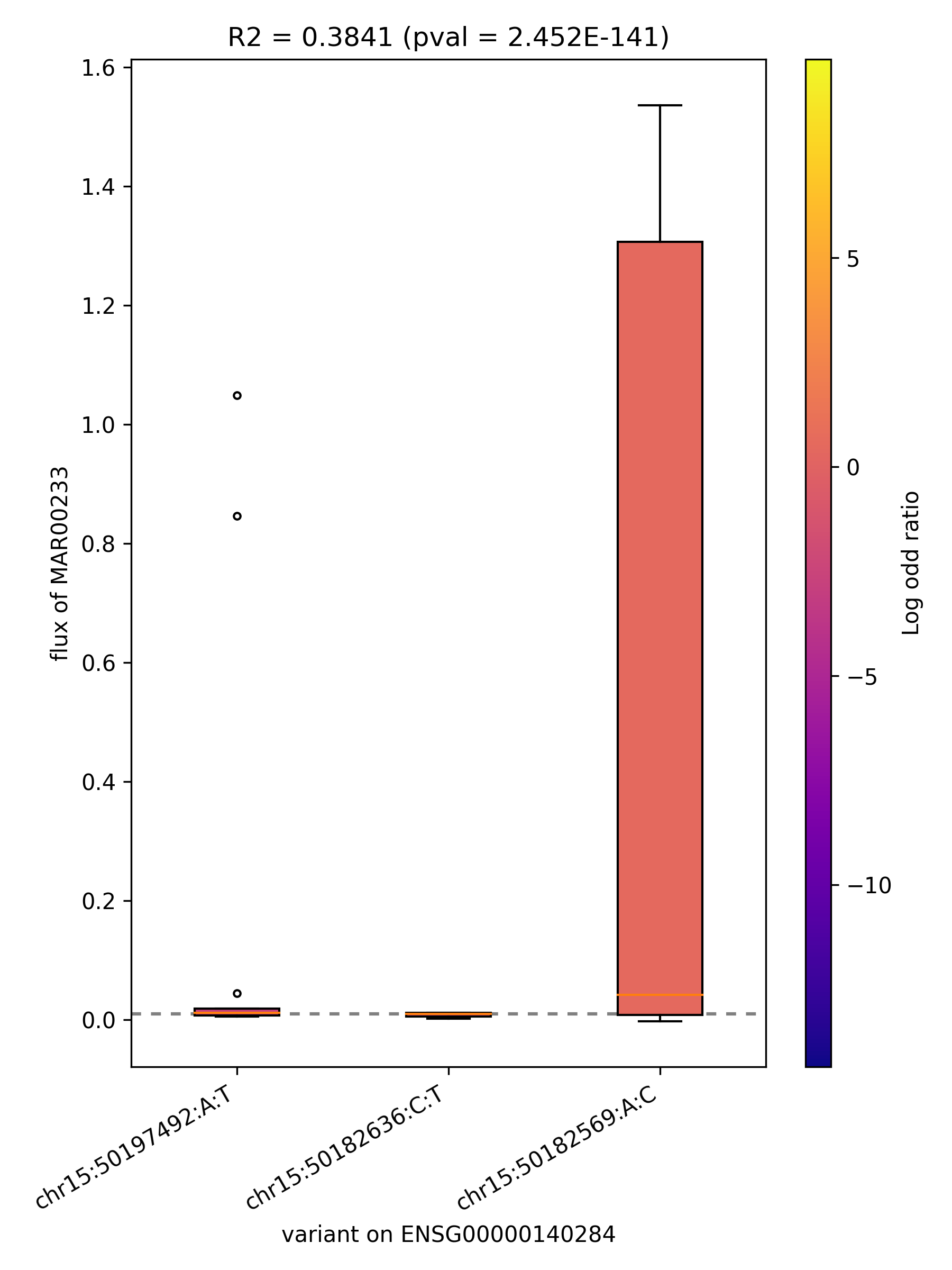

### MAR00233_ENSG00000197142.png

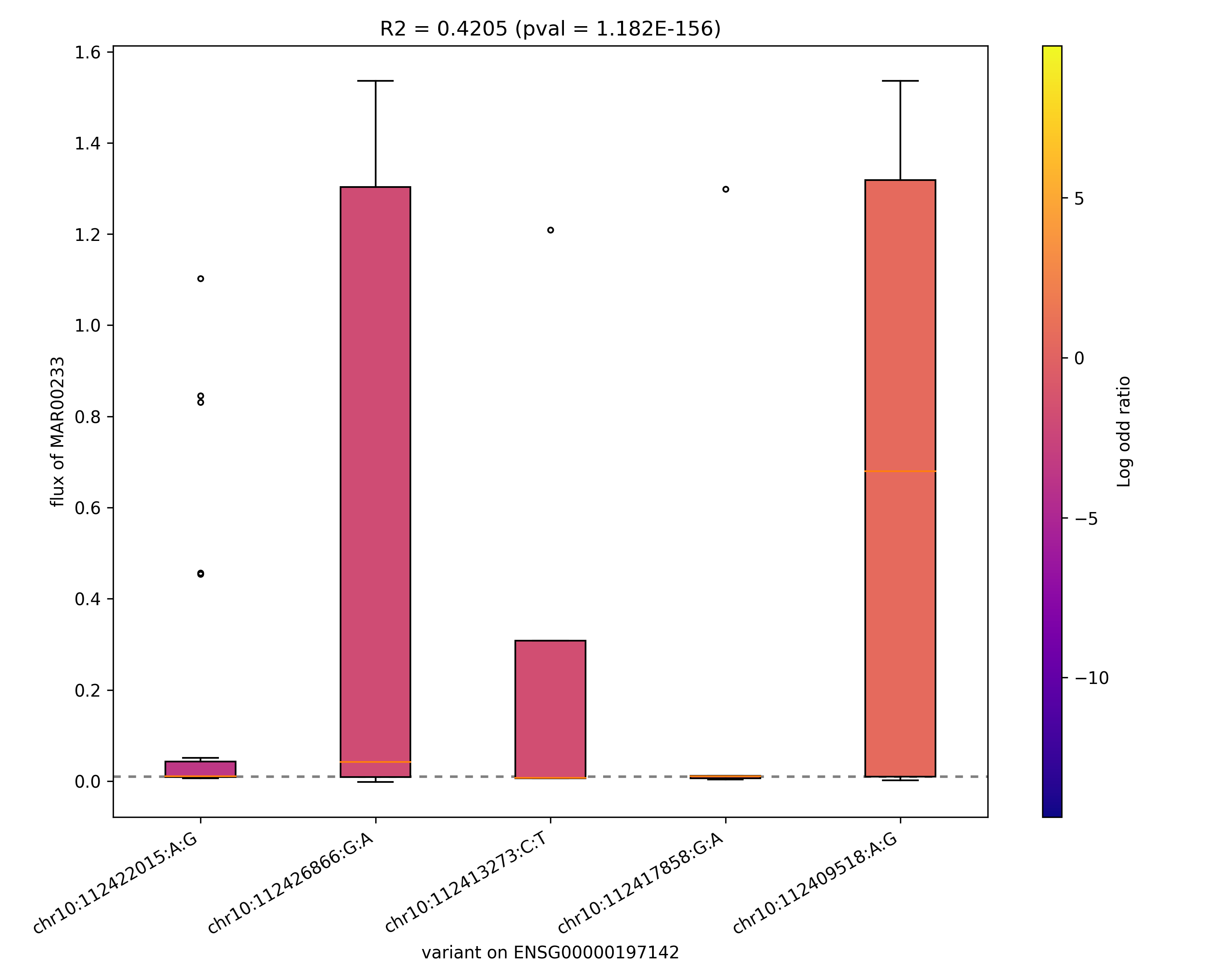

### MAR00235_ENSG00000165029.png

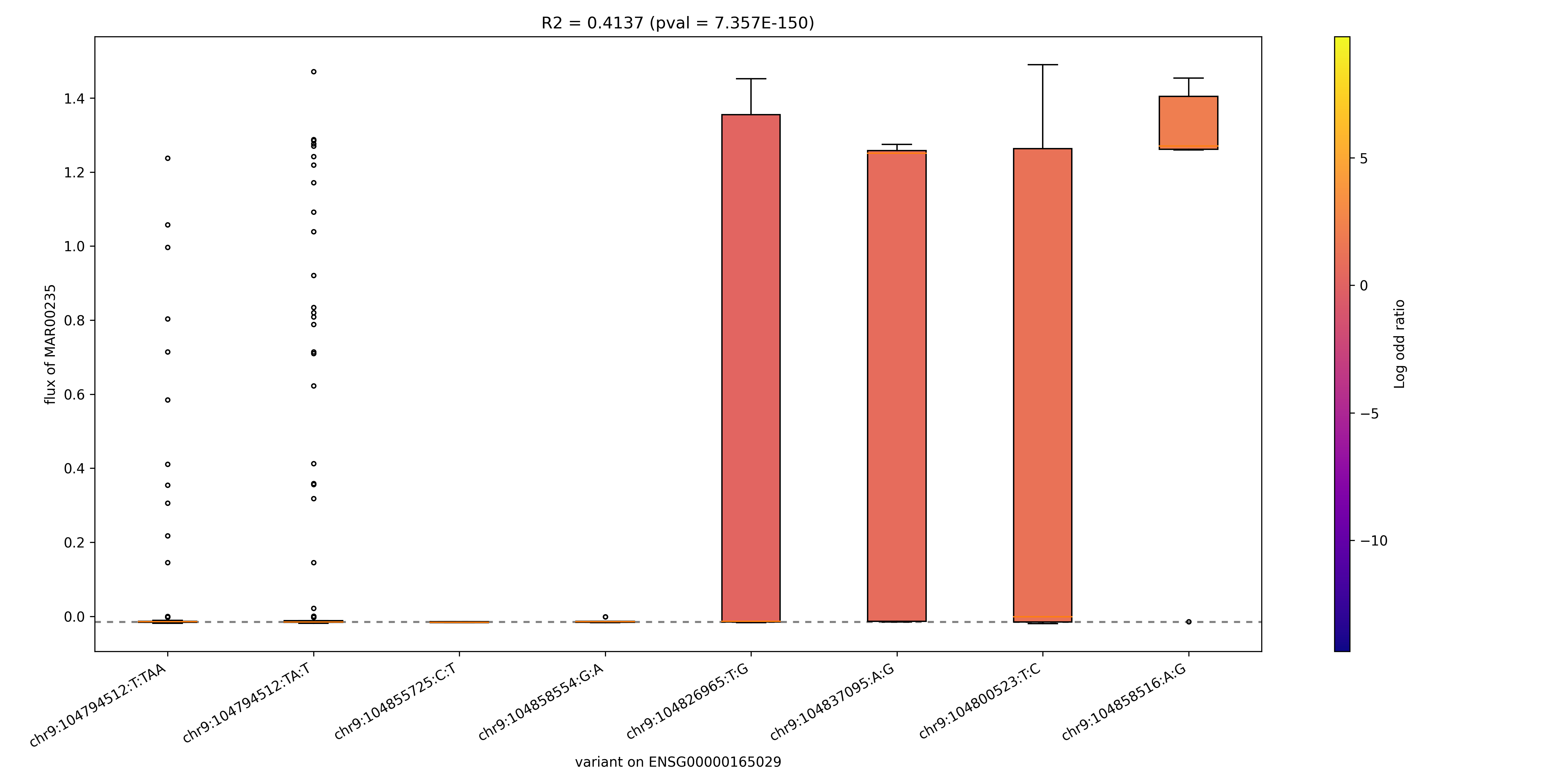

### MAR00255_ENSG00000123983.png

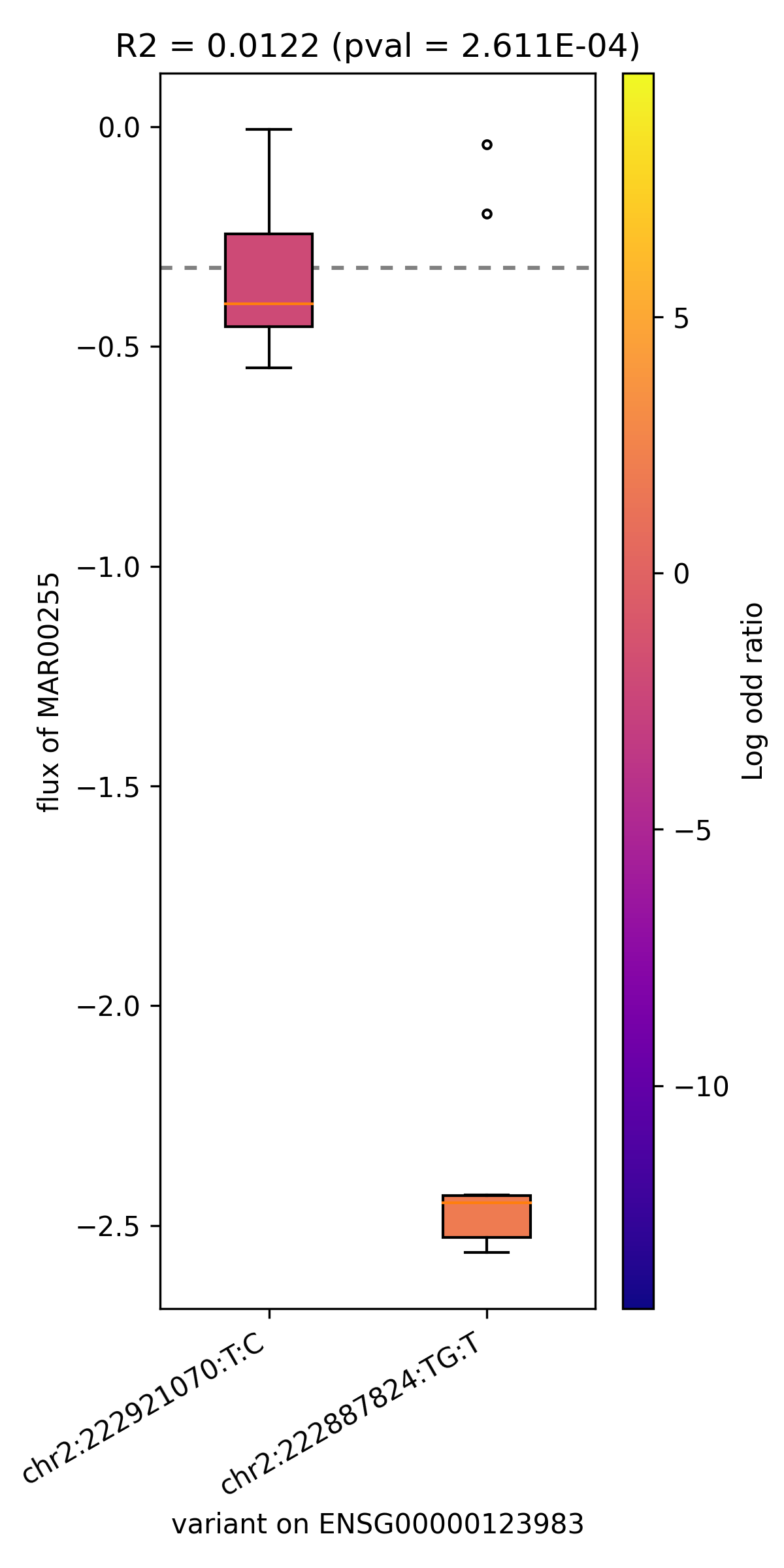

### MAR00255_ENSG00000151726.png

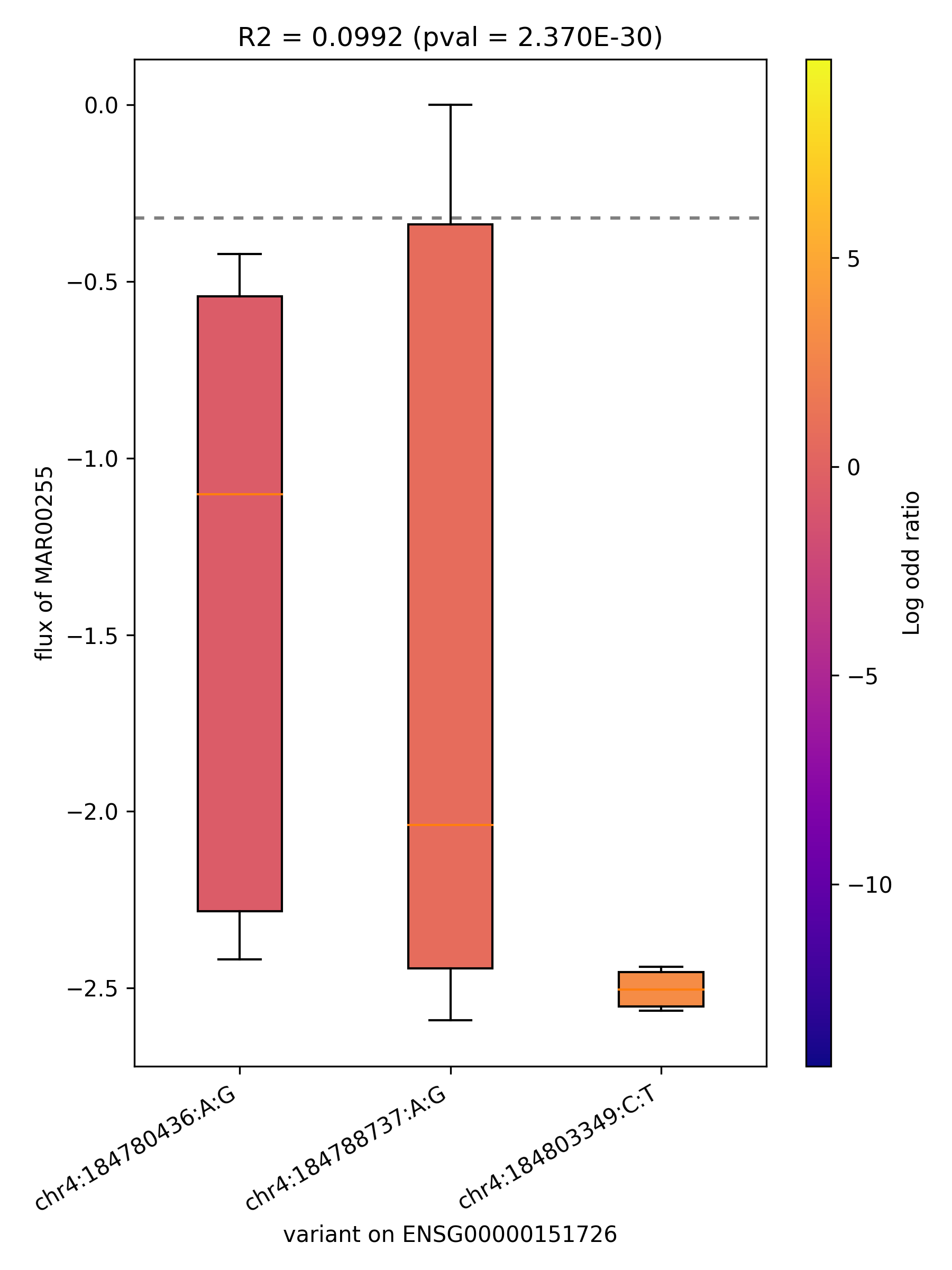

### MAR00255_ENSG00000164398.png

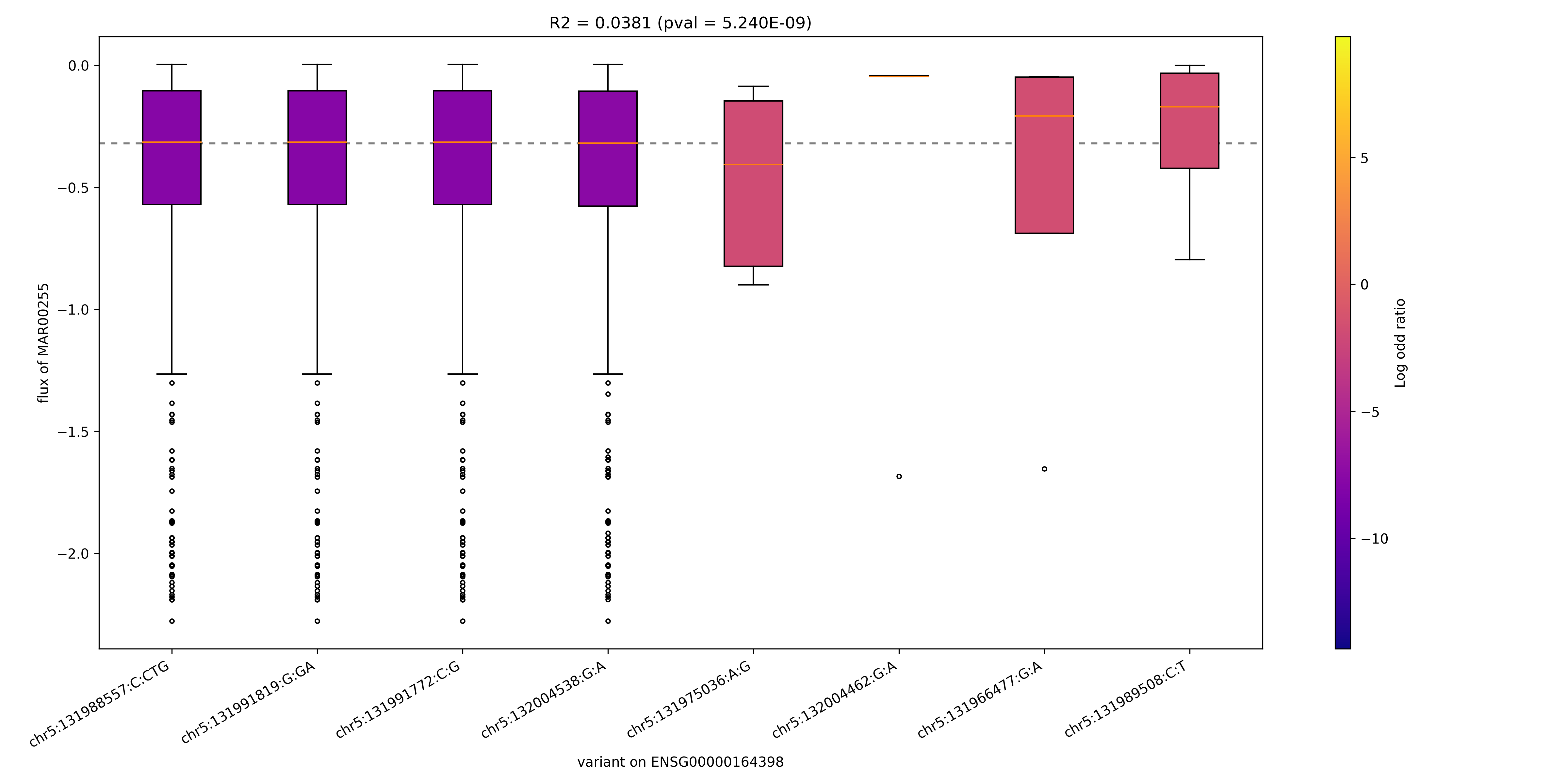

### MAR00255_ENSG00000239642.png

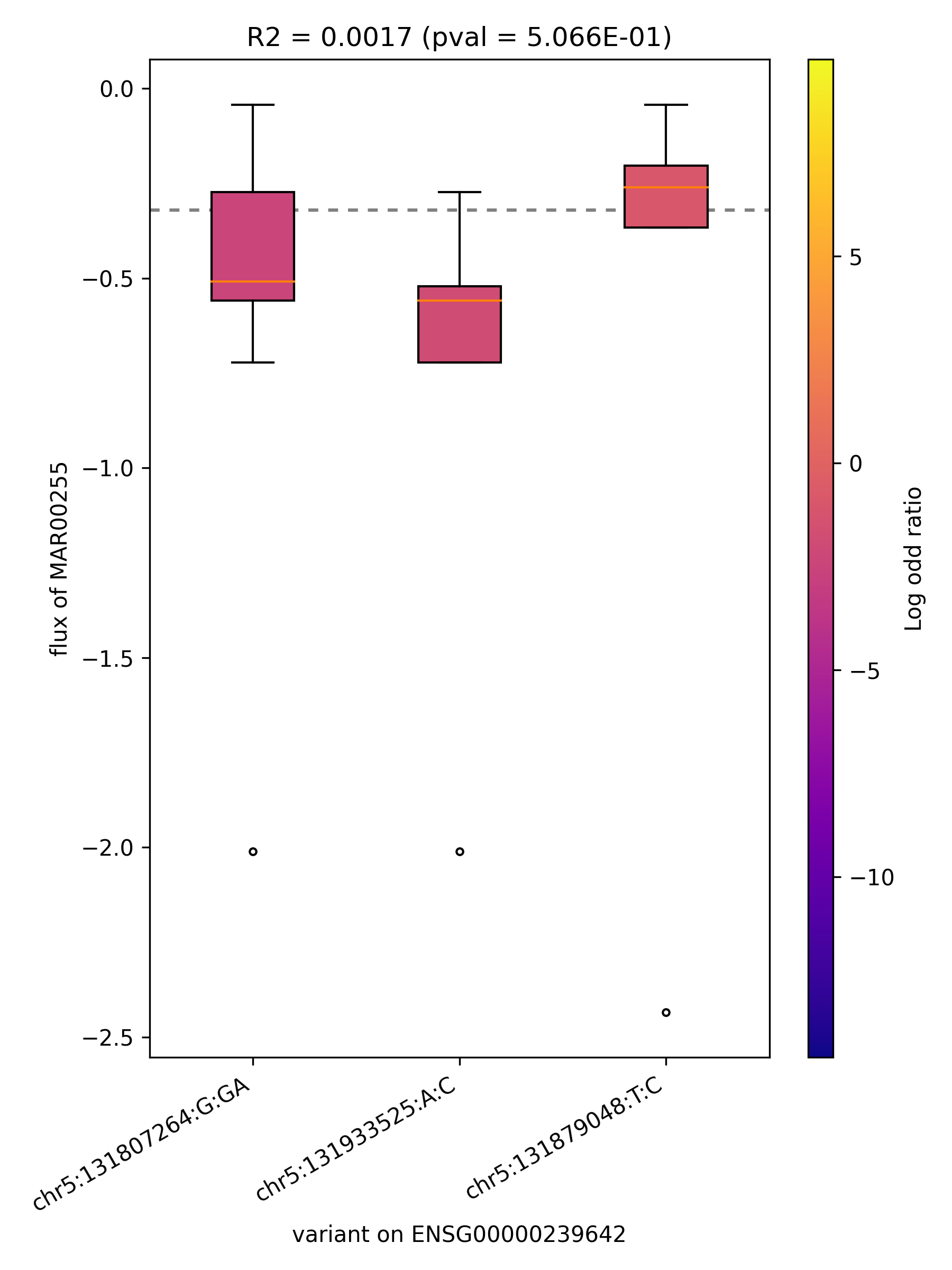

### MAR00267_ENSG00000103740.png

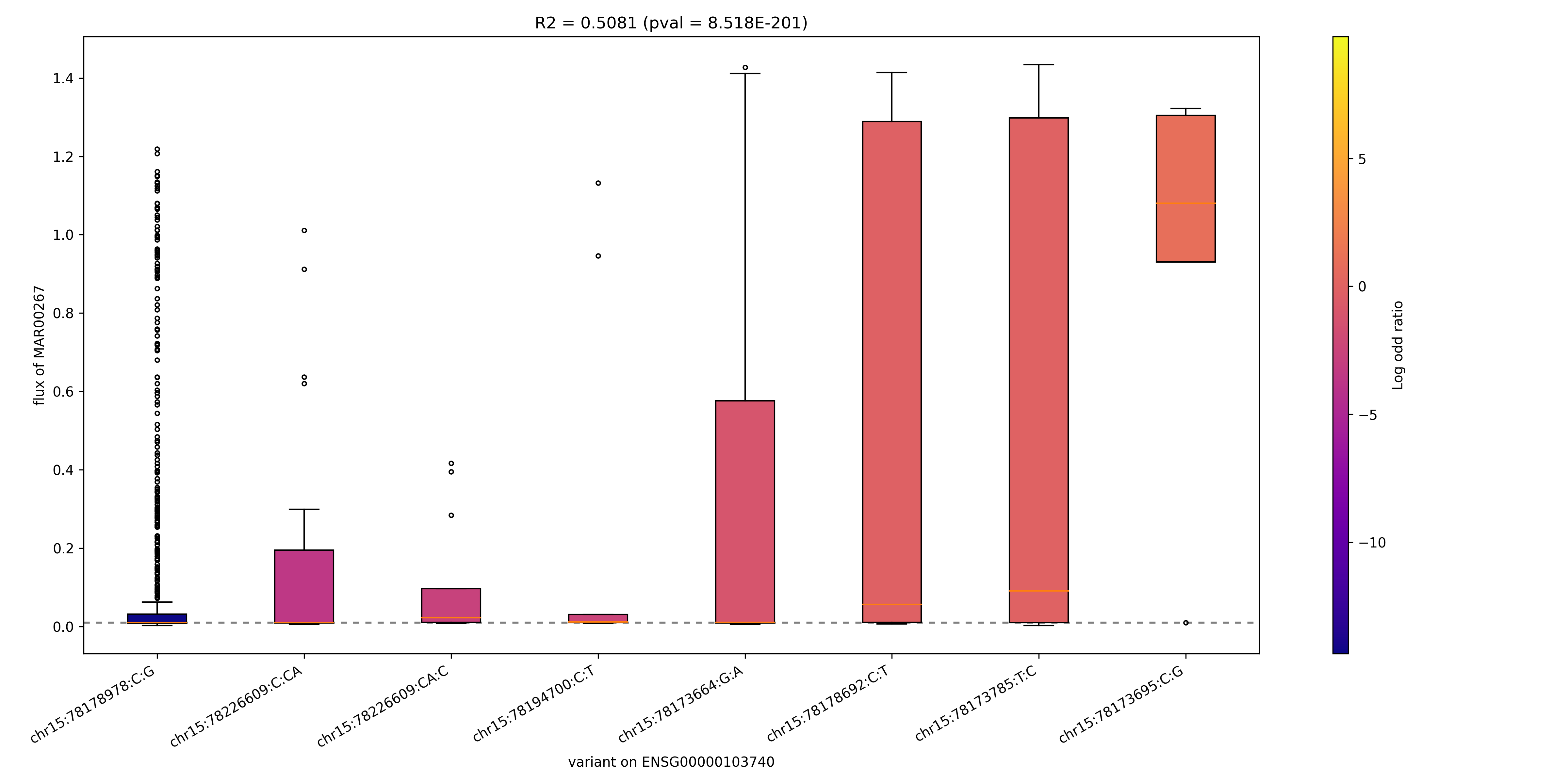

### MAR00267_ENSG00000130377.png

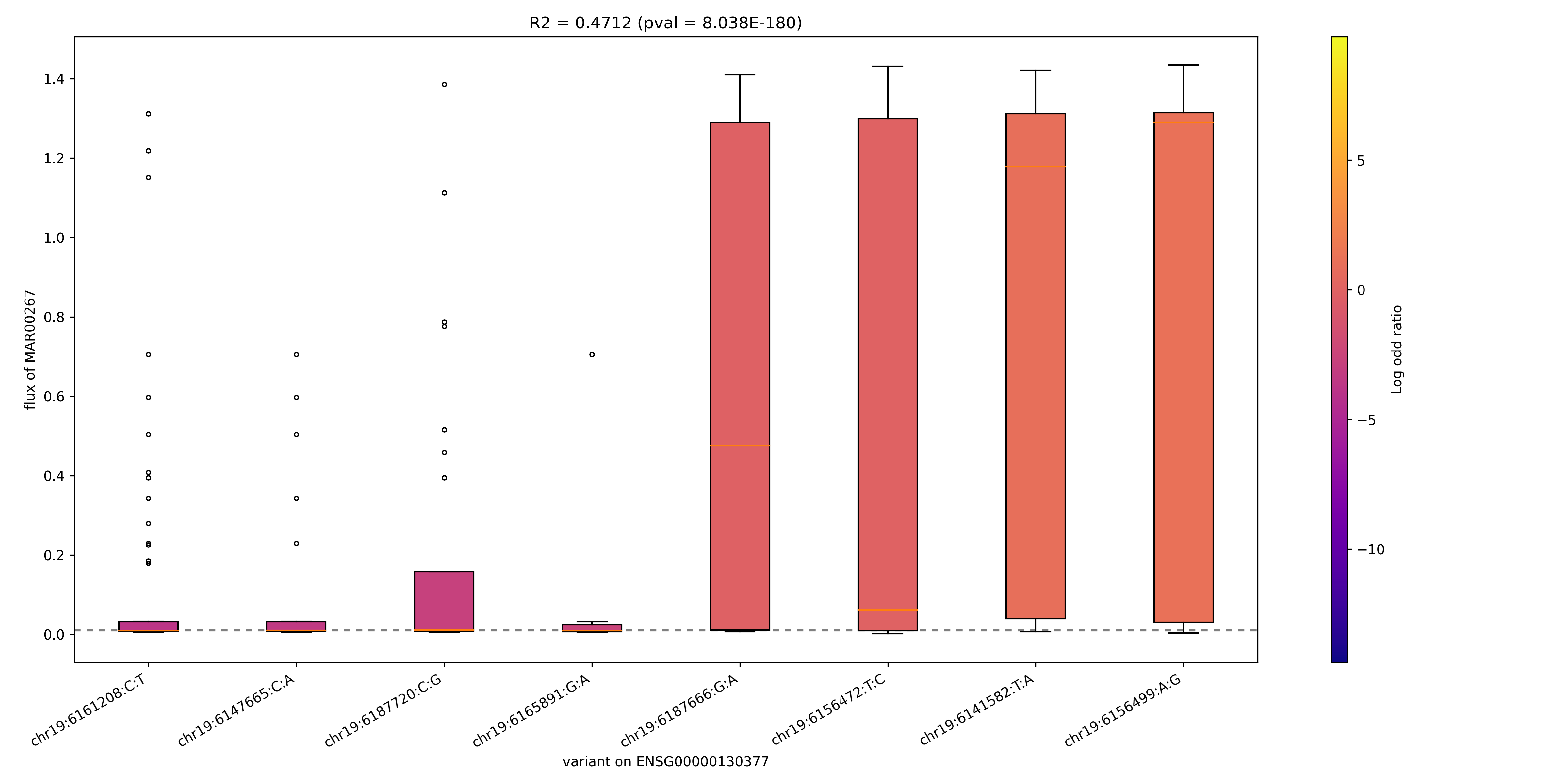

### MAR00267_ENSG00000140284.png

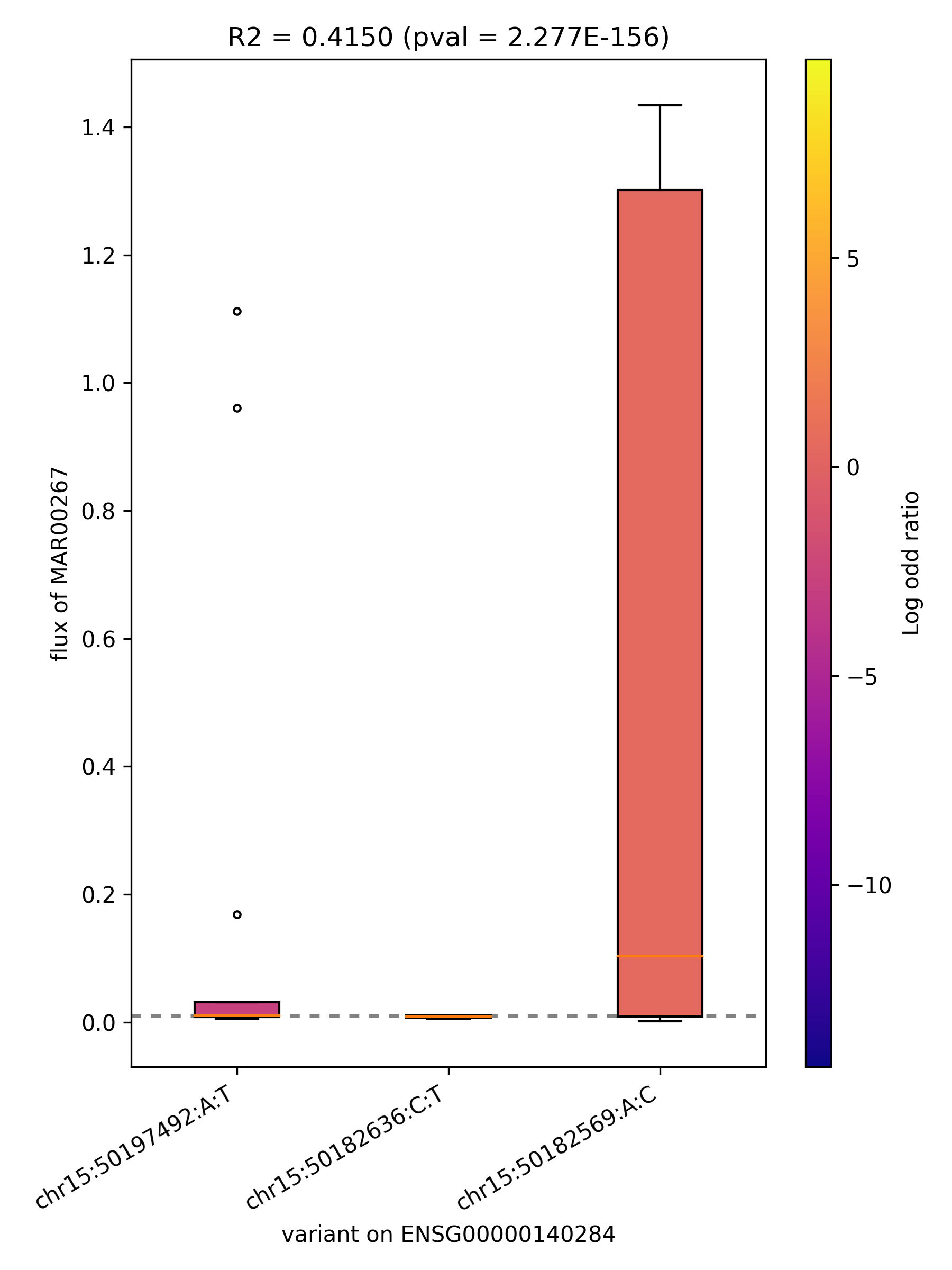

### MAR00267_ENSG00000197142.png

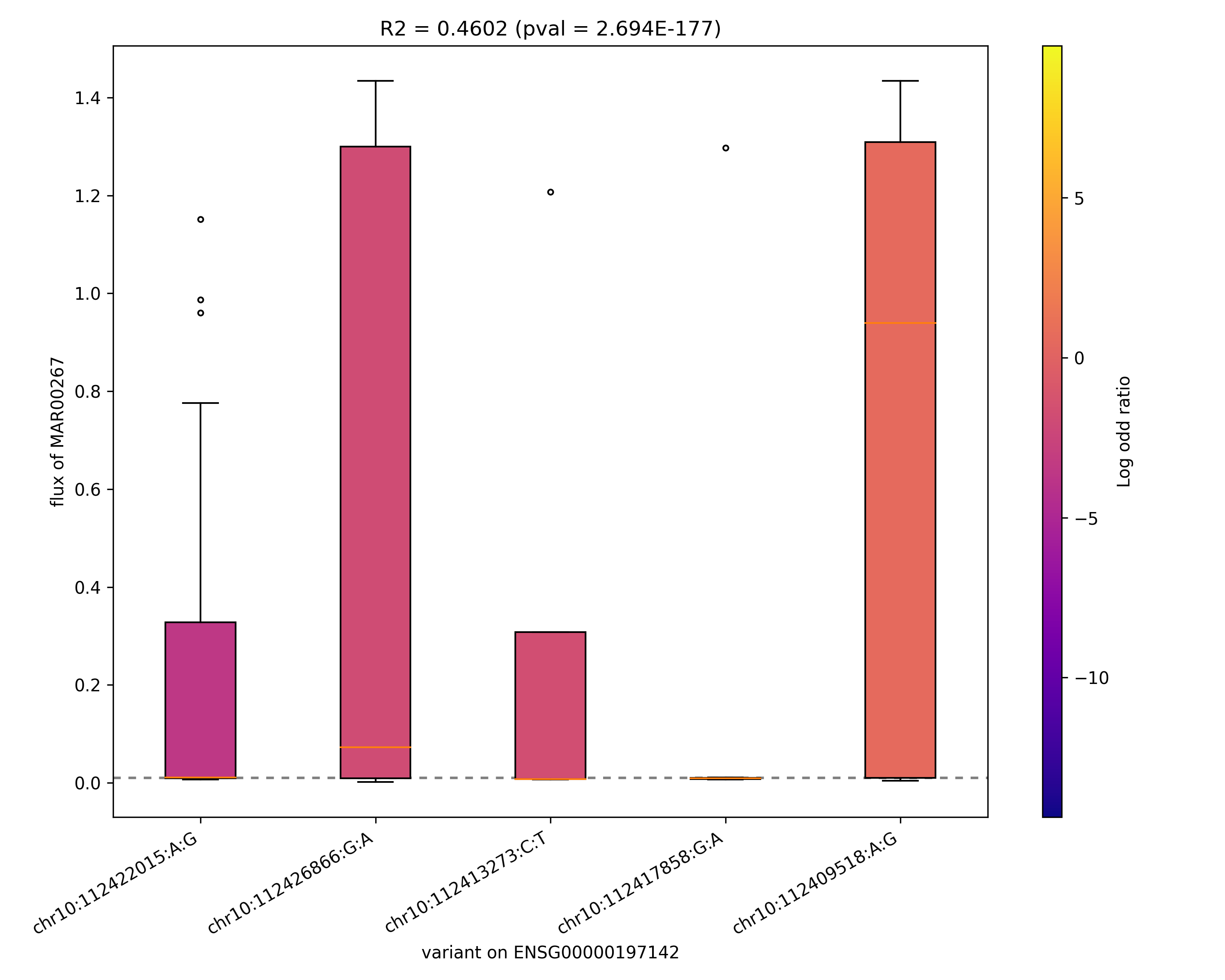

### MAR00269_ENSG00000165029.png

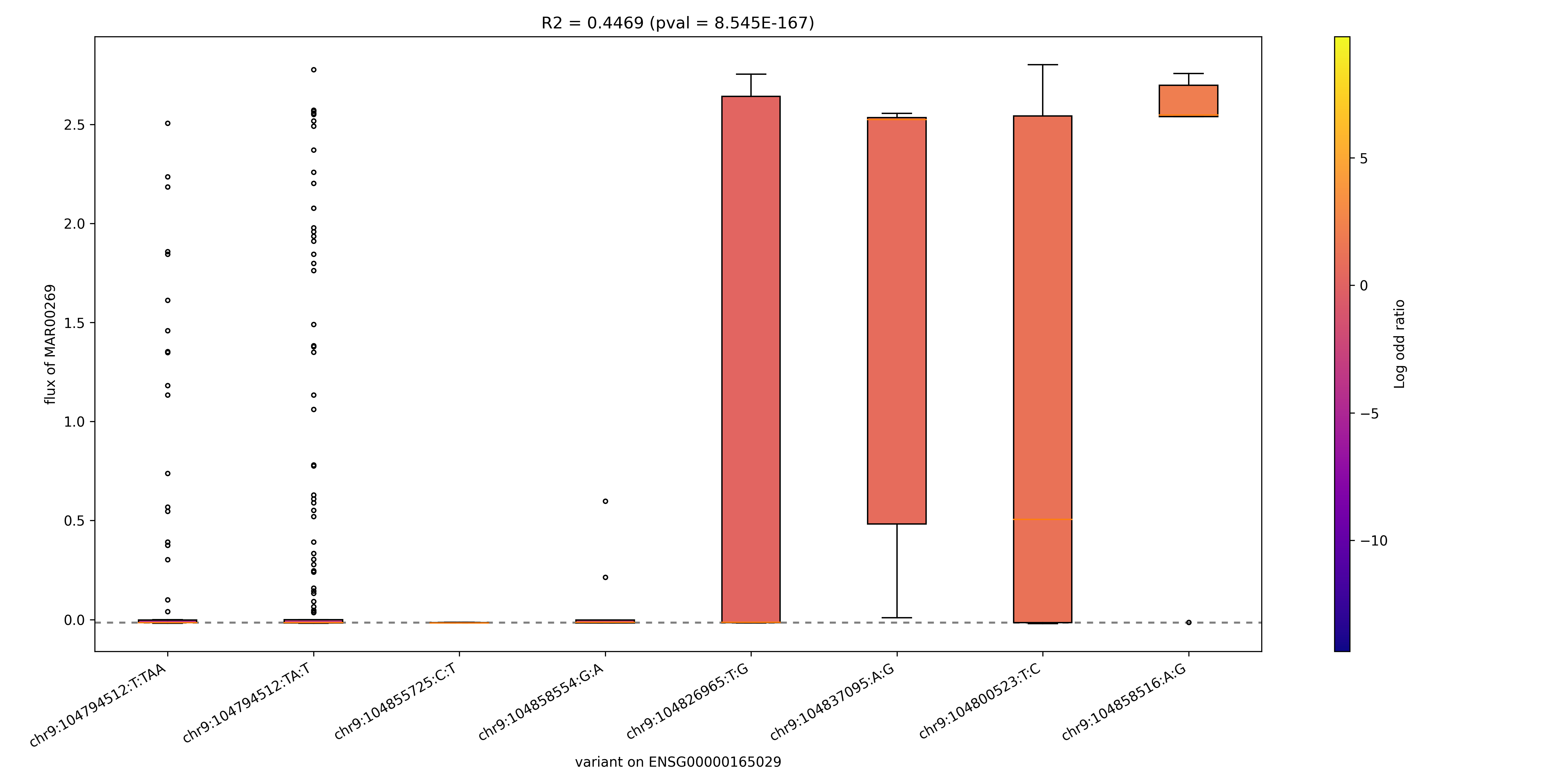

### MAR00271_ENSG00000103740.png

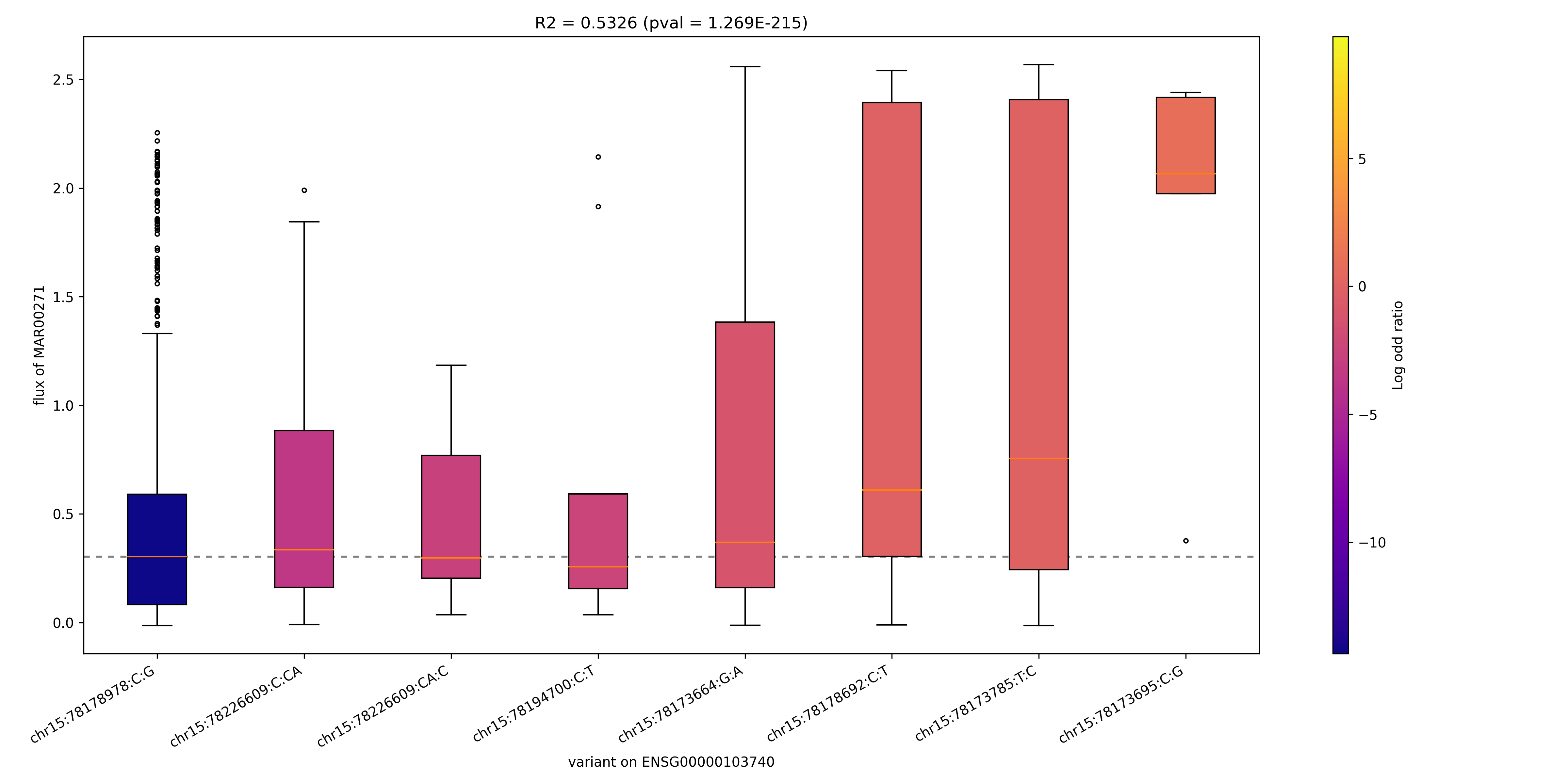

### MAR00271_ENSG00000130377.png

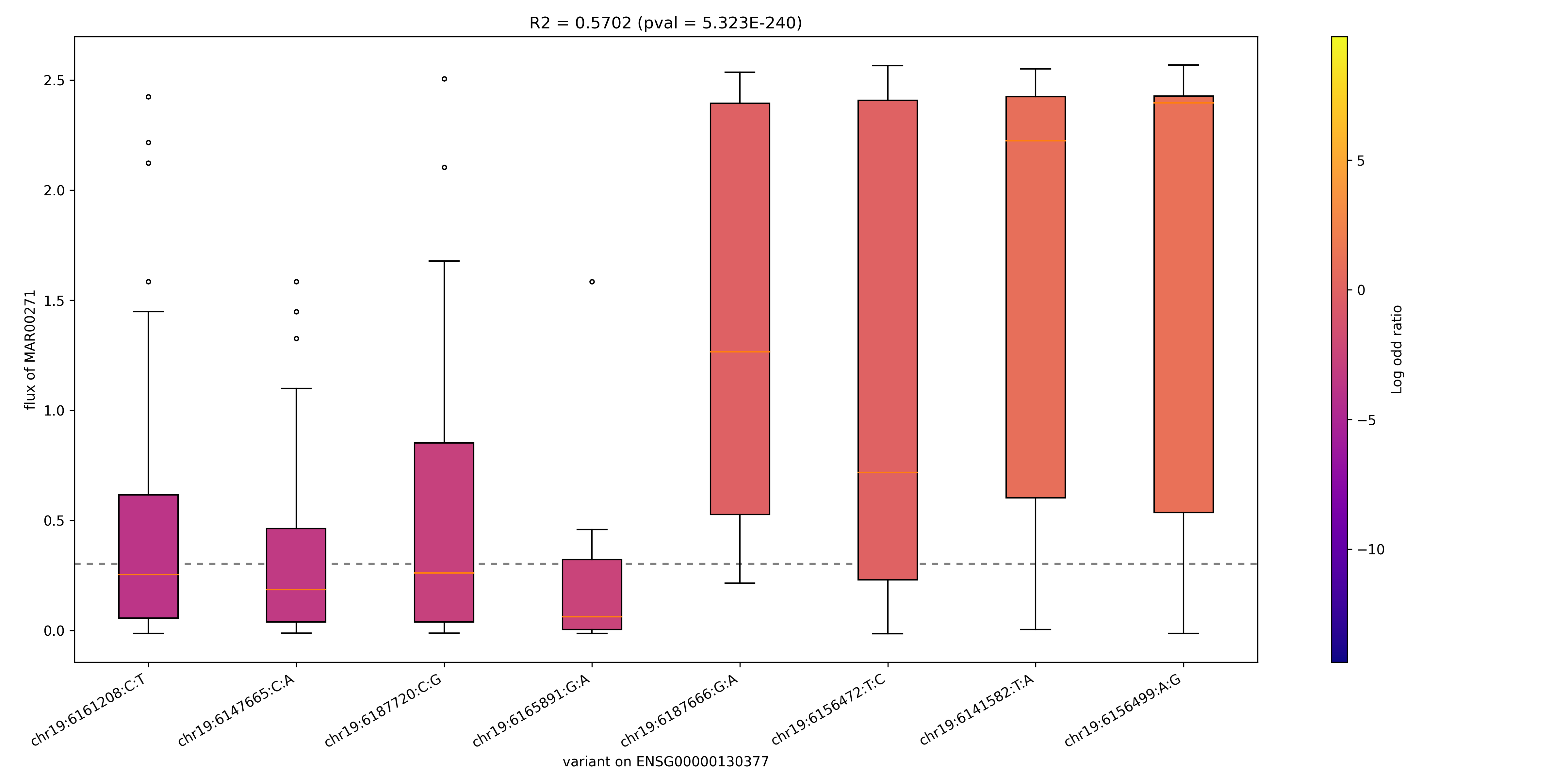

### MAR00271_ENSG00000140284.png

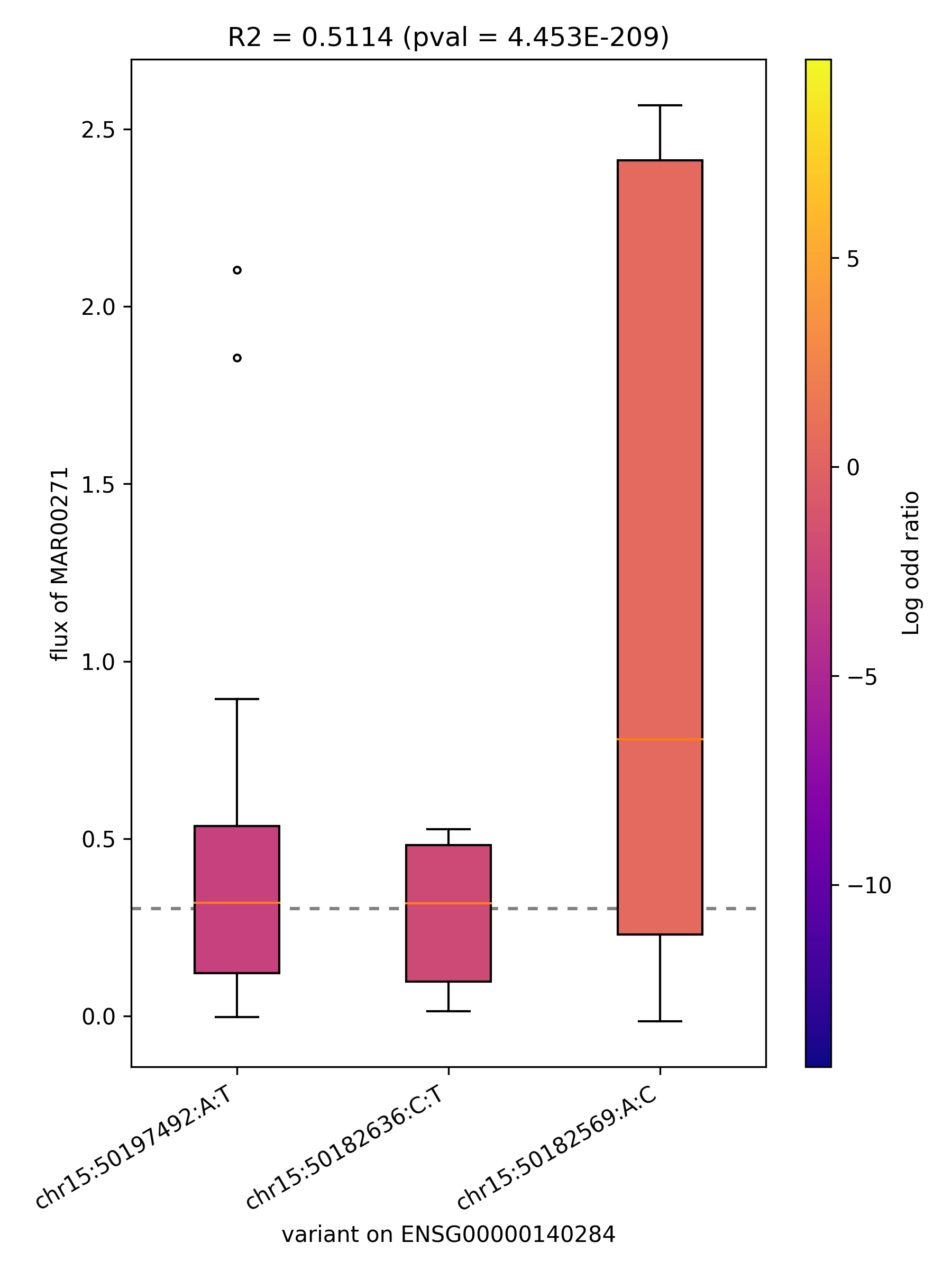

### MAR00271_ENSG00000197142.png

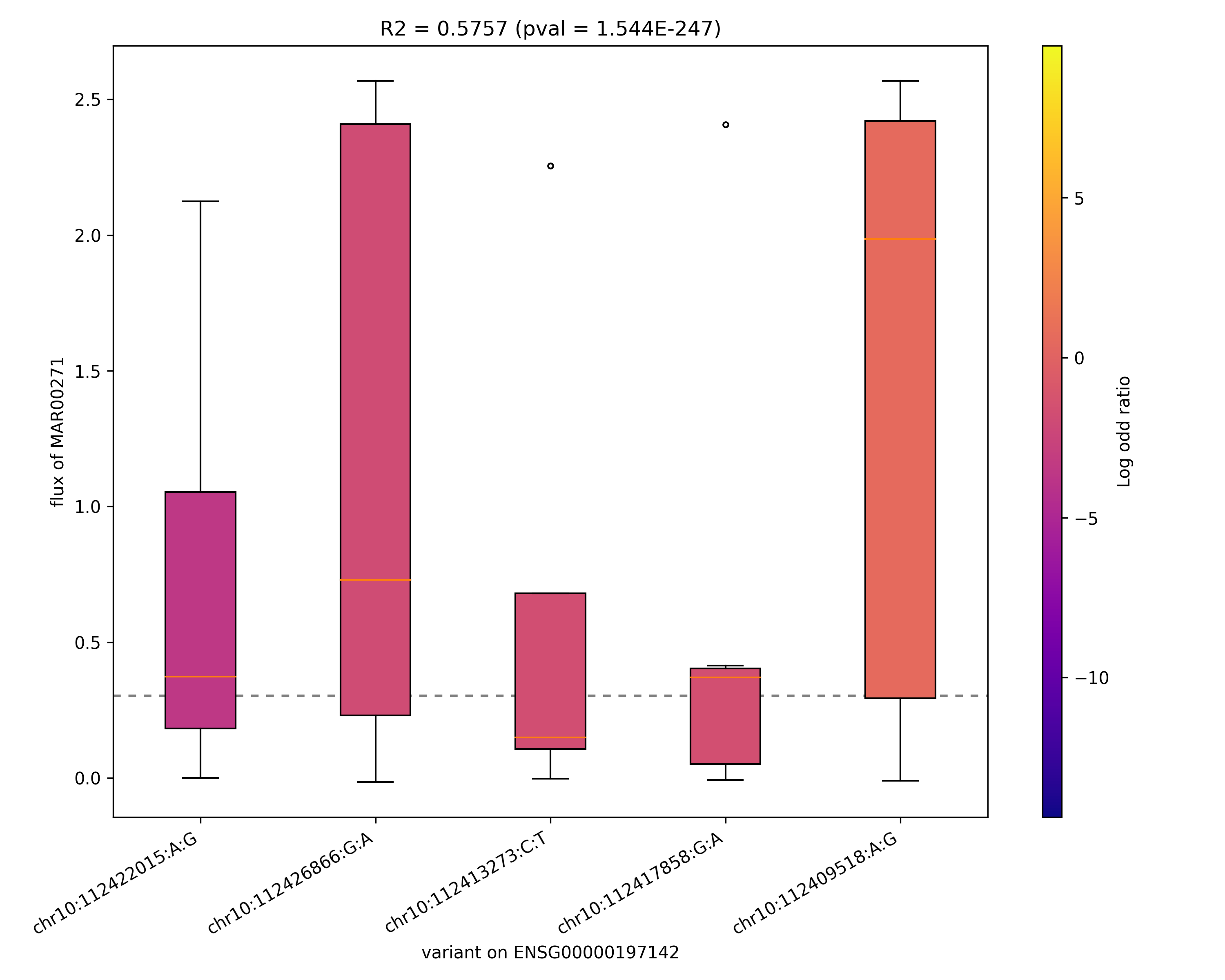

### MAR00273_ENSG00000165029.png

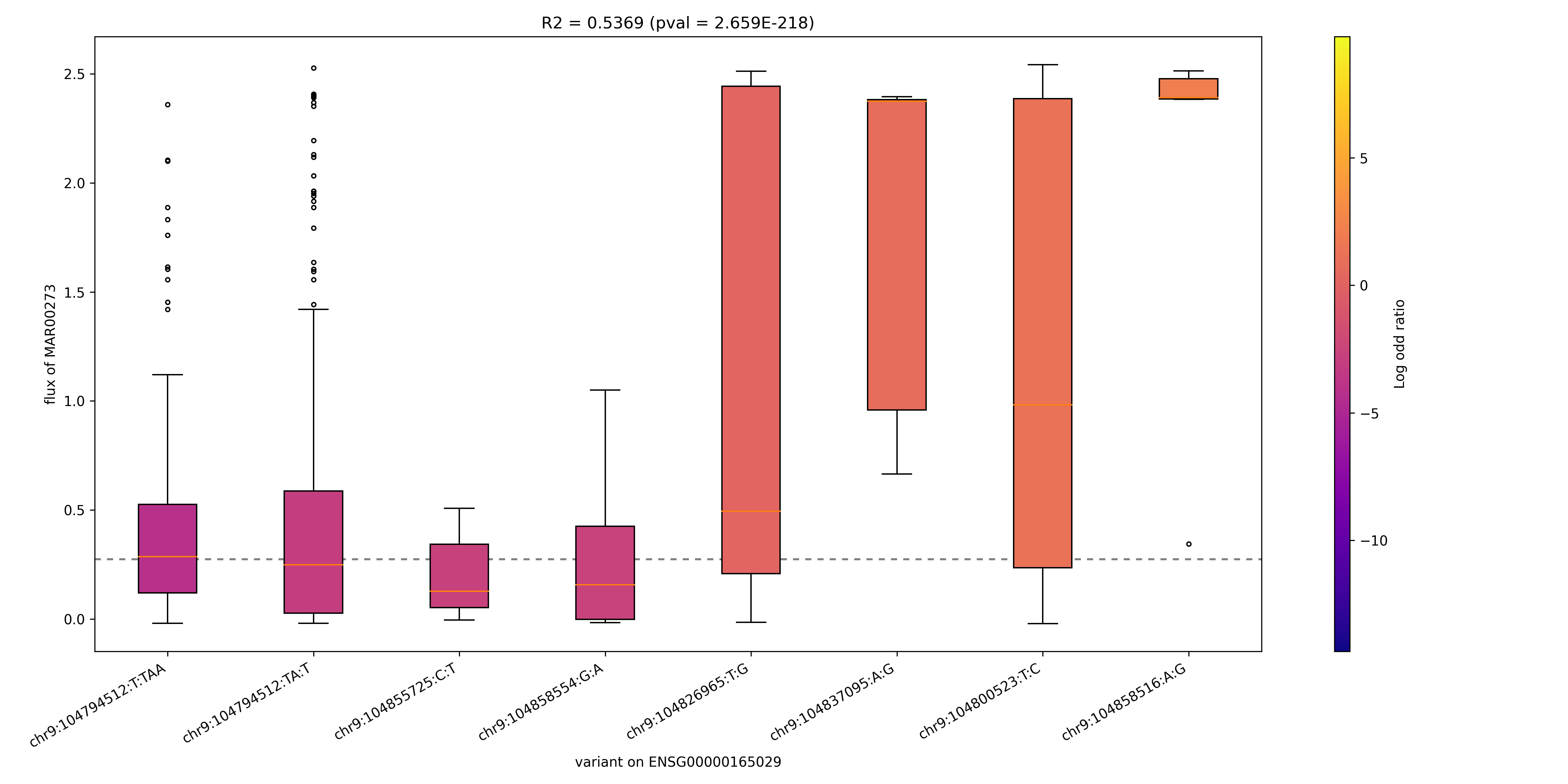

### MAR00452_ENSG00000110090.png

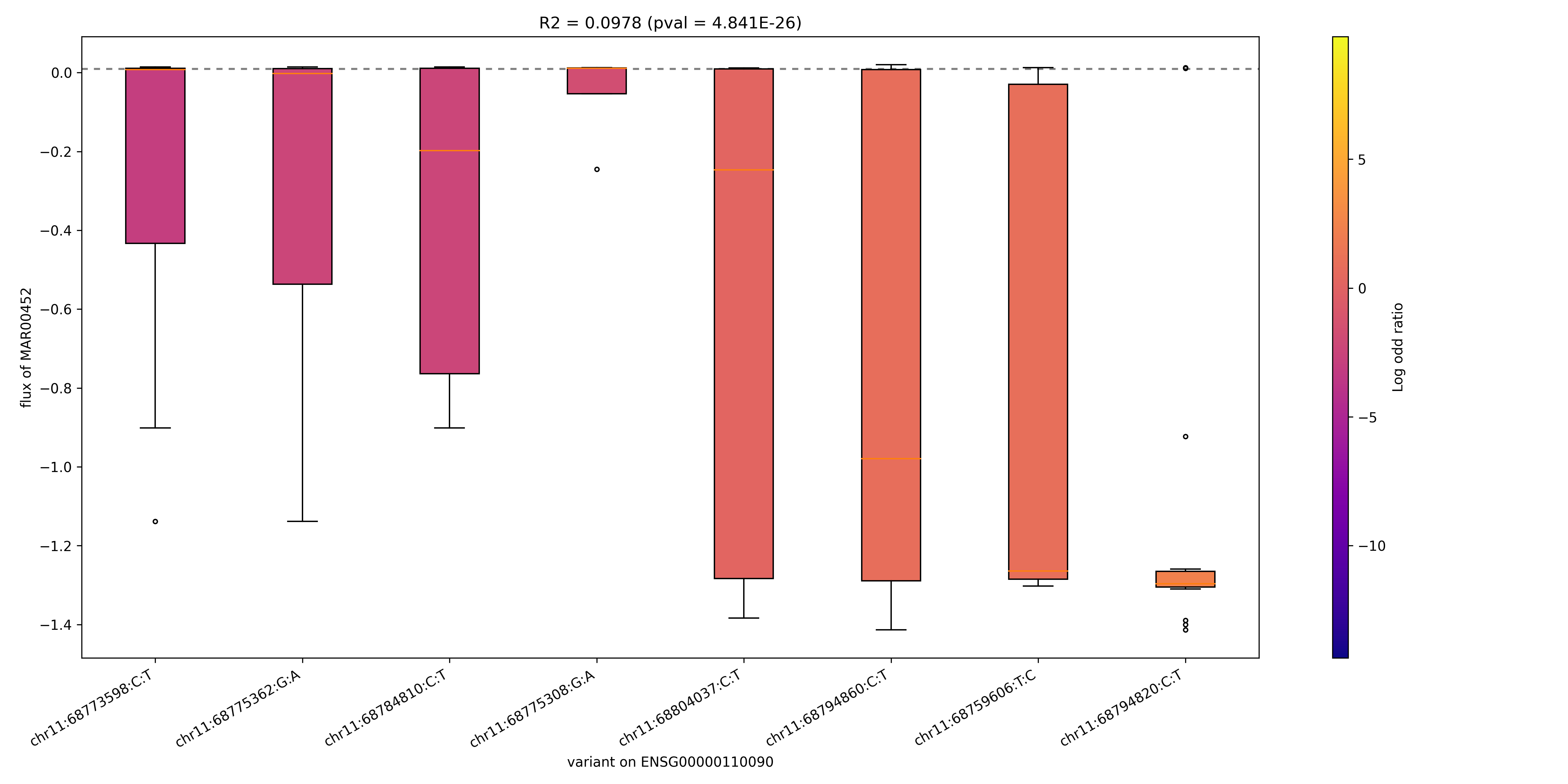

### MAR00452_ENSG00000169169.png

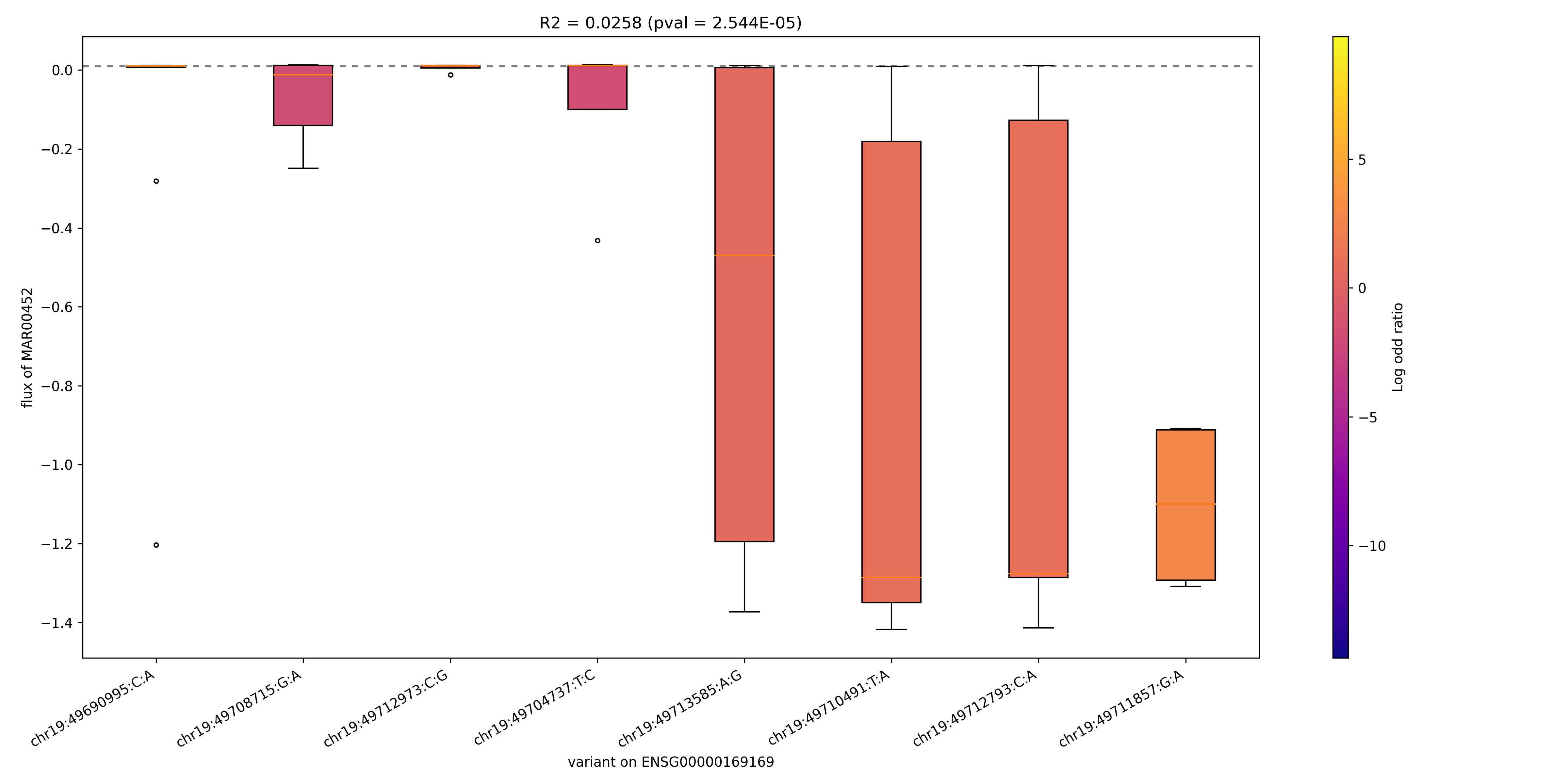

### MAR00742_ENSG00000166743.png

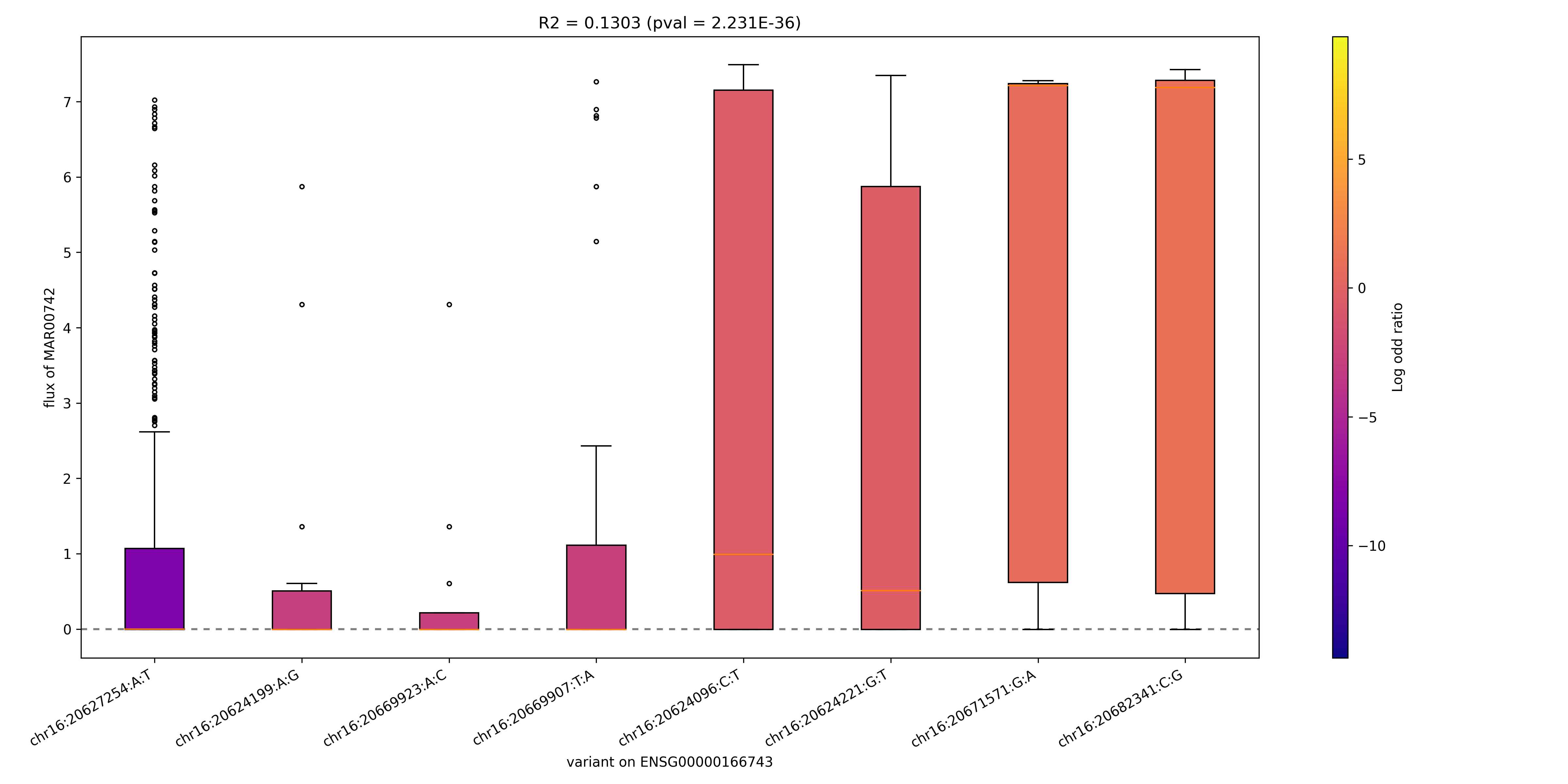

### MAR01231_ENSG00000159228.png

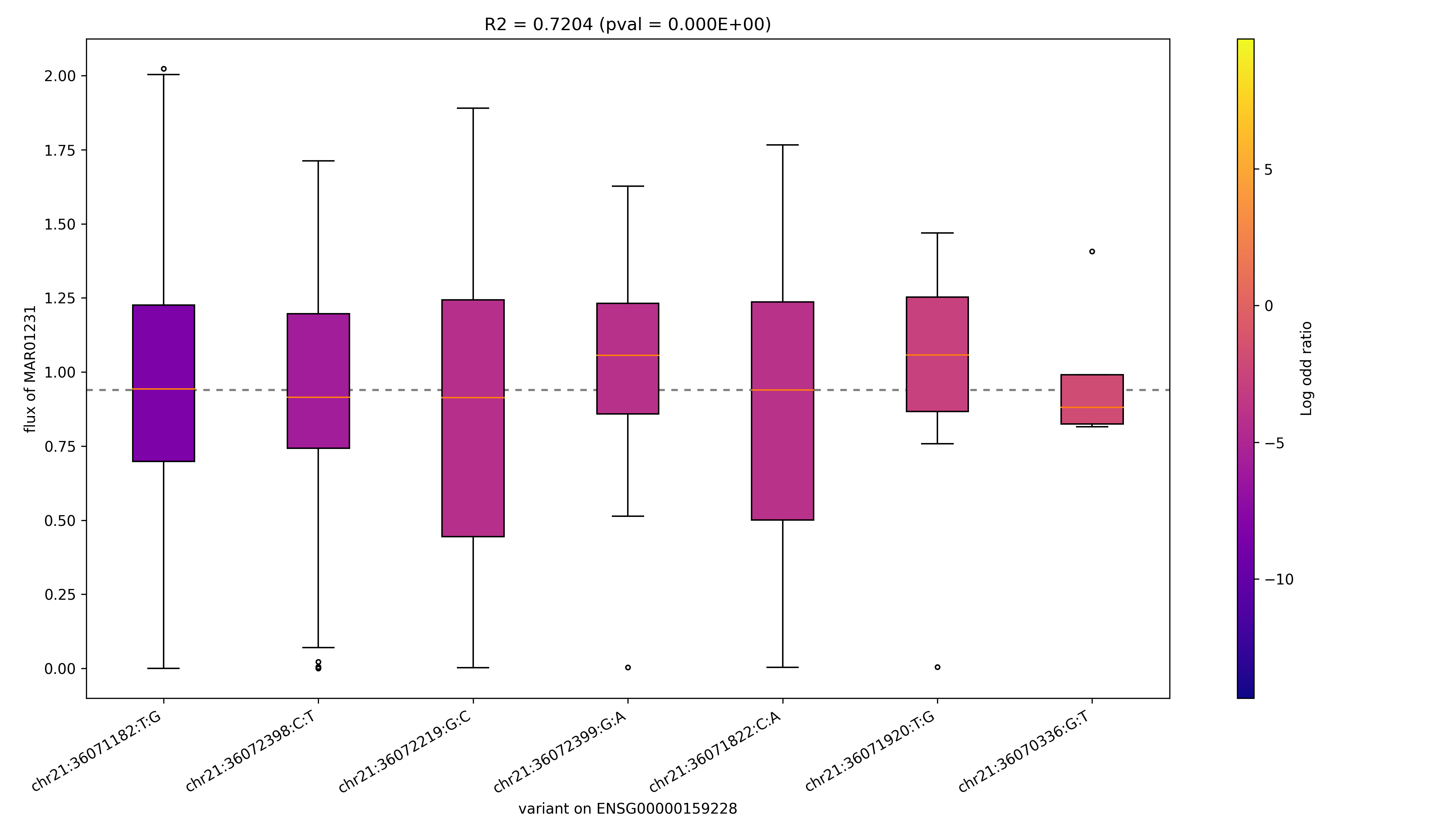

### MAR01263_ENSG00000103740.png

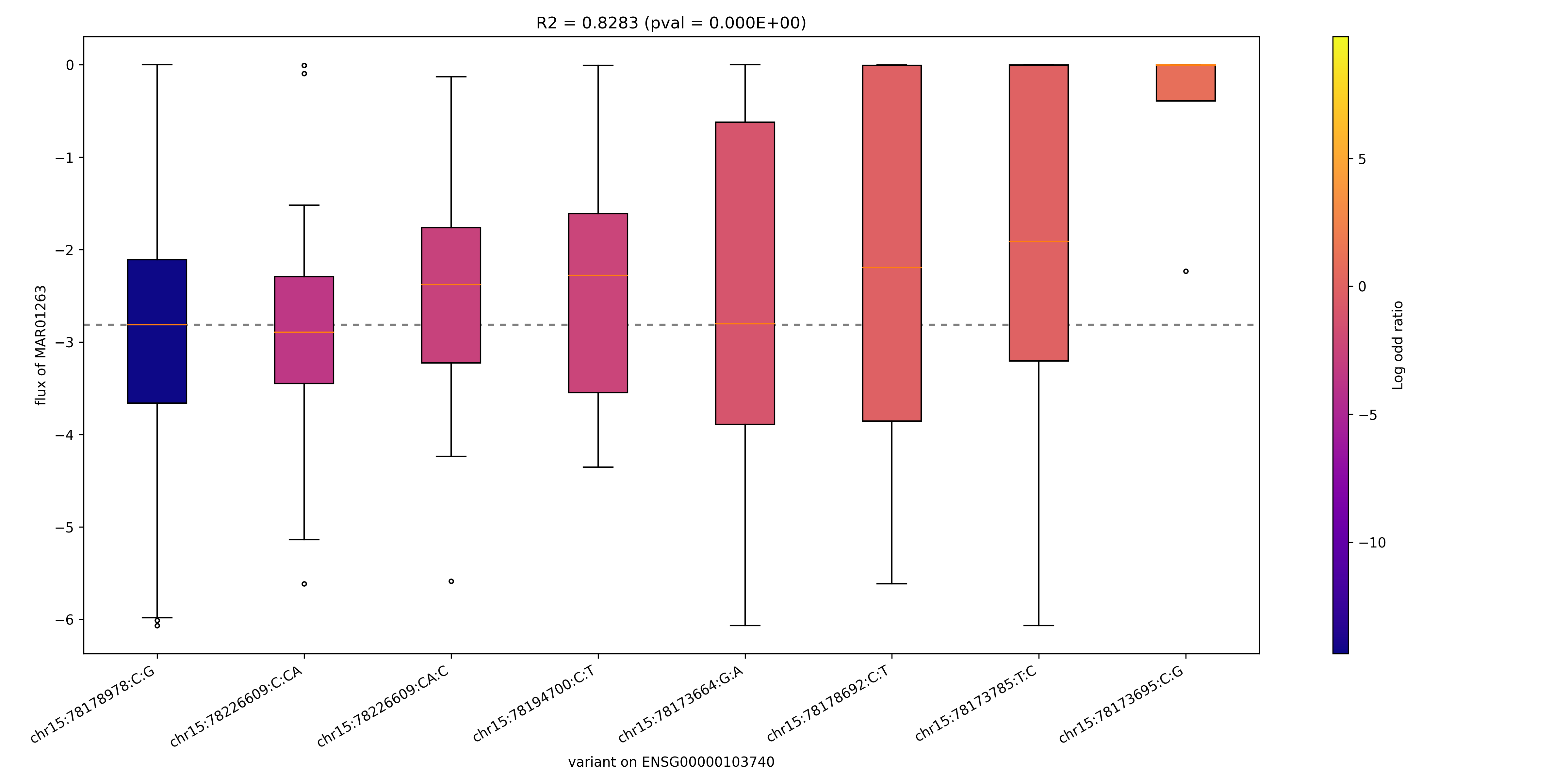

### MAR01263_ENSG00000130377.png

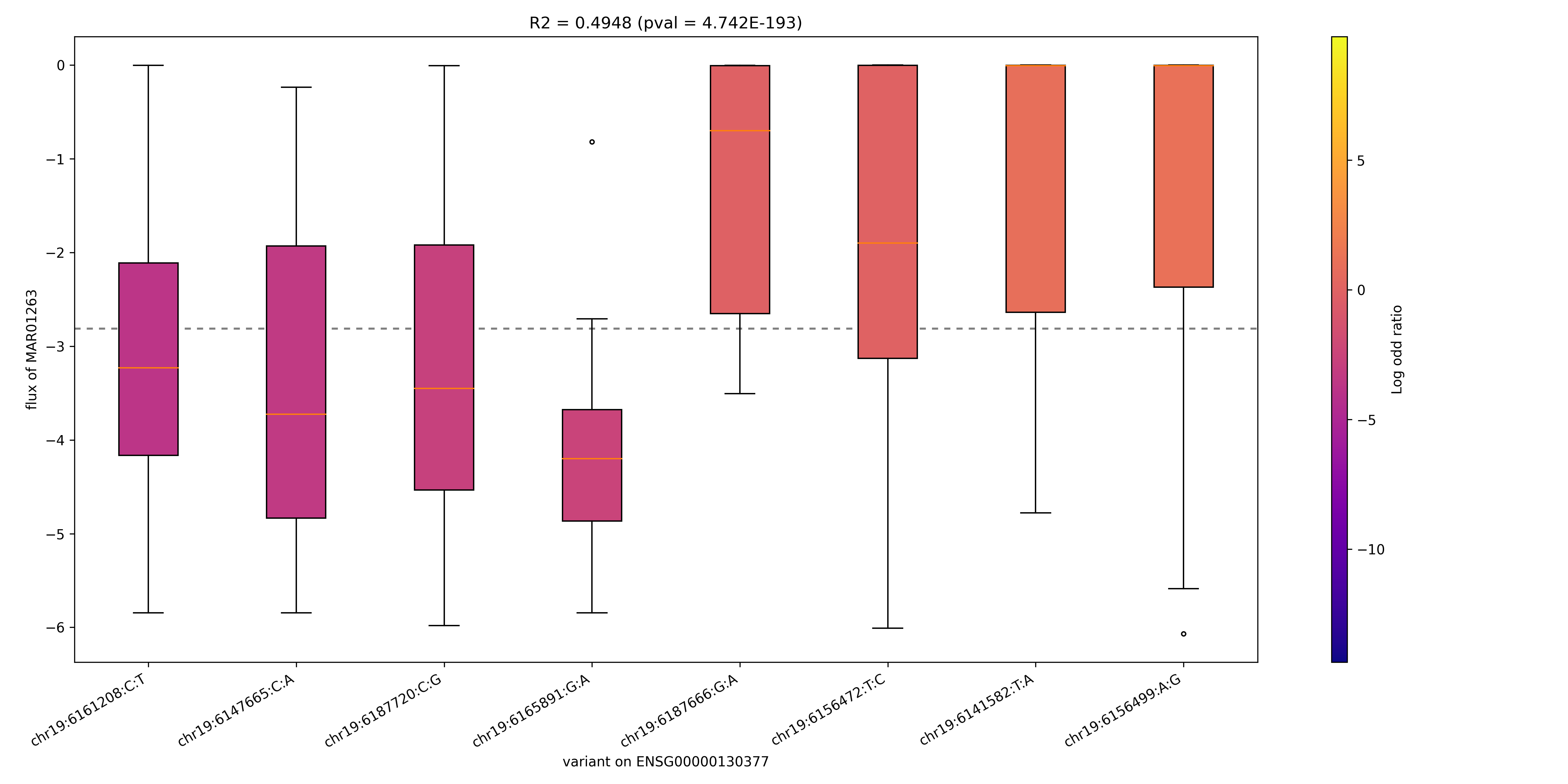

### MAR01263_ENSG00000140284.png

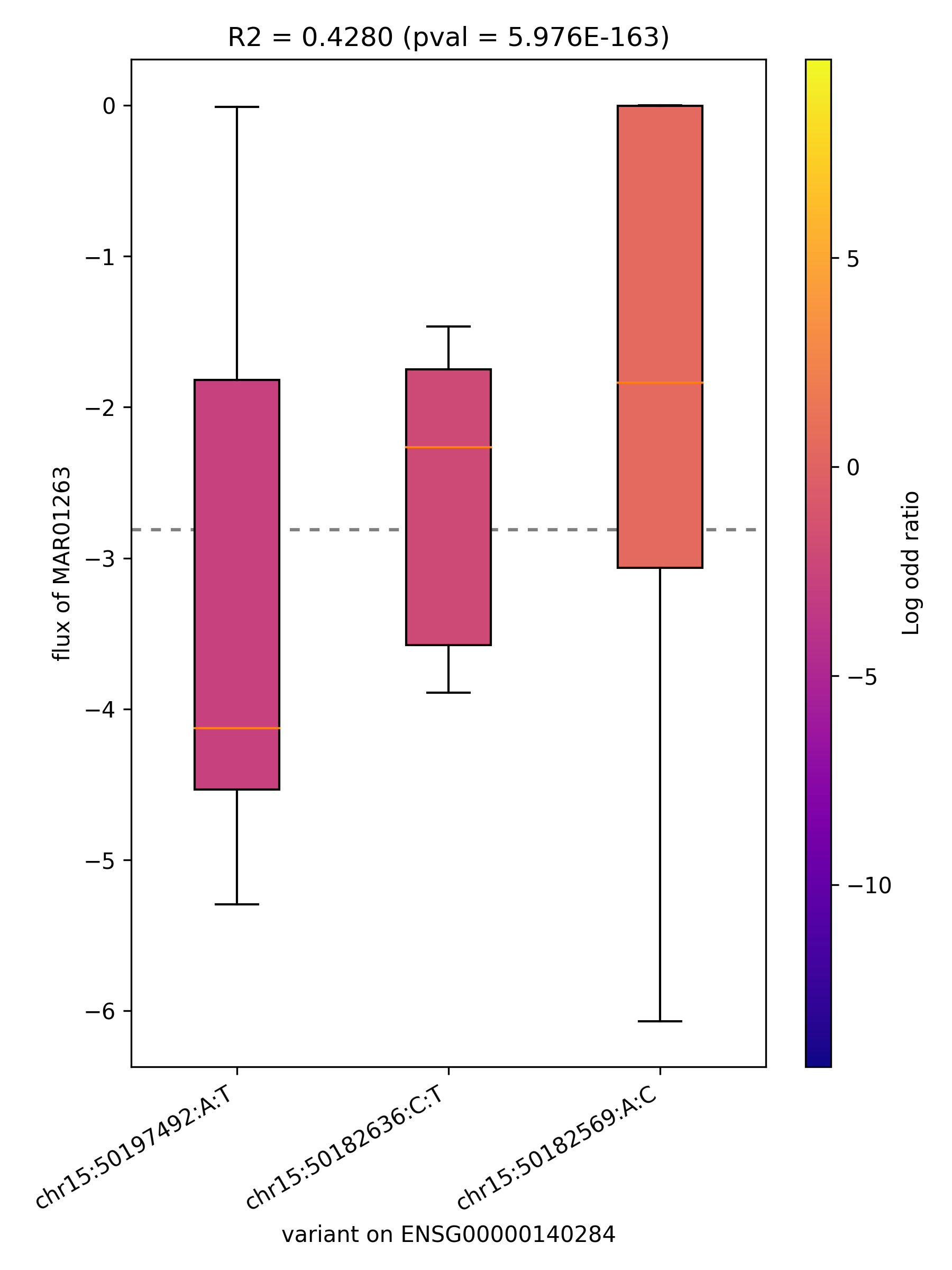

### MAR01263_ENSG00000197142.png

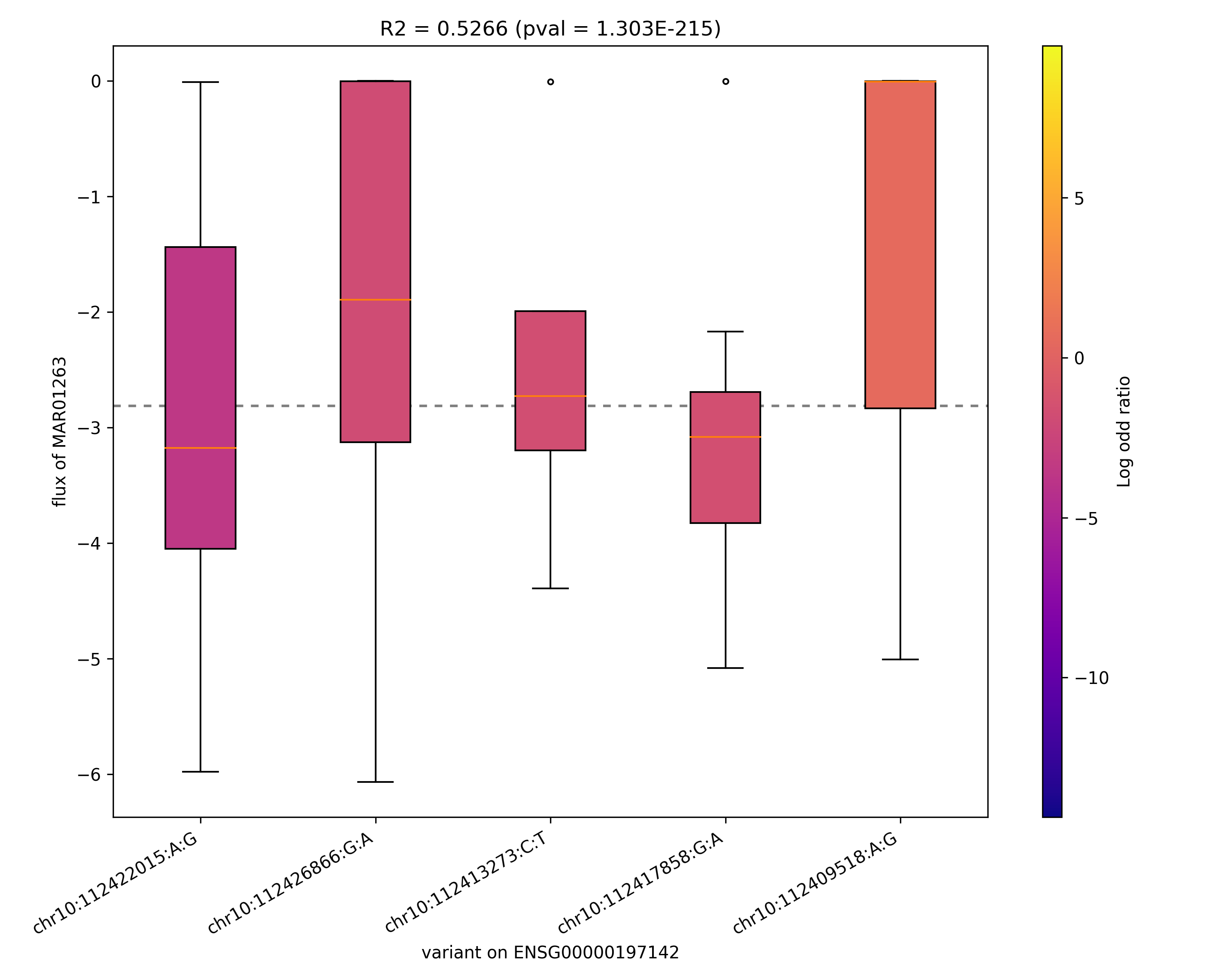

### MAR01280_ENSG00000159228.png

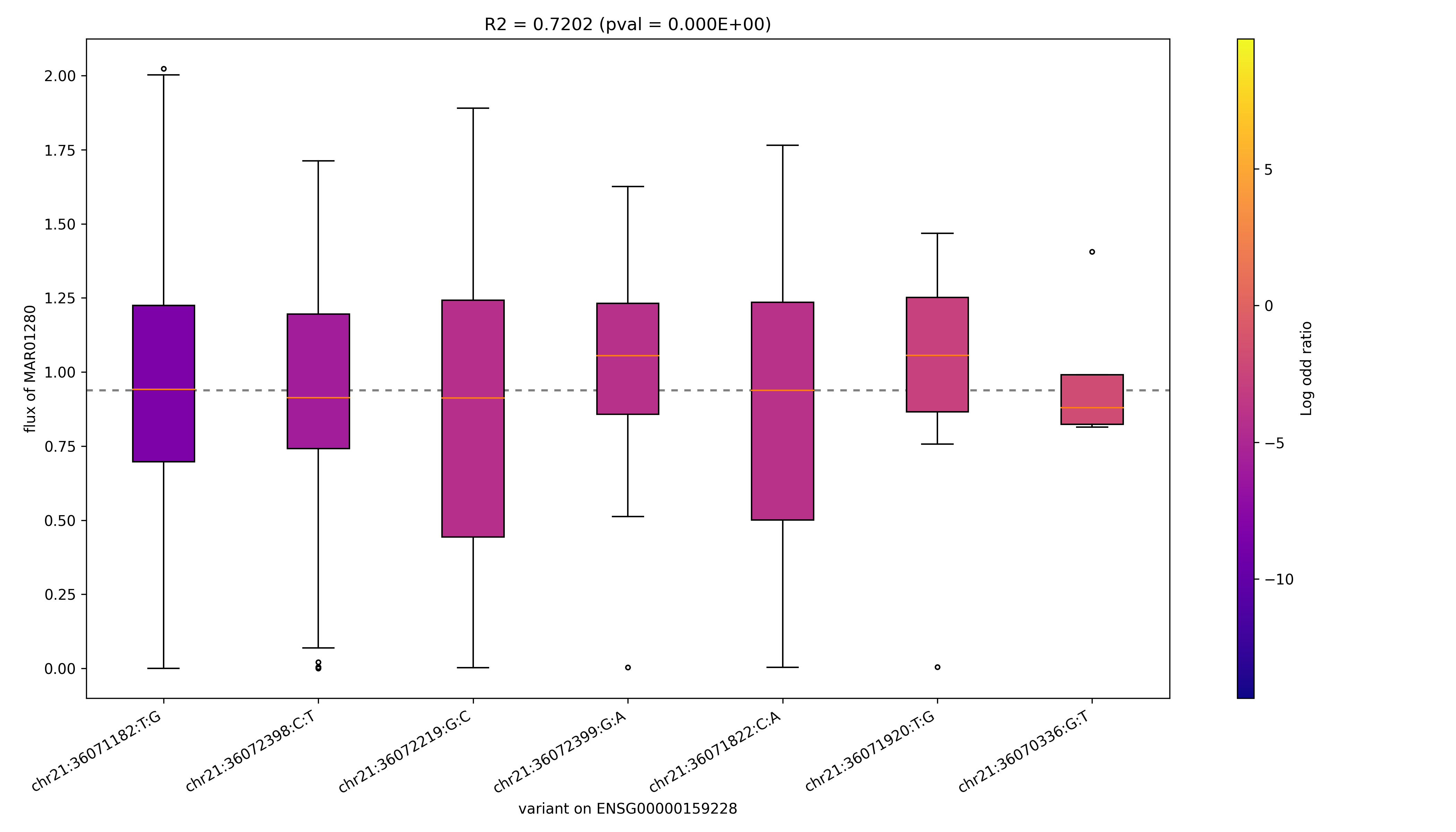

### MAR01812_ENSG00000118094.png

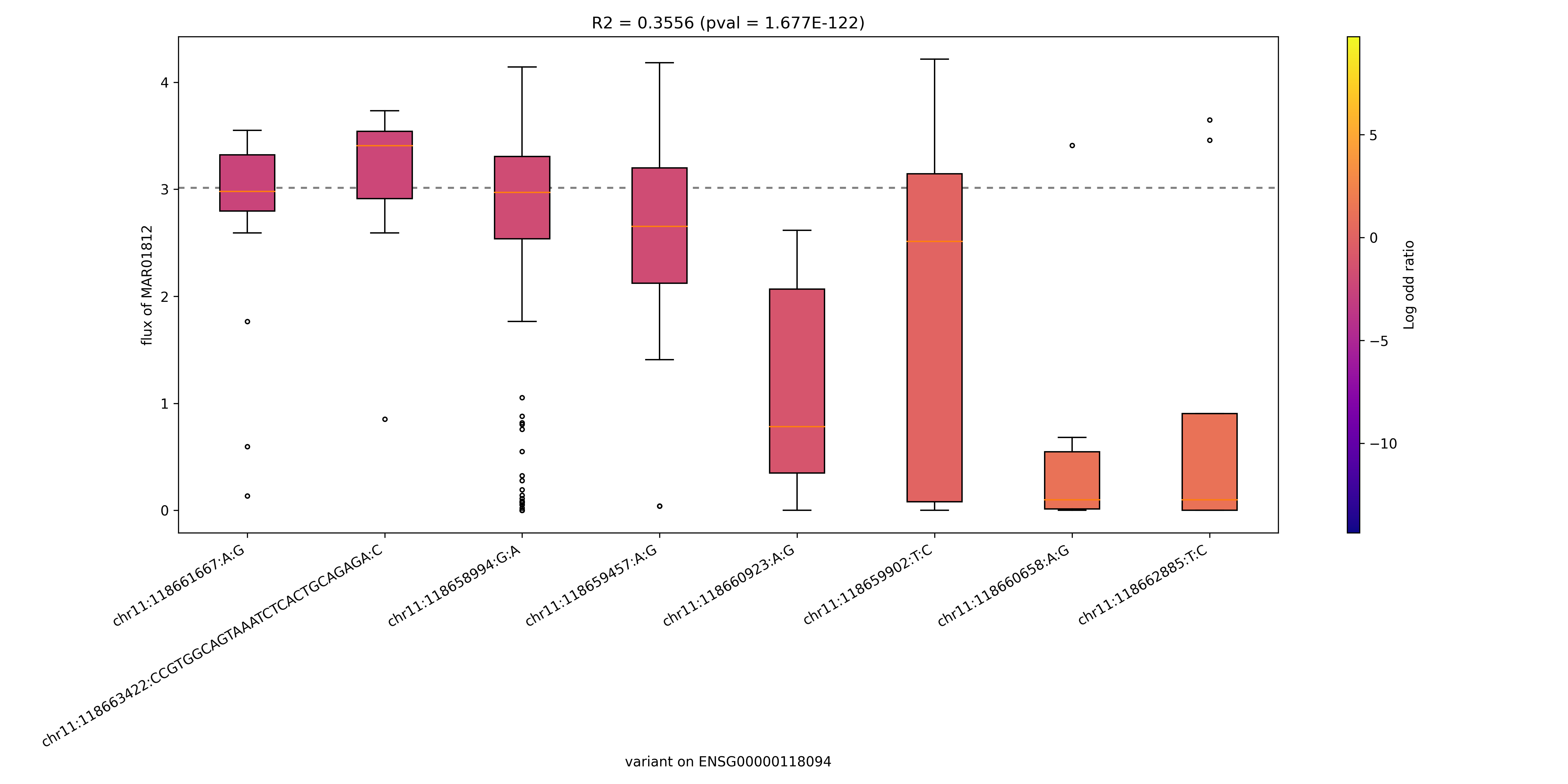

### MAR01812_ENSG00000142102.png

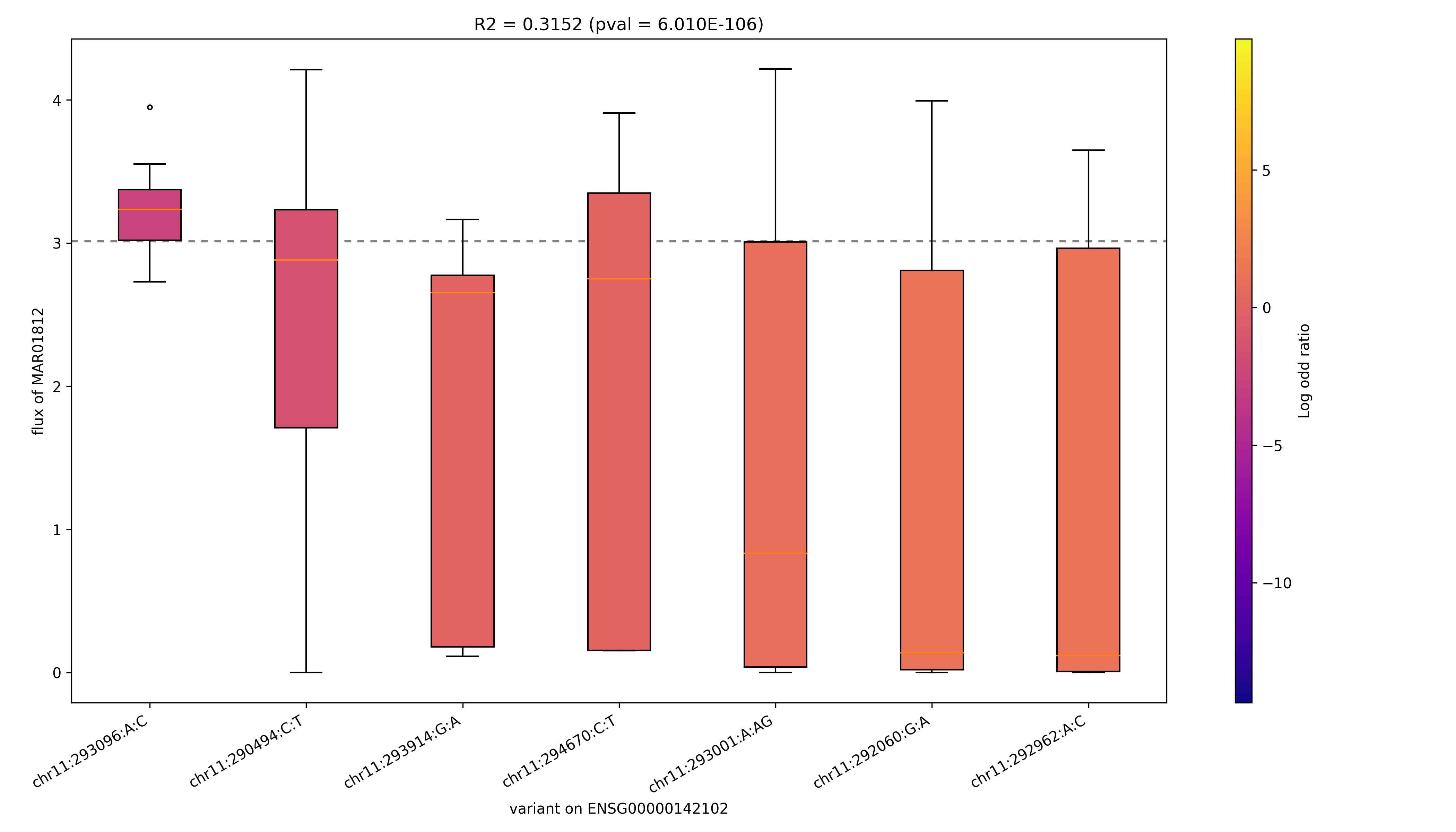

### MAR02611_ENSG00000110090.png

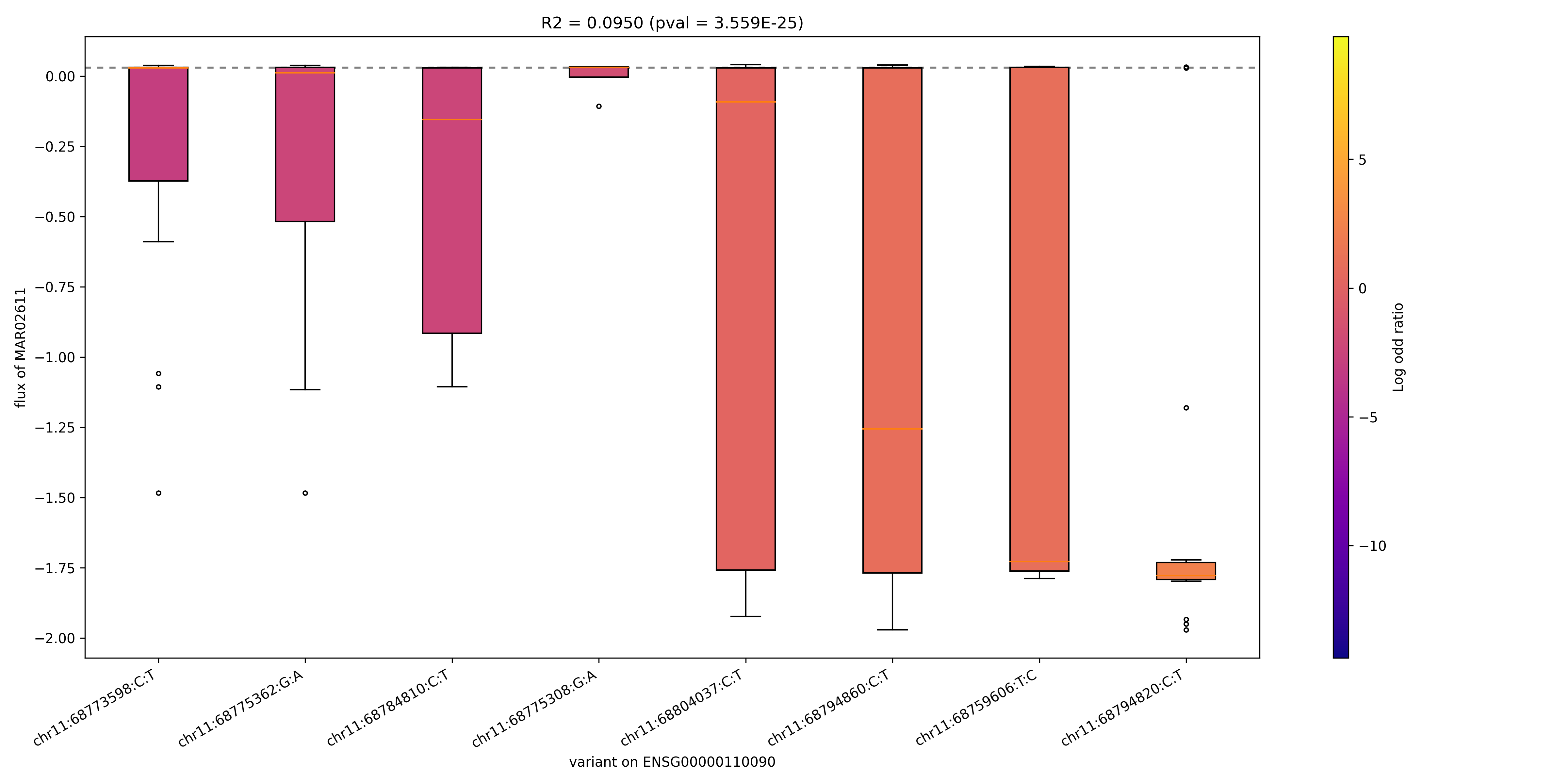

### MAR02611_ENSG00000169169.png

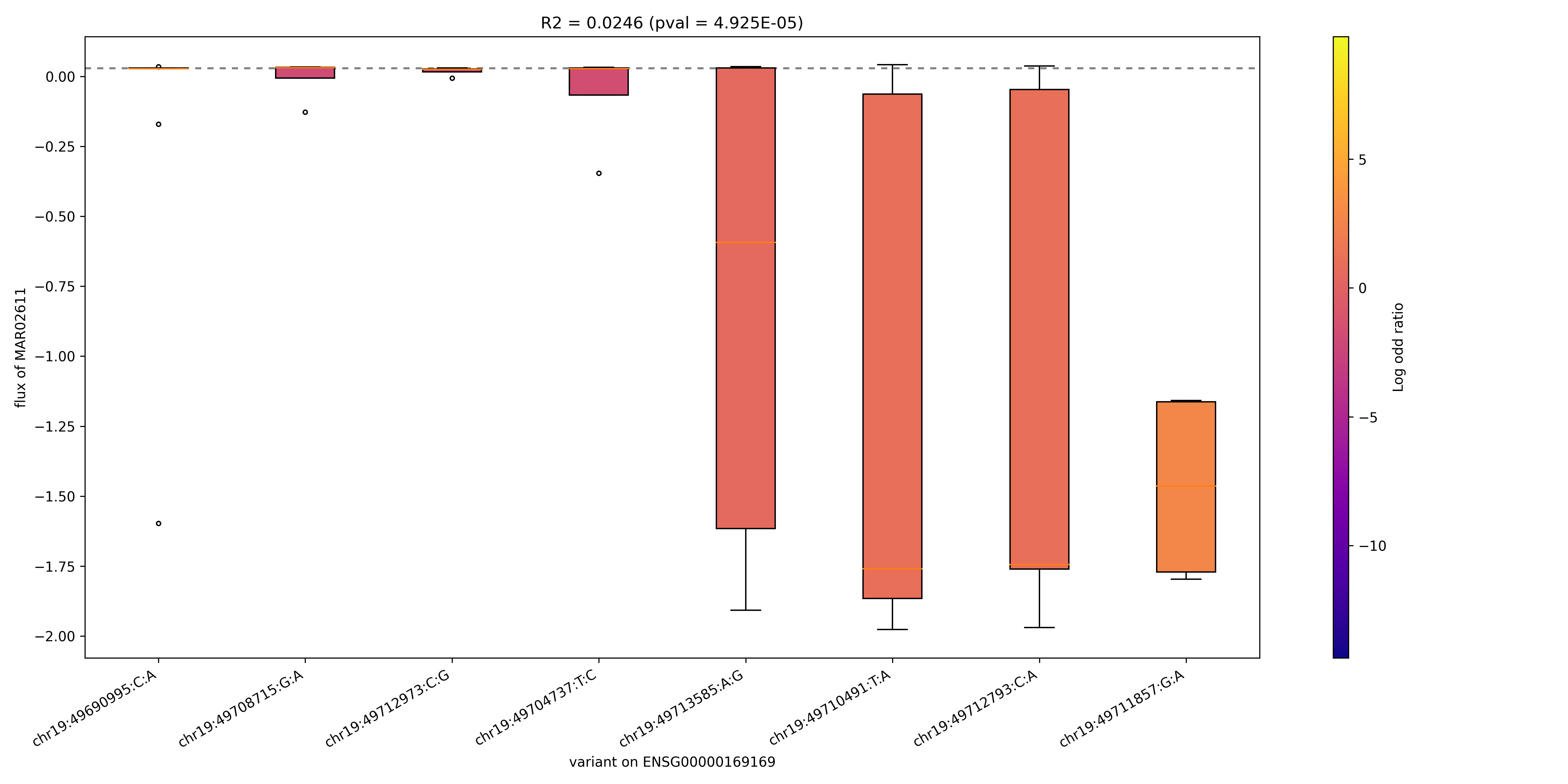

### MAR02633_ENSG00000205560.png

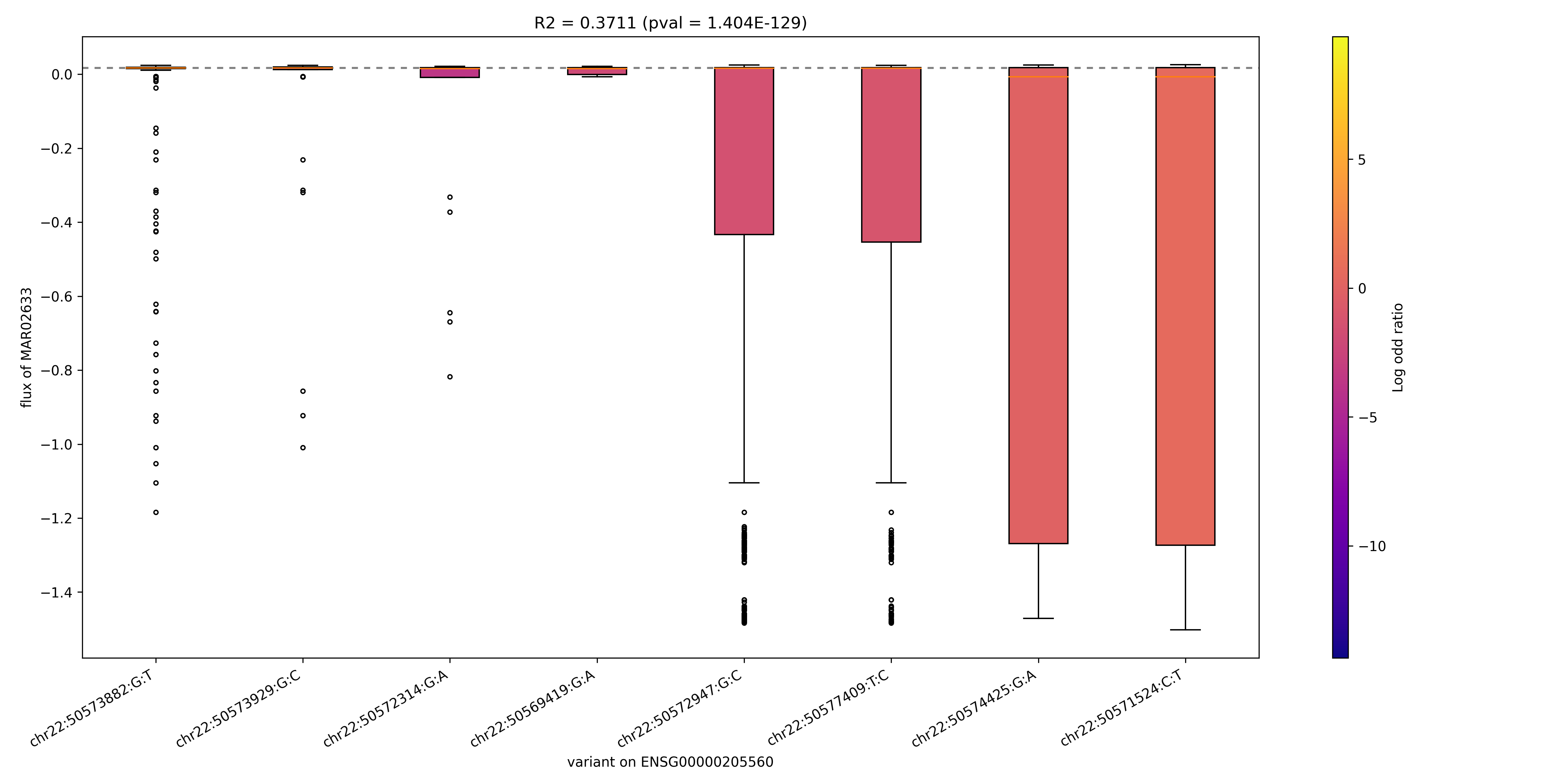

### MAR02640_ENSG00000157184.png

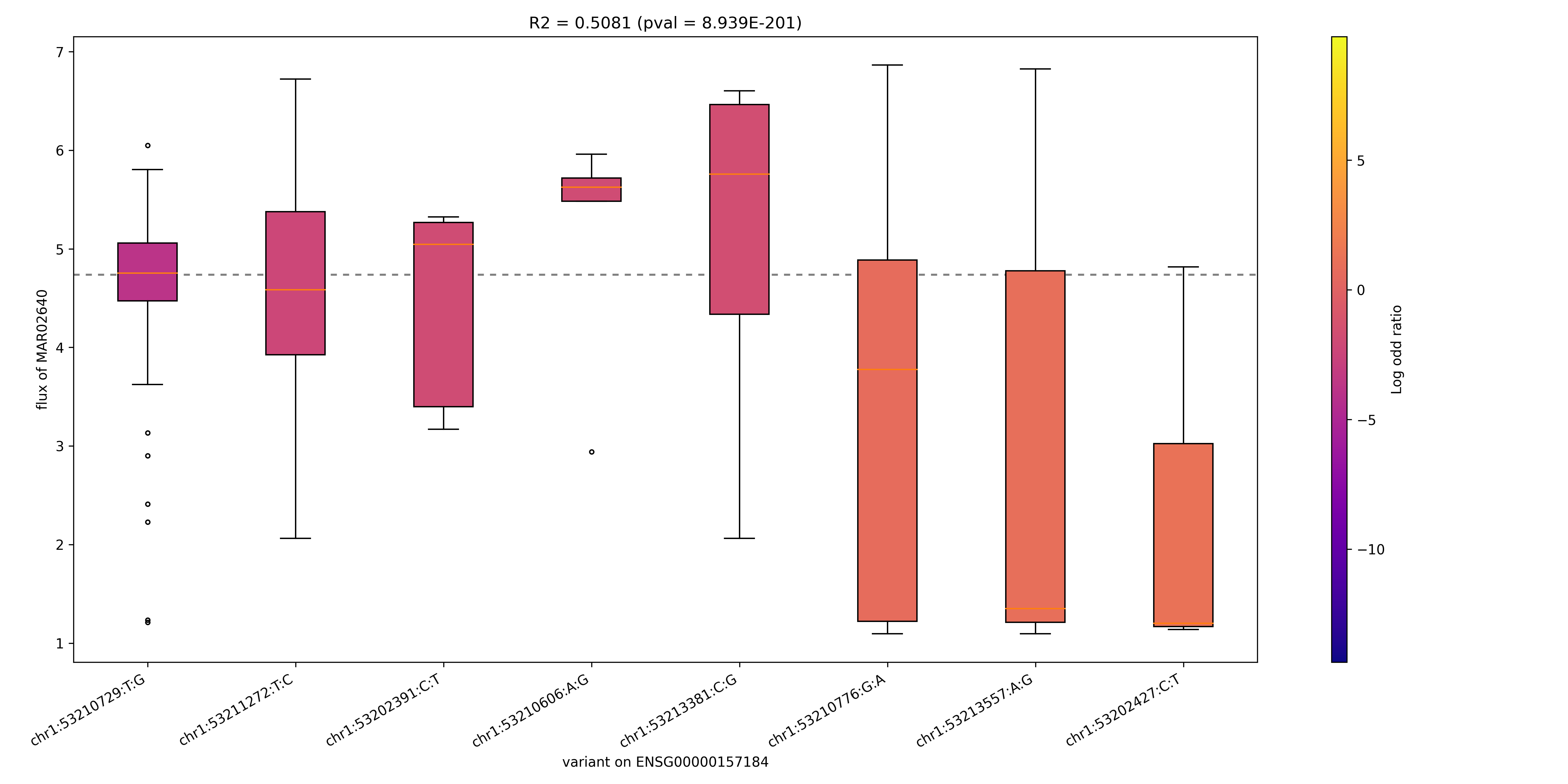

### MAR02657_ENSG00000110090.png

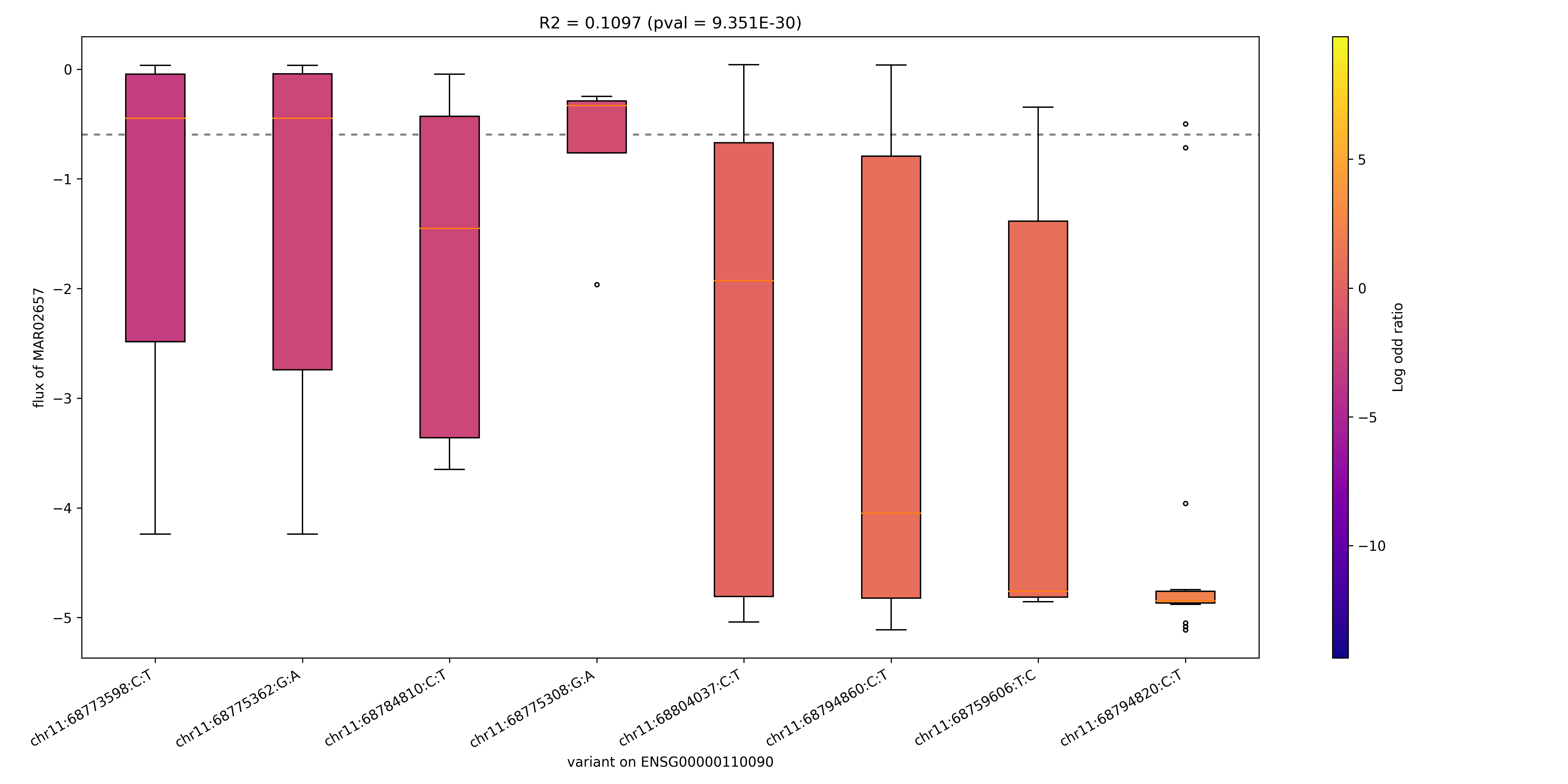

### MAR02657_ENSG00000169169.png

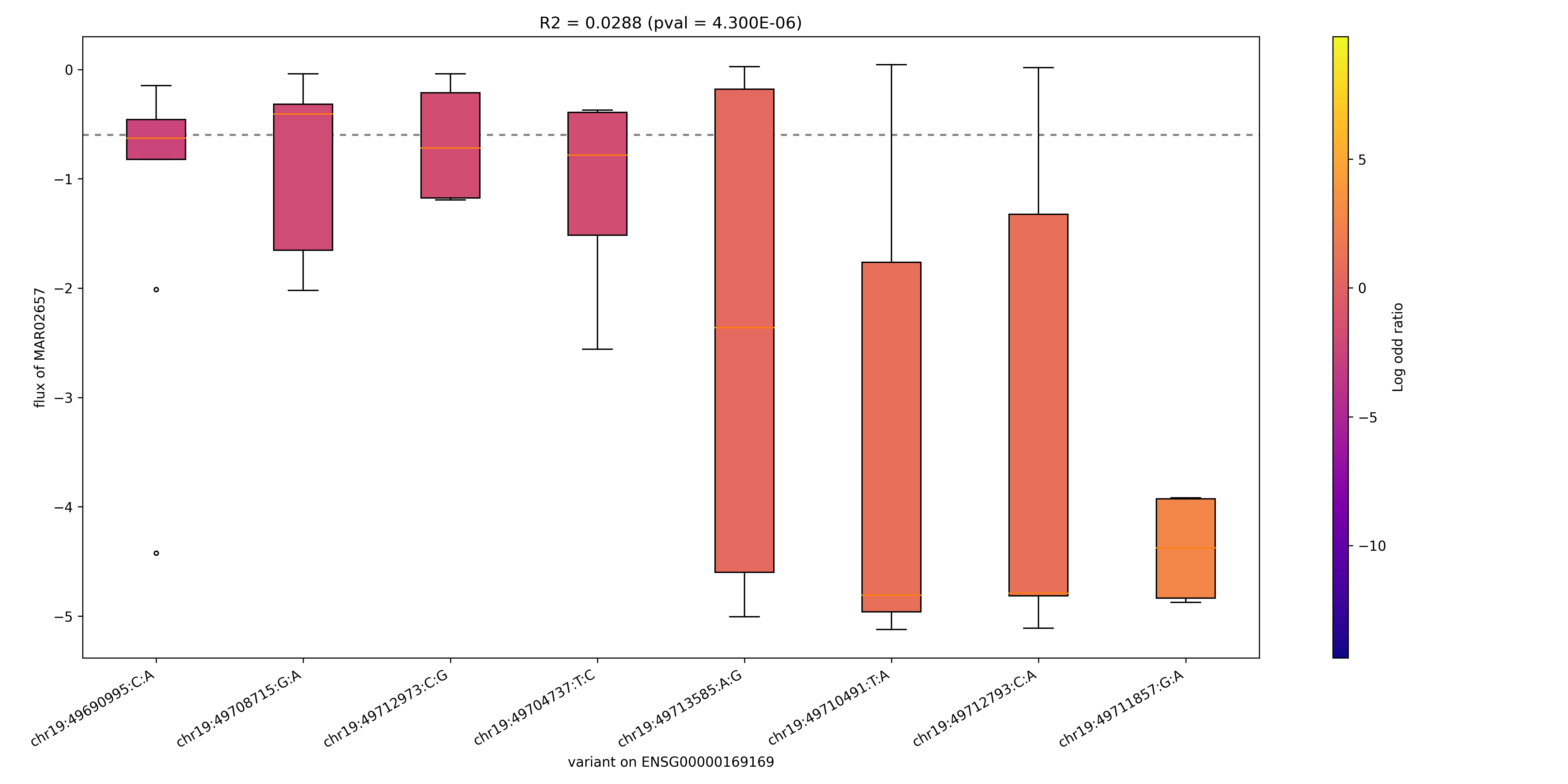

### MAR02666_ENSG00000205560.png

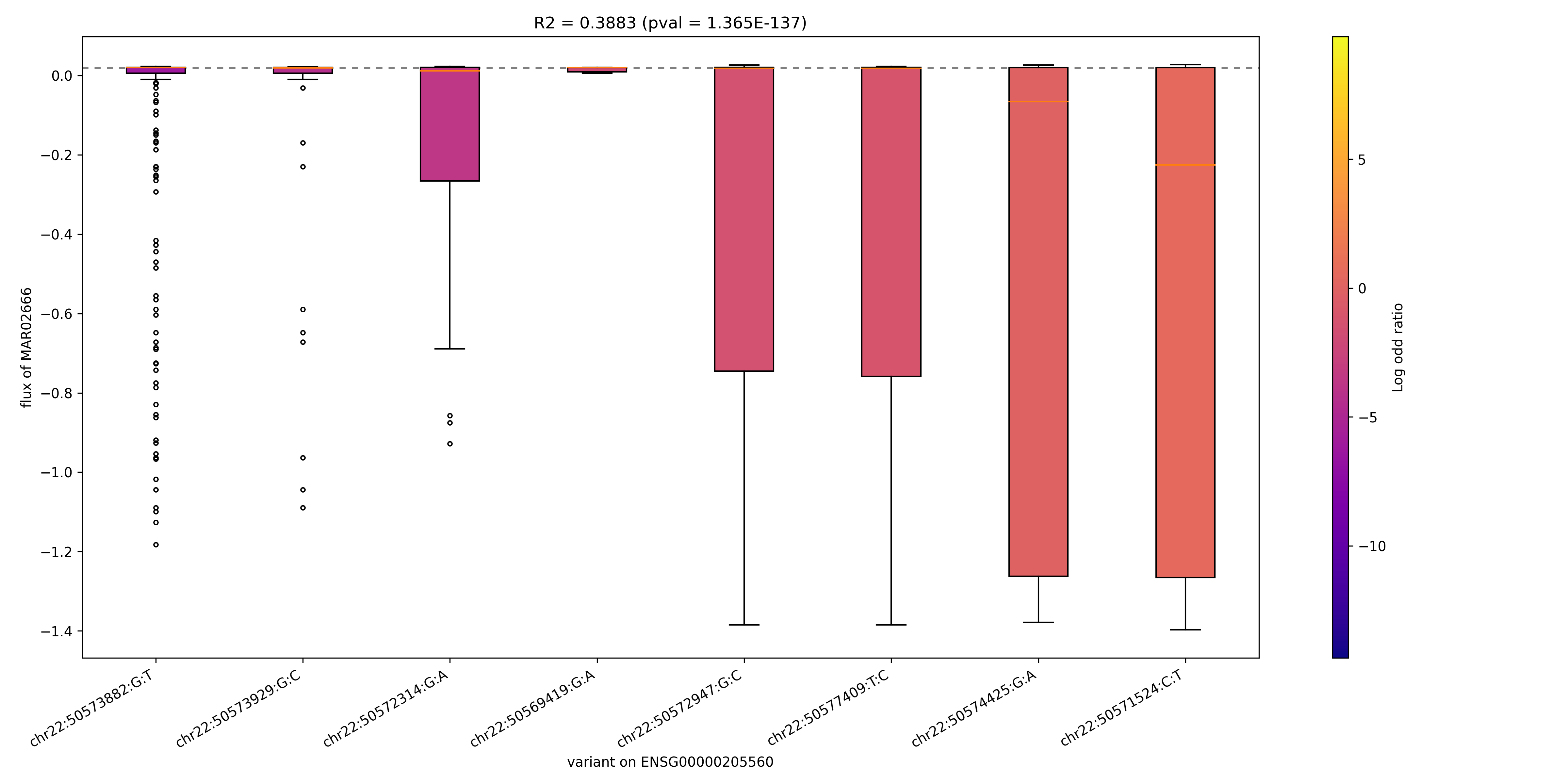

### MAR03226_ENSG00000072778.png

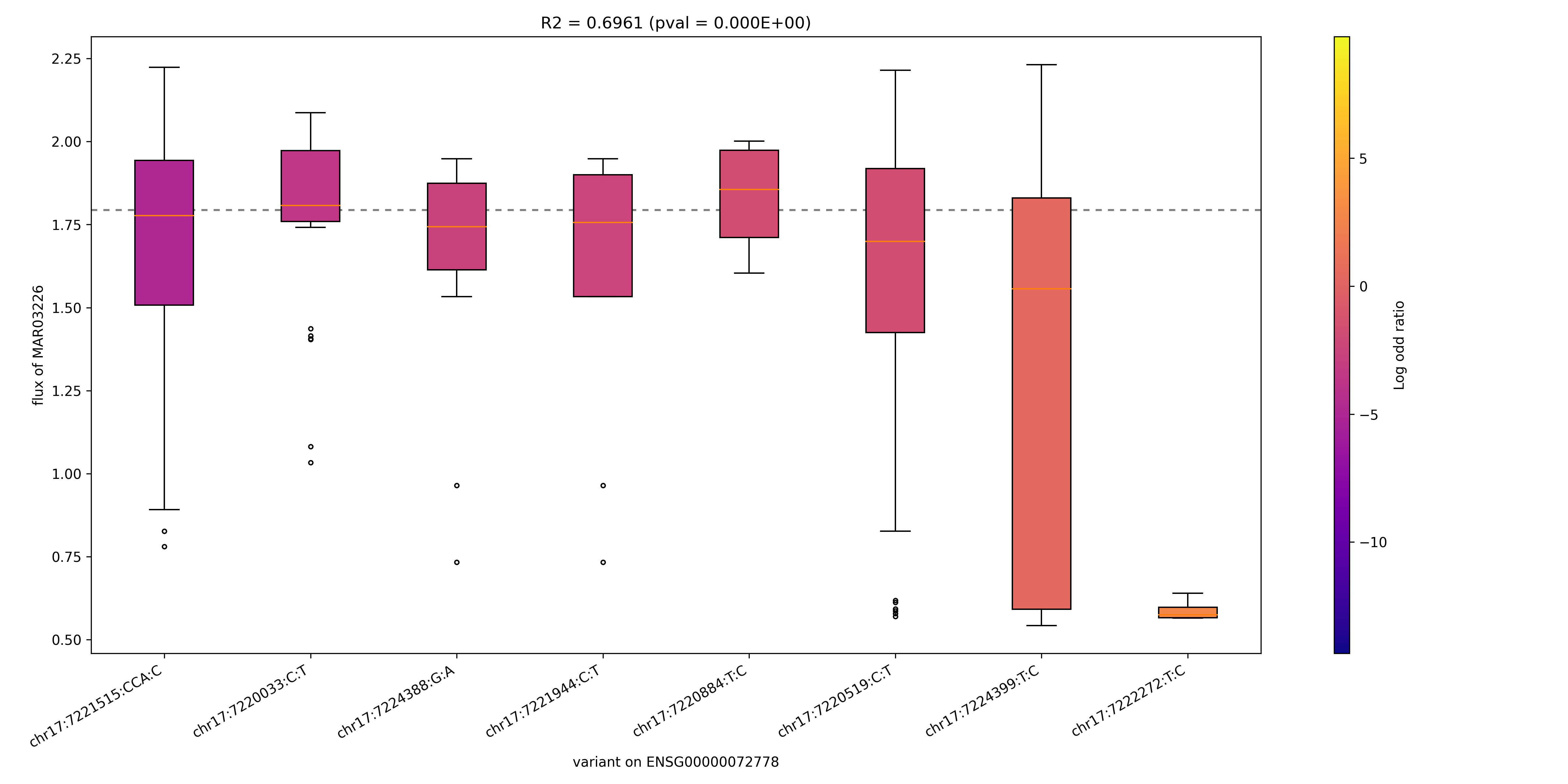

### MAR03226_ENSG00000117054.png

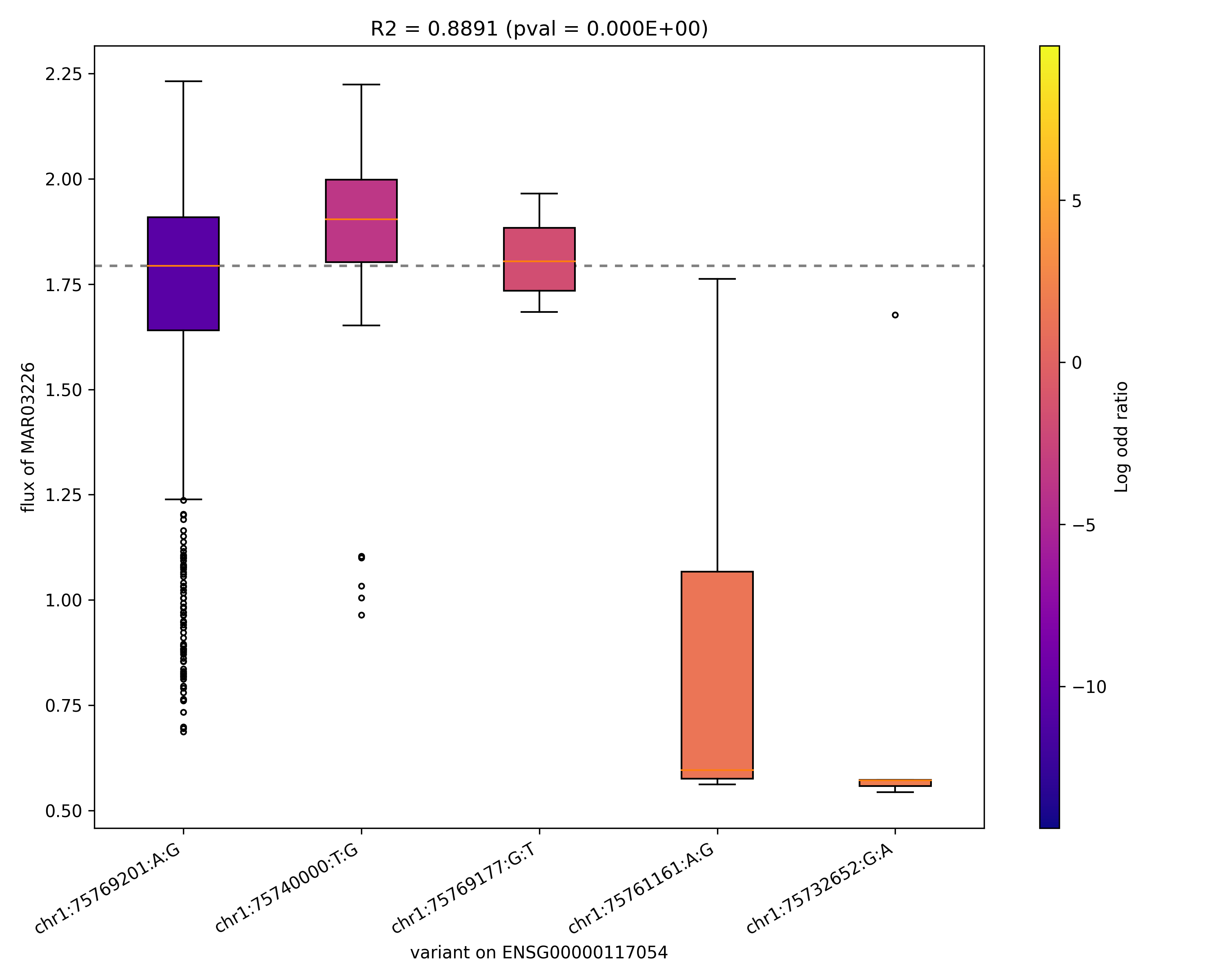

### MAR03227_ENSG00000113790.png

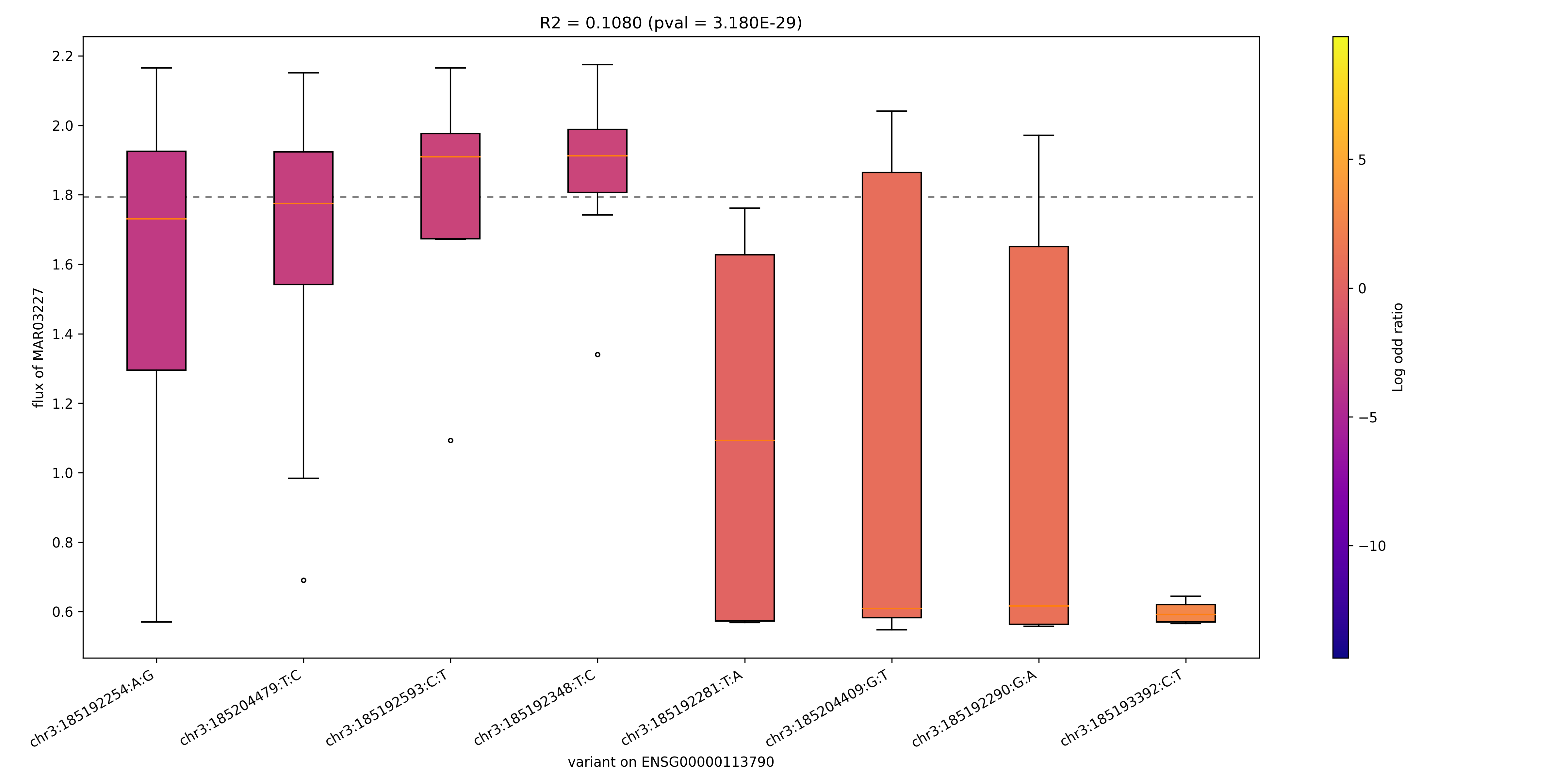

### MAR03273_ENSG00000197142.png

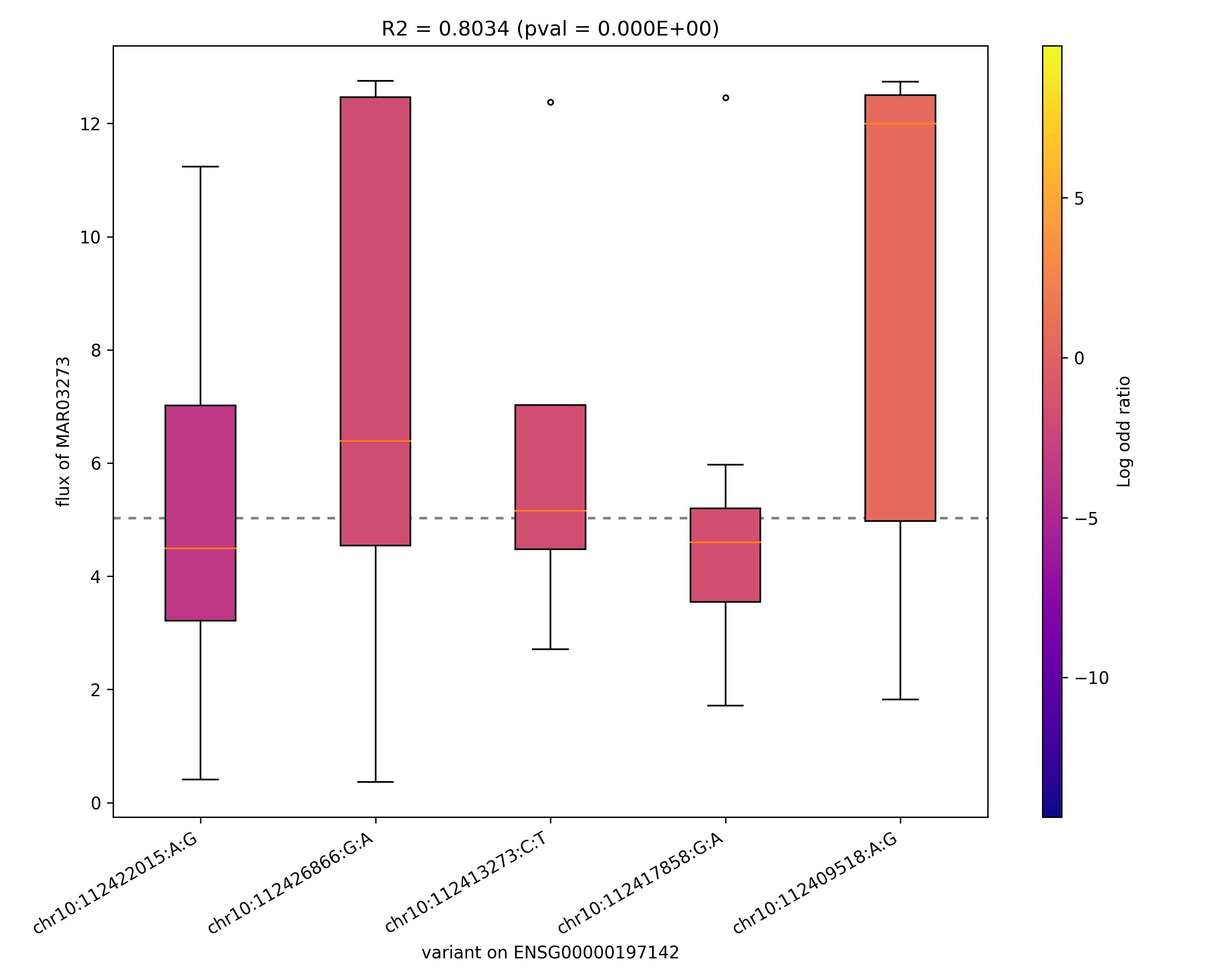

### MAR03846_ENSG00000123983.png

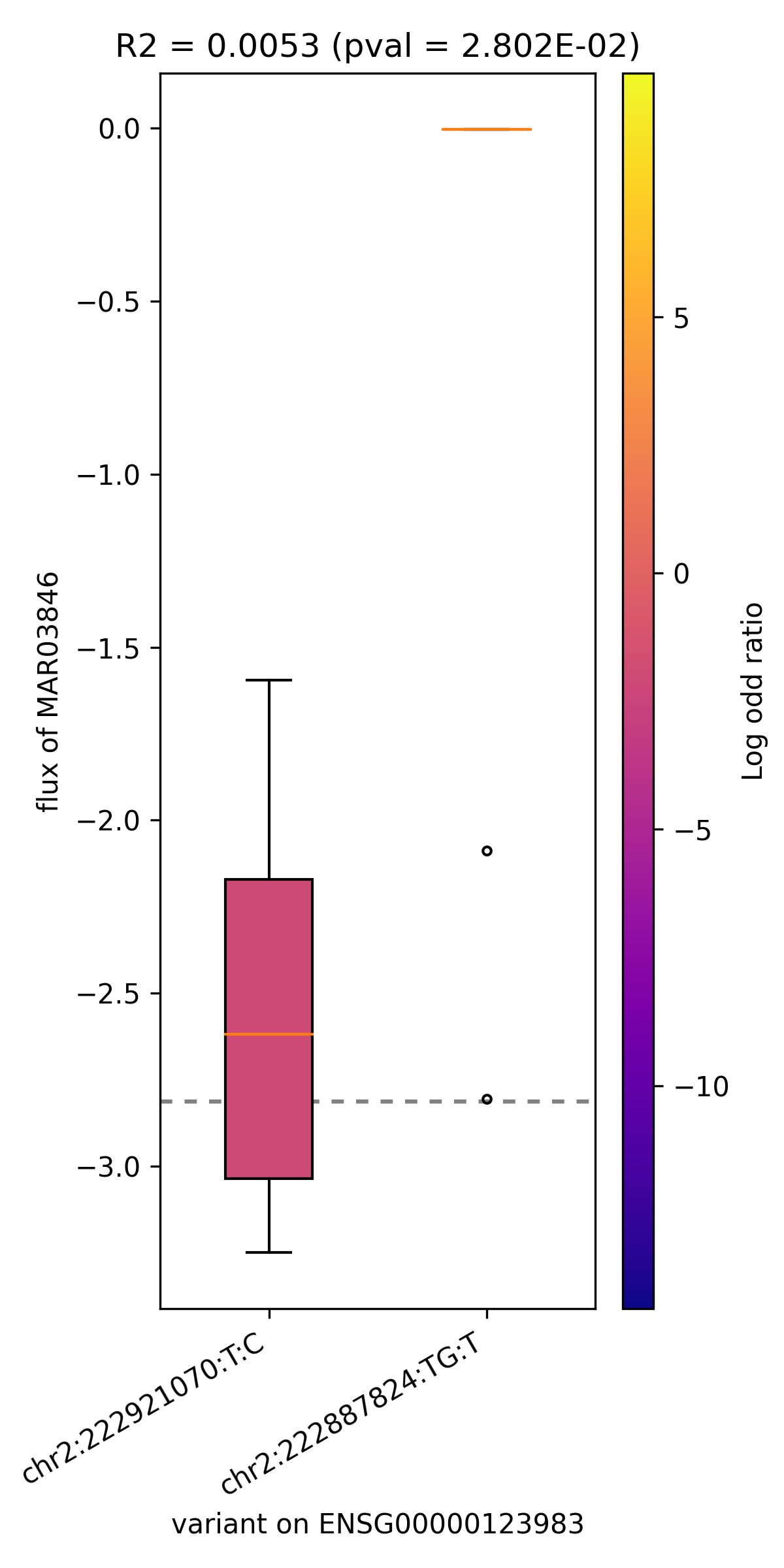

### MAR03846_ENSG00000151726.png

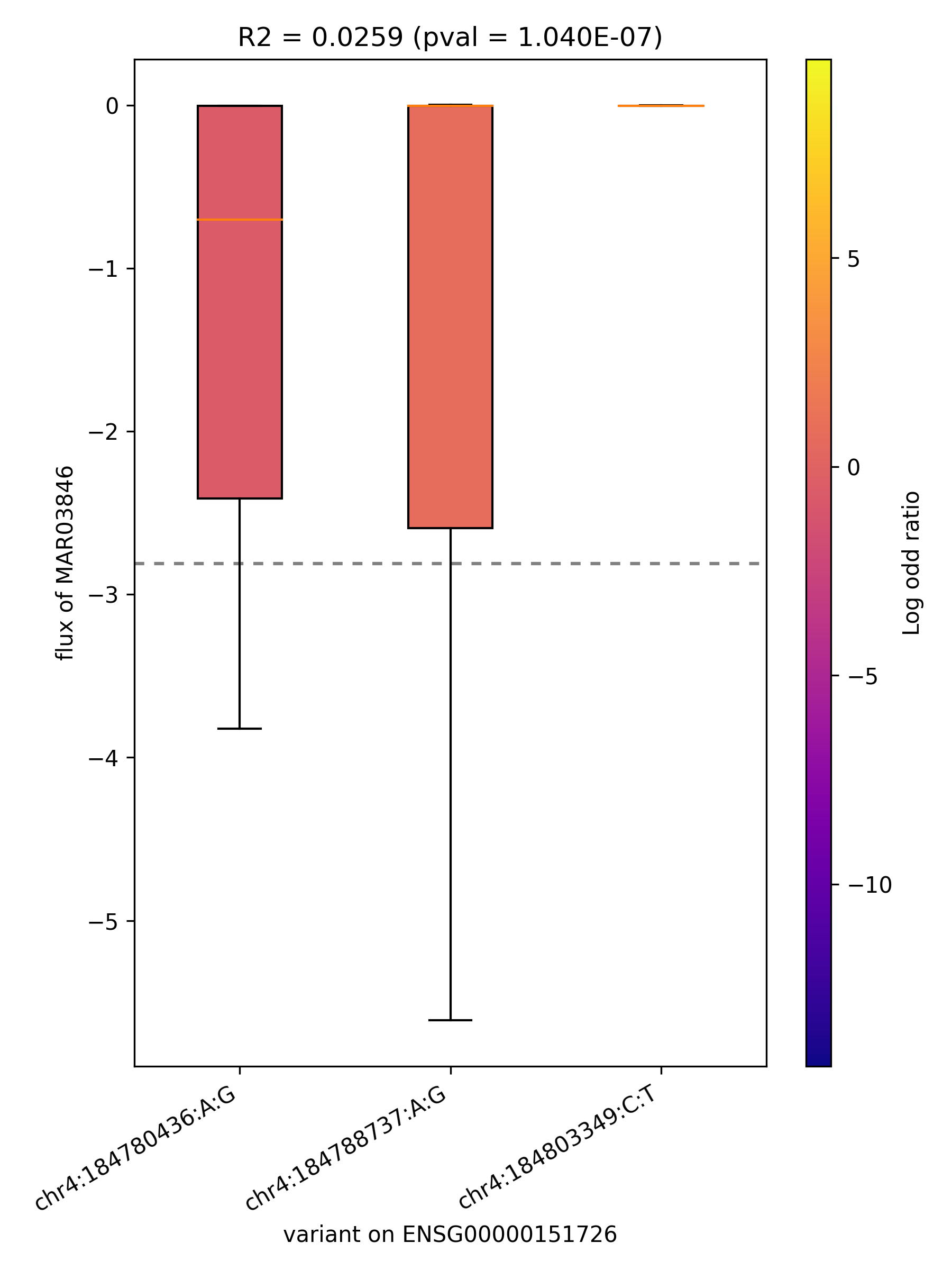

### MAR03846_ENSG00000164398.png

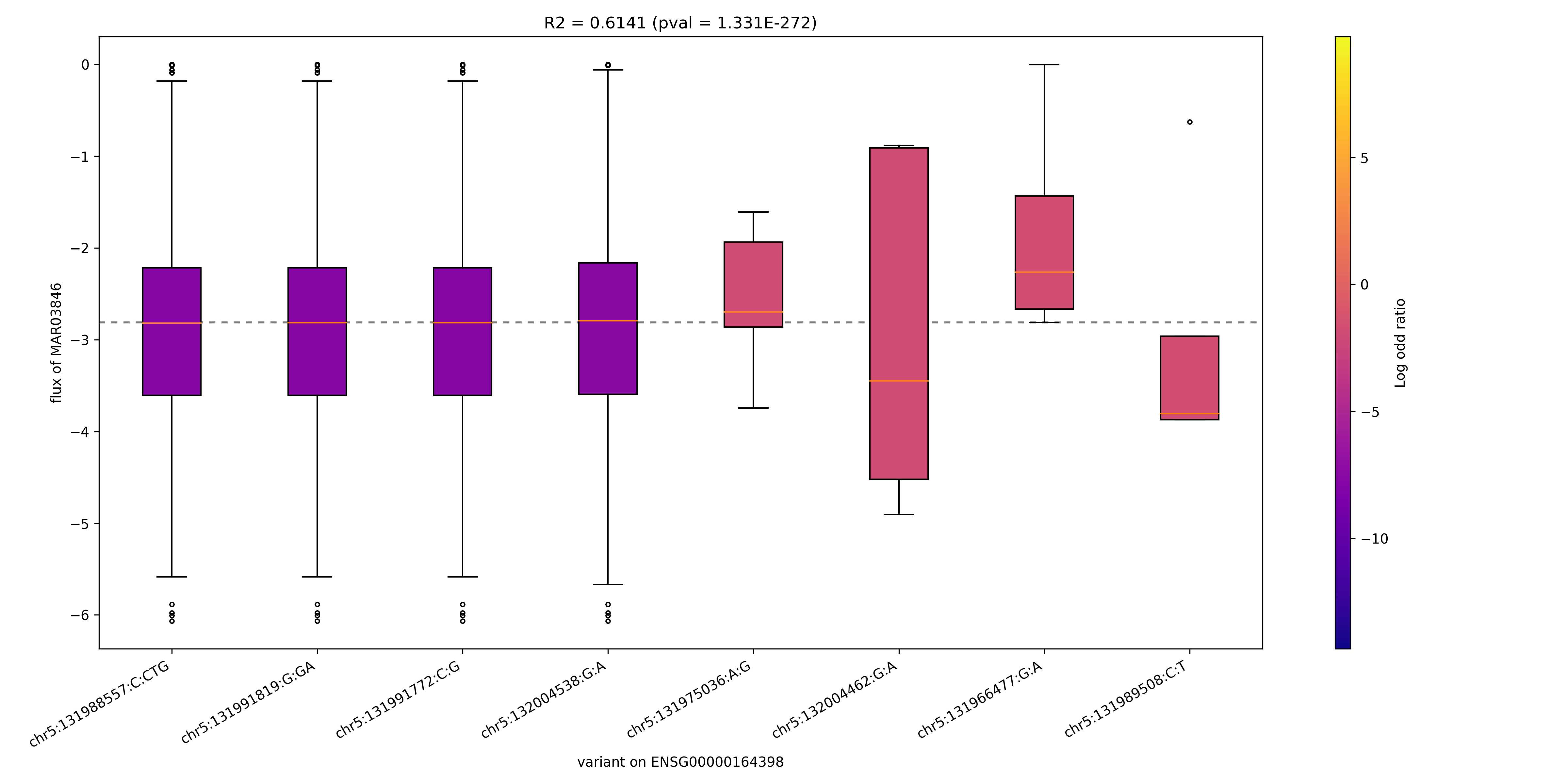

### MAR03999_ENSG00000138185.png

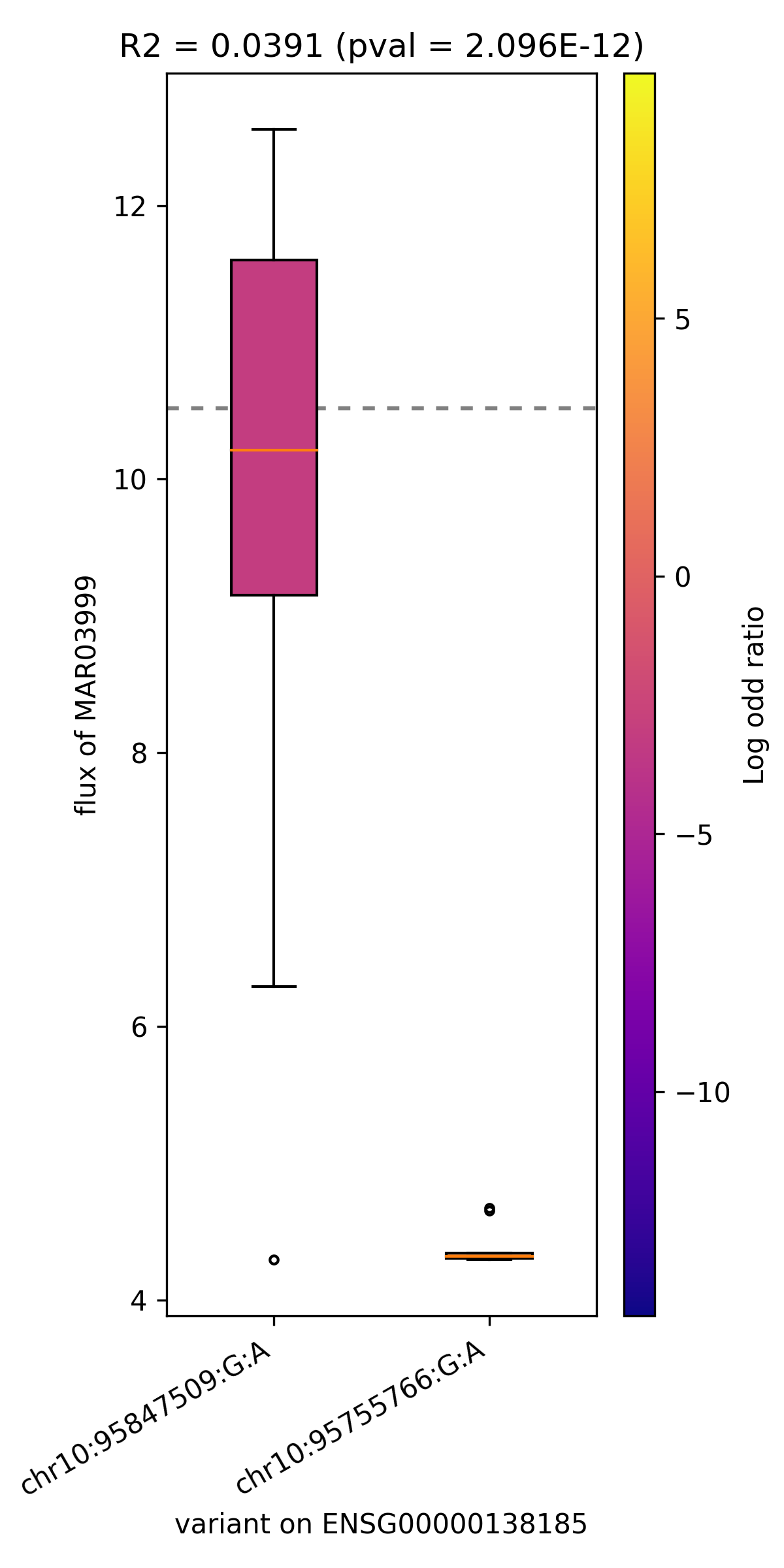

### MAR04004_ENSG00000004455.png

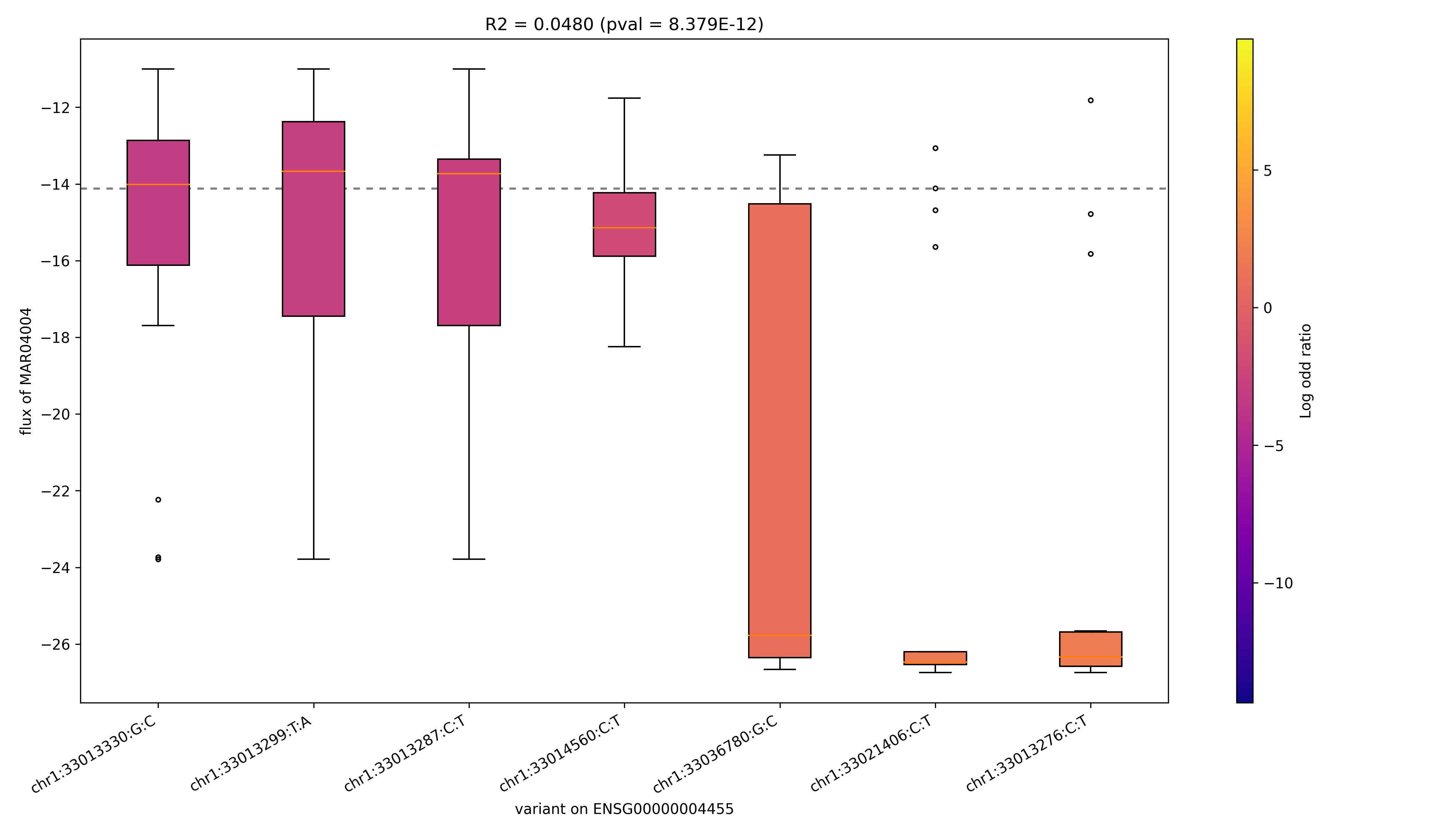

### MAR04004_ENSG00000155085.png

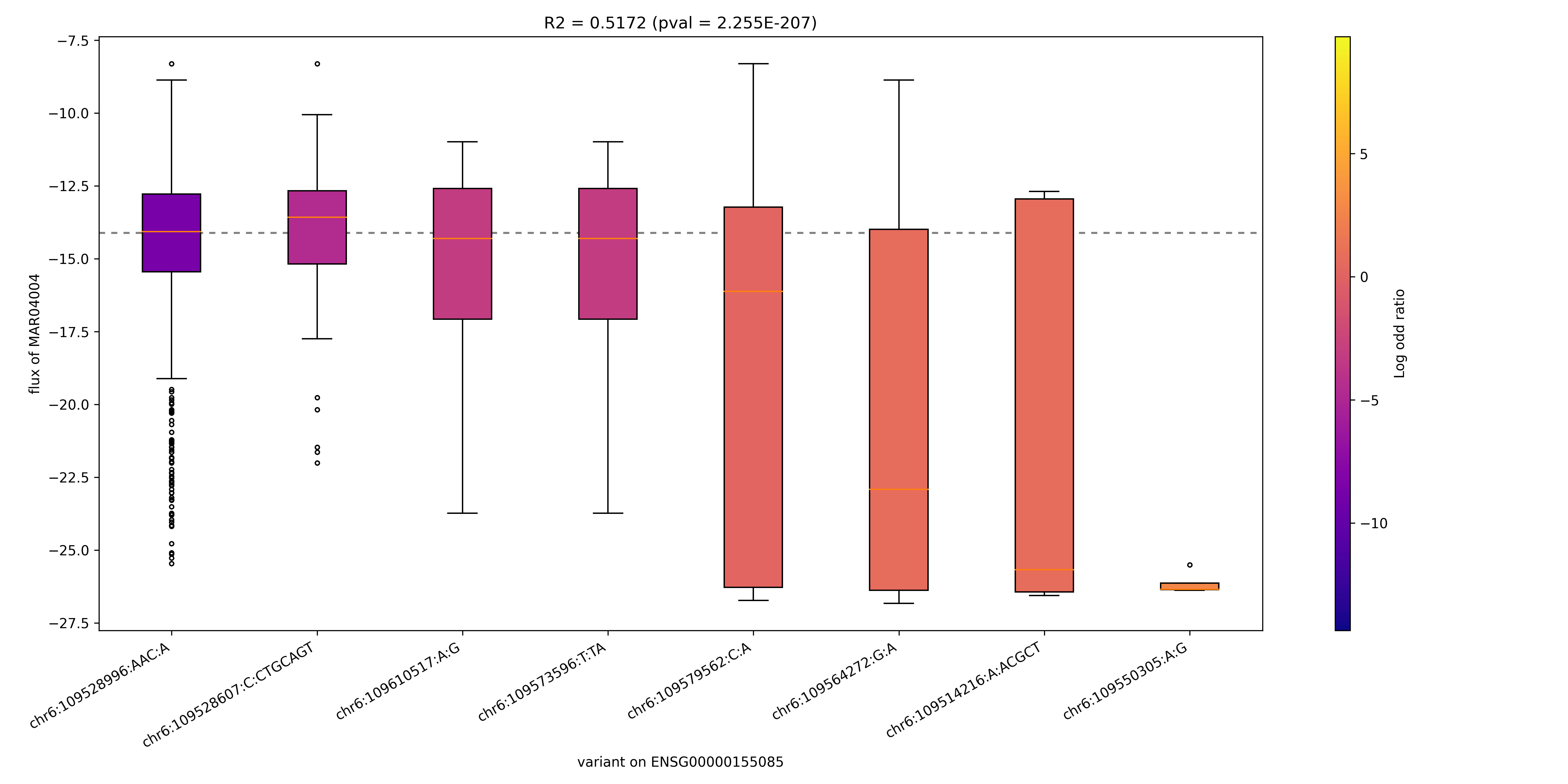

### MAR04004_ENSG00000165695.png

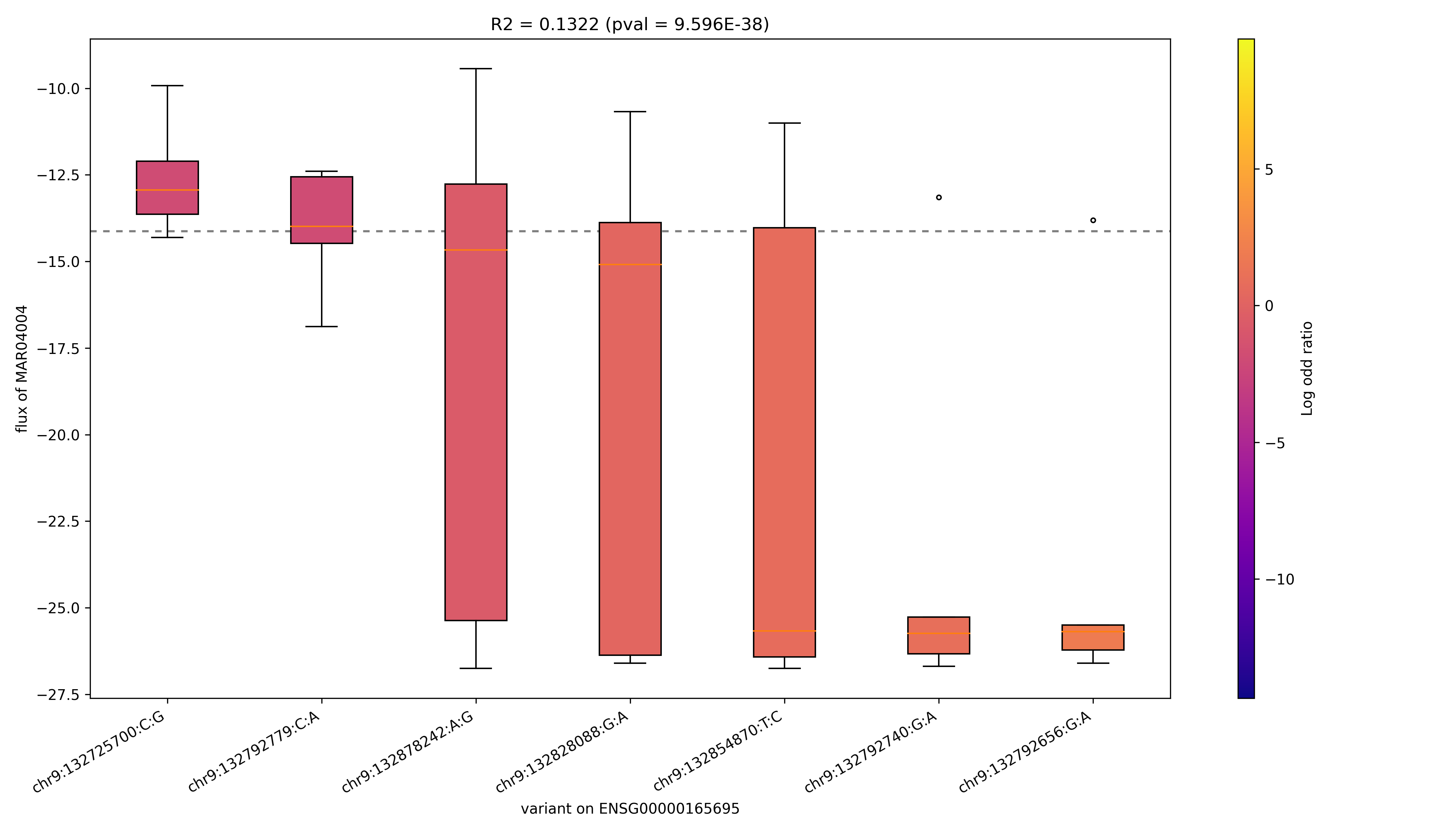

### MAR04004_ENSG00000273841.png

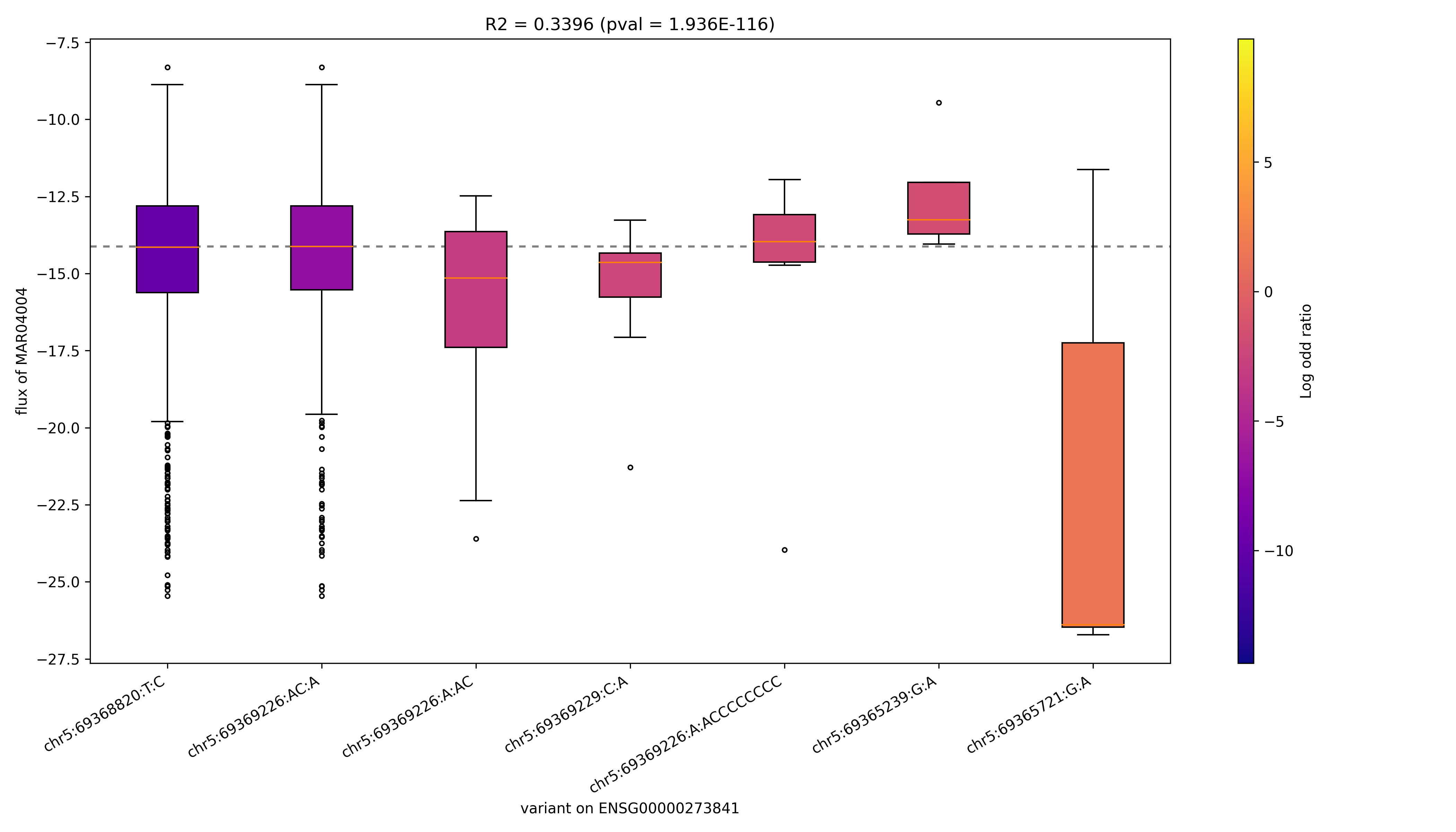

### MAR04012_ENSG00000156110.png

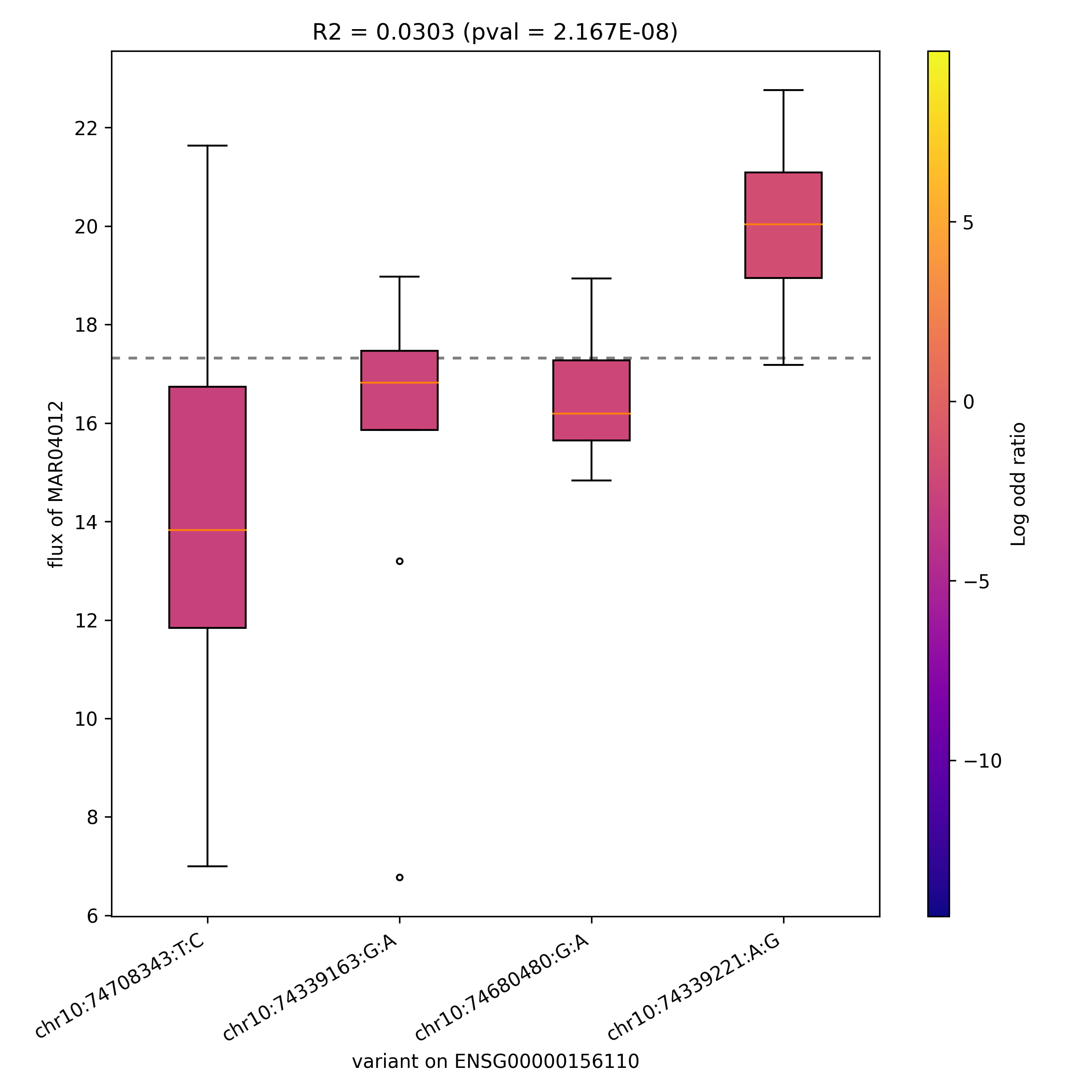

### MAR04171_ENSG00000067225.png

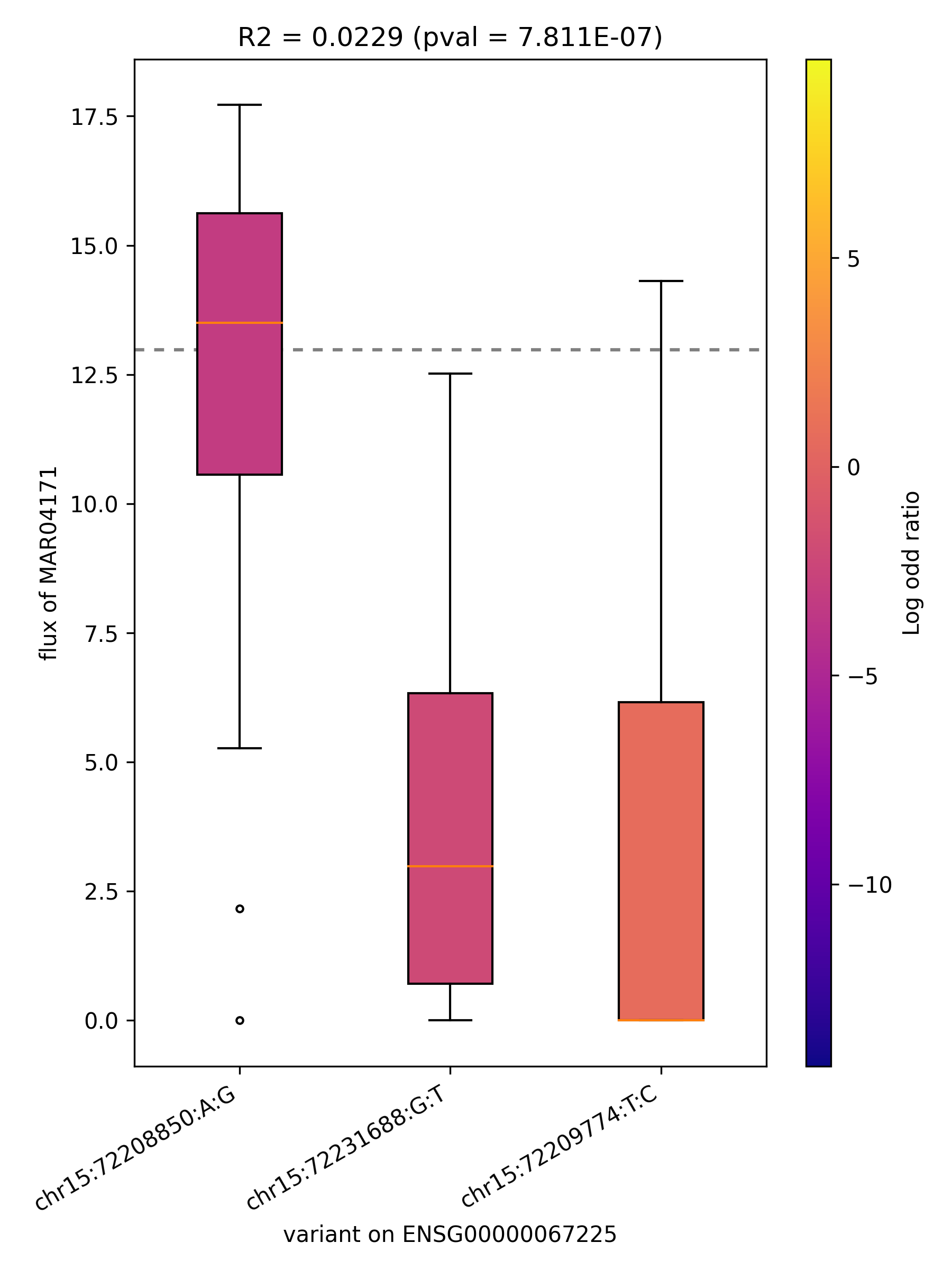

### MAR04171_ENSG00000143627.png

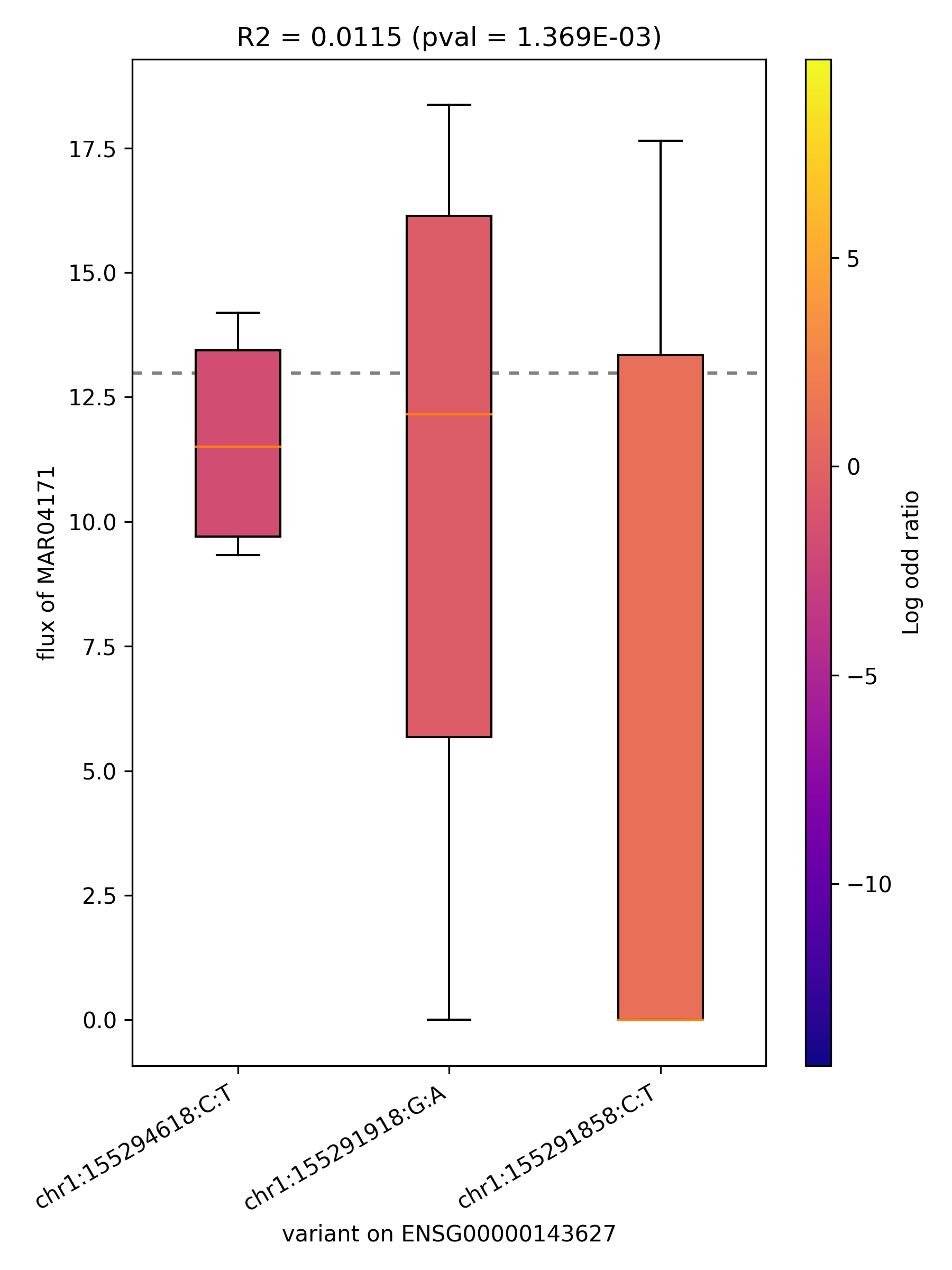

### MAR04267_ENSG00000157064.png

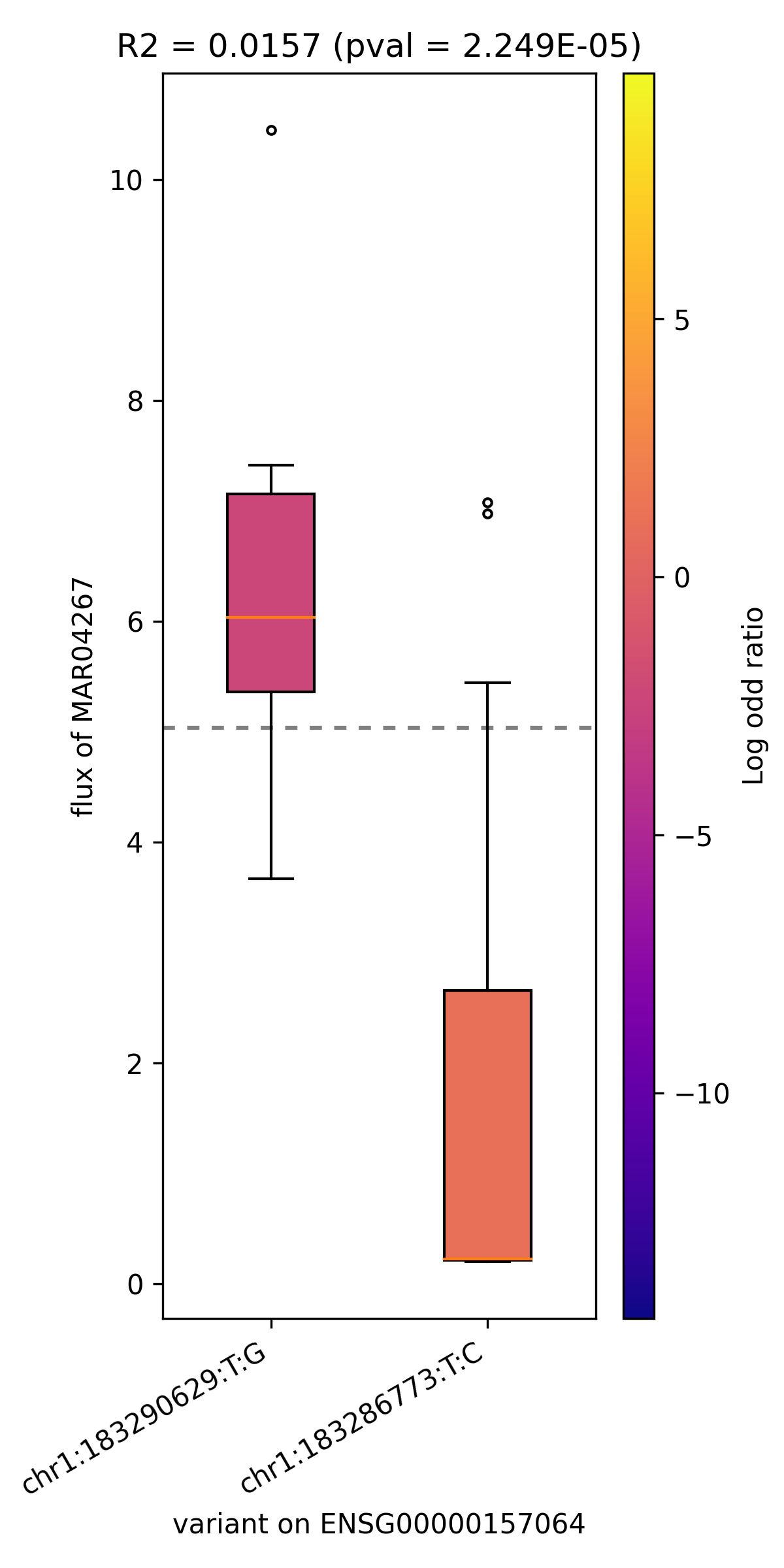

### MAR04394_ENSG00000106633.png

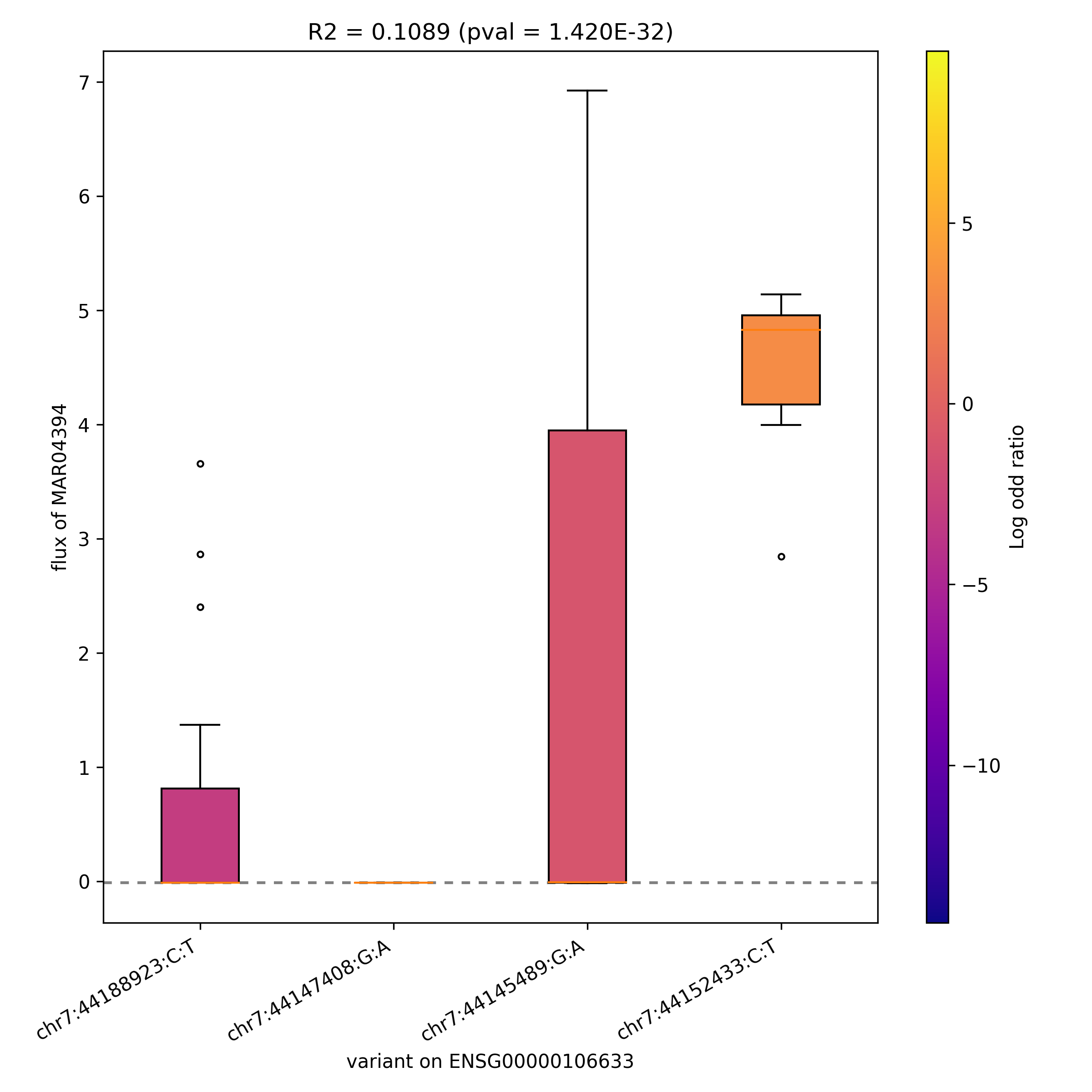

### MAR04394_ENSG00000156510.png

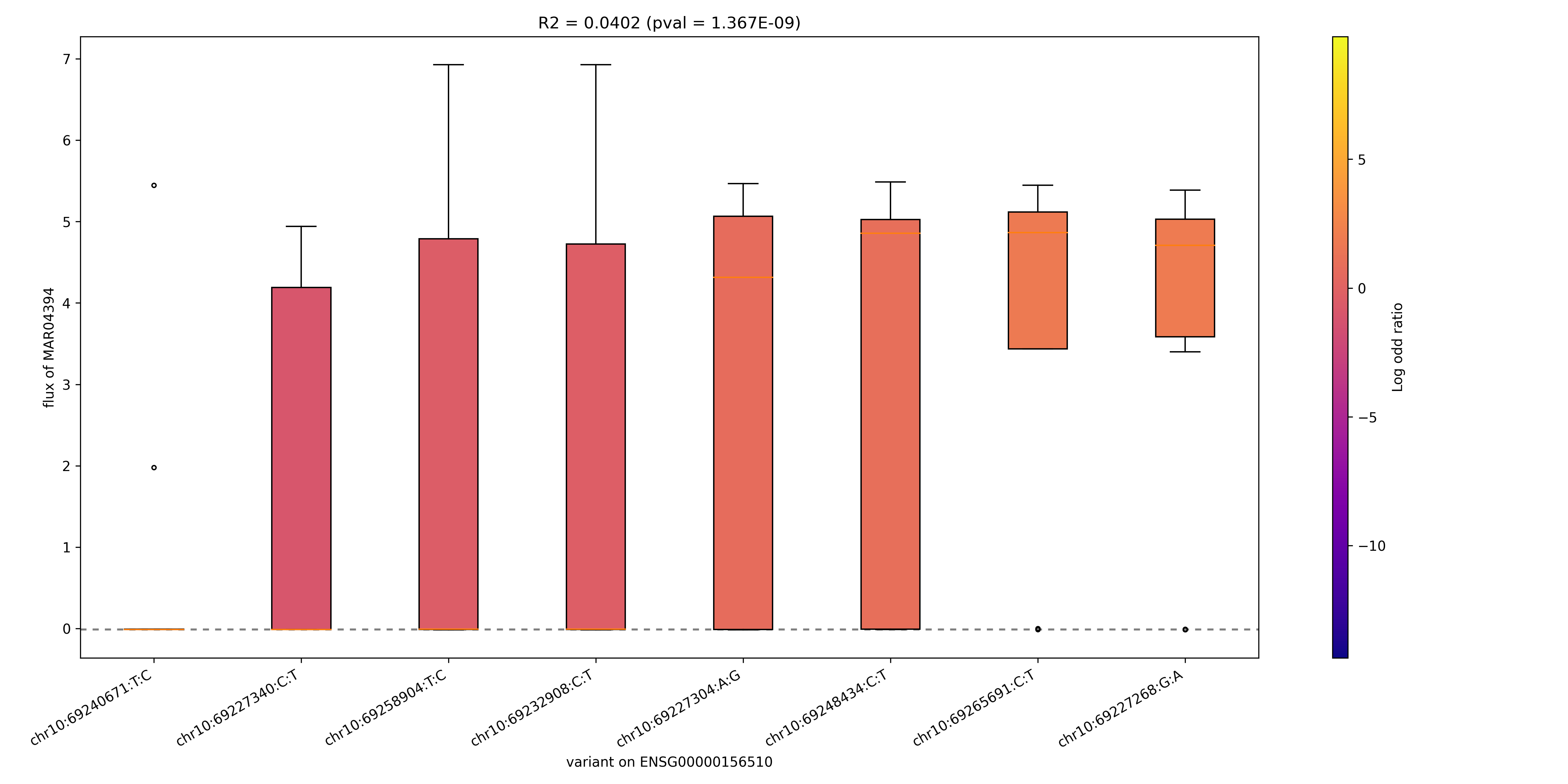

### MAR04394_ENSG00000156515.png

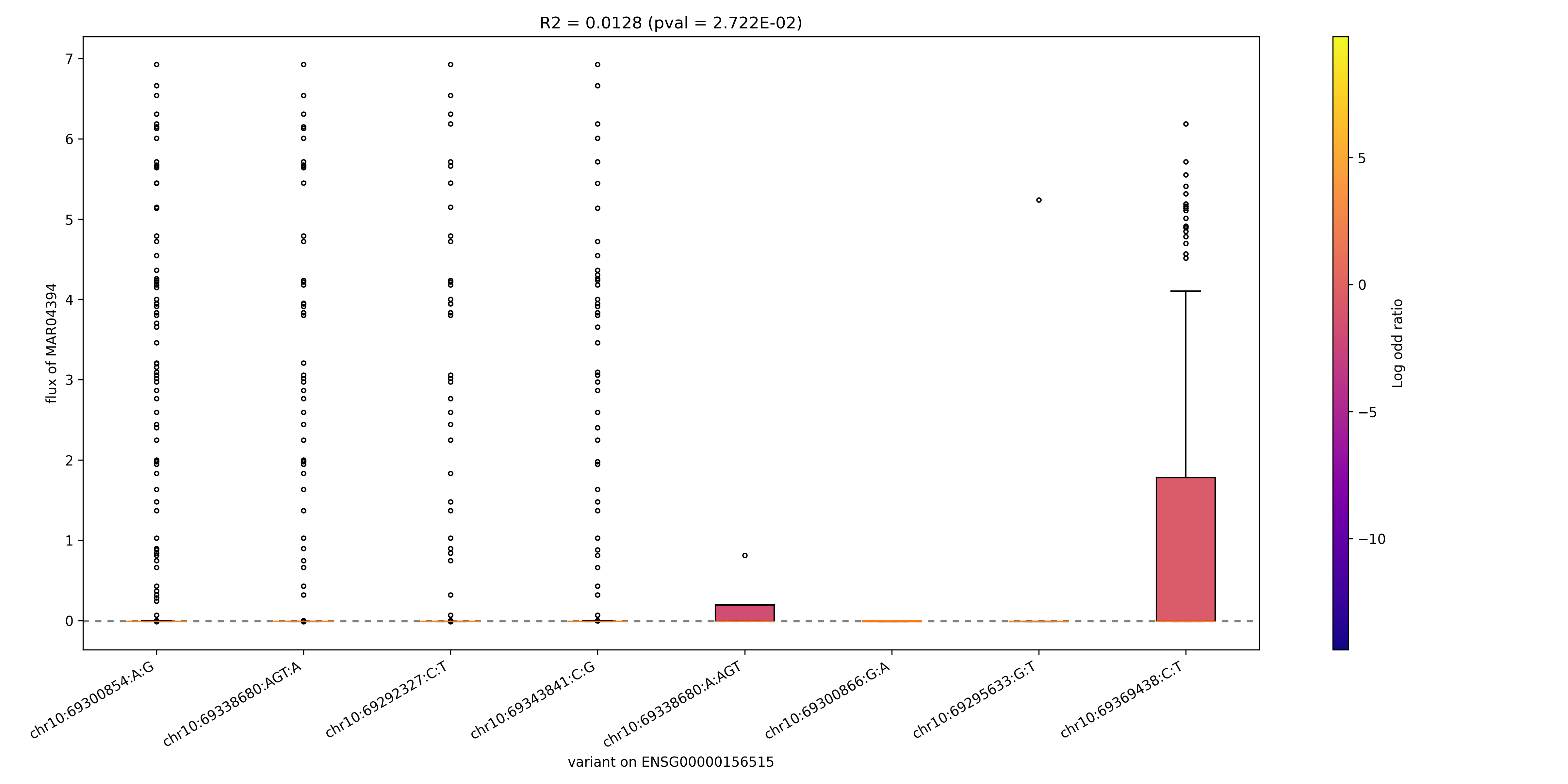

### MAR04394_ENSG00000160883.png

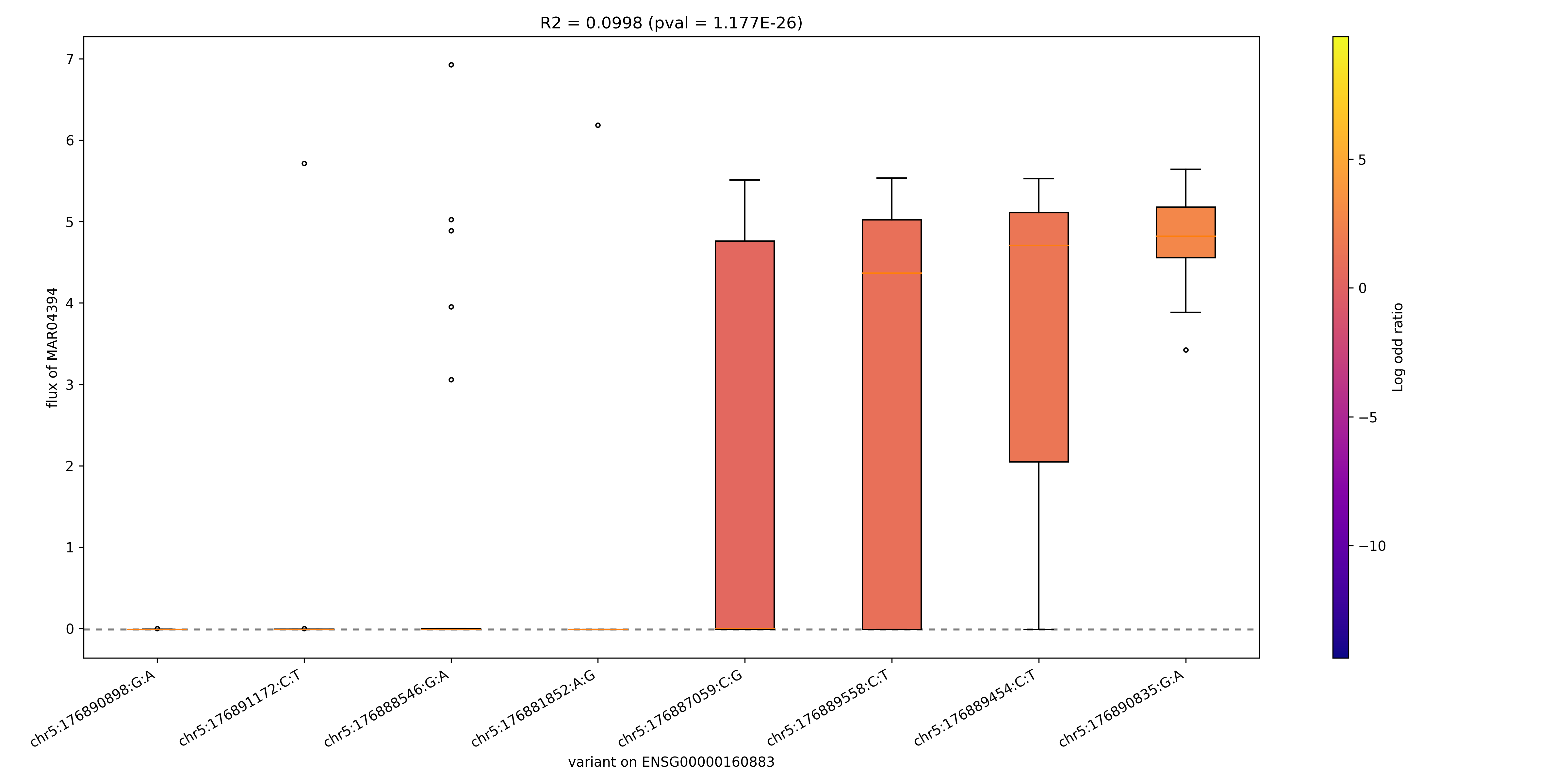

### MAR04473_ENSG00000142657.png

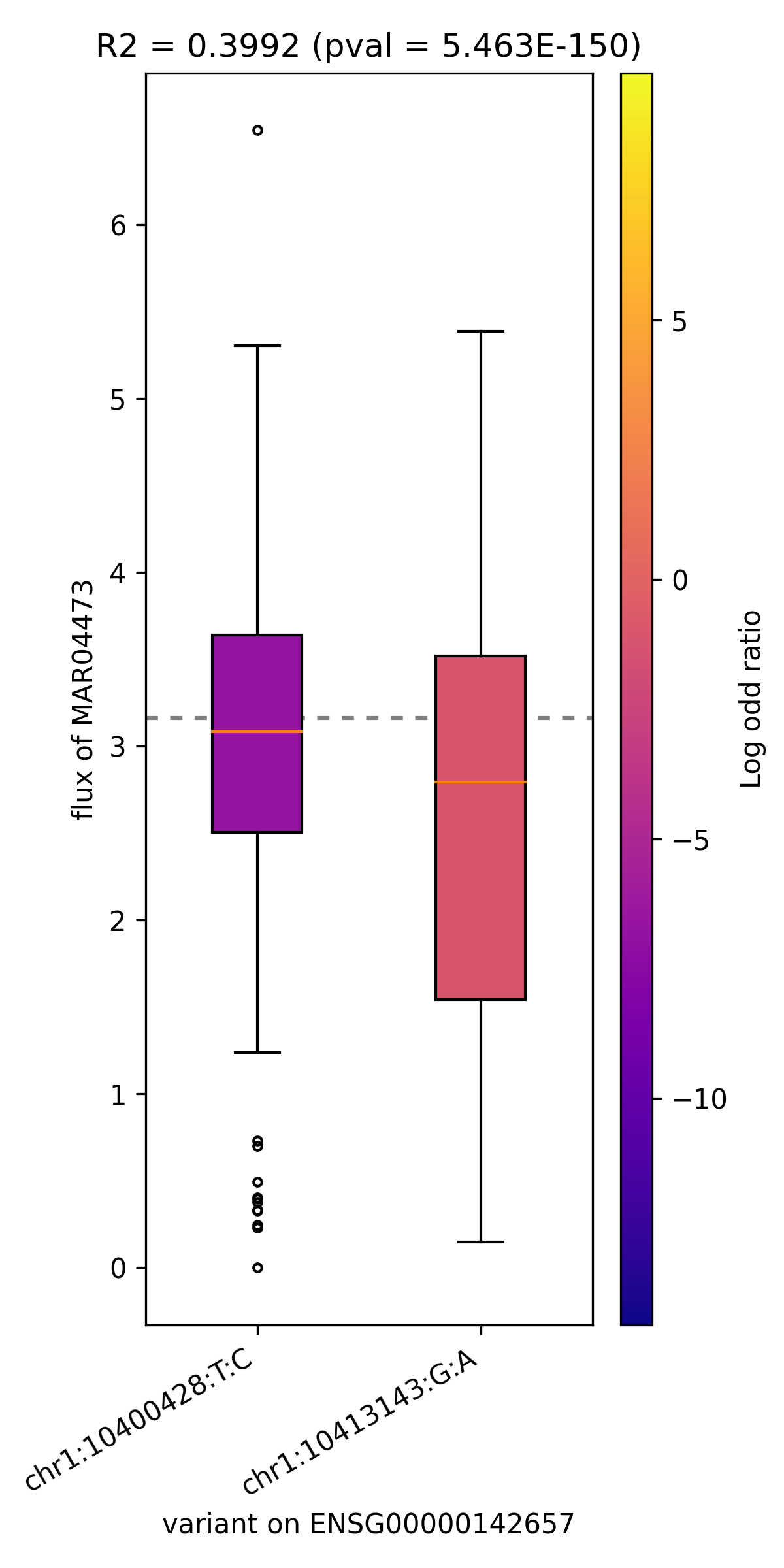

### MAR04489_ENSG00000159399.png

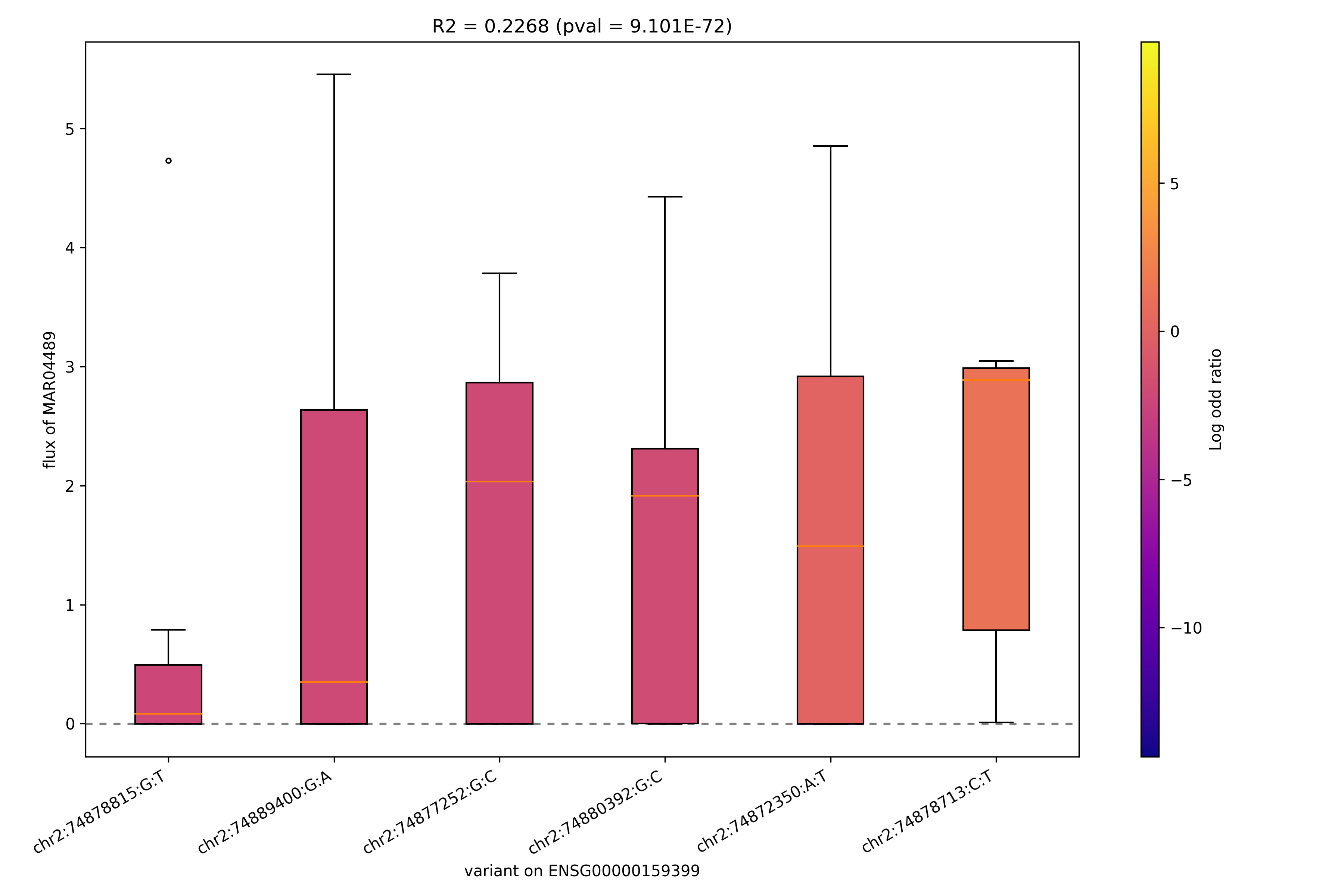

### MAR04490_ENSG00000159399.png

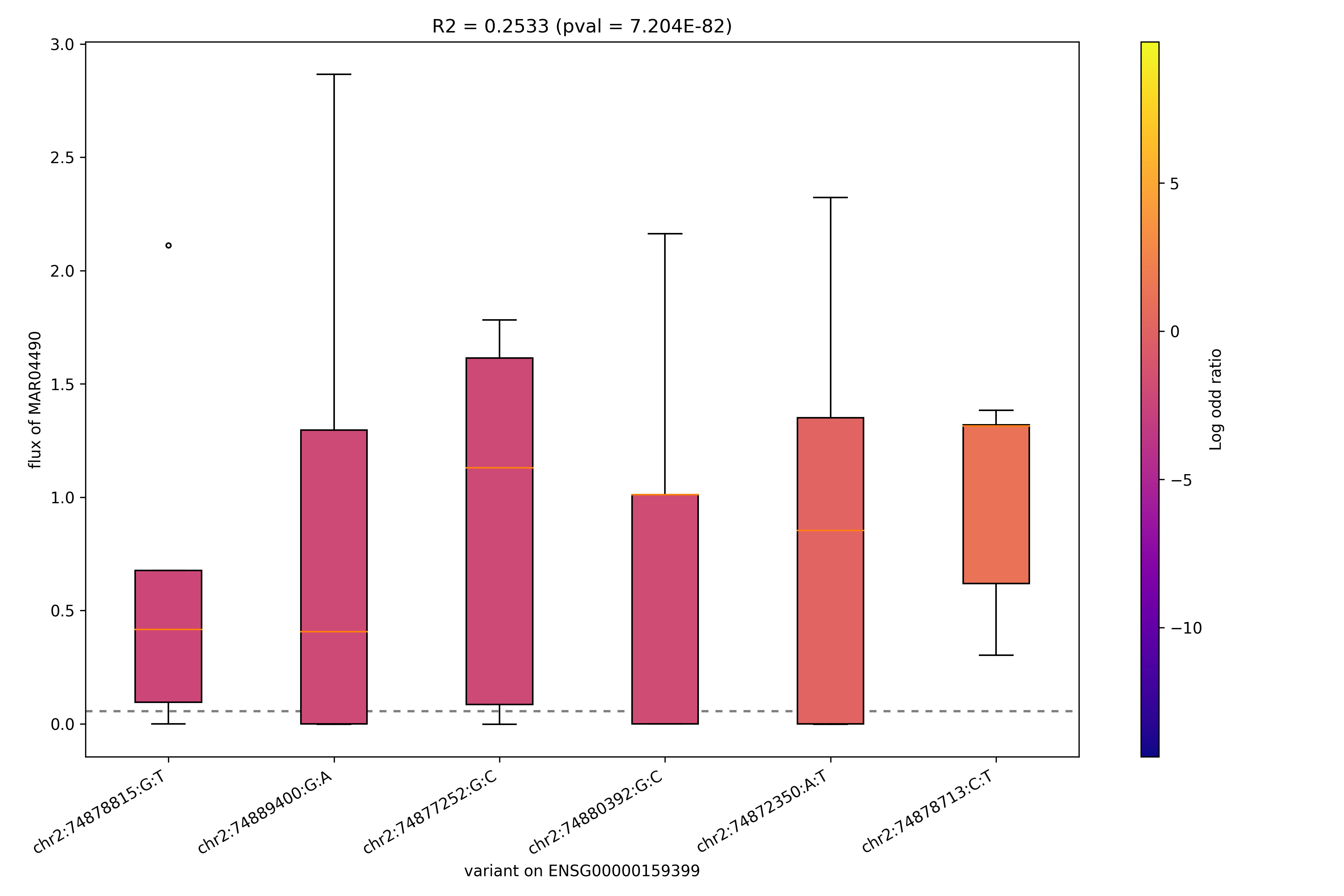

### MAR04513_ENSG00000156136.png

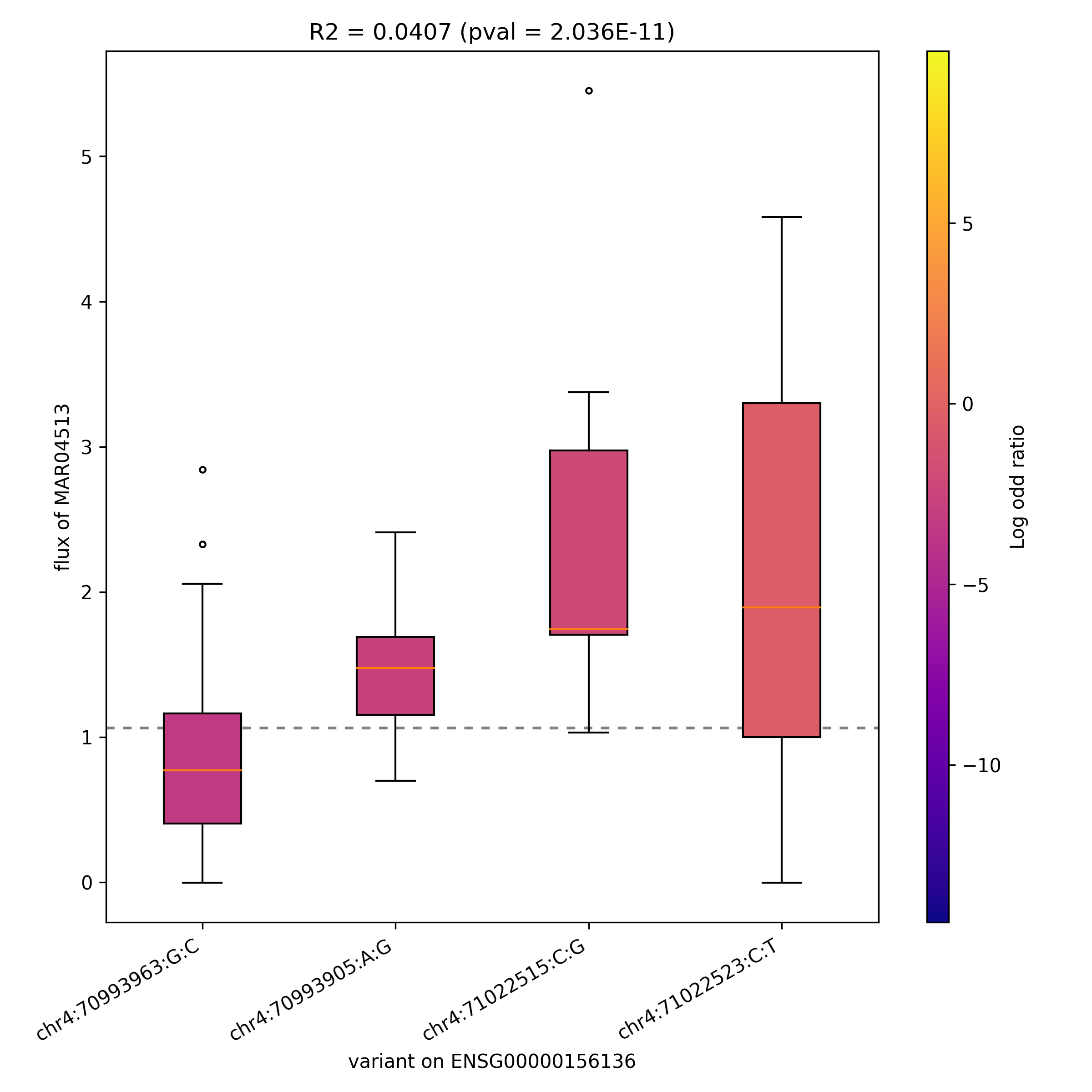

### MAR04611_ENSG00000171848.png

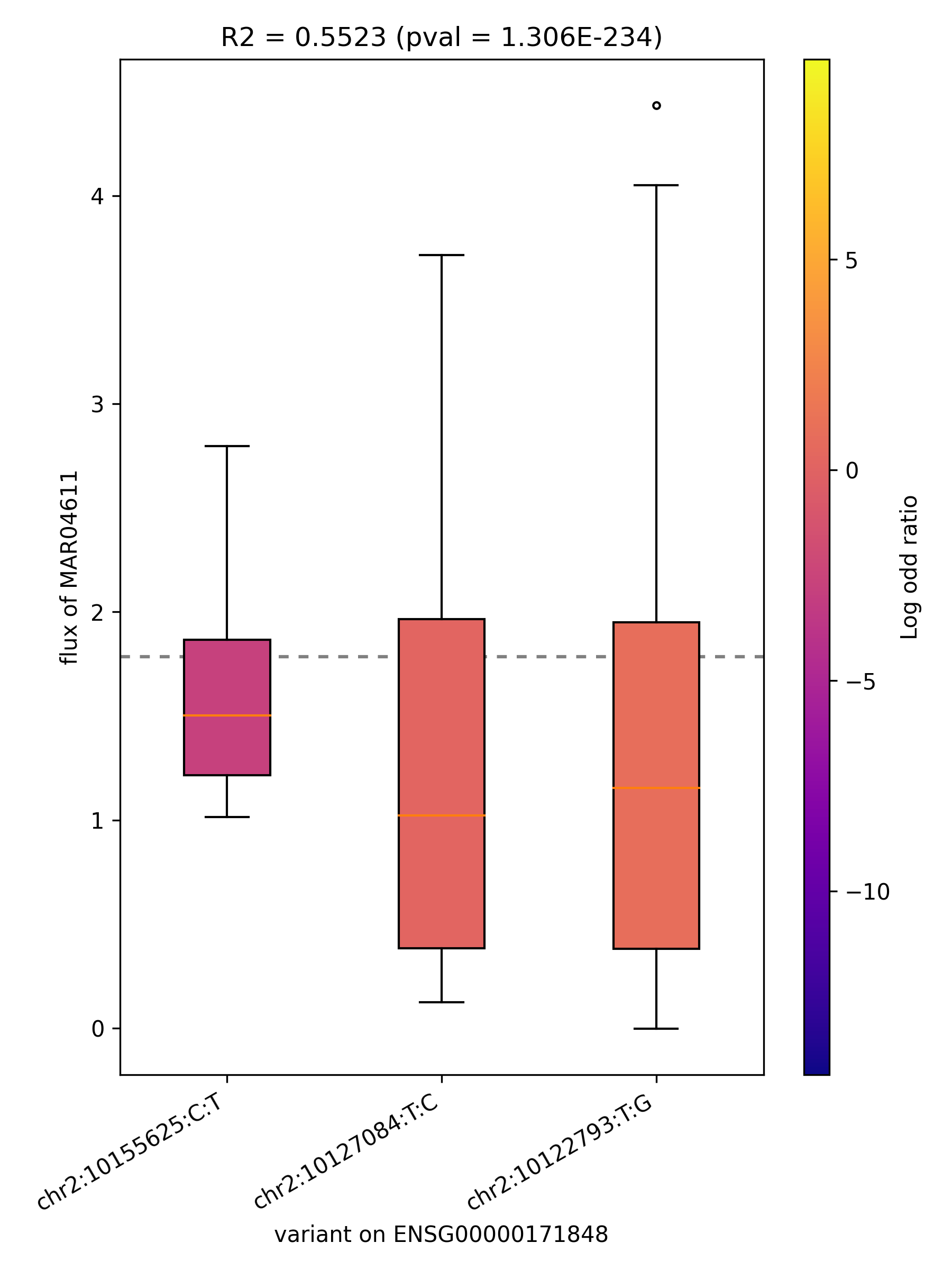

### MAR04615_ENSG00000167325.png

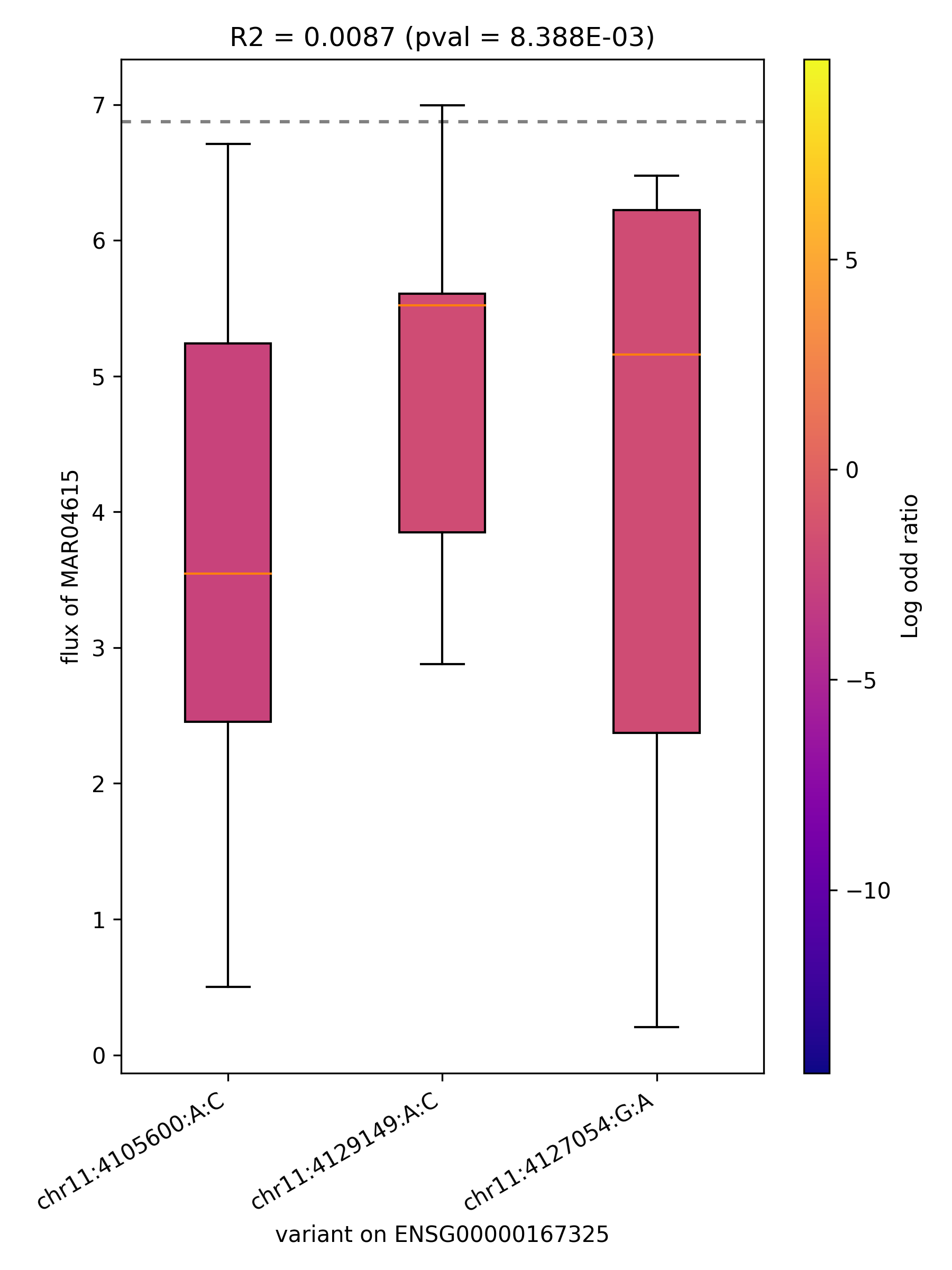

### MAR04615_ENSG00000171848.png

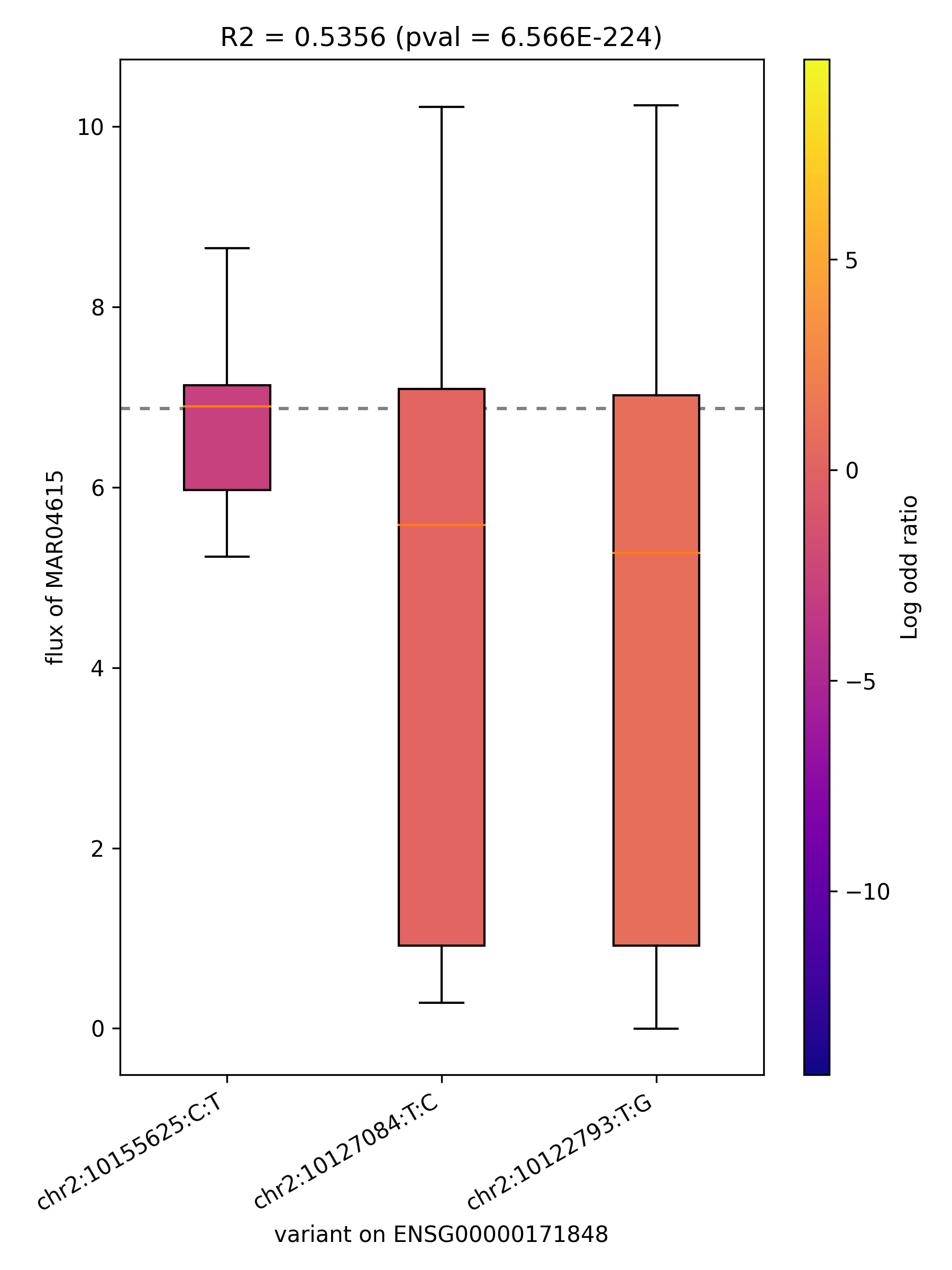

### MAR04632_ENSG00000076685.png

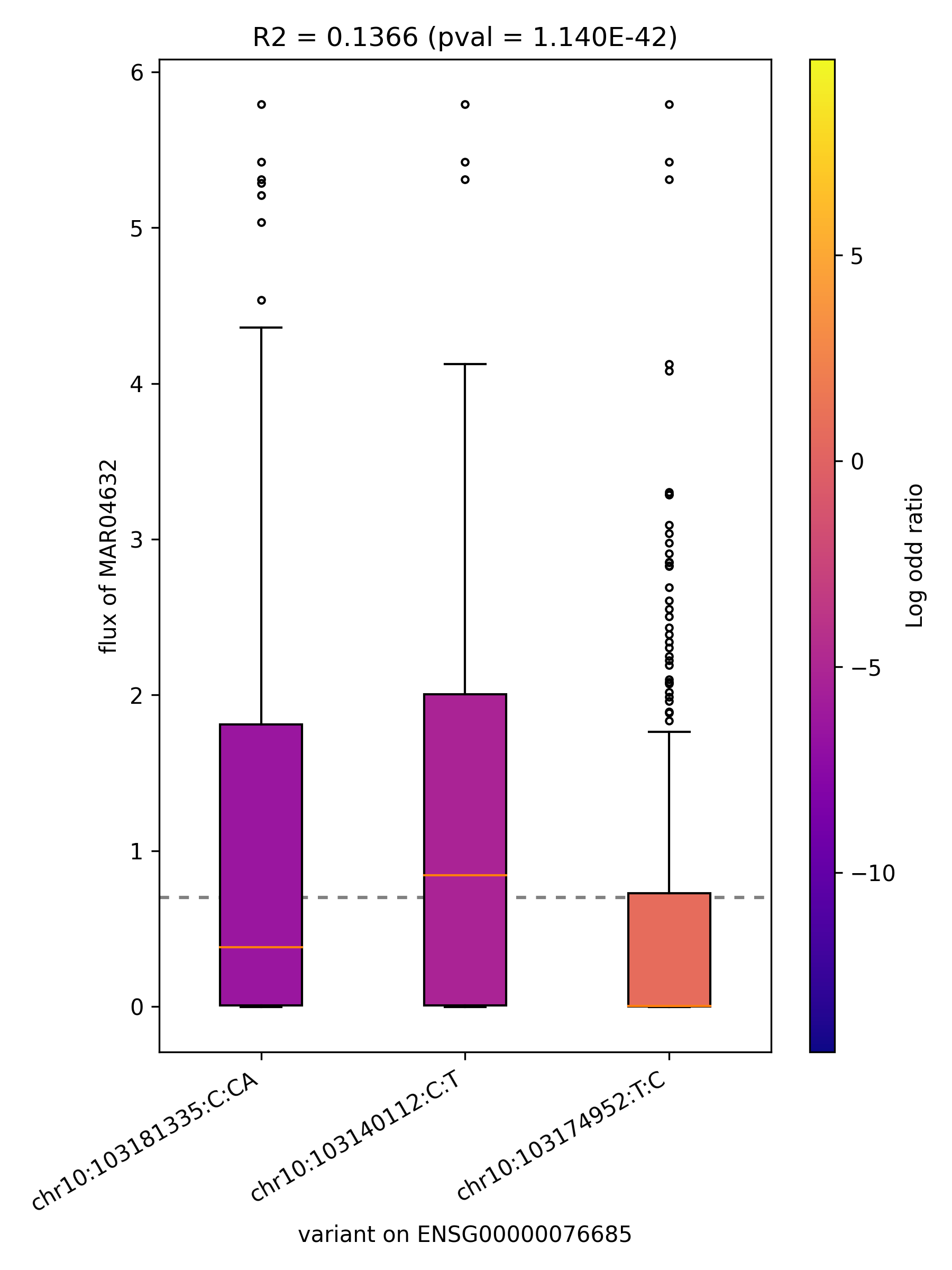

### MAR04632_ENSG00000116981.png

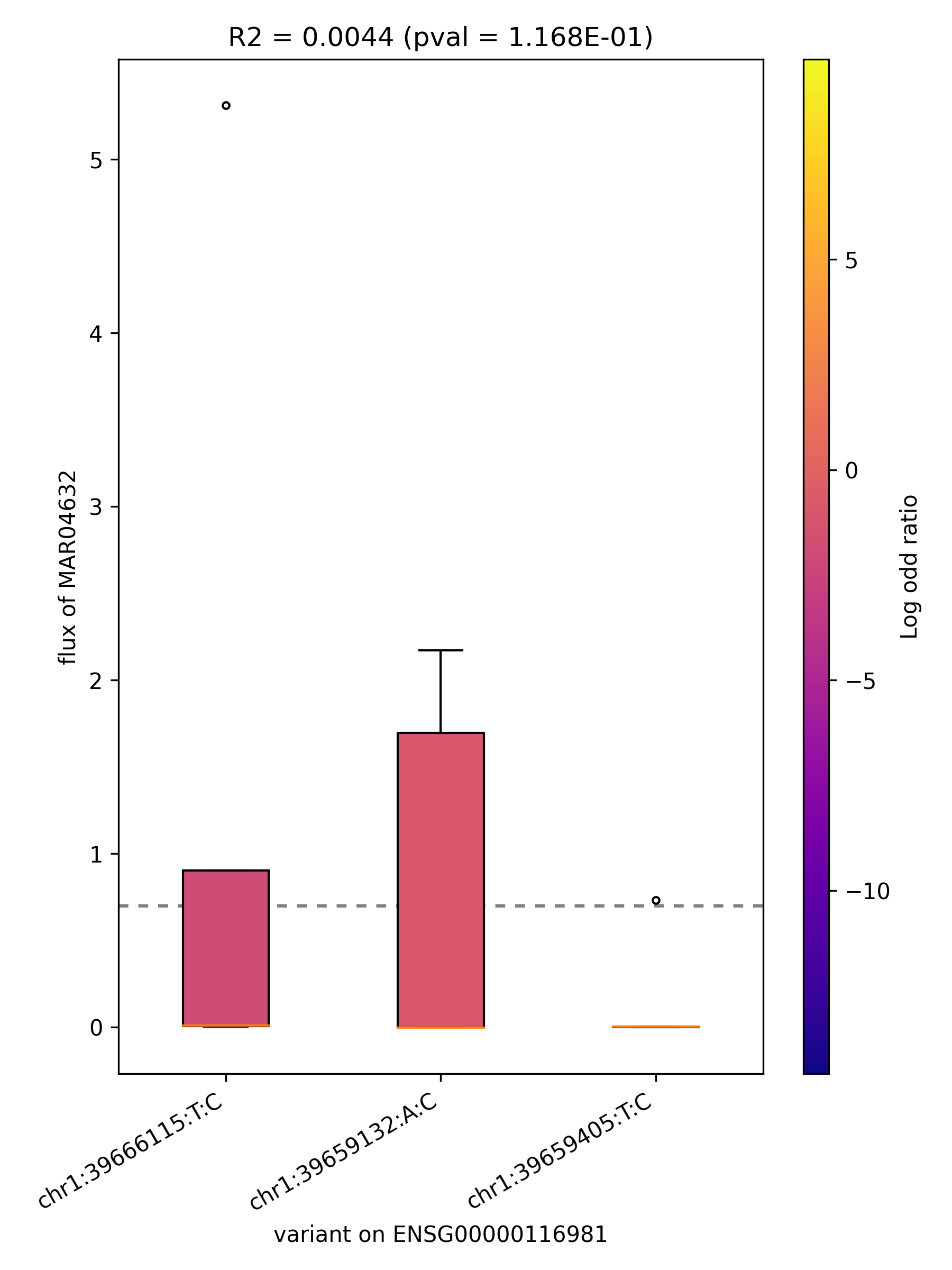

### MAR04632_ENSG00000135318.png

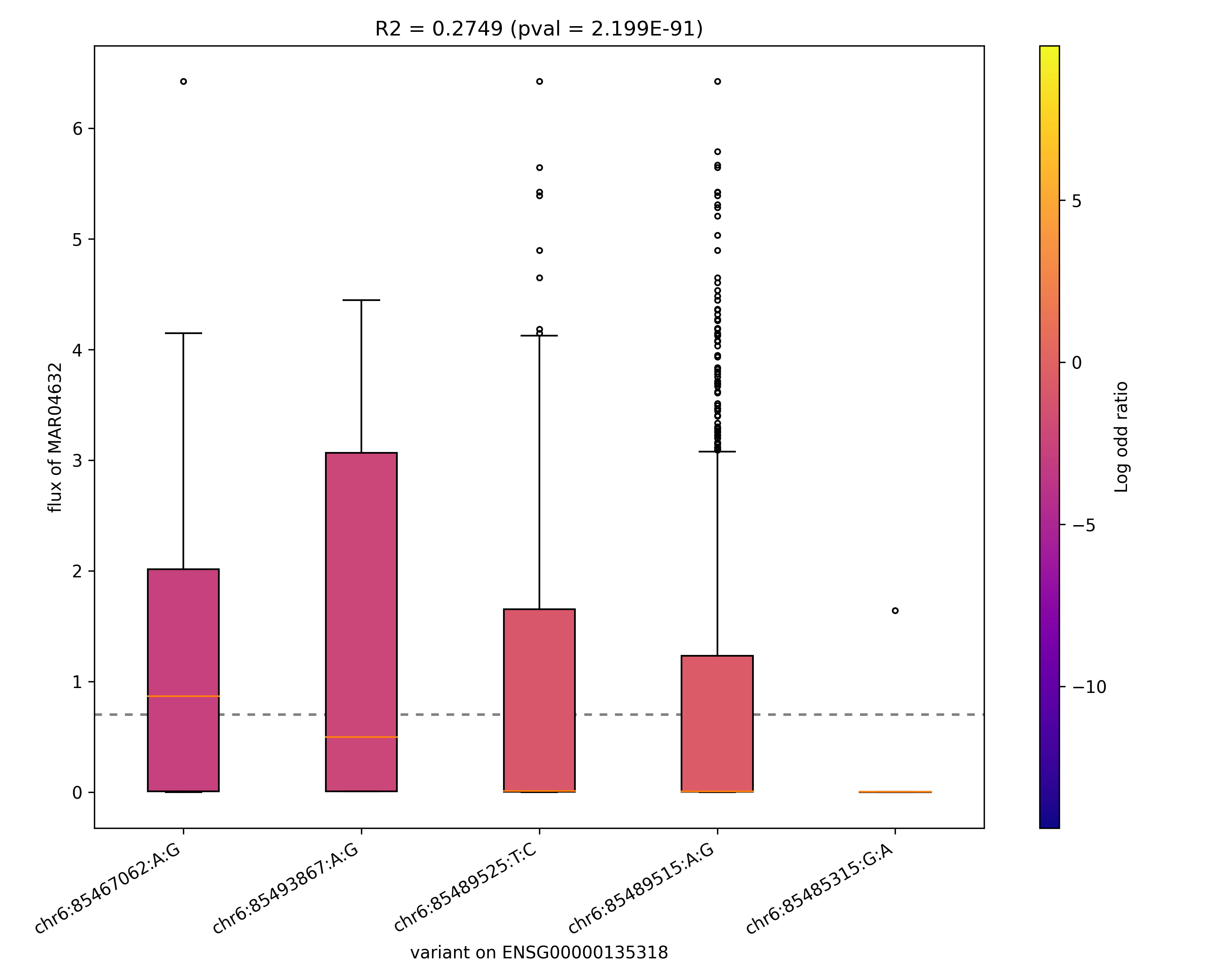

### MAR04632_ENSG00000141698.png

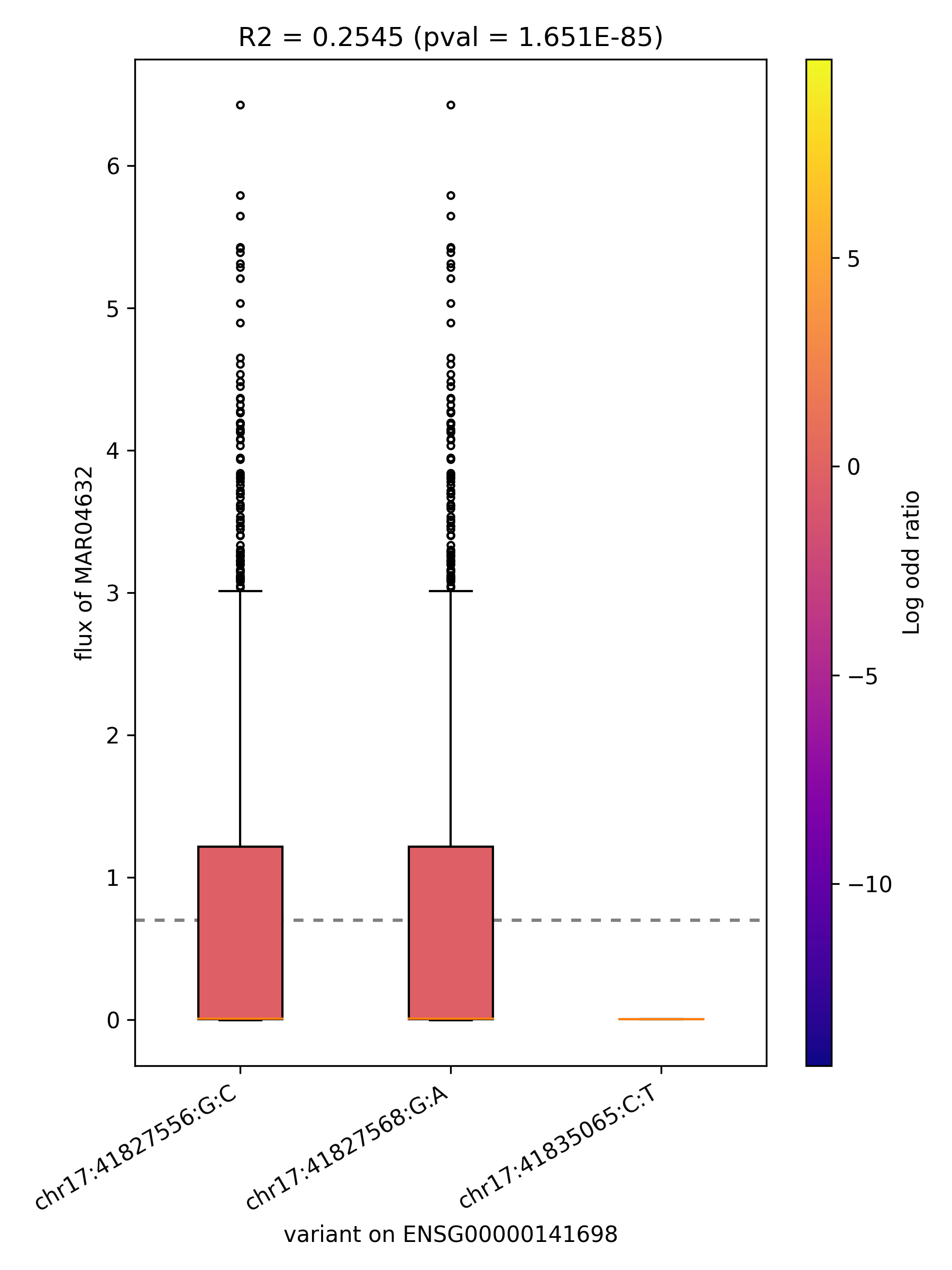

### MAR04632_ENSG00000146587.png

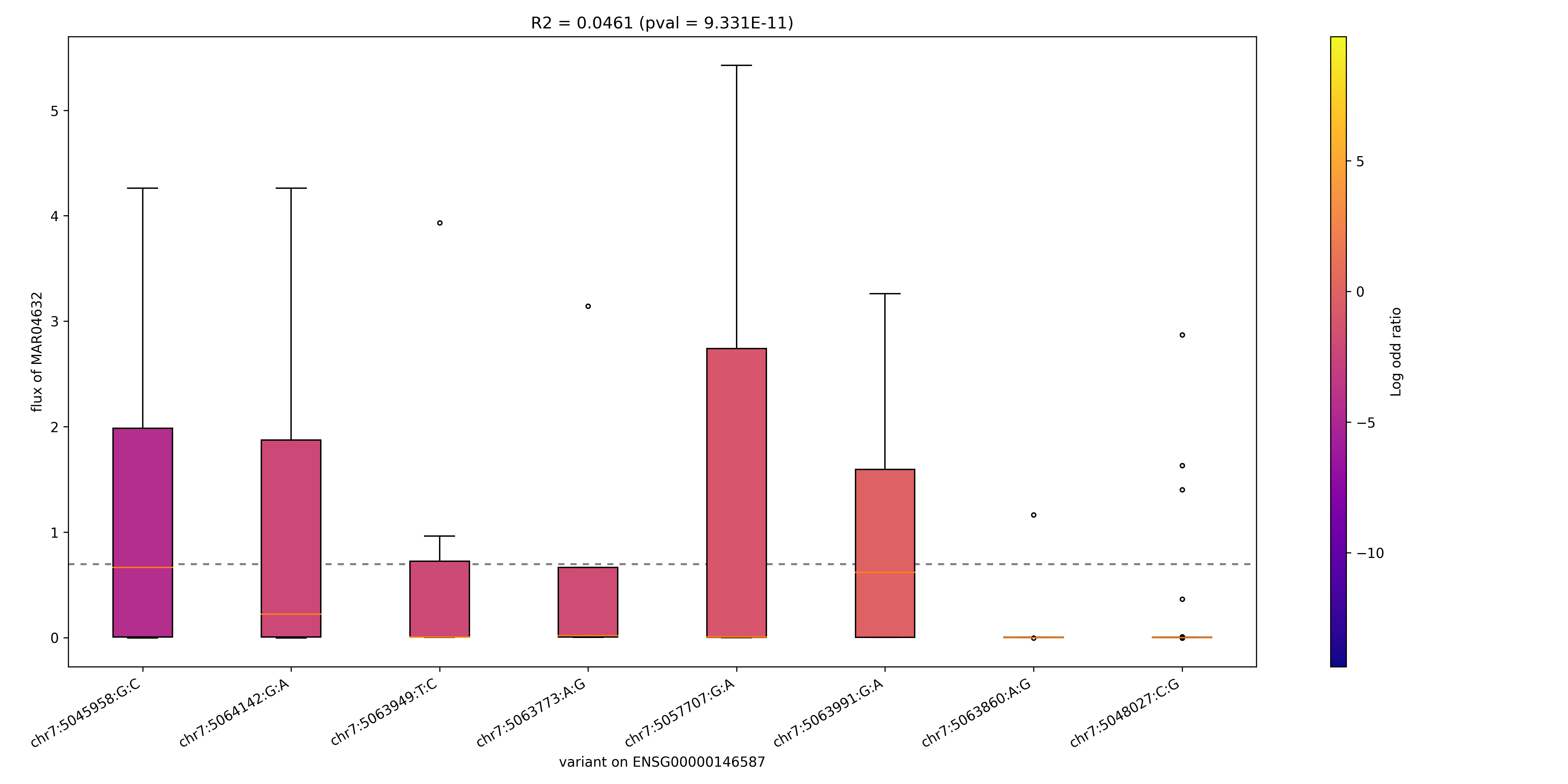

### MAR04660_ENSG00000139344.png

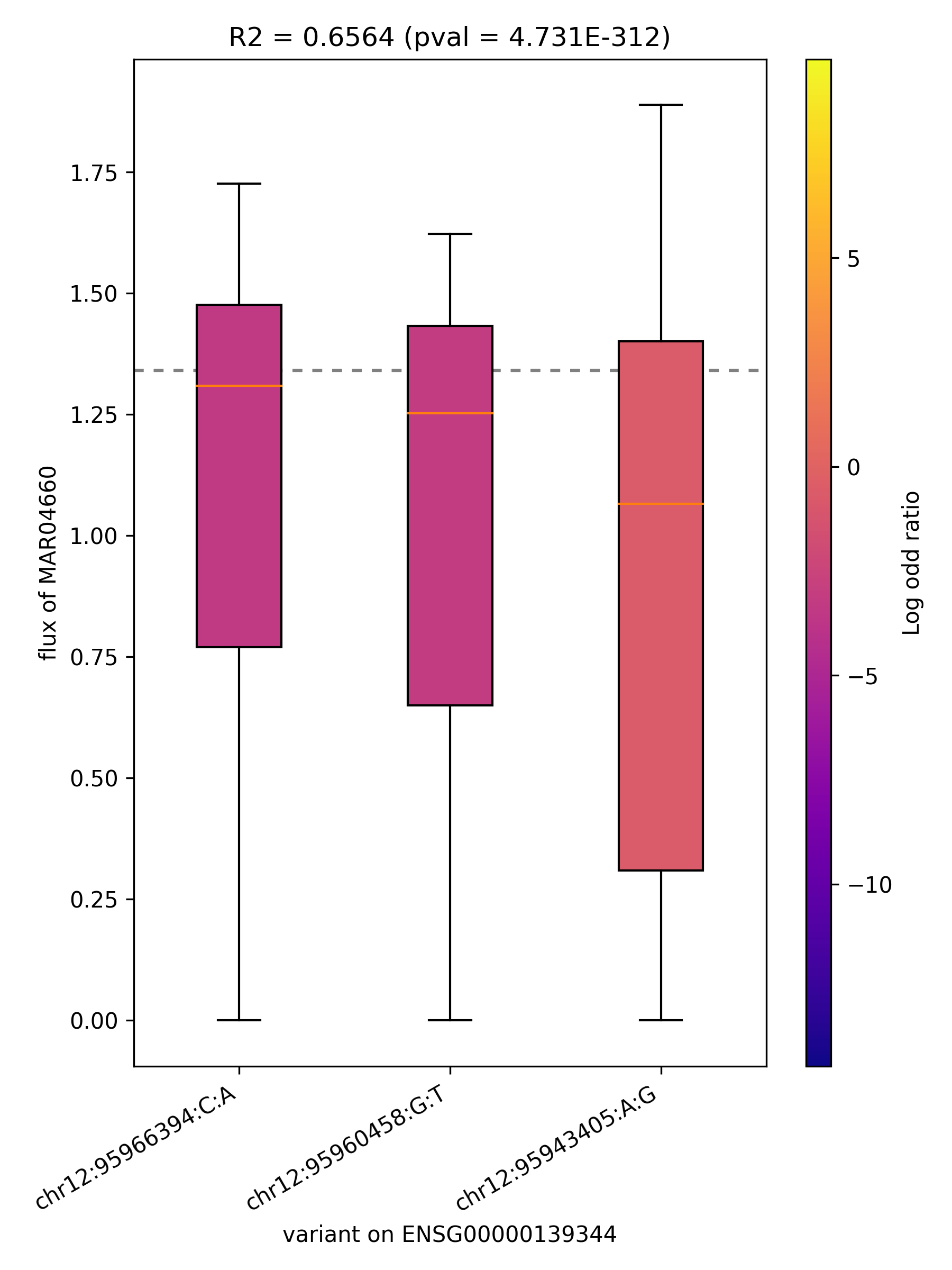

### MAR04676_ENSG00000130717.png

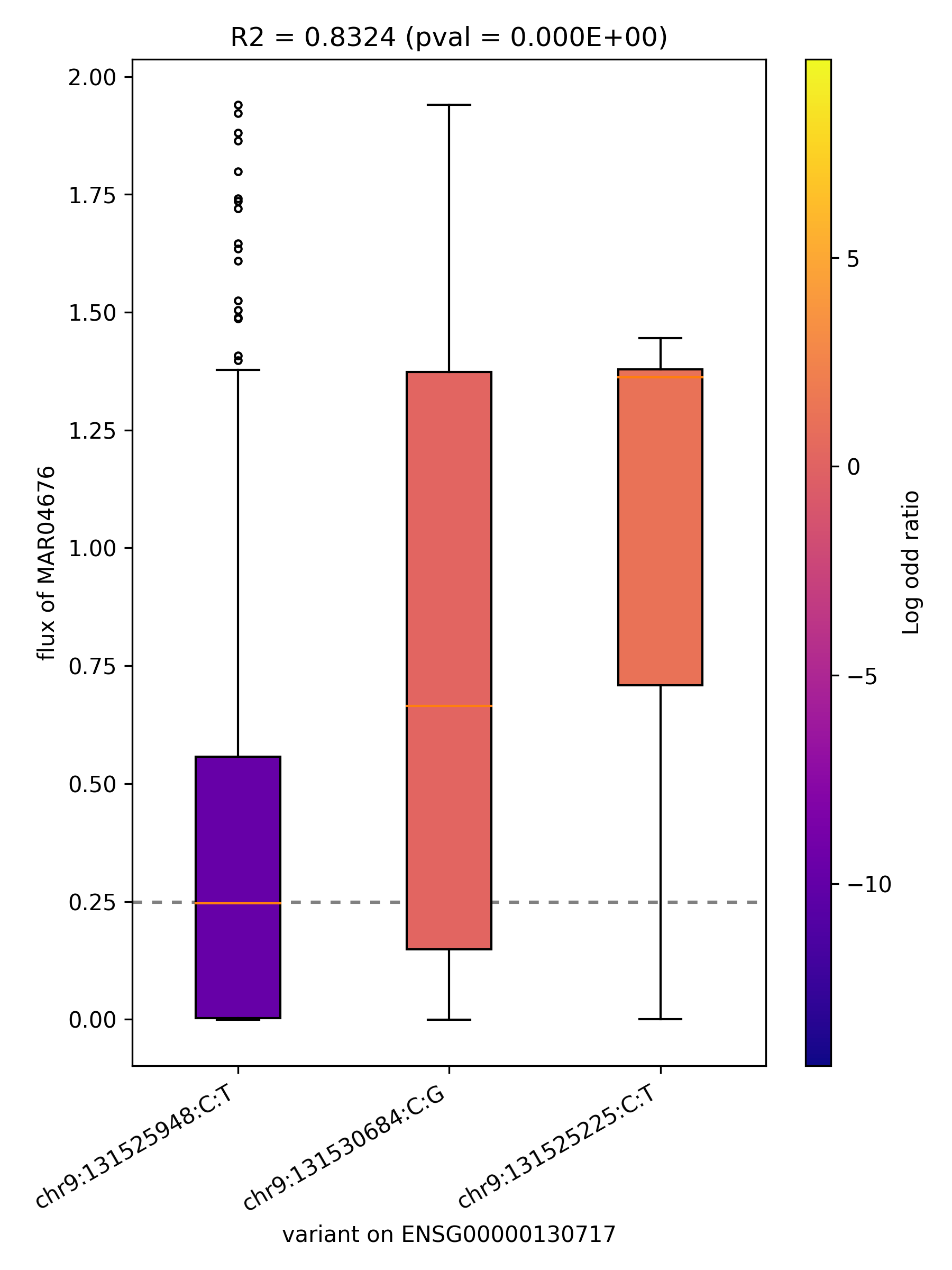

### MAR04676_ENSG00000198276.png

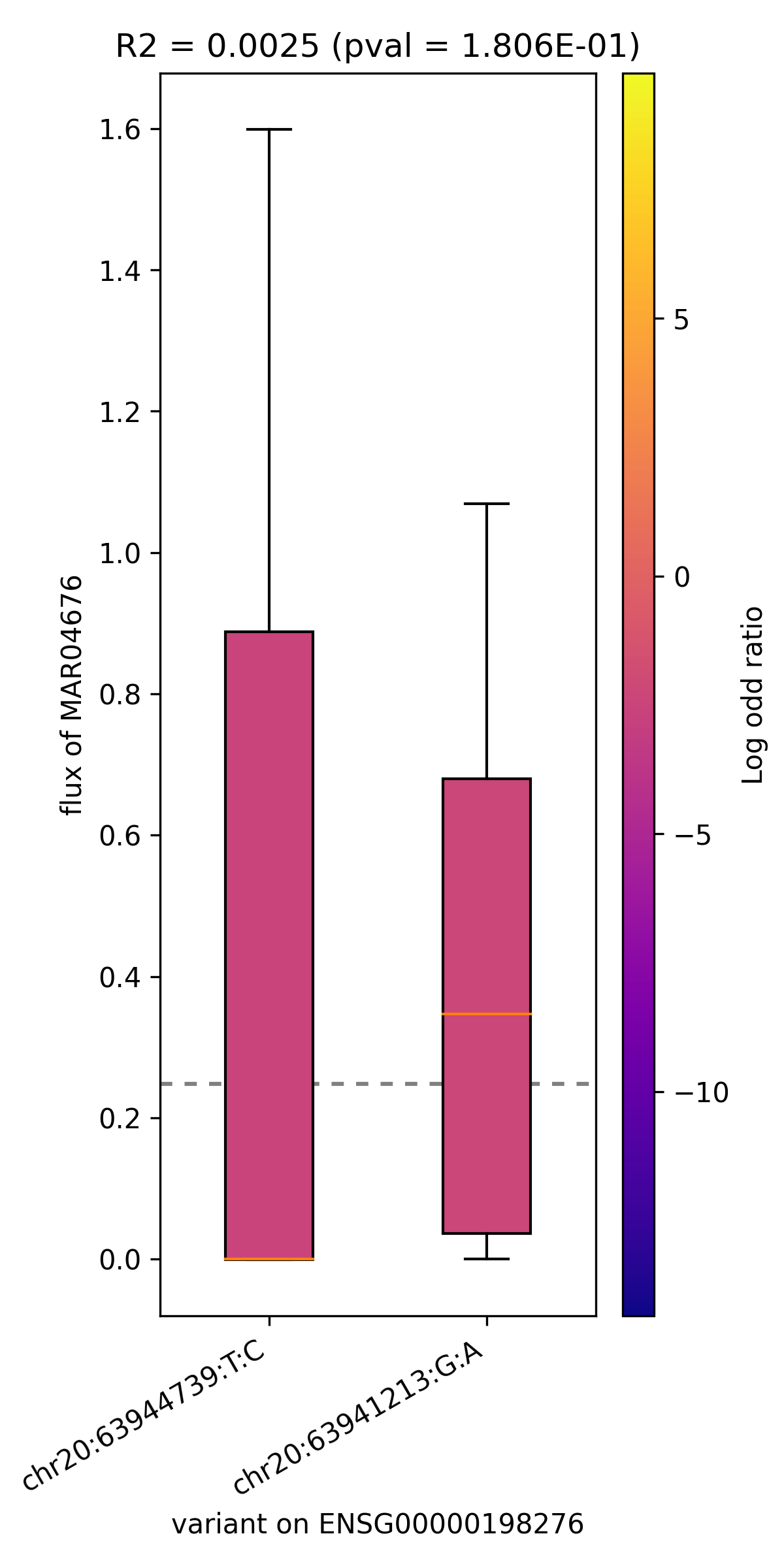

### MAR04973_ENSG00000059804.png

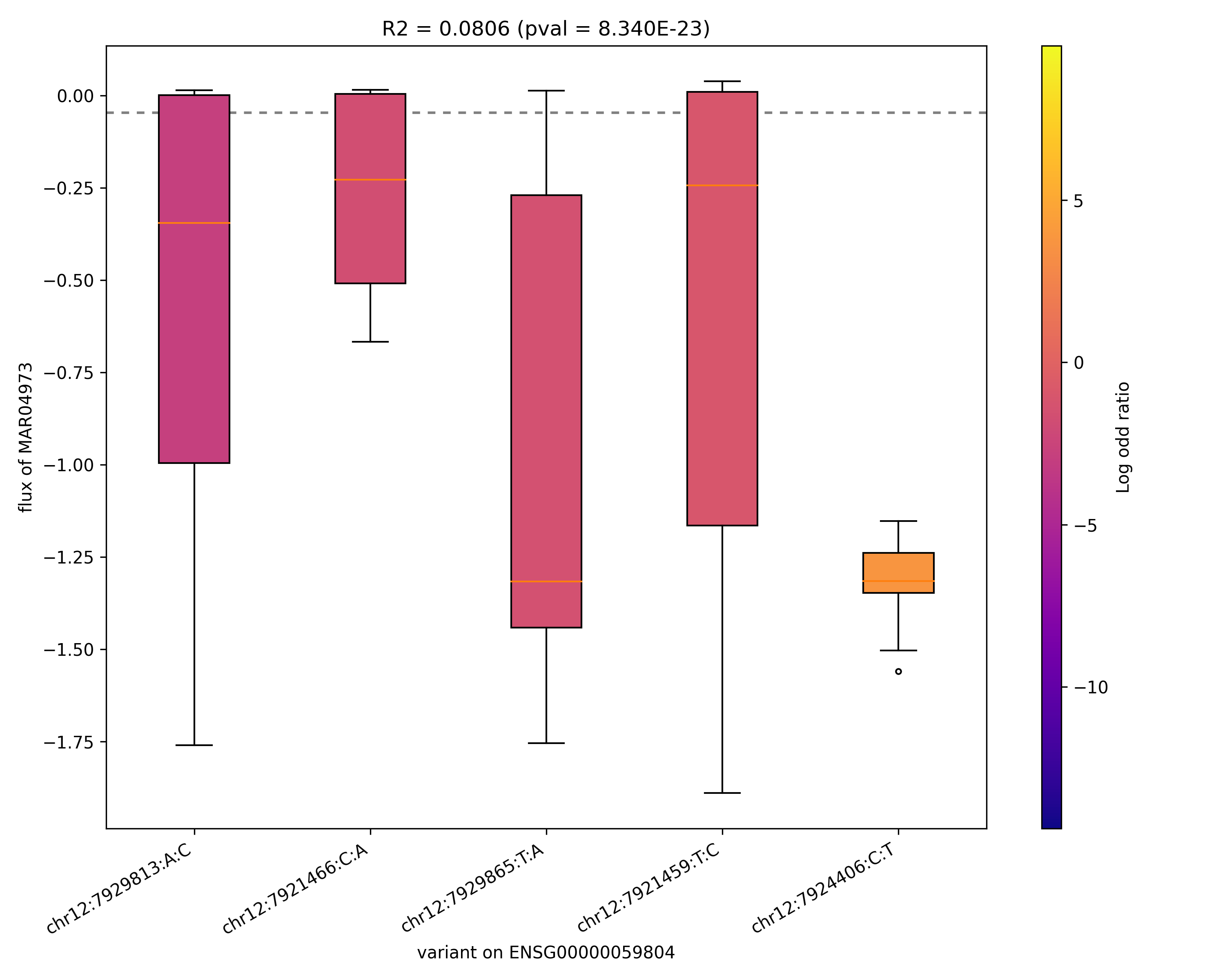

### MAR04973_ENSG00000163581.png

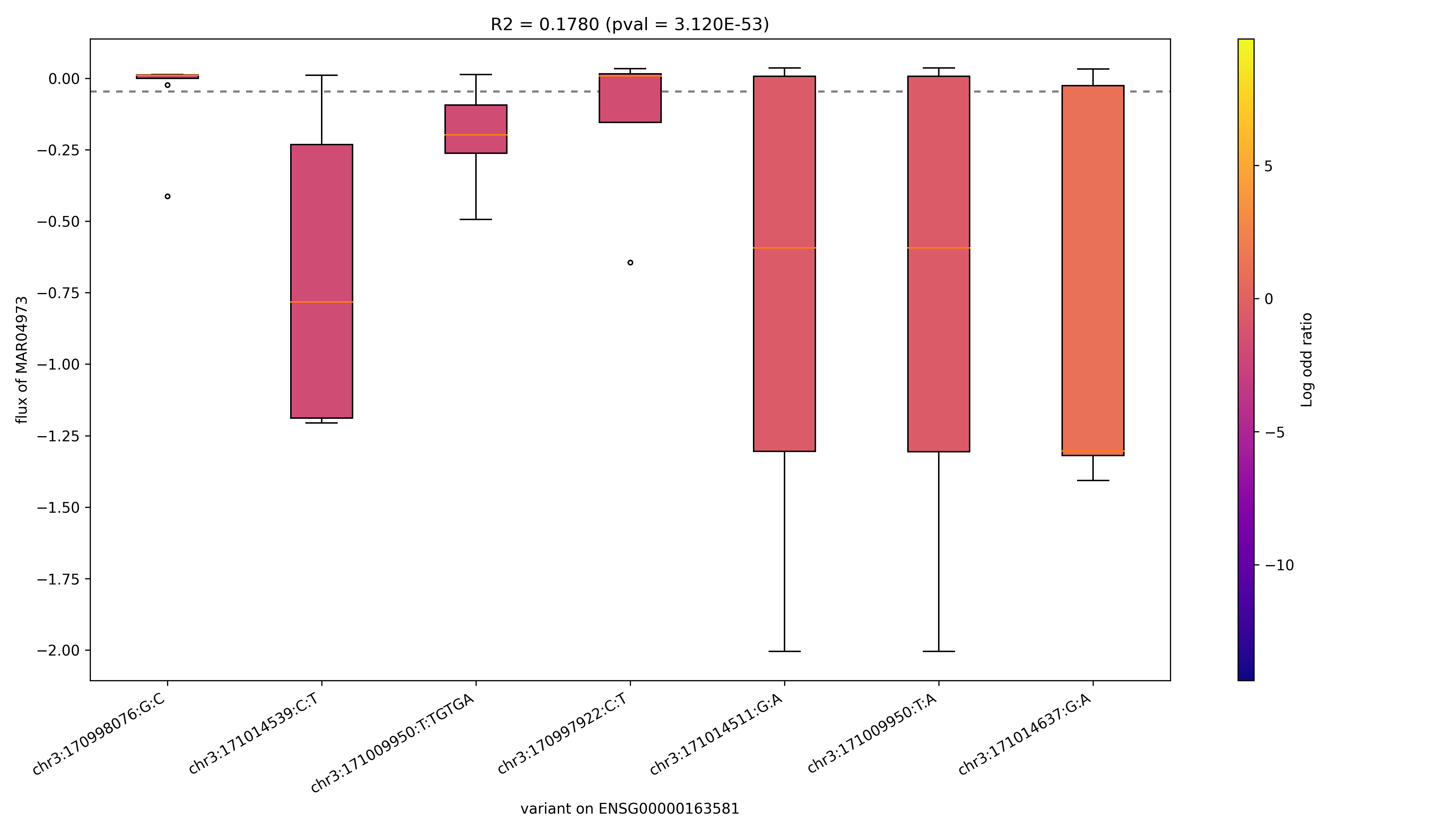

### MAR05301_ENSG00000198276.png

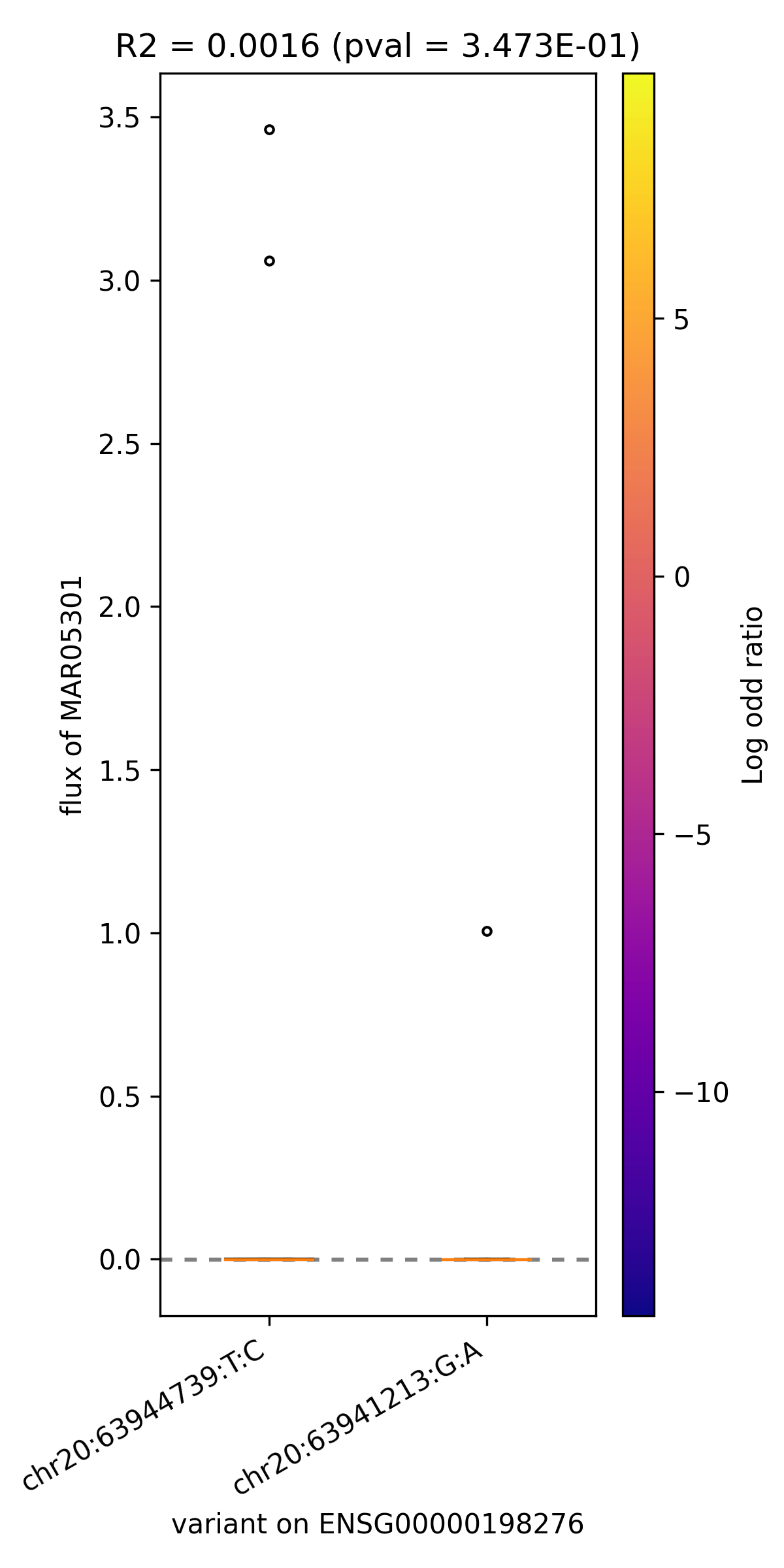

### MAR05456_ENSG00000110090.png

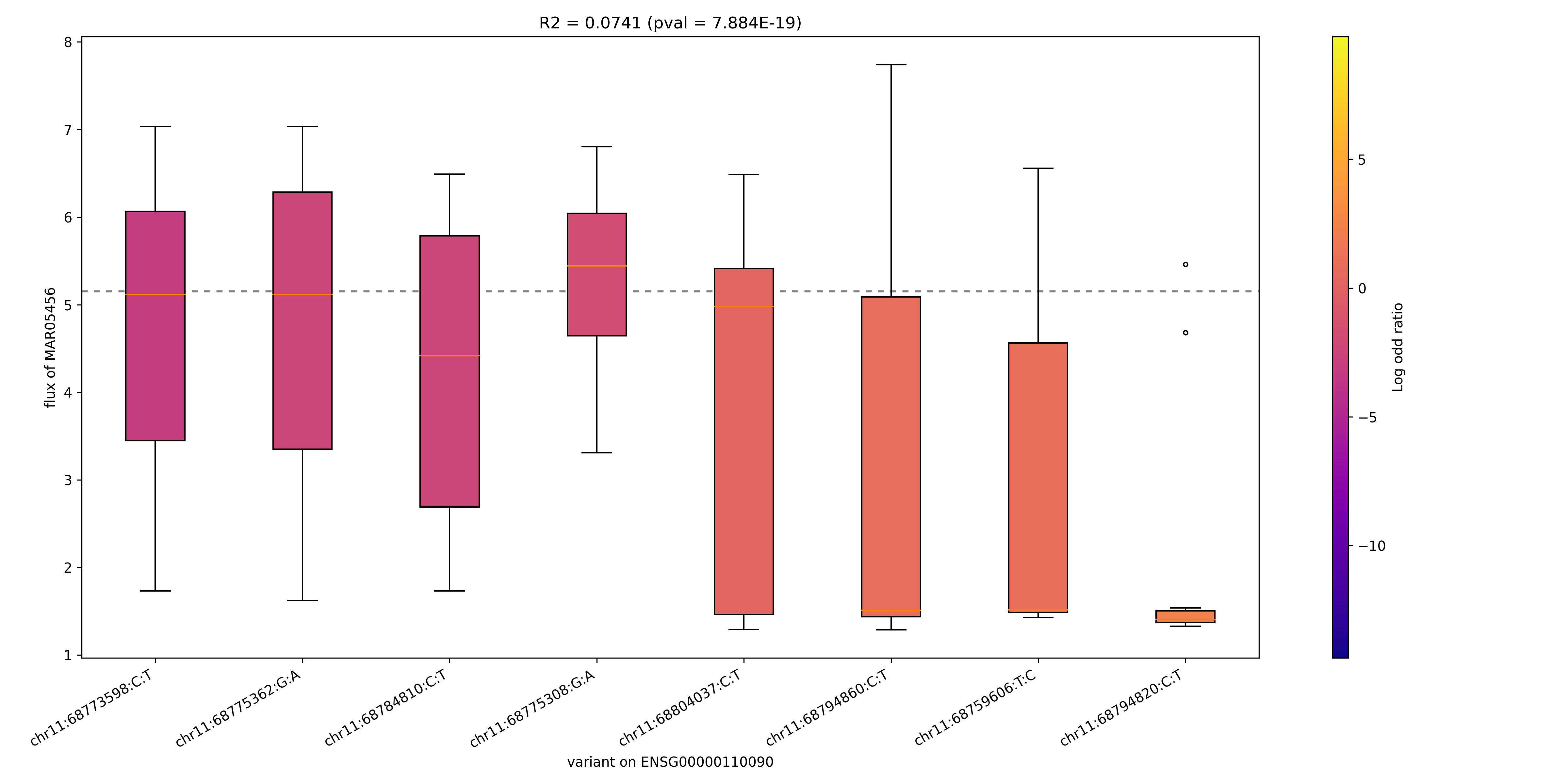

### MAR05456_ENSG00000169169.png

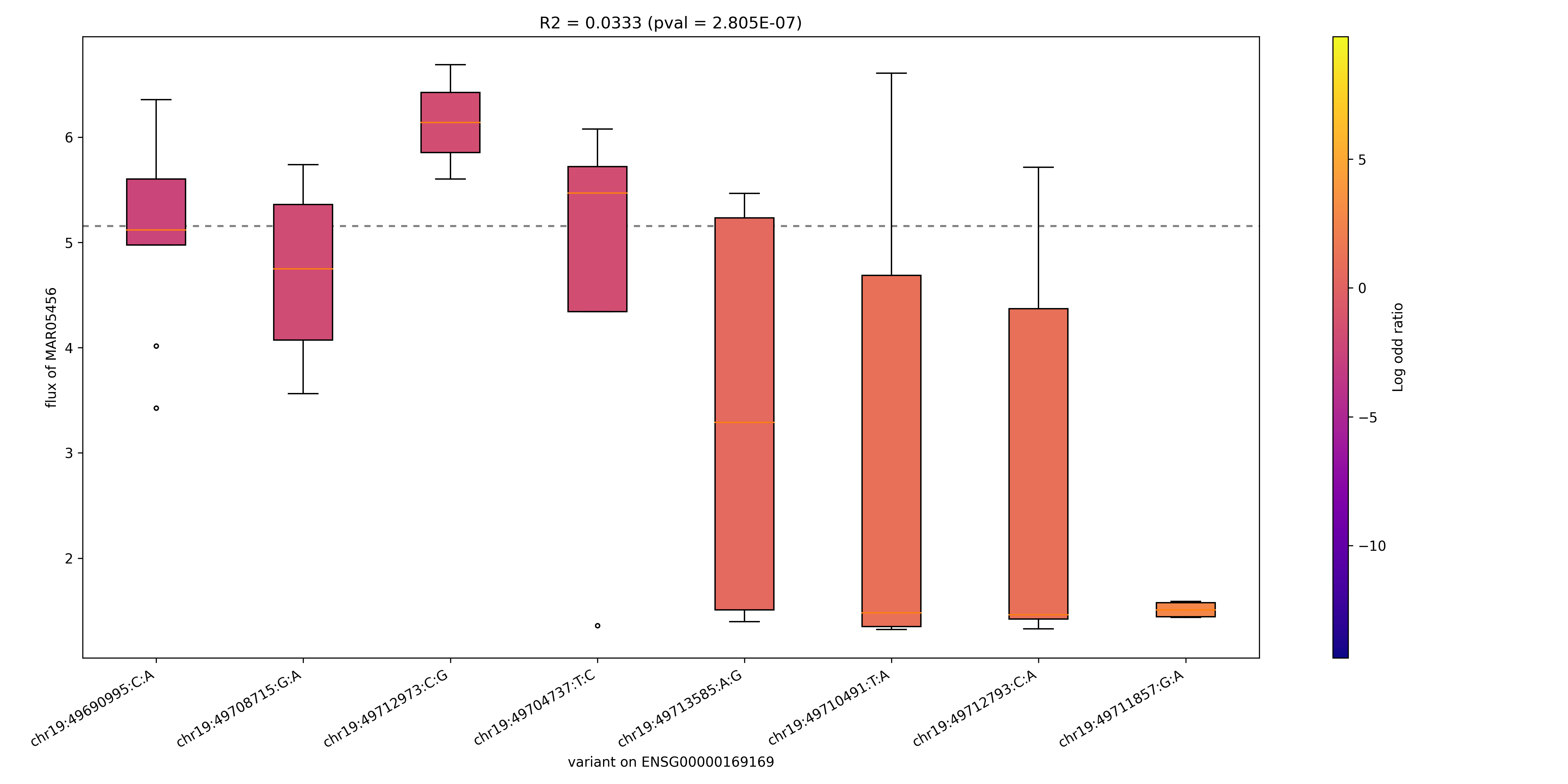

### MAR06278_ENSG00000205560.png

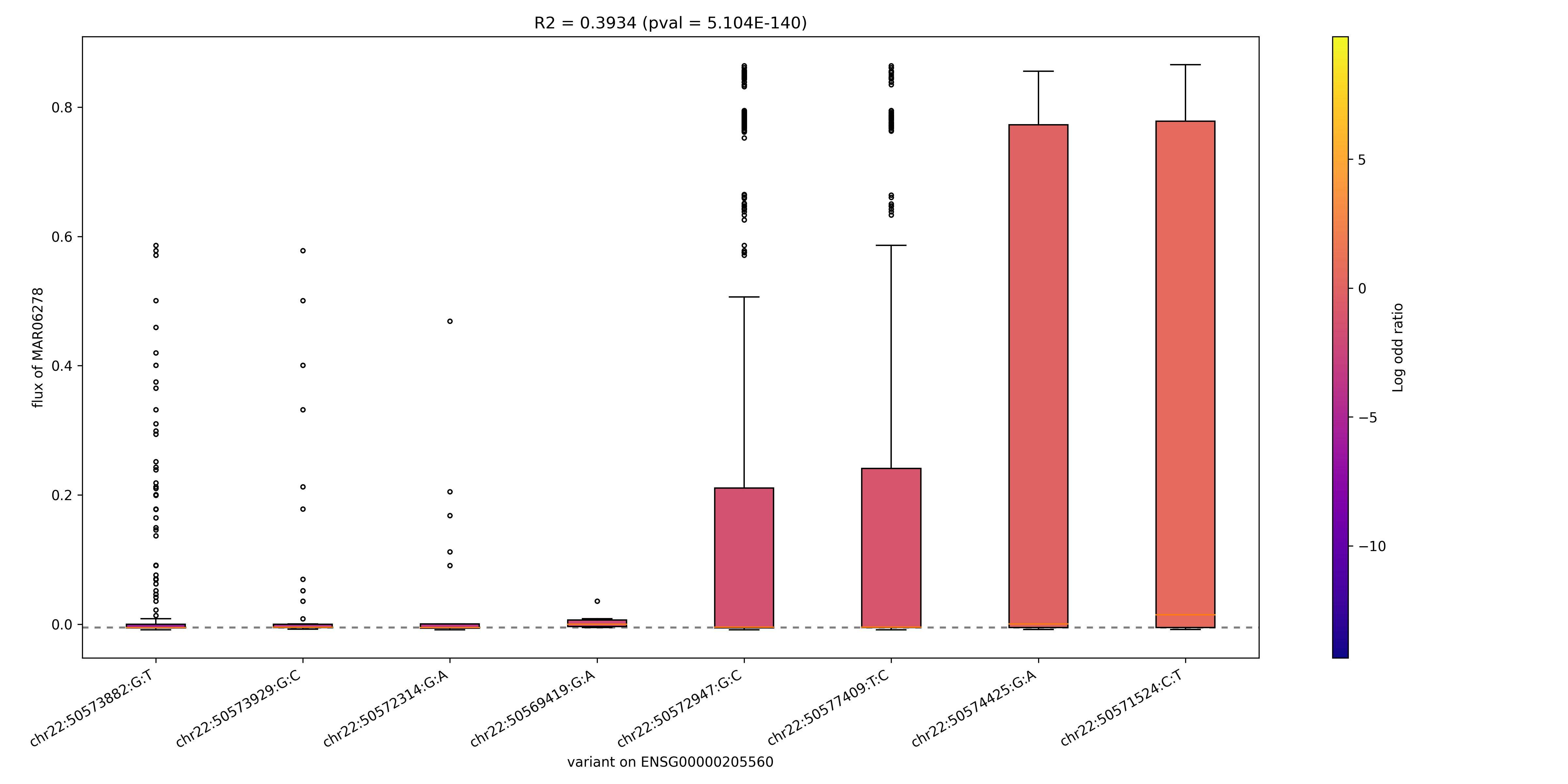

### MAR06533_ENSG00000225697.png

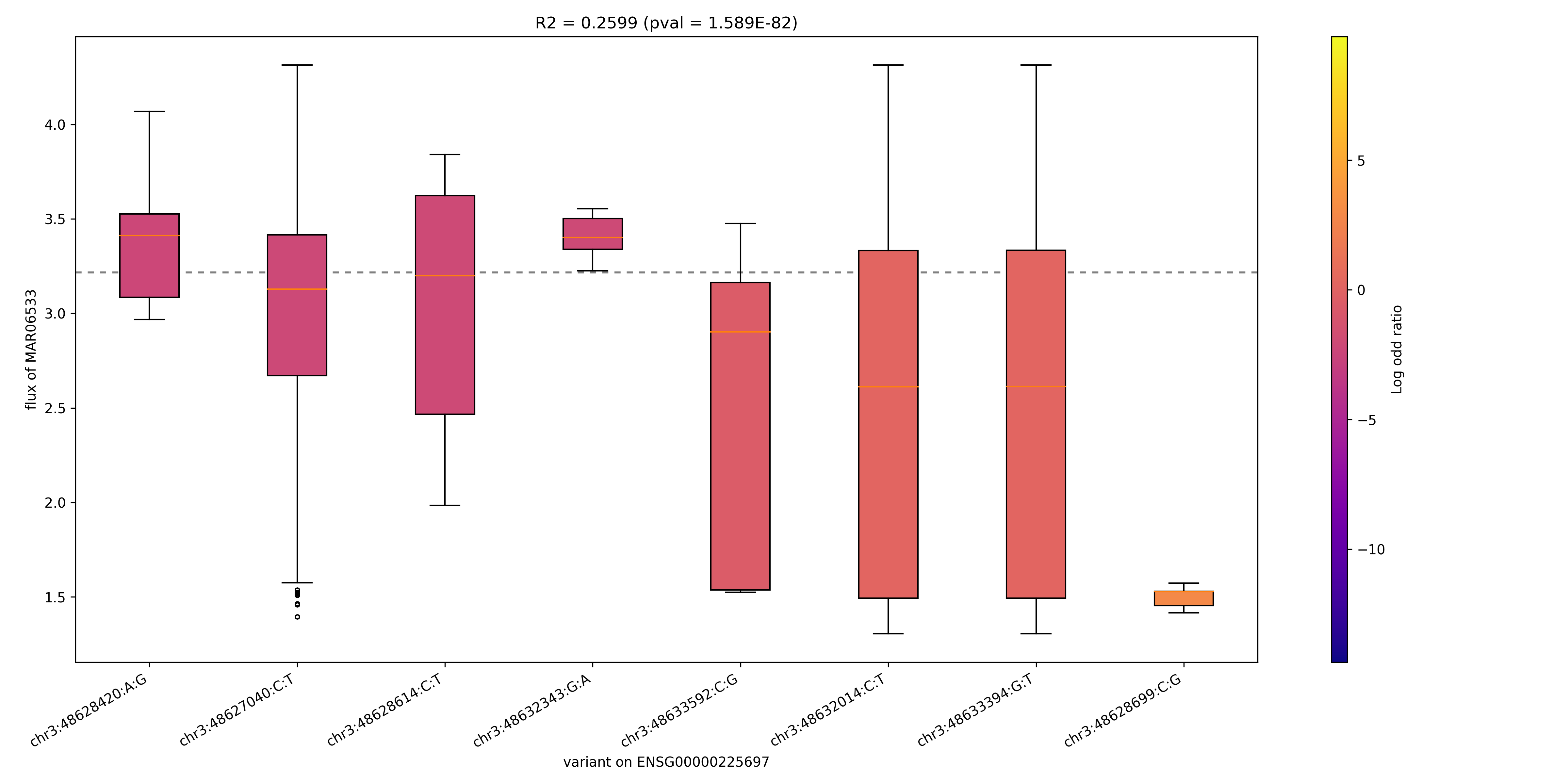

### MAR07143_ENSG00000109743.png

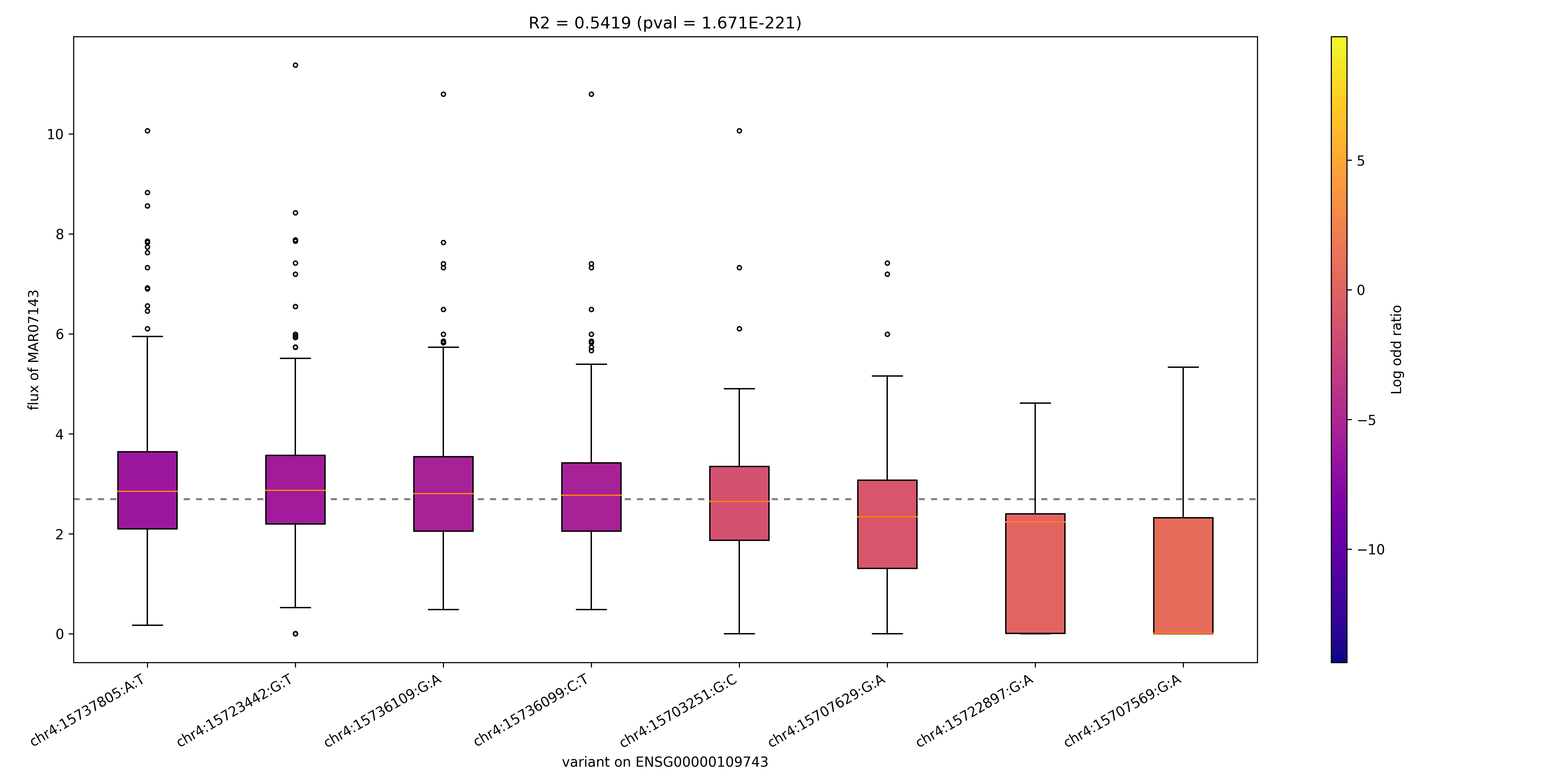

### MAR07144_ENSG00000142185.png

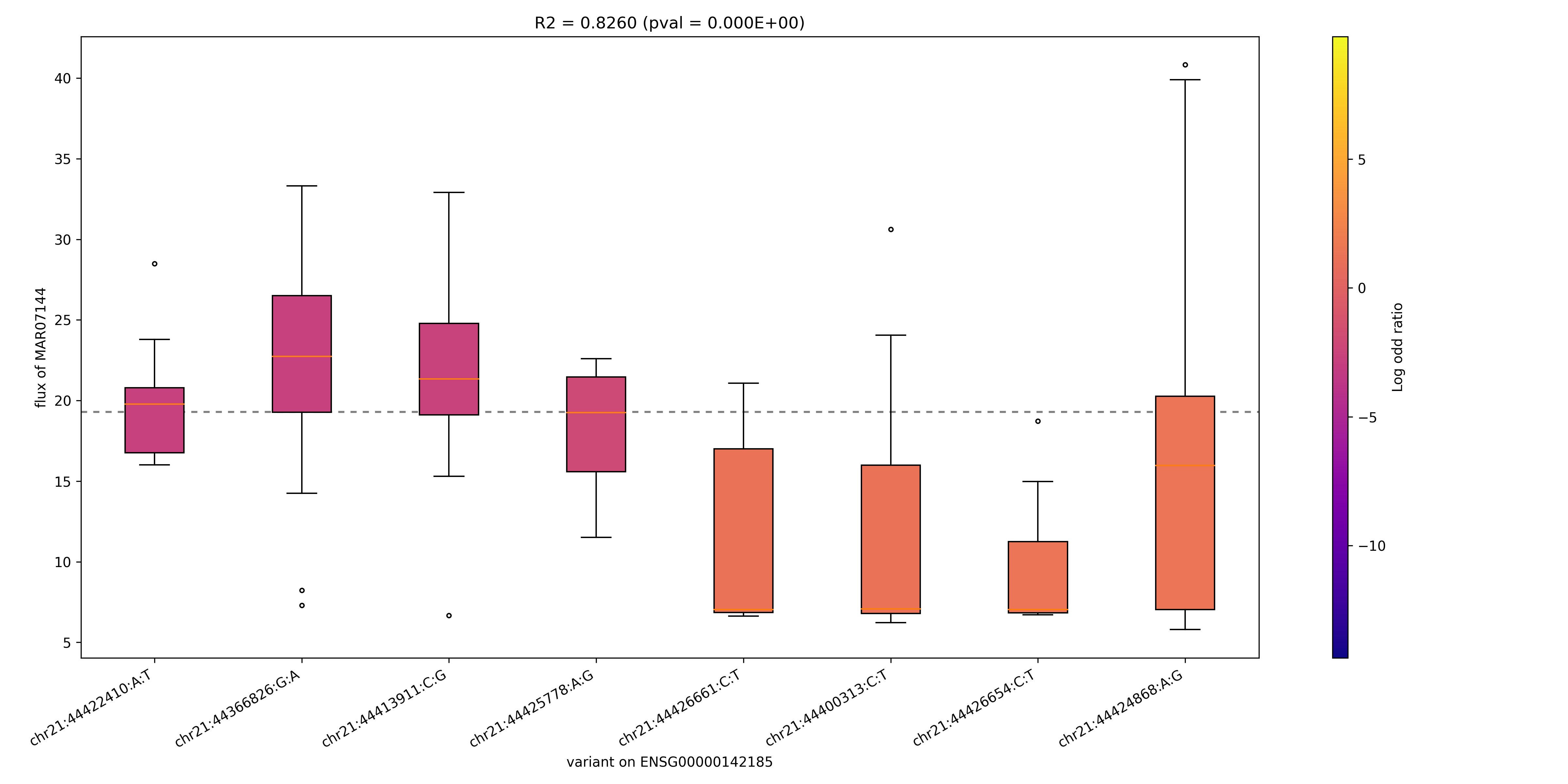

### MAR07144_ENSG00000170222.png

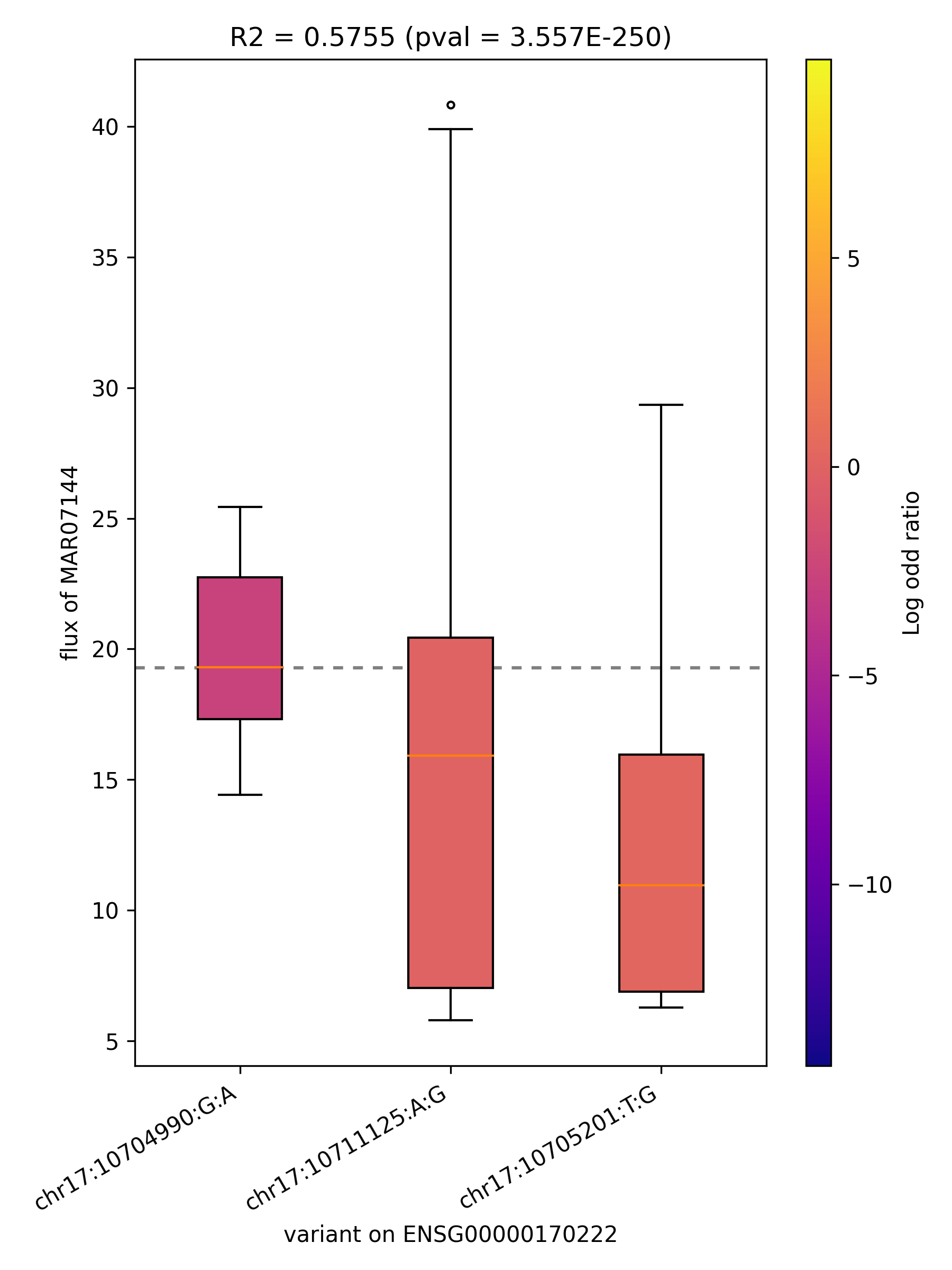

### MAR07623_ENSG00000077463.png

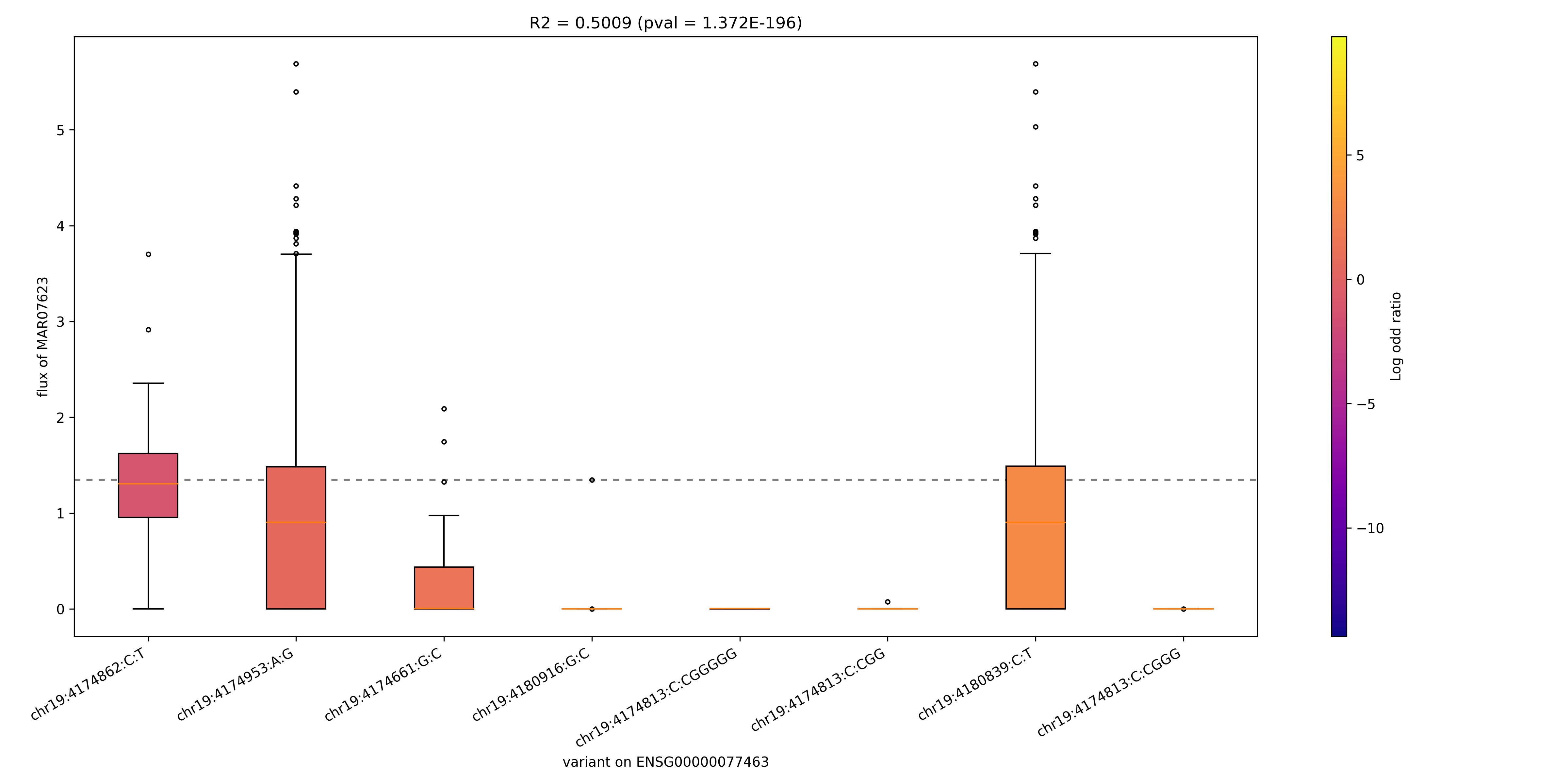

### MAR07623_ENSG00000129744.png

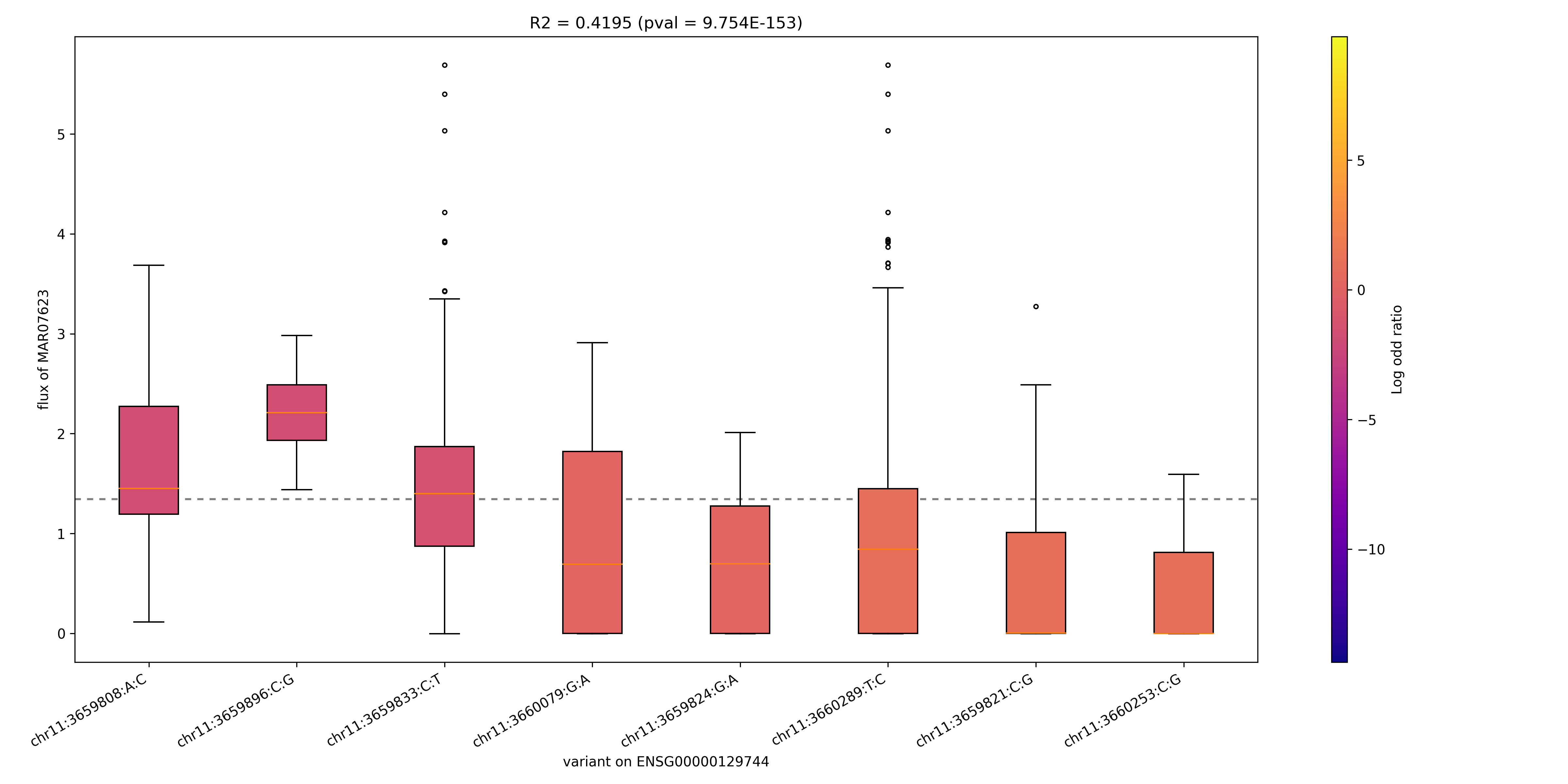

### MAR07623_ENSG00000156219.png

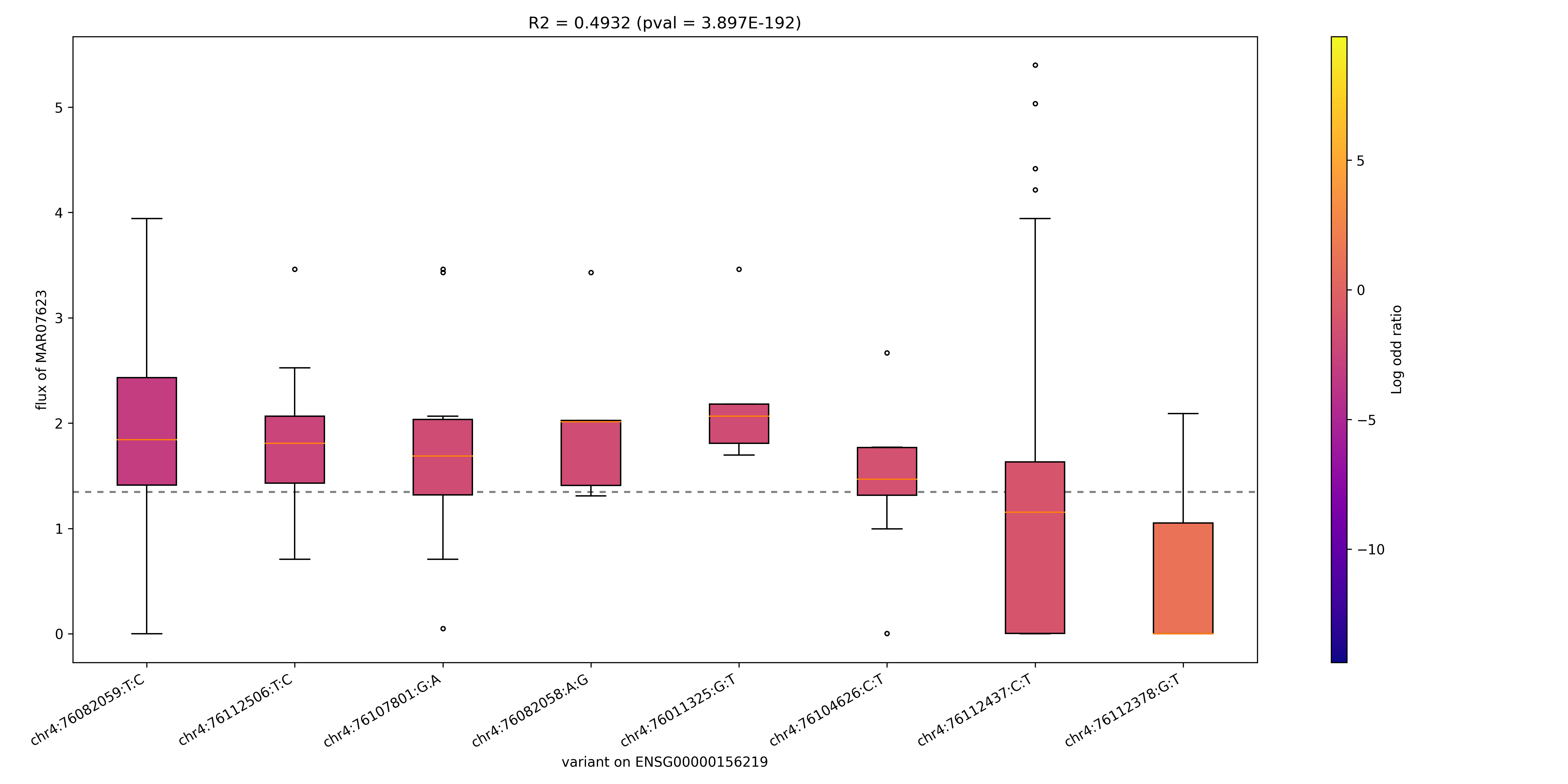

### MAR07623_ENSG00000167311.png

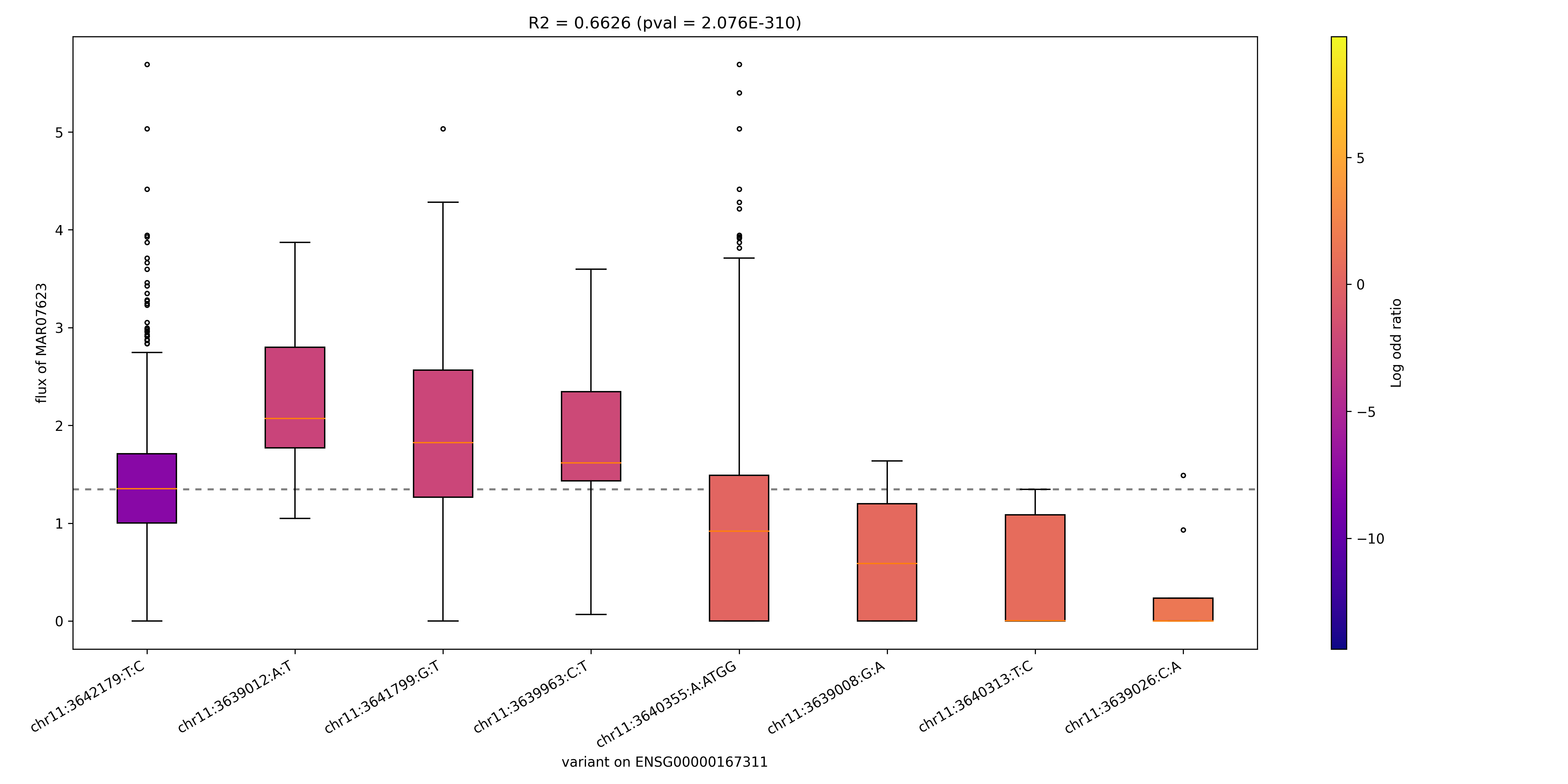

### MAR07625_ENSG00000077463.png

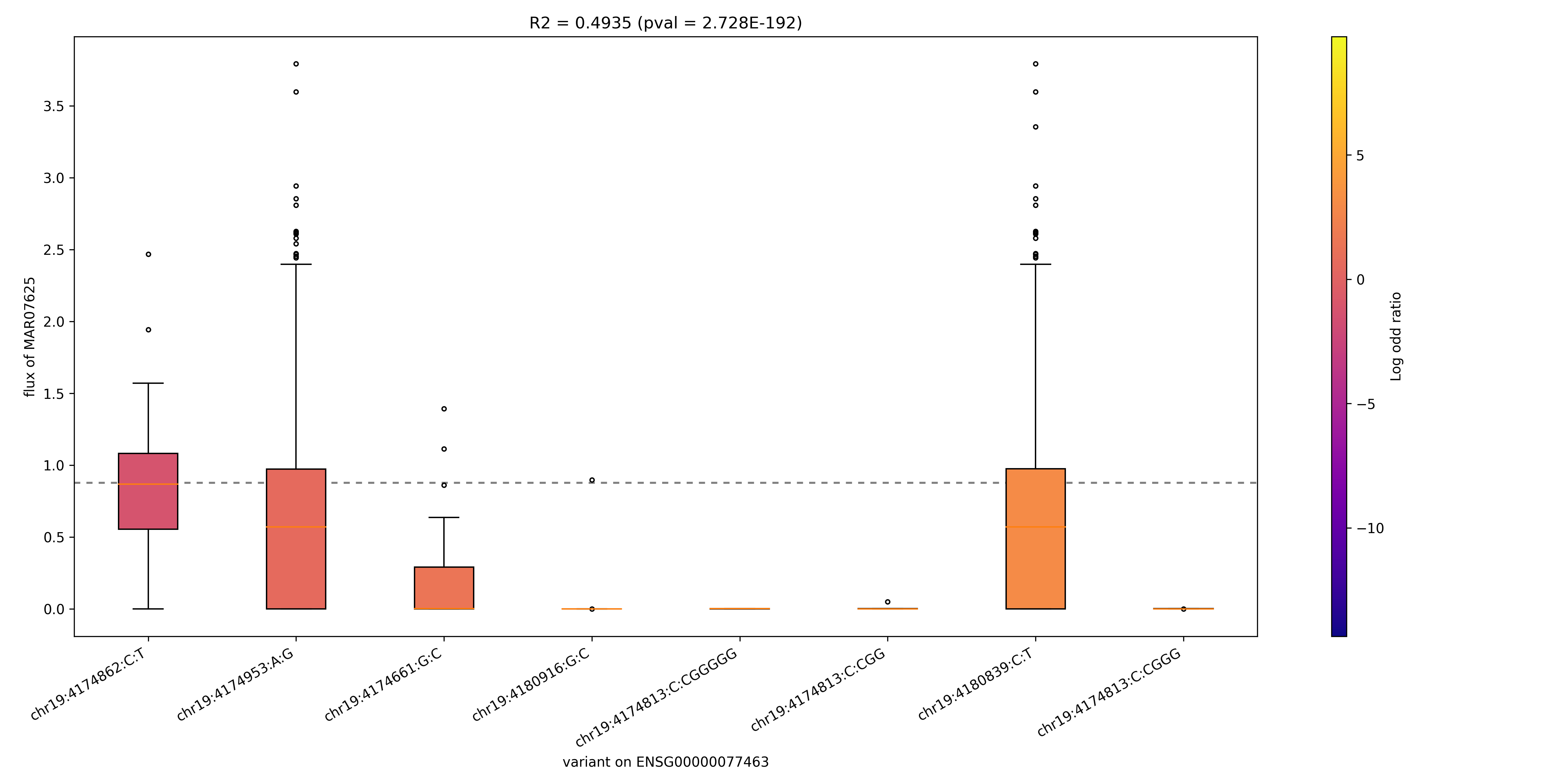

### MAR07625_ENSG00000129744.png

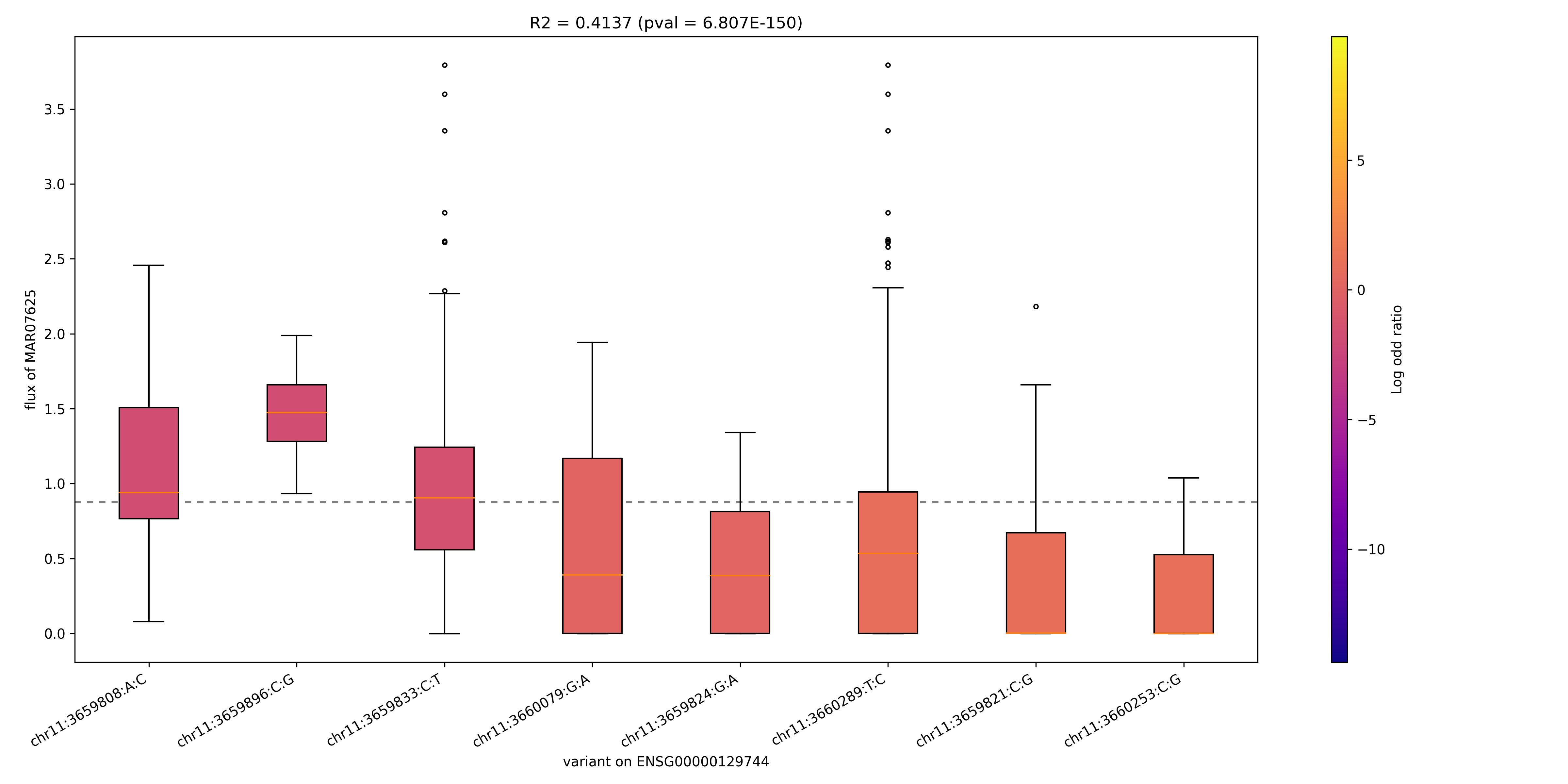

### MAR07625_ENSG00000156219.png

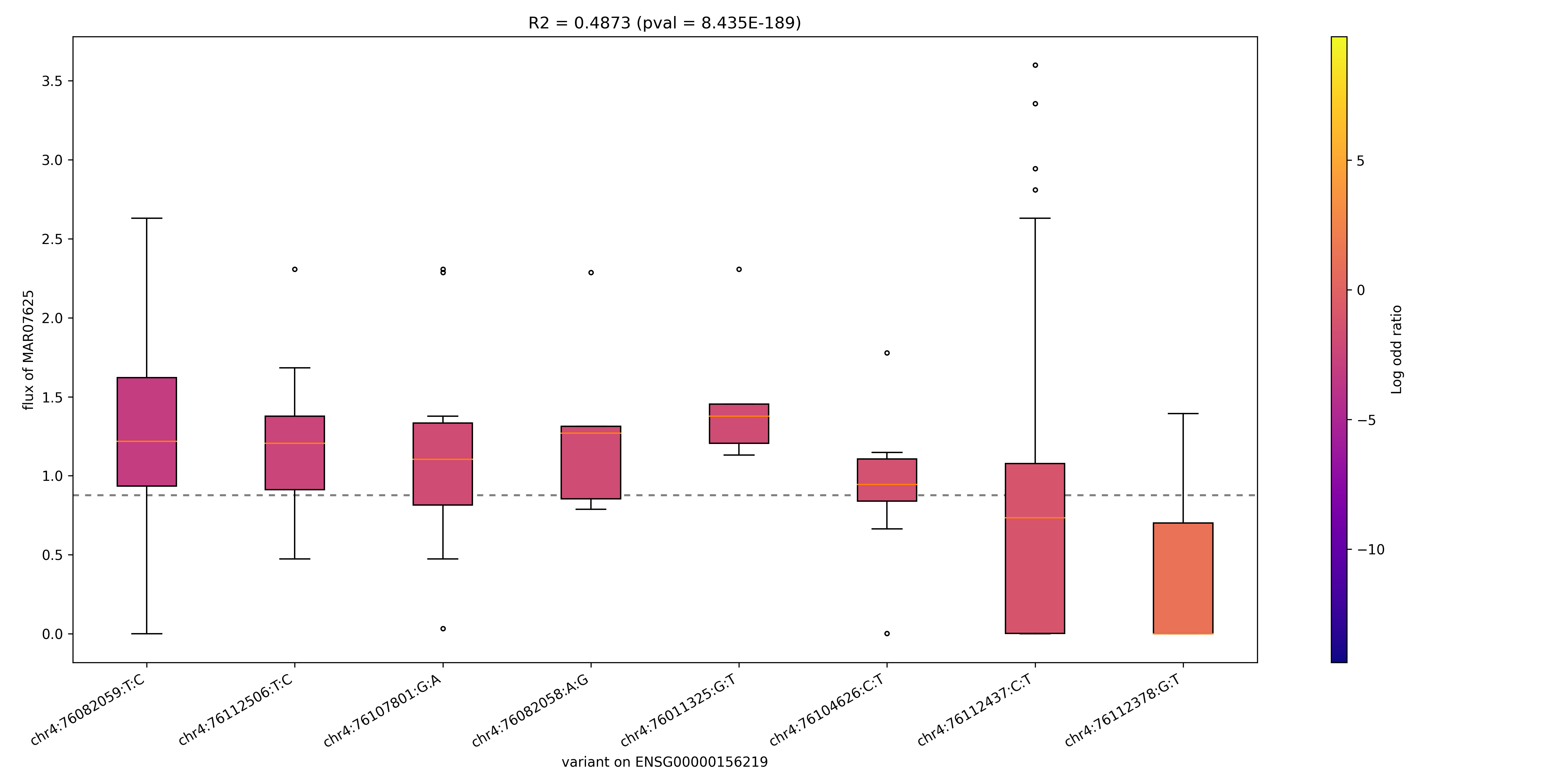

### MAR07625_ENSG00000167311.png

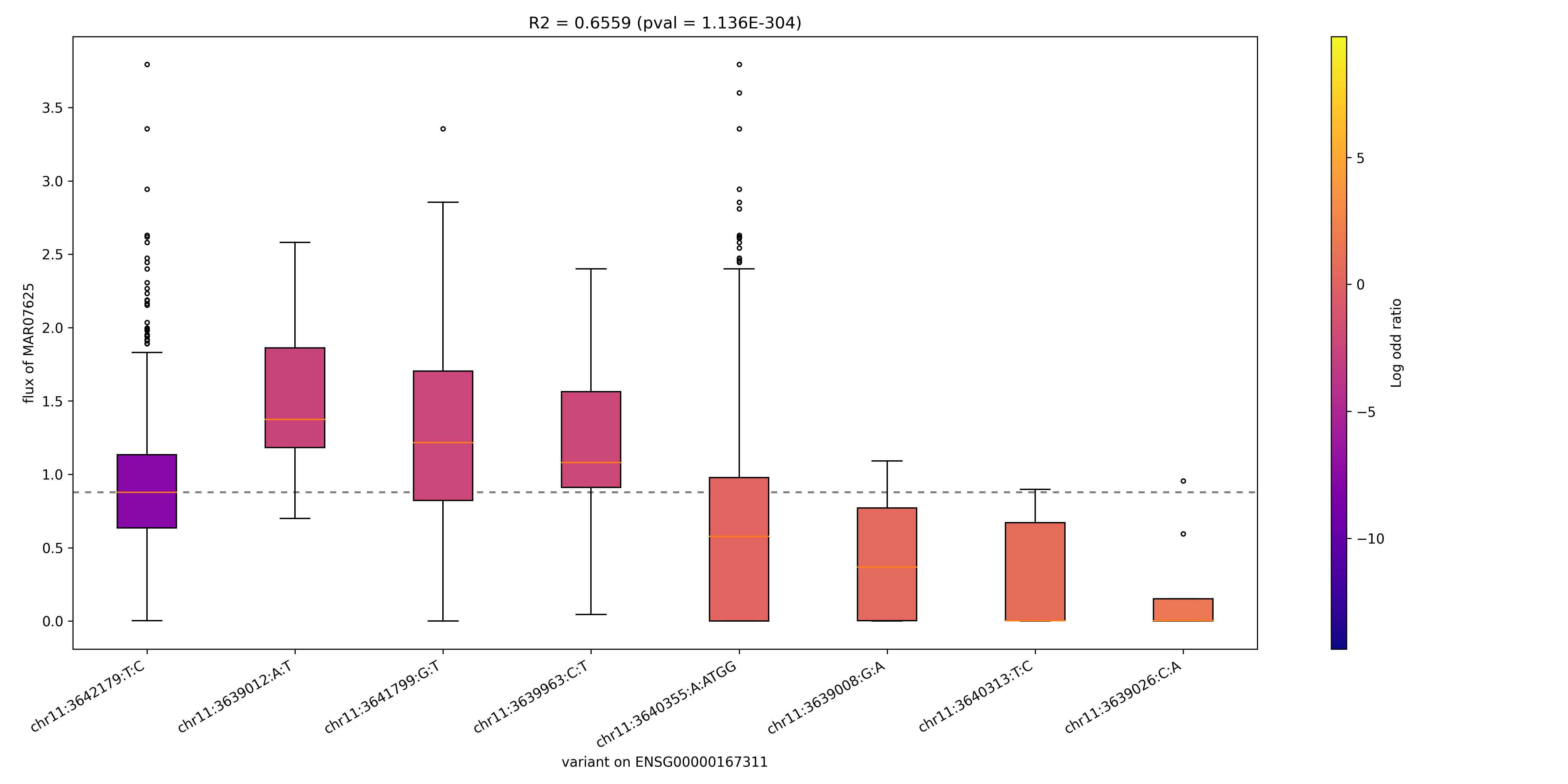

### MAR07687_ENSG00000173638.png

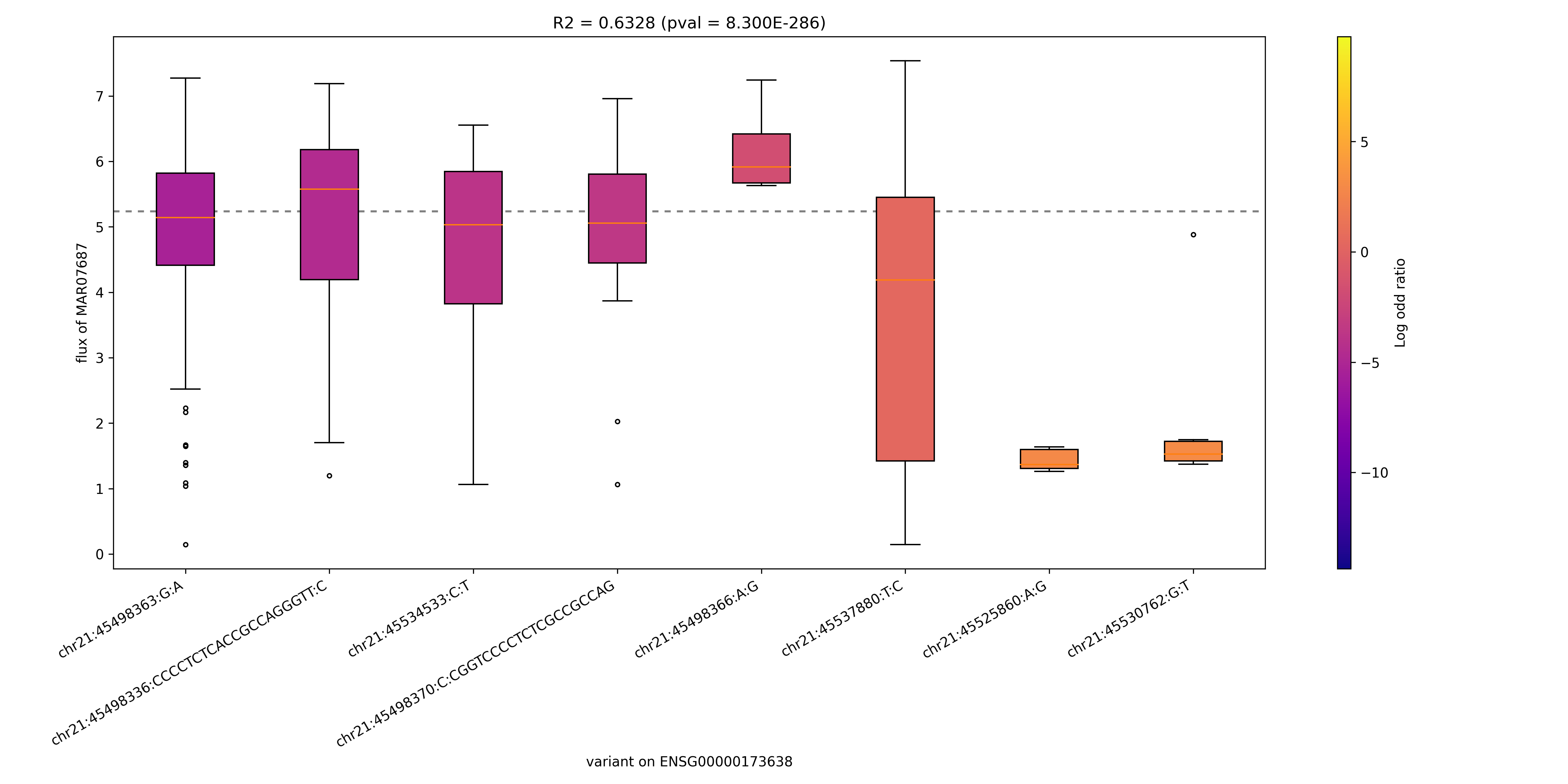

### MAR08487_ENSG00000135318.png

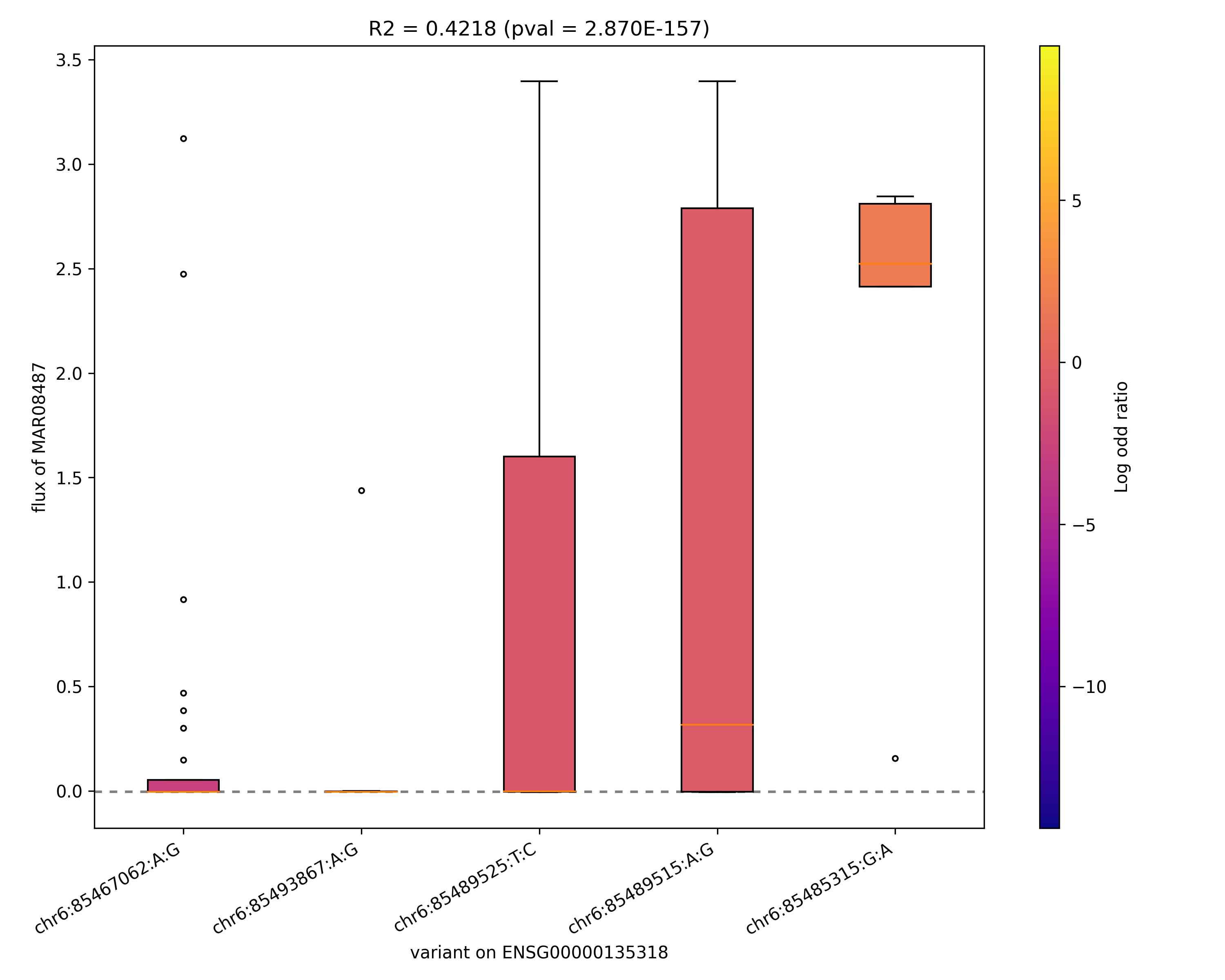

### MAR08586_ENSG00000059804.png

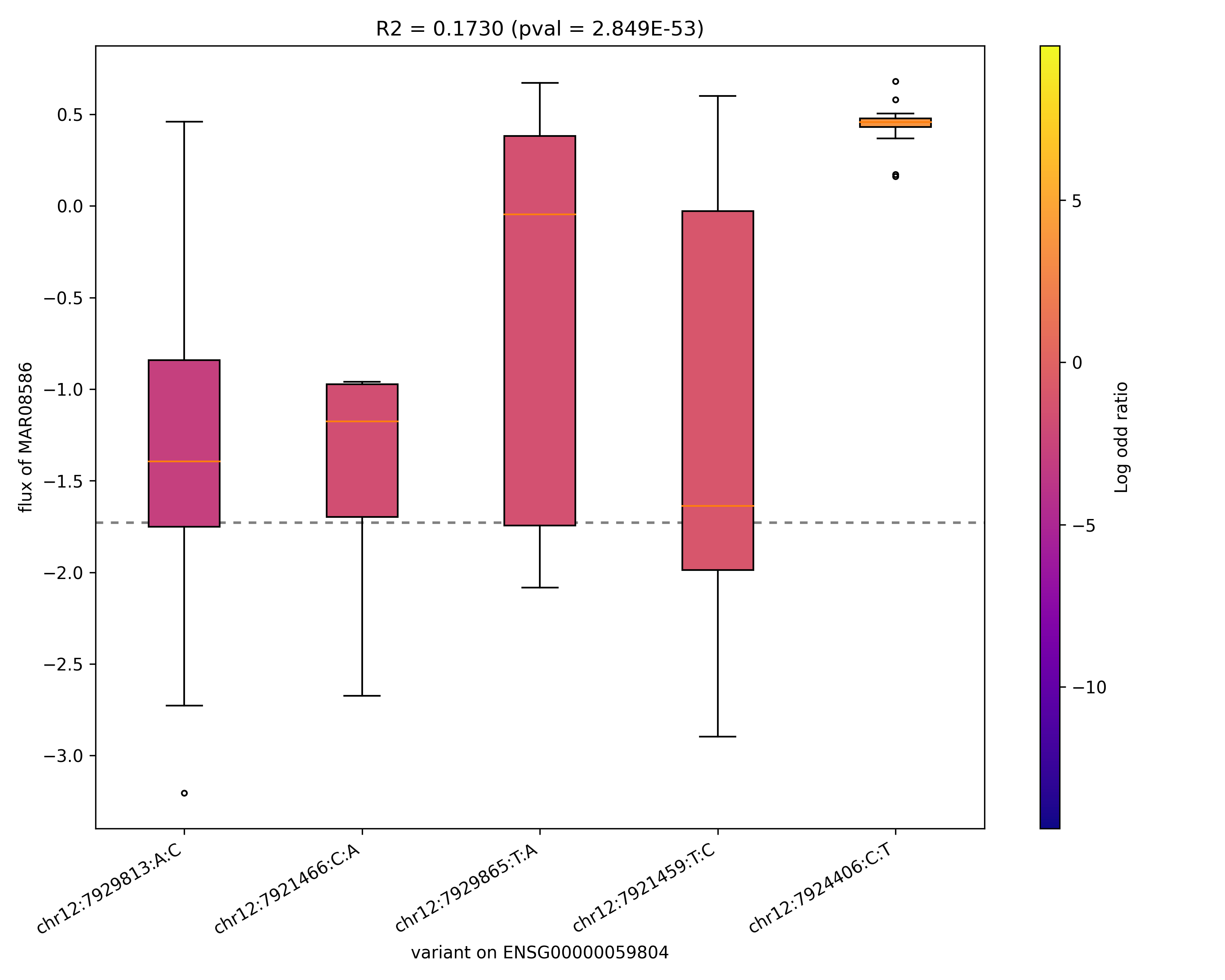

### MAR08587_ENSG00000214013.png

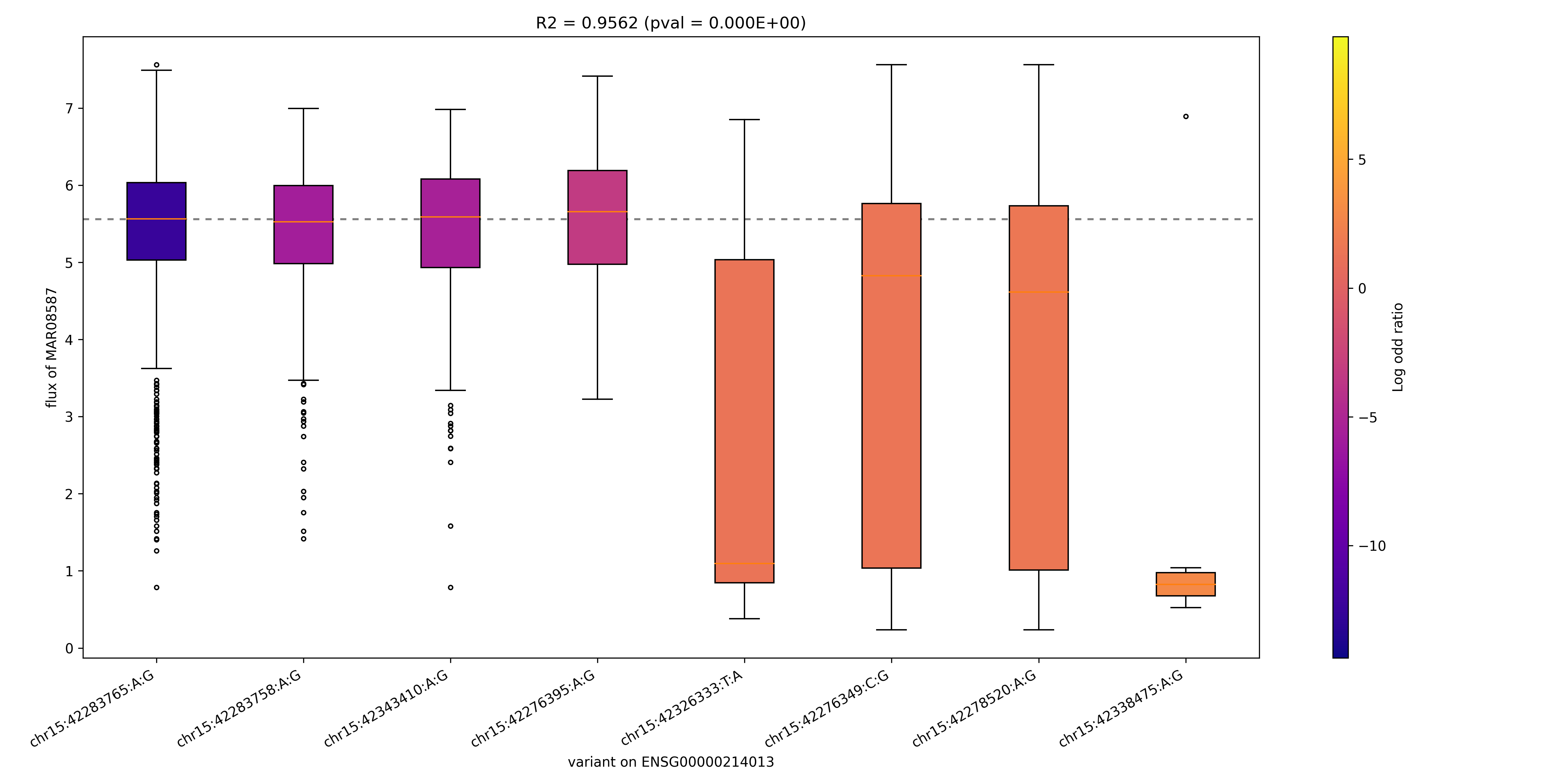

### MAR08591_ENSG00000214013.png

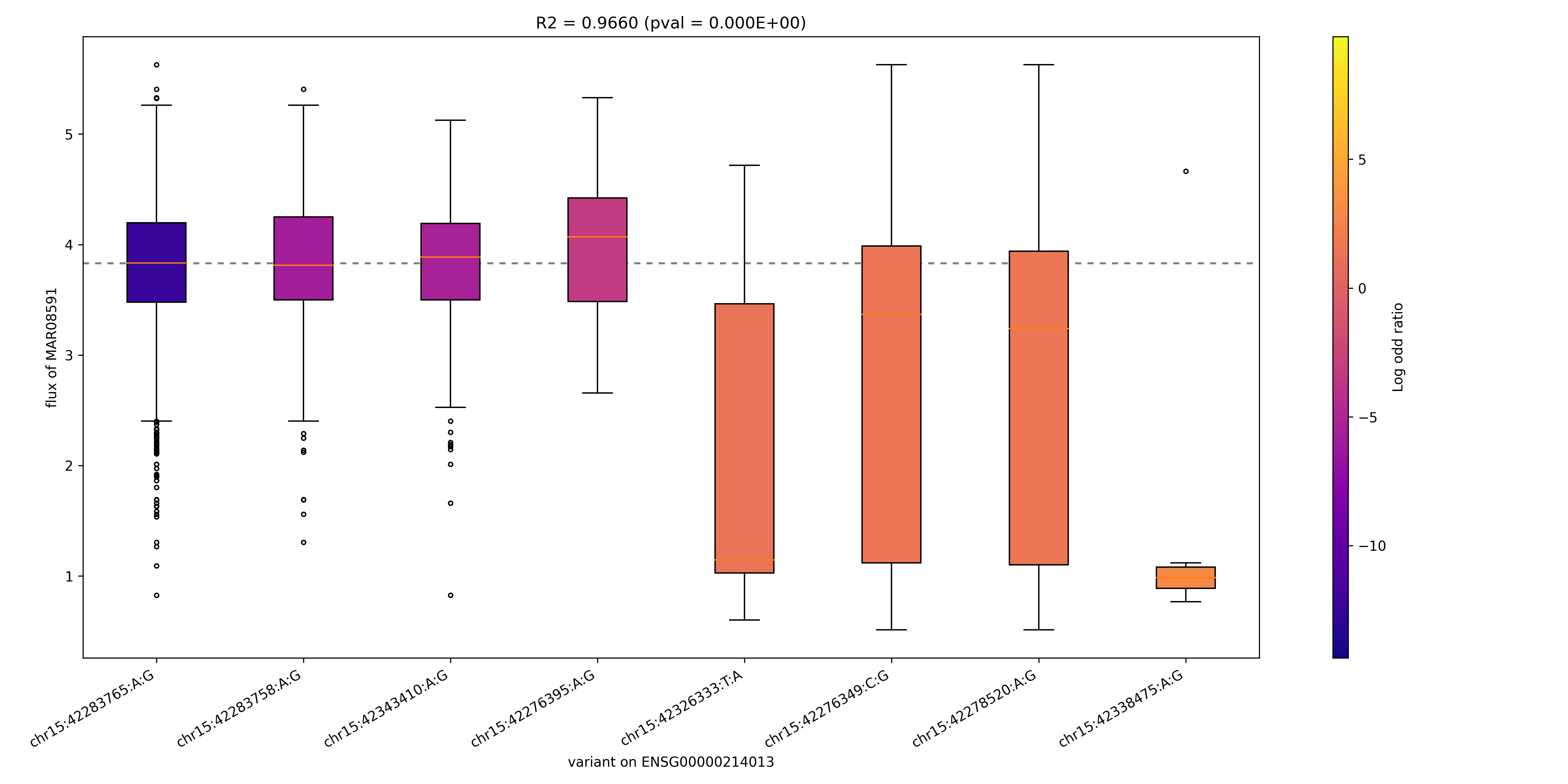
